# Assessing the Replicability of Scientific Claims in Drosophila Immunity Through an Analysis of 400 Publications

**DOI:** 10.1101/2025.07.07.663442

**Authors:** Hannah Westlake, Fabrice David, Yao Tian, Kenan Krakovic, Prince Kumar Sah, Nathan Klotz, Asya Dolgikh, Liza Juravlev, Thomas Esmangart de Bournonville, Alexia Carboni, Claudia Melcarne, Tisheng Shan, Yang Wang, Yizhu Mu, Akshata Kotwal, Nadia Pirko, Jean Philippe Boquete, Fanny Schüpfer, Samuel Rommelaere, Mickael Poidevin, Zhonggeng Liu, Shu Kondo, Girish S. Ratnaparkhi, Sveta Chakrabarti, Guiqing Liu, Florent Masson, Li Xiaoxue, Mark A. Hanson, Haobo Jiang, Francesca Di Cara, Estee Kurant, Bruno Lemaitre

**Affiliations:** Global Health Institute, School of life science, EPFL, Lausanne, Switzerland; BioInformatics Competence Center of EPFL-UNIL, School of Life Sciences, Station 19, EPFL, 1015 Lausanne, Switzerland; Department of Human Biology, Faculty of Natural Sciences, University of Haifa, Haifa 34988, Israel; Department of Entomology and Plant Pathology, Oklahoma State University, Stillwater, OK, 74078, USA; Dalhousie University, Department of Microbiology and Immunology, Halifax, NS, Canada; Indian Institute of Science Education and Research (IISER), Dr. Homi Bhabha Road, Pashan, Pune, India; Department of General and Medical Genetics, The Institute of Biology and Medicine, Taras Shevchenko National University of Kyiv, 64 Volodymyrska St., 01601 Kyiv, Ukraine; Institute for Integrative Biology of the Cell (I2BC), INSERM U1280, CEA, CNRS, Université Paris-Saclay, Gif-sur-Yvette, France; College of Plant Science and Technology, Huazhong Agricultural University, Wuhan, Hubei Province, China; Department of Biological Science and Technology, Tokyo University of Science, Research Building 11F Tokyo University of Science 6-3-1, Niijuku, Katsushika-ku, Tokyo 125-8585, Japan; Manipal Institute of Regenerative Medicine, Bengaluru, Manipal Academy of Higher Education, Manipal, India; Institute of Zoology, Guangdong Academy of Sciences, Guangzhou, China; Micropolis, 268 Route de Micropolis, Saint-Léons, France; Centre for Ecology and Conservation, University of Exeter, Penryn Campus, Penryn, Cornwall, TR10 9FE, UK

## Abstract

*Drosophila* immunity has been the focus of intense study and has impacted other research fields including innate immunity and agriculturally or epidemiologically relevant investigations of insect pests and vectors. We have performed a reproducibility project that analyses the conceptual replicability of claims from articles published on *Drosophila* immunity before 2011. To assess replicability, we extracted claims from 400 articles on the *Drosophila* immune response to bacteria and fungi and performed preliminary verification by comparing these claims to other published literature in the field. Using alternative approaches, we also experimentally tested some ‘unchallenged’ claims, with no published follow-up. The intent of this analysis was to centralize evidence on insights and findings to improve clarity for scientists that may base research programs on these data. All our data are published on a publicly available website associated with this article (https://ReproSci.epfl.ch/) that encourages community participation. This article provides a short summary of claims that were found to have contradictory evidence, which may help the community to assess past findings on *Drosophila* immunity and improve clarity going forward. Statistical analysis of this reproducibility study and metascience insights obtained from this approach are discussed in a companion article.

## Introduction

Innate immune mechanisms have become central to our understanding of immunology (Danilova, 2006; Medzhitov and Iwasaki, 2024; Pradeu et al., 2024), although their study was long overshadowed by interest in adaptive immunity. Innate immune mechanisms are now widely recognized as having key importance as a first layer of defense, in the recognition of infectious agents, and in subsequent polarization of the immune response. These responses are activated by a specific set of receptors that sense microbe-specific molecules, virulence factors and/or infection-induced damage. Complex signaling pathways integrate this information to induce specific innate immune modules adapted to the characteristics of the encountered pathogen. Characterization of the innate immune response has greatly benefitted from the use of model organisms such as *Drosophila,* which is amenable to genetic studies and relies exclusively on innate immunity for defense (Buchmann, 2014; Westlake et al., 2024).

The *Drosophila* immune system is composed of several immune modules, some of which are conserved in vertebrates (Buchon et al., 2014; Liegeois and Ferrandon, 2022; Westlake et al., 2024). The systemic immune response is best characterized by the production and secretion into the hemolymph of host-defense peptides by the fat body and hemocytes. The best-characterized of these effectors are the antimicrobial peptides (AMPs). AMPs are small cationic peptides with antimicrobial activity, whose expression is induced to incredibly high levels in response to infection (Hanson and Lemaitre, 2020). The systemic antimicrobial response is predominantly regulated at the transcriptional level by the Toll and Imd (Immune-deficiency) signaling pathways. In *Drosophila*, the Toll pathway is activated by microbial cell wall components (fungal glucans and Lysine-type peptidoglycan), and microbial proteases. Toll signaling culminates in the activation of two NF-κB transcription factors, Dif (Dorsal-related immunity factor) and Dorsal, which regulate a large set of immune genes such as the Bomanins, the antifungal peptide Drosomycin, and many other proteins (Clemmons et al., 2015; De Gregorio et al., 2002; Troha et al., 2018). Flies lacking Toll pathway activity are viable but highly susceptible to infection by Gram-positive bacteria and fungi (Rutschmann et al., 2002). The Imd pathway is activated by DAP-type peptidoglycans produced by Gram-negative bacteria and a subset of Gram-positive bacteria (*e.g. Bacillus*) (Kaneko et al., 2004; Leulier et al., 2003). Binding of peptidoglycan to receptors of the Peptidoglycan Recognition Protein family (PGRP-LC, PGRP-LE, PGRP-SD) initiates an intracellular signaling cascade with homology to the TNFα-Receptor pathway, which ultimately activates the NF-κB factor Relish (Choe et al., 2002; Gottar et al., 2002; Iatsenko et al., 2016; Rämet et al., 2002; Takehana et al., 2004). The Imd pathway regulates the expression of many genes encoding antibacterial peptides, serine proteases, and the iron sequestration protein Transferrin, among others. Imd deficient flies are viable, but susceptible to Gram-negative bacterial infection.

Complementary to these transcriptional programs, melanization is an arthropod-specific immune response resulting in rapid deposition of the black pigment melanin at wound or infection sites and concomitant production of microbicidal reactive oxygen species (Cerenius et al., 2008). Melanization relies on the activation of enzymes called prophenoloxidases (PPOs), which catalyze the oxidation of phenols, resulting in melanin polymerization (Dudzic et al., 2015). This process also hardens clots with melanin polymer plugs that prevent blood loss, akin to the mechanical function of fibrin scabs. Clotting, which involves both humoral factors and hemocytes, contributes to wound healing and has been implicated in host defense against nematodes (Theopold et al., 2014; Yadav and Eleftherianos, 2018).

In addition to soluble immune effectors, cellular components also play a crucial role in immunity. In adult flies, two hemocyte types known as plasmatocytes and crystal cells are present in circulation or as sessile cells attached to the inner cuticle (Banerjee et al., 2019; Lanot et al., 2000; Sanchez Bosch et al., 2019). Plasmatocytes are macrophage-like cells with phagocytic activity, while crystal cells produce and store PPO enzymes and contribute to melanization. Use of hemocyte deficient flies has shown that hemocytes contribute to survival upon systemic infection to certain bacterial species, and several phagocytic receptors have been identified (Charroux and Royet, 2009; Defaye et al., 2009; Melcarne et al., 2019a; Stephenson et al., 2022; Ulvila et al., 2011).

Natural infection with microbes also triggers a local immune response in epithelia, which has been extensively characterized in the gut (Buchon et al., 2013; Tafesh-Edwards and Eleftherianos, 2023; Westlake et al., 2024). In this organ, a range of defenses including the peritrophic matrix, an acid pH, peristalsis, and epithelial repair through stem cell proliferation all contribute to tolerating ingested pathogens. Local production of AMPs and ROS provide two complementary inducible defense mechanisms in the gut (Liehl et al., 2006; Ryu et al., 2006). It is now clear that each organ contributes individual input to the immune response in an integrated manner, which vary according to the route of infection and the nature of the pathogens (Westlake et al., 2024). Finally, extensive studies have highlighted the role of the immune system upon mating and in the reproductive organs to prevent infection (Schwenke et al., 2016; Siva-Jothy, 2009).

*Drosophila* immunity has been a topic of intense research, with thousands articles and reviews published on this topic (Westlake et al., 2024). Characterization of the *Drosophila* immune system has contributed significantly to fields including innate immunity and agriculturally or epidemiologically relevant investigations of insect pests and vectors. As expected in such a large body of work, some published results were later found to be irreplicable. Although some of these results have been contradicted in the literature, many have no published follow-up, either due to a lack of research or low motivation to publish negative or contradictory results. We have addressed this by performing an extensive reproducibility project that analyses the conceptual replicability of claims from articles on *Drosophila* immunity accepted for publication before 2011. This time frame allows opportunity for publication of follow-up results. We extracted major and minor claims from 400 articles on the *Drosophila* immune response to bacteria and fungi and performed verification where possible by comparing to other published literature in the field. We also experimentally tested 55 ‘unchallenged’ claims with no published follow-up, a process that will be continuously expanded with the help of the community.

The aim of the ReproSci project is to assess the replicability of claims made in articles published in the field of *Drosophila* immunity. This is in no way an assessment of the ‘scientific value’ of the research. Although a lack of replicability is often viewed negatively, papers that include non-replicable observations may nevertheless have a positive impact by developing new techniques, introducing interesting concepts, providing a foundation for future discoveries, or presenting results that are useful despite incorrect interpretation. Similarly, articles that include only well-supported claims may be of low scientific value if they only confirm previous observations and present little in the way of innovation. The leading authors (H.W., B.L.) acknowledge that the selection of claims within papers and assessments of those claims cannot be fully objective (see limitation section). Although conflicting data are often discussed at conferences or at less formal meetings, diffusion of this knowledge is limited due to lack of publications. Our hope is to provide a dataset with community-sourced proofs that allow a wider range of scientists to easily identify claims with contradictory evidence. In association with this article, we have published the *ReproSci* website, which describes and presents assessments of claims from 400 articles and encourages community participation (https://ReproSci.epfl.ch/). Opening the ReproSci database in July 2023 to comment by community members was intended to provide a greater degree of objectivity and consensus and generate ongoing discussion beyond initial assessments. Although our study has revealed that most claims published in the field of *Drosophila* immunity before 2011 are conceptually replicable, this article synthesizes the main results on a subset of claims of particular interest that may be considered as challenged or as more complex than initially reported. Although we cannot entirely rule out the possibility that our verification experiments contain errors, our findings call for caution regarding the replicability of certain claims in the field of *Drosophila* immunity. These include claims published in high profile journals that have been contradicted or have tenuous support. This article is accompanied by 28 supplementary materials that include new experimental data from various laboratories that support or contradict some unresolved claims that are discussed in this article. In many cases, re-analysis of these claims is made possible by advances in understanding of immune mechanisms and technology since publication of the initial articles. The website associated with this article provides a more complete analysis. Statistical analysis of this reproducibility study and metascience insights obtained from this approach are discussed in a companion article (Lemaitre et al., 2026).

## Results

### Overview of the reproducibility project

400 research articles focusing on *Drosophila* immunity published between 1959 and 2011 were selected by PubMed keyword retrieval and manual curation (**Supplementary Table 1**). Articles dealing with population genetics, virulence of entomopathogens, gut homeostasis, and the immune response to viruses and wasps were excluded. The main (1 per article), major (2-4 per article) and minor (0-4 per article) claims were extracted along with a summary of evidence supporting the claims and the methods of each article. The replicability of each claim was then assessed by analyzing subsequent publications on the topic from the same or other laboratories. Claims were assessed as verified, unchallenged, challenged, mixed, or partially verified (see Methods). All article summaries and assessments were released to the community on July 11^th^, 2023, via the *ReproSci* website, which allows users to comment on the claims and provide confirmatory or contradictory evidence. 45 major and 11 minor claims (many of which are discussed in the Supplement) were also experimentally tested in various laboratories using alternate approaches. For instance, a result initially obtained using RNAi may be tested using newly generated CRISPR mutants. Our project focuses on assessing the strength of the claims themselves (conceptual replication/inferential/indirect reproducibility) rather than testing whether the original methods produce repeatable results (results/direct reproducibility). Thus, our conclusions do not directly challenge the initial results leading to a claim, but rather the general applicability of the claim itself. This article presents some of the main observations from the *ReproSci* project and is intended to allow the reader to quickly grasp key claims in the *Drosophila* immunity field with low replicability (challenged claims) or not as simple as proposed in the initial articles.

**Figure 1.**
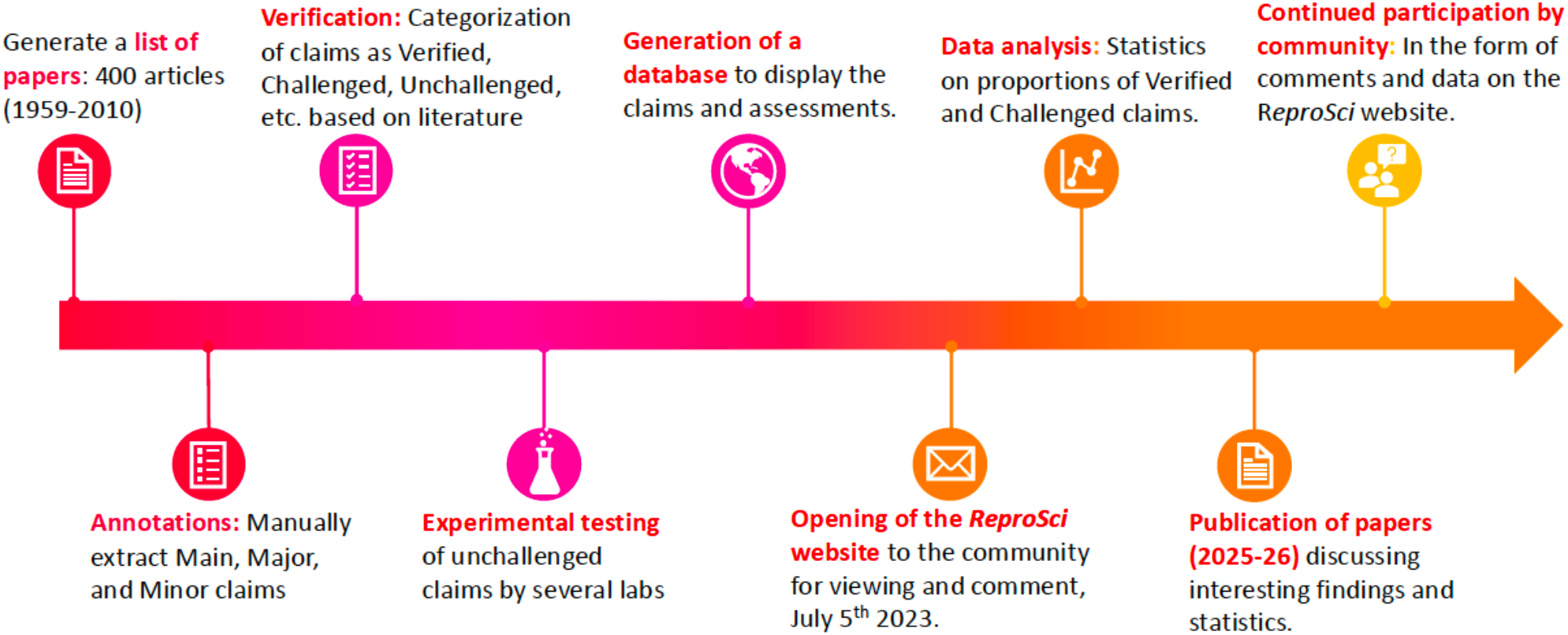
The timeline of the *Drosophila* immunity *ReproSci* project.

### Insights from the *ReproSci* project on the Toll pathway

#### Toll is not regulated by 18-wheeler

Following the characterization of the Toll receptor (Toll-1) as a key regulator of *Drosophila* AMP genes (Lemaitre et al., 1996), the Toll homolog 18-wheeler (Toll-2) was reported to have a complementary role in LPS sensing and regulation of the antibacterial response, similar to mammalian TLR4 (Williams et al., 1997). This paper was initially widely cited, but was directly challenged by a follow-up study revealing that 18-wheeler does not have a direct role in immunity, but rather indirectly affects survival and AMP gene expression through a general negative impact on fly development (Ligoxygakis et al., 2002).

#### GNBP1 is not a sensor of fungal glucan

GNBP1 (Gram-Negative Binding Protein 1) was initially characterized for its ability to bind LPS and fungal glucan, based on overexpression in cell culture and *in vitro* binding studies (Kim et al., 2000). Subsequent genetic studies revealed that this secreted protein instead cooperates with PGRP-SA in sensing of Gram-positive bacteria, and found no evidence that GNBP1 acts as an immune sensor for LPS or glucan (Gobert et al., 2003; Pili-Floury et al., 2004). GNBP3, a *Drosophila* homolog of GNBP1, does have glucan binding activity and activates immune responses to fungi *in vivo* (Gottar et al., 2006).

#### GNBP1 does not process peptidoglycan

Gram-positive bacteria stimulate the Toll pathway through cooperative action of PGRP-SA and GNBP1, which activate the apical serine protease ModSP upon sensing of peptidoglycans (Buchon et al., 2009c; Gobert et al., 2003; Pili-Floury et al., 2004). Structural studies revealed that PGRP-SA can bind peptidoglycan, consistent with its demonstrated role as pattern-recognition receptor *in vivo* (Chang et al., 2004). However, one study suggested that GNBP1 can also digest peptidoglycan, and that this hydrolytic function of GNBP1 is required for full activation of PGRP-SA and ModSP (Wang et al., 2006). Reported enzymatic activity of GNBP1 has never received follow up. Studies in other insects such as *Tenebrio molitor* and *Manduca sexta* suggest that the primary function of GNBP1 is as an adaptor between PGRP-SA and the apical serine protease ModSP (Kim et al., 2008; Park et al., 2007; Wang et al., 2022). *In vitro* experiments performed as part of *ReproSci* found that GNBP1 does not process peptidoglycan in the absence or presence of PGRP-SA (**Supplementary S1**). Similarly, pre-incubation of GNBP1 with peptidoglycan did not enhance ModSP autoactivation. These results suggest that contrary to the initial claim, GNBP1 has no enzymatic activity against peptidoglycan. The initial results may have been obtained due to residual impurities in preparations of recombinant GNBP1.

#### Spirit, Sphinx and Spheroide are not key components of the Toll pathway

An RNAi screen identified five serine proteases involved in the activation of the Toll ligand Spatzle during the immune response (Kambris et al., 2006a). The roles of two of these genes, *SPE* and *grass*, were confirmed in subsequent studies, although *grass* was shown to affect the whole pattern recognition branch of the Toll pathway and not only the PGRP-SA/GNBP1 branch as suggested in the initial study (El Chamy et al., 2008; Jang et al., 2006). Data from *ReproSci* using newly generated viable loss-of-function mutations reveals that of the remaining three genes, neither the serine protease homologs Spheroide and Sphinx1/2 nor the serine protease Spirit are strictly required for Toll activation (**Supplementaries S2, S3, S4**). It is likely that Sphinx1/2 has no role in the immune response, while the immune-inducible gene Spirit may have a redundant or a subtle role. Non-replicable results on the roles of Spirit, Sphinx and Spheroide in Toll pathway activation may be due to off-target effects common to first-generation RNAi tools (Ueda, 2001).

#### The Toll immune pathway is not negatively regulated by wntD

Gordon et al., (2005) found that *wntD* acts as a feedback inhibitor of *dorsal* but not *Dif* during the innate immune response. *wntD* mutants had elevated expression of *Diptericin* but not *Drosomycin* in response to septic infection with the Gram-positive bacterium *Micrococcus luteus*, and increased susceptibility to *Listeria monocytogenes* that was rescued in *wntD, dorsal* double mutants. Data from FlyAtlas (Thurmond et al., 2019) show strong expression of *wntD* in the early embryo but not at the adult stage at which the experiments were performed by (Gordon et al., 2005). Using an independent null mutant of *wntD*, we did not observe any impact of *wntD* on expression of *Diptericin* and *Drosomycin,* or increased sensitivity to *Listeria monocytogenes* infection (**Supplementary S5**). Thus, we conclude that *wntD* is likely not a negative regulator of the Toll pathway during the immune response. The initial claim concerning *wntD* may be explained by a genetic background effect independent of *wntD*.

#### Toll is not strongly activated by overexpression of GNBP3 or Grass

Experiments performed for *ReproSci* find that contrary to previous reports, overexpression of *GNBP3* (Gottar et al., 2006) or *Grass* (El Chamy et al., 2008) in the absence of immune challenge does not effectively activate Toll signaling or only at a very modest level (**Supplementaries S6, S7**).

### Insights from the reproducibility project on the Imd pathway

#### PGRP-SD is a component of Imd and not Toll signalling

PGRP-SD was initially identified as a secreted pattern-recognition receptor involved in the Toll pathway in response to certain Gram-positive bacteria (Bischoff et al., 2004; Wang et al., 2008). This was inconsistent with subsequent observations that **i)** PGRP-SD binds DAP-type peptidoglycan found in Gram-negative bacteria which typically activate Imd signaling (Basbous et al., 2011; Leone et al., 2008; Lim et al., 2006); and that **ii)** this gene is regulated by the Imd pathway (De Gregorio et al., 2002). Using isogenic fly stocks and newly generated mutants, Iatsenko et al., subsequently showed that PGRP-SD is a positive regulator of the Imd pathway upstream of PGRP-LC that promotes sensing of DAP-peptidoglycan (Iatsenko et al., 2016). Additional *in vitro* experiments performed in the framework of *ReproSci* reveal that PGRP-SD does not contribute to Toll activation by peptidoglycan under our experimental conditions (**Supplementary S1**). This suggests that the original results assigning PGRP-SD to the Toll pathway may be due to a genetic background effect.

#### PGRP-LE is exclusively intracellular and is not critical for cuticular melanization

(Takehana et al., 2004, 2002) found that intracellular PGRP-LE contributes to the activation of the Imd pathway together with the transmembrane receptor PGRP-LC. While its role as an intracellular sensor has been confirmed (Bosco-Drayon et al., 2012; Kaneko et al., 2006; Neyen et al., 2012), the initial papers claimed that an extracellular form of PGRP-LE could function akin to CD14 (Takehana et al., 2004, 2002). Follow-up papers have not addressed the existence of this extracellular form. Using a new proteomic approach, we did not detect PGRP-LE in adult fly hemolymph (Rommelaere et al., 2024). Moreover, use of a GFP tagged version of PGRP-LE (Bosco-Drayon et al., 2012) did not reveal an extracellular form in larvae (**Supplementary S8**). We conclude that it is likely PGRP-LE is an exclusively intracellular sensor, with a prominent role regulating the gut immune response as reported by (Bosco-Drayon et al., 2012; Neyen et al., 2012). In addition, experiments performed for *ReproSci* found no critical role for PGRP-LE in cuticular melanization, contrary to (Takehana et al., 2004, 2002) although we cannot exclude subtler role of PGRP-LE in hemolymph melanization (**Supplement S9**). The results of the initial papers are likely due to the use of ubiquitous overexpression of *PGRP-LE*, resulting in melanization due to overactivation of the Imd pathway and resulting tissue damage. We were also unable to find a role for PGRP-LE in survival to *Bacillus* infections (**Supplementary S10**) and conclude that the initial result may be due to genetic background and comparison to a robust wild-type as the baseline.

#### LPS does not activate the Imd pathway

In mammals, LPS is a strong inducer of TLR4-NF-κB pathway (Hoshino et al., 1999; Poltorak et al., 1998). Studies in *Drosophila* using LPS from Sigma^TM^ initially provided evidence that LPS can stimulate the Imd pathway (Imler et al., 2000). However, use of highly purified bacterial cell wall components showed that DAP-type peptidoglycan and not LPS stimulates Imd pathway activity (Kaneko et al., 2004; Leulier et al., 2003). Furthermore, in contrast to an initial claim by Leulier et al., 2003, monomeric DAP-type peptidoglycan (notably tracheal cytotoxin, TCT) is also a strong elicitor of the Imd pathway, in addition to polymeric peptidoglycan (Kaneko et al., 2004; Stenbak et al., 2004).

#### Naked DNA is not immunogenic in Drosophila

In mammals, free DNA can activate IFN-mediated inflammatory responses through sensors such as cGAS (Decout et al., 2021). In 2002, an article reported that DNA is immunogenic in *Drosophila* after observing elevated expression of *Diptericin* genes in *DNase II^lo^* hypomorphic mutant flies (Mukae et al., 2002). However, experiments performed for *ReproSci* using the original *DNAse II^lo^* hypomorph show that elevated *Diptericin* expression in the hypomorph is eliminated by outcrossing of chromosome II, and does not occur in an independent *DNAse II* null mutant, indicating that this effect was likely due to genetic background **(Supplementary S11)**. Furthermore, elevated *Diptericin* expression in *DNAse II^lo^* flies is not observed in germ-free conditions, indicating that it is not due to DNA excess (West, 2018). Moreover, experiments showed that injection of naked bacterial and eukaryotic DNA does not activate an immune response in wild-type or *DNAse II* mutant flies. These results indicate that naked DNA unlikely to be an elicitor for AMP genes in *Drosophila* as originally claimed, although it may activate the recently discovered cGLR-STING pathway (Cai et al., 2022). Subsequent studies indicated that cGLR ligands are RNA molecules, not DNA (Holleufer et al., 2021; Slavik et al., 2021). Thus, it remains possible that such receptors may be able to detect native rather than naked DNA. The possibility that DNA-associated proteins such as histones are immunogenic also remains to be investigated.

#### JNK signaling is not directly required for AMP expression

Several studies pointed to a direct role of the JNK pathway in the regulation of AMPs (Delaney et al., 2006; Kallio et al., 2005; Kleino et al., 2005) while other studies did not identify any major role (Boutros et al., 2002; Kenmoku et al., 2017; Park et al., 2004; Silverman et al., 2003). Experiments performed for *<u>ReproSci</u>*show that inhibition of JNK (RNAi against *hep* or overexpression of *bsk^DN^*) in the adult fat body does not significantly suppress Diptericin expression in response to infection with the Gram-negative bacterium *Ecc15* (**Supplementary S12**). This contradicts conclusions of (Delaney et al., 2006; Kallio et al., 2005; Kleino et al., 2005) and supports those of (Boutros et al., 2002; Park et al., 2004; Silverman et al., 2003). This is in agreement with the more recent *in vivo* results of (Kenmoku et al., 2017). In the absence of further evidence, we conclude that JNK is unlikely to have a strong direct or systematic role on regulation of *Drosophila* AMPs *in vivo,* and is not required in the adult fat body for AMP expression in response to infection. Previous results finding a direct role for JNK in AMP production are likely due to a combination of i) use of cell culture systems that do not fully reflect *in vivo* systems; ii) the conclusion that kinase-dead JNK kinase (TAK1) is biologically equivalent to a null mutant; and iii) underappreciation for the overarching role that JNK plays in cytoskeletal remodeling, pathway activation, and cell differentiation, which are not specific to immunity.

#### Caspar is not a canonical negative regulator of Imd signaling

*caspar* encodes an orthologue of human Fas associated factor 1 (FAF1) which was reported to interact with key immune proteins including NF-κB. Kim et al., (2006) found that a null mutation of *caspar* produced high constitutive expression of *Diptericin* and increased resistance to Gram-negative infections (Kim et al., 2006). Using flies deficient for *caspar*, we did not detect any increased *Diptericin* expression in unchallenged flies as initially reported (**Supplementary S13**). We conclude that while Caspar might affect the Imd pathway in certain tissue-specific contexts, it is unlikely to act as a generic negative regulator of the Imd pathway.

### Insights from the reproducibility project on immune-induced peptides

#### Edin does not significantly contribute to lifespan or resistance against L. monocytogenes

Gordon et al., (2008) identified *edin,* a host defense peptide gene highly expressed upon *L. monocytogenes* infection. They found that overexpression of *edin* markedly reduced lifespan and fly fitness whereas RNAi knockdown of *edin* in the fat body increased susceptibility to *L. monocytogenes* infection. Contrary to (Gordon et al., 2008), experiments performed for *ReproSci* using a novel *edin* null mutant indicate that loss of *edin* does not increase susceptibility to *L. monocytogenes* infection compared to appropriate controls. Moreover, overexpression of *edin* did not cause any major reduction in lifespan (**Supplementary S14**).

#### Listericin overexpression is not protective against L. monocytogenes infection

(Goto et al., 2010) found using transfection and RNAi in S2 cells that the *Listericin* gene encodes a novel antibacterial peptide-like protein whose induction is cooperatively regulated by PGRP-LE and the JAK-STAT pathway. However, a microarray indicates that this gene is regulated by the Imd pathway with a minor impact of Toll (De Gregorio et al., 2002). Experiments performed for *ReproSci* suggest that *Listericin* expression does not specifically depend on PGRP-LE but rather is induced through both PGRP-LC and –LE as other Imd-regulated AMPs are (**Supplementary S15**). Silencing STAT92A in the fat body by using an *in vivo* RNAi approach did not confirm a role of JAK/STAT in induction of *Listericin* following infection as claimed by (Goto et al., 2010). Overexpression of Listericin using a newly generated construct did not reveal any protective effect against *L. monocytogenes* as found by (Goto et al., 2010). Thus, Listericin should rather be considered as a host defense peptide regulated by the Imd pathway whose activity remains to be confirmed.

### Insights from the reproducibility project on phagocytosis and encapsulation

#### Dscam1 does not directly contribute to phagocytosis through opsonization

*Drosophila Dscam1* has three immunoglobulin-like domains and encodes thousands of isoforms through alternate splicing. An early paper proposed that secreted Dscam1 isoforms can bind bacteria and enhance phagocytosis, an attractive hypothesis due to its potential to produce multiple ‘recognition’ isoforms akin to an adaptive immune system (Watson et al., 2005). Dscam1 homologs have independently been implicated in crustacean and mosquito immunity, systems that are not easily amenable to genetic studies (Dong et al., 2025, 2006; Ng et al., 2014; Ng and Kurtz, 2020). However, no follow-up studies were published on the role of Dscam1 proteins in *Drosophila* immunity. Subsequent studies showed that *Dscam1* splicing is not affected by septic injury (Armitage et al., 2014), and contrary to a role for secreted Dscam1 isoforms in *Drosophila* opsonization, a proteomic analysis of adult hemolymph did not detect any secreted Dscam1 isoforms (Rommelaere et al., 2024). Furthermore, all Dscam1 isoforms encode the transmembrane domain. Experiments performed for *ReproSci* did reveal a mild decrease in phagocytosis of *E. coli* and *S. aureus* by hemocytes of *hml>Dscam1-RNAi* larvae: a 30% reduction in comparison to 50% in the initial study (Watson et al., 2005) (**Supplementary S16**). Dscam1 plays a key role in neuronal dendrite arborization and affects how neurons influence peripheral hematopoiesis, and RNAi against *Dscam1* in neurons reduces hemocyte numbers (Ouyang et al., 2020). It is likely that reduced phagocytosis in *Dscam1* depleted animals is due to an effect on hematopoiesis rather than a direct immune effect. In the absence of additional studies, *Dscam1* should not be considered a *Drosophila* immune gene *sensu stricto*.

#### PGRP-LC is not a phagocytic receptor

(Rämet et al., 2002) reported a role of the Imd receptor PGRP-LC in phagocytosis of Gram-negative bacteria in an extensive RNAi screen (Rämet et al., 2002). Subsequent studies did not reveal any critical role of PGRP-LC in bacterial engulfment (Choe et al., 2002; Melcarne et al., 2019). It is likely that PGRP-LC indirectly (and modestly) affects phagocytosis through its role in the Imd pathway and associated upregulation of phagocytic receptors (Wong et al., 2017), but this pattern recognition receptor should not be considered a phagocytic receptor.

#### Croquemort is not a phagocytic engulfment receptor for apoptotic cells

The CD36 scavenger member Croquemort was initially identified as a macrophage-specific phagocytic receptor for apoptotic cells. Use of an overlapping set of deficiencies removing *croquemort* revealed that Croquemort was critical for engulfment of apoptotic cells during embryogenesis (Franc, 1999; Franc et al., 1996). However, using a clean *croquemort* null mutant, experiments performed for *ReproSci* reveal that *croquemort* is not required for removal of apoptotic cells in embryos (**Supplementary S17**). *croquemort* mutant phagocytes do display a general defect in phagosome maturation and accumulate acidic vacuoles (similar to *draper* mutants) (Guillou et al., 2016; Han et al., 2014). It is not known why *croquemort* affects phagosome maturation, but could be due to its role in lipid uptake (Woodcock et al., 2015). The available data indicate that Croquemort is likely not a key phagocytic receptor required for initial uptake but contributes to sustaining phagocytosis by promoting lysosomal degradation. Of note, another CD36 member, Santa Maria, was recently implicated in the phagocytosis of apoptotic cells by glia (Hilu-Dadia et al., 2025).

#### dSR-CI contributes to hemocyte binding to bacteria but is not required for phagocytosis

The scavenger receptor dSR-CI was initially identified as a hemocyte-restricted receptor with high affinity for low density lipoprotein (Pearson et al., 1995). *Ex vivo* and RNAi experiments suggested a significant role for dSR-CI as a pattern-recognition receptor (Rämet et al., 2001). Experiments for *ReproSci* using a newly generated *dSR-CI* null mutant and a *dSR-CI*, *dSR-CIII* double mutant confirmed that *SR-CI* promotes binding to *E. coli* and *S. aureus* bacteria as initially proposed by (Rämet et al., 2001) (**Supplementary S18**). Although this was not assessed in the initial study, SR-CI is often cited as a phagocytic receptor, but our experiments did not reveal any major requirement of dSR-CI in phagocytosis of *E. coli* and *S. aureus ex vivo*. We confirm the (Rämet et al., 2001) observation that loss of *SR-CI* has no apparent effect on the inducible AMP response. The precise role of *dSR-CI* in the *Drosophila* immune response remains to be determined.

#### Retinophilin/Undertaker has no major role in phagocytosis

Using overlapping deficiencies in embryos and RNAi in S2 cells, (Cuttell et al., 2008) found that Undertaker (uta/rtp), a Junctophilin, was required for phagocytosis of apoptotic cells and bacteria and links Draper-mediated phagocytosis to Ca^2+^ homeostasis. However, expression of Undertaker is restricted to the adult eye (Thurmond et al., 2019), which argues against a major role in phagocytosis by embryonic macrophages. Using a *rtp^1^* null allele, we did not observe any effect of Undertaker on phagocytosis of apoptotic cells or bacteria by larval hemocytes (**Supplementary S19**). The published data are most consistent with a scenario in which RNAi generated off-target knockdown of a protein related to *retinophilin/undertaker,* while Undertaker itself is unlikely to have a role in phagocytosis. In contrast, independent data support the notion put forth by (Cuttell et al., 2008) that calcium flux and store-operated calcium entry (including all other components mentioned in this paper) are required for phagocytosis and blood cell activation in animals (Etchegaray et al., 2012; Gronski et al., 2009; Moon et al., 2020; Mortimer et al., 2013; Weavers et al., 2016).

#### NimC1 has a co-operative role in phagocytosis of Gram-negative bacteria

An *in vivo* RNAi approach suggested that NimC1 is required for the uptake of Gram-negative bacteria (Kurucz et al., 2007). In contrast, a null mutation in NimC1 did not reproduce this phenotype (Melcarne et al., 2019). However a *NimC1, eater* double mutant revealed that these two Nimrod receptors function co-operatively in phagocytosis of Gram-negative bacteria (Melcarne et al., 2019). It is possible this disparity relies on a cryptic effect, such as genetic background differences. Although the initial claim that NimC1 alone is essential for phagocytosis of Gram-negative bacteria was not replicable, the claim that NimC1 is involved in phagocytosis of Gram-negative bacteria has been validated.

#### Eater has a co-operative role in phagocytosis

An initial study pointed to a role for the transmembrane receptor Eater in the phagocytosis of both Gram-negative and Gram-positive bacteria (Chung and Kocks, 2011; Kocks et al., 2005). The authors used an overlapping set of deficiencies to remove Eater *in vivo* and RNAi in S2 cells to reach this conclusion. Subsequent studies have confirmed that Eater is indeed involved in the phagocytosis of Gram-positive bacteria (Bretscher et al., 2015; Chung and Kocks, 2011) While nearly all of the claims in the original article have been confirmed, subsequent studies showed that a null mutation of *eater* does not impact phagocytosis of Gram-negative bacteria (Bretscher et al., 2015; Claudia Melcarne et al., 2019). Reduced phagocytosis of Gram-negative bacteria was observed only in *NimC1, eater* double mutants, indicating that Eater is involved in this process but not strictly necessary.

#### Psidin is not required for induction of Defensin

Brennan et al., (2007) found that Psidin acts non-autonomously in the hemocytes to control induction of *Defensin* in the fat body, and suggested that a cytokine produced by hemocytes is required to activate systemic *Defensin* expression. In contrast to this, experiments performed for *ReproSci* using two independent *psidin* mutants show that *Defensin* expression is similar to wild type in these lines (**Supplementary S20**). Furthermore, this result, and other instances where mutations have been found to specifically eliminate Defensin expression, is likely due to segregating polymorphisms within Defensin that disrupt primer binding in some genetic backgrounds, leading to false negative results ((Brennan et al., 2007; Neyen et al., 2014) and see **Supplementary S20**). (Brennan et al., 2007) further found that Psidin is required for phagolysosomal maturation. Experiments performed for *ReproSci* confirm that phagolysosomal maturation is impaired in independent *psidin* mutants, and furthermore that *psidin* mutant hemocytes are abnormally large and multinucleate (**Supplementary S20**).

#### Hemese is not essential for encapsulation

(Kurucz et al., 2003) found that loss of Hemese, a hemocyte-specific transmembrane protein, causes overproduction of lamellocytes in permissive conditions. In contrast to this, using two fly lines, *Hemese^JP187^* and Hemese^JP828^, carrying a deletion in the central region of the gene, we find that loss of *Hemese* does not strongly affect lamellocyte number or encapsulation (**Supplementary S21**). The function of Hemese awaits further functional characterization.

### Insights from the reproducibility project on IRC, Duox, NOS and melanization

#### Immune Regulated Catalase (IRC) is not essential for survival to benign enteric infection

Using a ubiquitous and directed RNAi approach, (Ha et al., 2005b) found that adult flies depleted of the extracellular protein IRC (immune regulated catalase) displayed high mortality rates after ingestion of foods contaminated with normally benign microbes (Ha et al., 2005b). Contrary to this, we did not observe any increase in mortality upon feeding *IRC* null mutant flies with various microbial pathogens, although an effect of the mutation on motor function and overall fitness was observed (**Supplementary S22**). Moreover, IRC should not be considered as a bona fide catalase, but rather as a heme peroxidase, enzymes that uses hydrogen peroxide as a substrate to mediate some chemicals reactions such as protein cross-linking by forming dityrosine bonds (Kumar et al., 2010). The immune regulated gene *IRC* is likely to play a role in the immune response that remains to be characterized.

#### Duox-mediated ROS production is most likely a signalling mechanism rather than an antibacterial effector

Early experiments showed that ingestion of pathogenic bacteria by adult flies resulted in rapid upregulation of *Duox* expression in the midgut and an associated *Duox-*dependent reactive oxygen species (ROS) burst as shown by ubiquitous knockdown of this gene. Ha et al., (2005b) inferred that ROS produced by Duox, particularly HOCl, had highly bactericidal effects, as clearance of ingested bacteria was impaired in *Duox-RNAi* flies (Ha et al., 2005a). A role of Duox in the control of the gut microbiota has also been subsequently observed in other insects (Yao et al., 2016). Epithelium damage associated with Duox-mediated production of ROS was also thought to explain the increased epithelium renewal observed upon oral bacterial infection (Buchon et al., 2009a). However, most of these experiments were performed using a ubiquitous Gal4 driver to silence *Duox. Duox* is weakly expressed in the midgut compared to its expression in the adult crop, or larval hindgut and trachea (Thurmond et al., 2019), and is an essential gene required for development (Hurd et al., 2015). Experiments done in the framework of *ReproSci* were unable to reproduce a defect in bacterial clearance (*Ecc15-GFP*) in *Duox-RNAi* flies at the same time point as the initial publication (**Supplementary S23**). Furthermore, we did not observe any increase in *Duox* expression in the gut upon oral bacterial infection. Recent studies show that rather than Duox, the alternate *Drosophila* NADPH oxidase Nox stimulates stem cell proliferation upon bacterial infection (Iatsenko et al., 2018; Jones et al., 2013; Patel et al., 2019). Additionally, a recent article suggests that the impact of Duox on gut epithelial renewal is an indirect effect of Duox in the Malpighian tubules rather than the midgut (Liu et al., 2024). Other evidence suggests that lumenal ROS produced by Duox contributes to an early bacterial clearance via a signaling role that promotes visceral muscle contraction, regardless of its potential bacteriostatic activity (Benguettat et al., 2018; Du et al., 2016). We conclude that data on a bactericidal role for *Duox* in the gut context remain contradictory. More precise experiments are needed to clearly define the role of *Duox* in gut epithelial immunity, which may primarily play a signaling role rather than a microbicidal one.

#### Nitric oxide synthase (NOS) is not essential for systemic AMP expression upon oral infection

A study suggested that inhibition of nitric oxide synthase (NOS) by the inhibitor L-NAME increased larval sensitivity to Gram-negative bacterial infection, and reduced expression of a *Diptericin-LacZ* reporter in the fat body (Foley and O’Farrell, 2003). This study reported that the *NOS* gene was upregulated following infection in hemocytes, and that feeding with an NO donor induced systemic *Diptericin* expression, pointing to a role for NOS activity in mediating a robust innate immune response to Gram-negative bacteria. However, microarrays monitoring the immune response of larvae or adult flies to systemic or oral infections show that NOS is not an immune inducible gene (Buchon et al., 2009b; De Gregorio et al., 2001; Vodovar et al., 2005). Few follow-up studies have explored roles of NOS in activation of the Imd pathway (but see (Brown et al., 2009; Chakrabarti et al., 2012; Dijkers and O’Farrell, 2007; McGettigan et al., 2005)). Use of a viable null *NOS* mutant and RT-qPCR did not reveal a striking role for NOS in expression of the *Diptericin* transcript upon natural infection with the Gram-negative bacterium *Ecc1*5 (Chakrabarti et al., 2012, **Supplementary S24**). Loss of NOS may have a subtle impact on reporter expression by affecting fat body maturation, consistent with a role for NOS in regulation of the developmental hormone ecdysone (Jaszczak et al., 2015). The role of NOS in the gut to fat-body signaling, if any, requires further study.

#### MP1 is not required for wound melanization

RNAi-based studies identified two serine proteases, MP1 and SP7 (also known as MP2/PAE1), as contributors to cuticular melanization at the wound site (Tang et al., 2006). However, loss-of-function analyses have confirmed a role for SP7, but not for MP1, in this process (Ayres and Schneider, 2008; Castillejo-López and Häcker, 2005; Dudzic et al., 2015). Although MP1 is induced upon infection (De Gregorio et al., 2002), its precise function remains unclear. It may act in a different pathway, play a more subtle role in melanization, or function redundantly with other proteases.

### Insights from the reproducibility project on the roles of Eiger in immunity

#### Eiger has roles in melanization but minor effects on survival and AMP expression

A series of results implicating the TNF-related gene *eiger* in several aspects of *Drosophila* immune function (Brandt et al., 2004; Mabery and Schneider, 2010; Schneider et al., 2007) have been called into question due to the recent discovery that widely-used *eiger* mutants bore a secondary null mutation in the gene encoding the phagocytic receptor *NimC1* ((Kodra et al., 2020), which has independent roles in immunity (Melcarne et al., 2019)). Experiments for *ReproSci* using clean *egr^1^* and *egr^3^* mutants with a wild-type *NimC1* locus (Kodra et al., 2020), and *NimC1^1^* mutants for comparison, suggest that both *eiger* and *NimC1* have independent effects on survival to several pathogens, and that the effect of one or both mutations may explain several survival effects previously attributed solely to Eiger.

Contrary to previous results, we find that mutation of *eiger* does not increase tolerance to *Salmonella typhimurium* infection, and does not generally increase susceptibility to extracellular pathogens (**Supplementary S25**, consistent with (Geuking et al., 2009)). However, our results support a role for Eiger in hemolymph melanization in both adult flies and larvae consistent with (Bidla et al., 2007) (**Supplementary S26**). Experiments done in the framework of *ReproSci* did not reveal any induction of the *eiger* gene in the fat body, or a role in regulation of antibacterial peptide gene expression as initially described (**Supplementary S27**). Immune effects of *eiger* mutation are largely consistent with the reduced melanization observed in these mutants.

### Insights from the reproducibility project on the roles of Gr28b in immunity

#### Mutation of Gr28b does not produce anorexia, impair melanization, or affect humoral immunity

(Ayres and Schneider, 2009) found that a mutation in the gene encoding the gustatory receptor Gr28b caused constitutive anorexia and affected fly survival to a number of bacteria. Experiments for *ReproSci* instead suggest that the observed immune phenotypes are largely attributable to reduced melanization linked to the genetic background of the original *Gr28b* mutant, as an independent *Gr28b* mutant does not replicate these phenotypes (**Supplementary S28**). Furthermore, our results and those of (Sang et al., 2019) indicate that *Gr28b* mutants are not constitutively anorexic, and this result was likely obtained due to comparison to a wild-type line with unusually high feeding rates.

## Discussion

With thousands of papers published in the field of *Drosophila* immunity, it is difficult to summarize all we know of the fly immune system (Westlake et al., 2024). Today, the *Drosophila* immune system has been extensively characterized and serves as a model for other insects of medical or agricultural interest. The use of combined genetic approaches makes data obtained in the *Drosophila* model extremely reliable by current scientific standards. However, it is both impossible and undesirable to entirely avoid publication of controversial or irreplicable results, as science is an incremental and exploratory process where progress is made by constant course correction. That being said, contradictory data or claims that are propagated or not clearly refuted can cause a non-negligeable loss of time to scientists in research development. Taking into consideration the relevance of *Drosophila* immune research beyond the immediate community and the prestige associated with the use of sophisticated genetic approaches, it is valuable to create easily accessible assessments of established claims in the field, the roots of which may be buried deep in past literature.

In this paper, we have retrospectively analyzed the verifiability of claims from 400 articles accepted for publication before 2011 in the field of *Drosophila* immunity. We restricted our analysis to *Drosophila* host defense against bacteria and fungi, which the host laboratory has expertise in. It would be interesting to extend this analysis to *Drosophila* antiviral and anti-parasitoid immunity with collaboration from relevant laboratories. We expect that application of our approach to viruses and parasitoids would be more challenging, as co-evolved pathogens are expected to display more specific mechanisms that suppress the host immune system (Westlake et al., 2024).

Verifiability of claims was assessed either by analysing subsequent articles or through experimental testing, which we performed for 56 claims. This extensive assessment resource has been centralized on a website that can be accessed (https://ReproSci.epfl.ch/). We hope to continue updating this database as currently unchallenged claims are tested. Importantly, our study has revealed that most claims made in the field of *Drosophila* immunity have been supported by subsequent results over the years (Lemaitre et al., 2026). This article summarizes some of the main findings gained through this extensive reproducibility project with a specific focus on the rare claims that may be considered as challenged or more complex that initially reported. We cannot definitively exclude that some claims highlighted here as challenged could in fact be correct, and that our assessment was erroneous. Reproducibility projects are hampered by the fact that verification must be more stringent and convincing than the initial results to be widely accepted (Errington et al., 2021; Goodman et al., 2016; Shiffrin et al., 2018). However, we encourage the community to continue taking care when considering claims that have been challenged by *ReproSci*, which may be further clarified with future studies.

Except for the articles on the role of 18-wheeler in immunity (Ligoxygakis et al., 2002; Williams et al., 1997), PGRP-SD upstream of Toll (Bischoff et al., 2004; Iatsenko et al., 2016) and stimulation of the immune response by LPS (Imler et al., 2000; Kaneko et al., 2004; Leulier et al., 2003), few published results in the field of *Drosophila* immunity have been formally challenged. Even in the absence of published refutation, many of the challenged claims were already considered controversial by experts in the field. Others have quietly fallen out of the general current research interest. These may nevertheless still be a source of confusion for young scientists new to the field of *Drosophila* immunity or scientists with primary interests outside of *Drosophila* immunity that use the field as a reference. In our analysis, we were unable to reach definitive conclusions on the immune roles of IRC, Duox and NOS, although we suggest that their roles are not as simple as proposed in the initial articles.

Where possible, we have attempted to clarify the possible causes that led to irreplicable data or conclusions, as they emphasize challenges inherent to the field of *Drosophila* immunity and may point to methodologies that require cautious use (**Table 1**, see **Supplementary Table 2** for a list of challenged claims). A first major source of irreplicable results is where the genetic background is responsible for part or all the observed effect rather than the tested mutation. This is often due to use of inappropriate controls, for example where the wild-type comparison has a different genetic background than the mutant of interest. An example of this is the finding that WntD acts in the control of the Toll pathway (Gordon et al., 2005). Here, we obtained the same results as the authors of the claim when using the same mutant lines, but the result does not stand when using an independent mutant of the same gene, indicating the result was likely due to genetic background. Hemocyte numbers may also vary strongly according to genetic background and rearing conditions (Ramond et al., 2020; Sorrentino et al., 2004). This further complicates *in vivo* or *ex vivo* phagocytosis assays, which are often difficult to interpret as the effect of single mutations are often small. In contrast to developmental processes which are more tightly constrained, immune parameters are known to vary strongly among wild-type *Drosophila* lines (Lazzaro et al., 2004), making analysis of subtle effects difficult. This issue can be addressed in several ways: by using isogenized fly stocks with a controlled genetic background (Ryckebusch et al., 2025), by testing the phenotype of a mutation in an alternate background (e.g., over a deficiency) to see if it holds true, by validating loss of function phenotypes with RNAi, and by performing rescue experiments. None of these various ways offer a full guarantee, and caution in interpretation of the results is always required. A range of wild-type lines (Oregon-R, Canton S, etc.) can be tested to evaluate the natural variability of a phenotype and avoid one-to-one comparisons between a single wild-type and mutant that may be misleading. In some cases, natural polymorphisms affecting immune genes can cause apparent effects on gene expression by affecting primer binding, as suggested here for Defensin. Indeed, during this research we found that a few of published primers (Neyen et al. 2014) were not suited for testing AMP expression across genetic backgrounds.

**Table 1.**
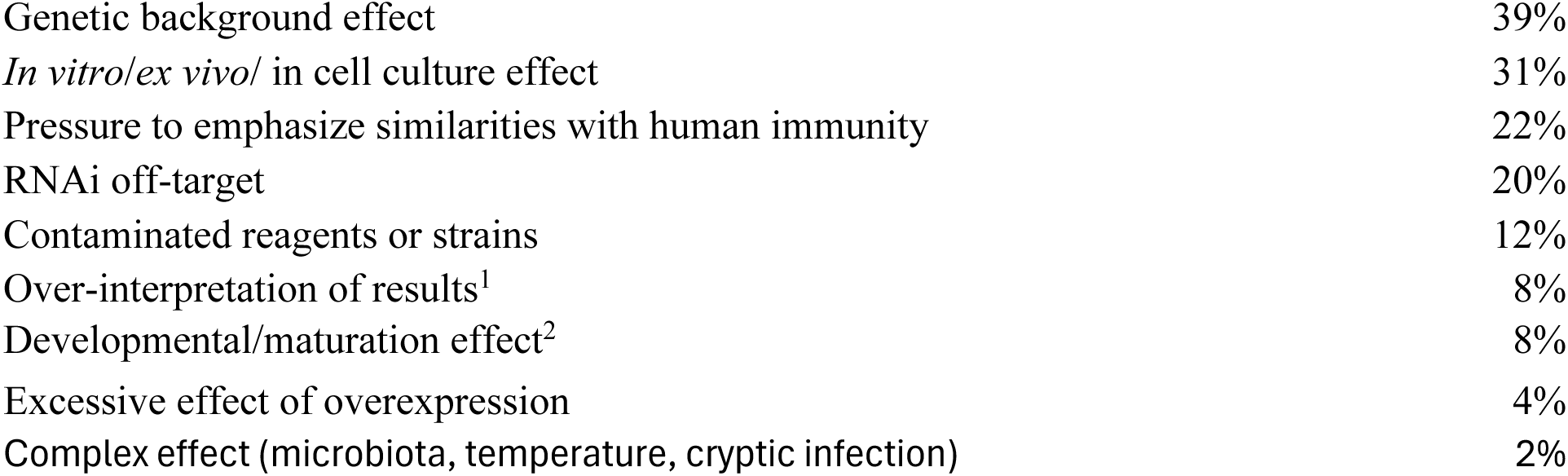
Statistics on methods associated with challenged claims.

A second source of irreplicable claims is linked to the use of *in vitro/ex vivo/*cell culture techniques that do not fully reflect results obtained using *in vivo* genetics approaches. An example is the initial implication of GNBP1 in the sensing of LPS; although this remains an important discovery, subsequent studies revealed that the role of GNBP1 was indirect, acting as an adapter between PGRP-SA and ModSP. Overexpression of genes can produce ectopic or excessive phenotypes (e.g, indicating a role for PGRP-LE in melanization) that are not supported using loss-of function mutations.

A third source of irreplicable claims is linked to the use of RNAi, which had many off-target effects when the technique was first introduced in the early 2000s (Seinen et al., 2011). The claims related to the serine proteases Spheroid, Sphinx and Spirit (Kambris et al., 2006b) and Undertaker (Cuttell et al., 2008) likely appear to be examples of this. Besides off-target effects, a limitation of the RNAi approach is that one can never be sure that it mimics a full null phenotype. As an example, the partial silencing of the serine protease gene *Grass* may explain why its role in the antifungal response was not detected using *in vivo* RNAi, but was revealed through the use of a null mutation (El Chamy et al., 2008; Kambris et al., 2006b). Newer generations of RNAi lines have greater specificity towards their target genes. Furthermore, use of several independent RNAi constructs for comparison, and inclusion of driver-only and UAS-only controls, helps to avoid off-target and background effects.

A fourth source of irreplicability is the use of contaminated or nonspecific reagents. The use of Sigma *E. coli* LPS solution as an immune elicitor has been a source of confusion in innate immune studies in both vertebrates and *Drosophila,* as effects attributed to LPS were found to be due to peptidoglycans or lipopeptides also present in the solution (Kaneko et al., 2004). Immune roles may also be attributed to genes that affect the development of immune organs, as exemplified by mutations of *18-wheeler* that delay fat body maturation (Ligoxygakis et al., 2002). Many mutations can have similar indirect effects on the immune system, as exemplified by genes involved with ecdysone signaling, which has both indirect and direct effects on immunity that are difficult to disentangle.

Irreplicability could also be due to differences in rearing conditions, such as flipping regimens that impact the microbiota, dietary composition (Lesperance and Broderick, 2021; Sannino and Dobson, 2023), and use of antifungal agents in food. Many fly stocks harbor cryptic endosymbionts such as *Wolbachia* or viruses such as Nora virus with subtle phenotypes that can regardless affect immune parameters under certain conditions (Hanson and Lemaitre, 2023; Teixeira et al., 2008). Nora virus, which can be present in up to 50% of laboratory fly lines, can significantly reduce lifespan and skew the outcomes of long-term survival studies (Franchet et al., 2025; Habayeb, 2006; Hanson and Lemaitre, 2023). The very high inducibility of AMP genes used to gauge intensity of the immune response can also be a source of confusion. These genes can be induced by 100-1000-fold upon infection, and their fold change is highly dependent on the basal expression level. If basal AMP expression is elevated in a subset of lines due to cryptic infection or differences in microbiota due to flipping frequency or geographic location, apparent differences in induction may simply result from variations in their baseline expression. The variable and high inducibility of AMP genes explains why they are often identified in microarray studies. This can however create a challenge when analyzing gene expression level, as a 5-10 fold induction is minimal for many AMPs. AMP induction can be contextualized by adding unchallenged animals as a negative control and bacteria-challenged animals as a positive control, which cover the range of induction of AMP genes. Selection of appropriate gene readouts and standards are also important (Neyen et al., 2014; Troha and Buchon, 2019). Lastly, another potential source of poor replicability is the variability in virulence among different strains of the same microorganism. This highlights the importance of carefully defining and standardizing the microbial strains used in experimental studies, a consideration that also applies to our own study. Finally, we observed that a number of challenged claims involve statements that draw parallels with mammalian immune systems. The tendency for granting agencies and journal editors to value *Drosophila* research only as a model for mammalian immunity insidiously pressures *Drosophila* studies to be framed in a human-centric way.

In this article we have not discussed a number of claims that had no clear follow-up studies and remain unchallenged (see **Supplementary Table 3** for a list of unchallenged claims). We encourage members of the *Drosophila* community to test these, or to contribute unpublished data that addresses these claims. Some examples of these include: the regulation of the Toll pathway by the p62 homolog *Drosophila* Ref(2)P (Avila et al., 2002), the regulation of AMPs independent of Toll and Imd pathways by the transcription factor FOXO (Becker et al., 2010), the biological significance of Imd pathway activation through cleavage of PGRP-LC by bacterial proteases (Schmidt et al., 2008), or the role of PGRP-LE in antimicrobial autophagy against intracellular bacteria (*L. monocytogenes, Mycobacterium marinum*) (Yano et al., 2008). In these studies and others, further research should clarify if the observed effect can be reproduced, and if the immune effects are direct or indirect.

In conclusion, our study has assessed the replicability of many claims published in the field of *Drosophila* immunity before 2011. Many articles published in this period were simpler in terms of the number of experiments and techniques used than modern studies, allowing for more efficient assessment. Assessing recent articles might be more useful to the community, to reduce the use of irreplicable data in current areas of research. However, this would likely be challenging as we would lose the benefit of hindsight and progress in the field that comes with time. Here we have favored analysis by claim rather than by article, which allows greater nuance and illustrates that articles may contain both replicable and irreplicable claims. This highlights the fact that many articles have scientific merit, despite containing challenged claims.

One important goal of this project was to make assessments of claims mainly debated in closed circles more widely accessible, and prevent time loss for scientists who may base research on claims that are widely (but quietly) considered irreplicable. We hope that this effort can assist our community by focusing attention on more promising areas of research, by providing a retrospective on foundational findings in the field, and by identifying common sources of irreplicability that can be controlled for in the future. In the framework of this project, we opened a web interface that allows the community to comment on these claims and contribute to their assessment. The program used to generate the database is open access, and could eventually be used by other communities to centralize post-publication assessments of articles by claims across research fields.

A companion article (Lemaitre et al., 2026) discusses the metascience aspect of this project by analyzing patterns of irreplicability and digging into some sociological reasons behind the publication of erroneous data.

### Limitations

This work has several limitations. First, the selection of claims and determination of replicability was influenced by the annotating authors (HW and BL). While the *ReproSci* website is open to the community, many PIs are reluctant or uninterested in the notion of testing old claims that represent scientific “cold cases”. Second, the leading author has published several articles that were included in the analysis, including 44 articles where B.L. is listed as co-author (4 as first author and 22 as last author.) Third, selection of the claims to be tested experimentally has the potential to create biases that may impact that experimental validation or simplify the complexity and nuance of the original findings. While we recognize that our approach has subjective elements, we hope that the community-accessible website will encourage researchers to contribute data and opinions from a range of sources to improve objectivity. Our intent is to honor the efforts of the *Drosophila* community while providing a complement that combats the bias against publishing negative results, which can regardless be essential and informative.

Where possible, the likely reasons behind major claims that were challenged were categorized (see full list in **Supplementary Table S2**). This table shows the likely reasons for major claims to be challenged, and the percentage of claims that were challenged due to that reason. Only one major claim per article was considered, as the same methodological issue often applied to multiple claims from the same article (Nb of articles=49). 1: The data were interpreted in a certain (erroneous) way due to a lack of available context at the time that the article was published, or alternative explanations were not effectively ruled out. 2: The effects observed were not due to a direct effect on immunity but instead due to effects on development of immune tissues or organs, including hemocytes.

## Methods

### Generation of the list of annotated articles

The purpose of the *ReproSci* reproducibility project is to determine whether a wide range of claims published in the field of *Drosophila* immunity are verifiable using independent sources. A list of publications was generated using a curated search string on the publicly available PubMed database. This search string (**Box1**) included keywords, MeSH terms and prominent authors in the *Drosophila* immunity field, and generated an initial list of approximately 2000 publications. This pool was then manually curated to remove 1) reviews and opinion pieces, 2) topics unrelated to *Drosophila* immunity captured by the search string, articles focusing mostly on virulence factors and gut homeostasis, and 3) immune topics outside the scope of Lemaitre lab expertise, including hematopoiesis, cell biology, immunity to viruses, immunity to parasitoid wasps, and immunity in other species. Select articles not captured by the search string but known and considered to be key articles in the field were manually added to this selection. The list was then restricted to the date range of interest (1959-2011). This limitation was imposed to allow a time frame in which follow-up studies may have been published that may verify, expand on or contradict the content of the publications selected for the study. These restrictions resulted in a primary list of 400 articles.

The list of articles delivered by the first search string, those that were removed and added, and the final selected list can be found in **Table S1**.

**Box 1.**
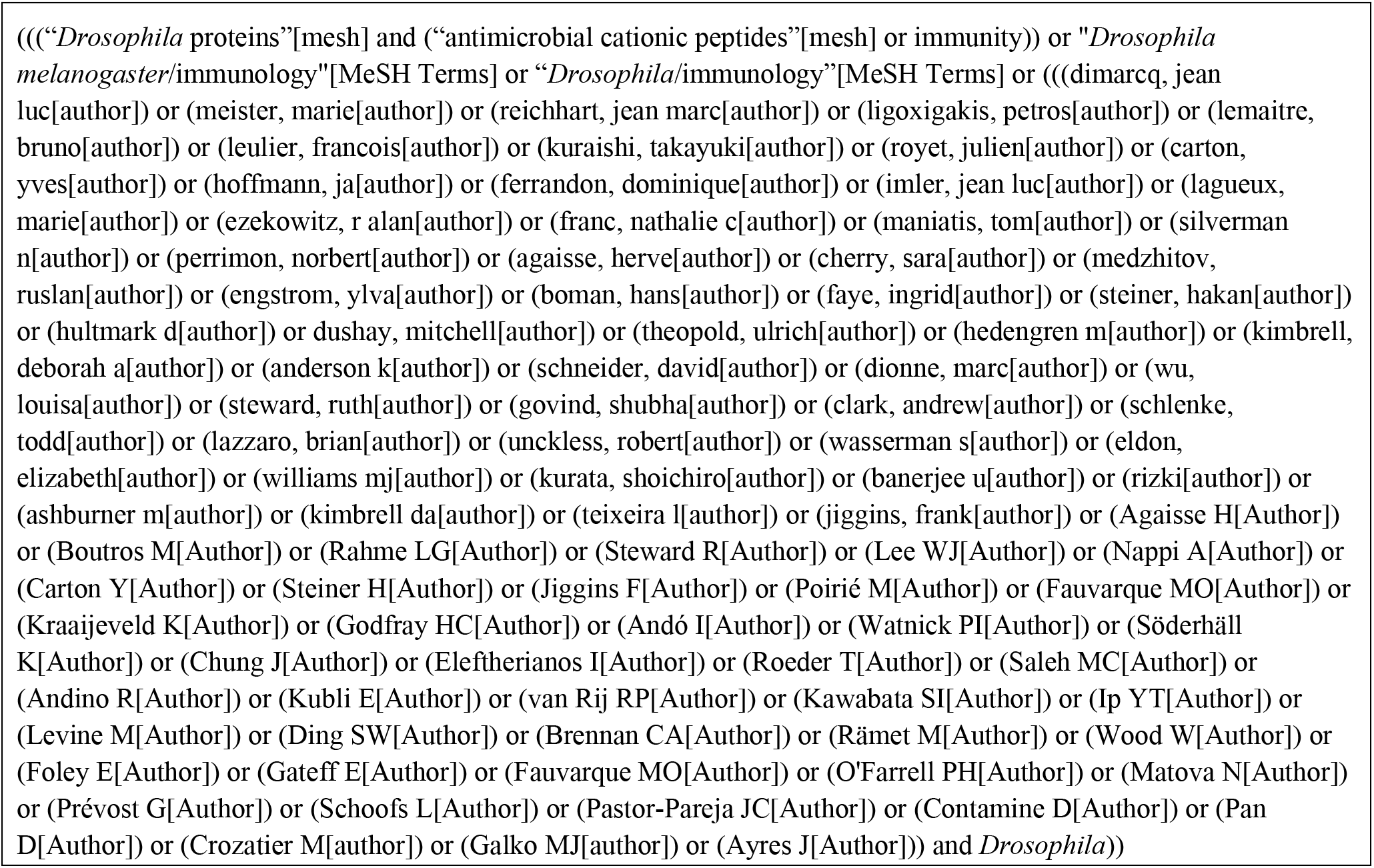
Search string used on PubMed to generate the initial pool of publications that was subsequently manually curated.

### Article annotation

Selected primary articles were annotated by a single researcher (HW) and reviewed by a second researcher (BL) to ensure consistency across the project. The *ReproSci* online database (https://ReproSci.epfl.ch/) was constructed to organize the annotations and provide an interactive and searchable interface. Claims were extracted through careful reading of the selected primary articles and organized according to their importance within the paper: a single **Main claim** representing the title claim of the paper or overarching finding; 2-4 **Major claims** emphasized in the abstract or those contributing to the overarching finding; and 0-4 **Minor claims** representing other findings or note not emphasized in the abstract. Wording of the claims was paraphrased and structured in an attempt to represent claims made directly by the authors, rather than interpretations made by the reader. Other supporting data such as the methods used in the paper and evidence used to support claims was also summarized.

Claims were then cross-checked with evidence from previous, contemporary and subsequent publications and assigned a verification category (see **Box 2**). A short statement explaining the evidence and citing sources was written to justify the categorization and provide additional context for the claim; this assessment was linked to each claim and can be viewed on the *ReproSci* website. Supplementary documents summarizing complex or ambiguous topics were also written where necessary and can be viewed on the *ReproSci* website.

**Box 2.**
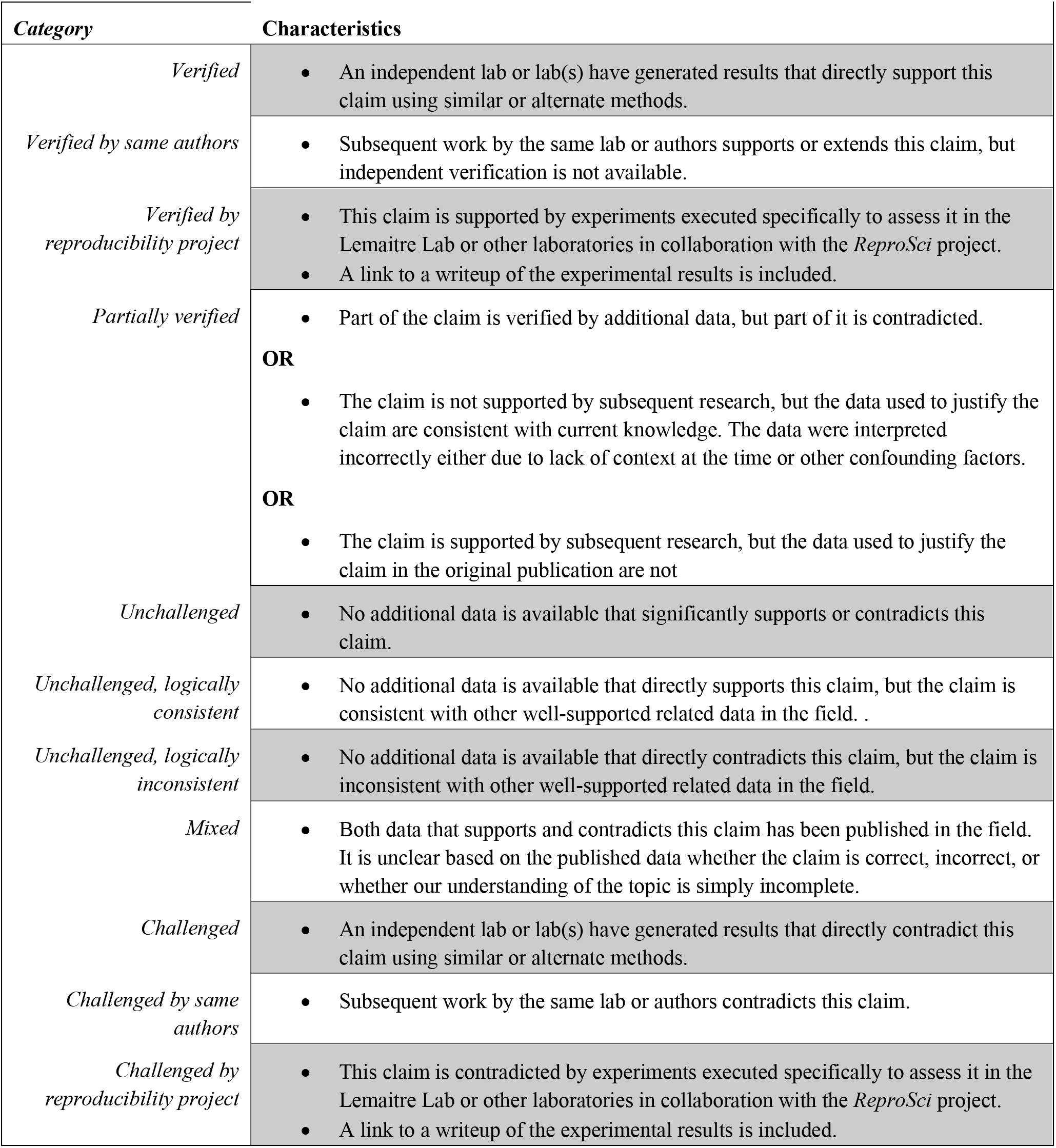
Verification categories used to assess claims.

### Experimental validation

A subset of claims that lacked follow-up in the published literature were selected for direct experimental reproduction by various authors from different laboratory with relevant expertise. This project is concerned with the accuracy of the conclusions themselves rather than the reliability of methods, so we employed indirect/inferential reproducibility/replicability, where experimental procedures that may be different from those originally used to reach a conclusion are used to independently verify it. When completed, a document was generated detailing the experiments, including methods and conclusions, conducted to verify an individual claim, which here are summarized as supplementary information. HW and BL take responsibility for the statements made following experimental validation of claims, and acknowledge that our verification experiments could also be erroneous. These short research projects were also uploaded to the *ReproSci* website, and the assessment of the claim was adjusted from ‘Unchallenged’ to ‘Verified/Challenged by reproducibility project’ as appropriate. The *ReproSci* website was made available to the *Drosophila* scientific community on July 11^th^ 2023, with an email to encourage community members to comment on annotations and verifications and contribute evidence. Experiments performed by various laboratories in the framework of the *ReproSci* project are summarized in supplemental documents S1 to S28 with their own materials and methods. Additional experimental validation of claims not highlighted in this article can be found on the *ReproSci* website.

### The *ReproSci* web interface

A versatile web interface was generated to display the 400 articles with annotations and assessments of the reproducibility of the claims made in these articles. This web interface allows i) easy generation of a defined database of articles including titles, authors, date, abstract; ii) an interface for annotation of each article in terms of major, main and minor claims, as well as fields for methods and additional supporting information, iii) a system that allows each claim or individual evidence to be assessed as verified, unchallenged, challenged, with an associated written assessment including references that were used to assign the claims to a chosen category, iv) outputs that tabulate and enumerate articles and claims according to assessment categories in the database, providing data that allow analysis of multiple parameters of reproducibility, and v) an interactive interface allowing community members to comment on and discuss claims and assessments, including uploading of documents to support their discussion. The implementation of the web interface involved the creation of a Ruby-on-Rails web application associated with a PostgreSQL database to store the data in a structured manner. At least 2 main tables store on one hand the annotations and on the other hand the relationships between them (assessments, evidence, comments). Annotations were indexed on-the-fly with the SOLR search engine, allowing subsequent free text search on them. A large part of Javascript code runs on the client side to assist users while typing annotations. Importantly, the application allows registered users to create new workspaces, each workspace focusing on a research subject. In a workspace, the owner of the project is able to moderate and to assign roles to certain users, such as the ability to add articles, enter annotations and assessments, or comment. Through moderation, community comments may be considered, and assessments updated with new information. The code of this web interface can be found at https://github.com/fabdavid/ReproSci/.

## Acknowledgments

We thank the Bloomington Stock Center in the USA and the Vienna *Drosophila* Resource Center (VDRC) for fly stocks; Paola Bellosta, Oren Schuldiner, Julien Royet, Michèle Crozatier and many members of the *Drosophila* community for providing fly strains or wasps; the BioImaging & Optics Platform (BIOP) in EPFL for confocal microscopy and the BioInformatics Competence Center of UNIL-EPFL for data analysis. We thank Neal Silverman for sharing information on Dnase II, Carolina Barillas-Mury on IRC, and Mika Rämet and Monty Krieger on Sr-C1.This work was supported by Swiss National Science Foundation 310030_189085 (BL) and the open-source program by ETH-Domain’s Open Research Data (ORD) Program (2022).

## Author contributions

Conceptualization: HW, BL

Methodology: HW, BL

Experimental Investigation: HW, YM, TEdB, AC, CM, TS, MP, ZL, YW, LJ, NP, AD, YT, AK, KK, FA, RH, SC, J-PB; FS, MAH, SR, SK, GR, GL, HJ, LX, FD (Francesca Di Cara), EK, NK, PSK, BL

Web interface design and coding: FD (Fabrice David)

Funding acquisition: BL, HW

Project administration: BL

Supervision: BL

Writing – original draft: BL, HW

Writing – review & editing: BL, HW, FM

## Supplementary materials

**Supplementary Table S1. List of the 400 articles on *Drosophila* immunity selected for analysis in the *ReproSci* study**

List of articles retrieved from PubMed with the search string (**Box 1**), list of articles manually removed including reviews, articles not focused on *Drosophila melanogaster*, articles related to viral and parasitoid immunity, and articles focusing on virulence factors.

**Supplementary Table S2: A selection of contradicted major claims discussed in this article.**

Major claims that were identified as contradicted and likely reasons that the study arrived at that conclusion. See the *ReproSci* website for a full list of contradicted claims, and updates to assessments.

**Supplement Table S3: A list of unchallenged major claims**

A list of unchallenged major claims that have received no follow-up studies directly assessing their reproducibility.

**Supplement Table S4: A list of newly generated mutations**

A list of mutants that were generated in the framework of the *ReproSci* project.

**Supplementary materials S1-S28**

Experimental validations performed by various laboratories are shown as supplements with their own methods sections and references.

