## Supplementary file for "Assessing the Replicability of Scientific Claims in Drosophila Immunity Through an Analysis of 400 Publications"

### Supplementary materials: Experimental tests of claims

### Supplementary S1: The roles of GNBP1, PGRP-SA, and PGRP-SD in *Drosophila* proModSP autoactivation

**Abstract**

*In vitro* experiments were carried out to verify the previous results that GNBP1 is needed to process and present peptidoglycans to PGRP-SA (Wang et al., 2006) and that PGRP-SD can interact with GNBP1 to enhance the interaction between GNBP1 and PGRP-SA (Wang et al., 2008). Contrary to these results, we found that GNBP1 did not process peptidoglycan in the absence or presence of PGRP-SA. Furthermore, pre-incubation of GNBP1 with peptidoglycans did not enhance proModSP autoactivation, nor did the co-presence of PGRP-SD influence this process under our experimental conditions.

**Introduction**

Wang et al. (2006) reported that recombinant GNBP1 hydrolyzed peptidoglycans from *Micrococcus luteus* and *Enterococcus faecalis* but not from *Staphylococcus aureus* or *Bacillus subtilis*. The authors also reported that PGRP-SD interacted with GNBP1 to stabilize the interaction between GNBP1 and PGRP-SA and increased the signaling capacity of GNBP1/PGRP-SA *in vivo* (Wang et al., 2008). Yet, Shan et al. (2023) recently showed that *S. aureus* and *E. coli* peptidoglycans caused partial autoactivation of freshly prepared proModSP in the presence of GNBP1 and PGRP-SA without PGRP-SD. This incomplete autoproteolysis also occurred after β-1,3-glucan was incubated with GNBP3 and proModSP. In this work, we adopted a highly sensitive method to determine whether GNBP1 could hydrolyze fluorescein-labeled *M. luteus* peptidoglycan. *In vitro* reconstitution tests were performed to test whether GNBP1 preincubation with peptidoglycans or co-presence of PGRP-SD could enhance proModSP autoproteolysis.

**Methods**

EnzChek® Lysozyme Assay Kit (E-22013, Thermo Fisher Scientific) and chicken egg white lysozyme (L-040-10, Gold Biotechnology), DAP-type peptidoglycan of *E. coli* K12 (tlrl-ksspgn, InvivoGen) and Lys-type peptidoglycan of *S. aureus* (tlrl-pgns2, InvivoGen) were purchased from the vendors. *M. luteus* peptidoglycan was purified from bacterial culture (Sumathipala and Jiang, 2010). *Drosophila* PGRP-SA, GNBP1, proModSP, and *M. sexta* PGRP4 were expressed in baculovirus-infected insect cells and purified as in previous studies (Shan et al., 2023; Kariyawasam et al., 2022). The coding region of PGRP-SD was amplified by PCR using primers j1553 (5′-CCCATGGGAATTCCGGAAGTACCGATCGTG) and j1554 (5′-GAAGCTTACTCGAGCAT TTCTTCGGACCAG) and the cDNA template (TA01534, *Drosophila* Genomics Resource Center). The purified product was TA cloned and confirmed by sequencing. The *Eco*RI-*Xho*I fragment was subcloned into the same sites in pMFH6 (Lu and Jiang, 2007). Bacmid and baculovirus generation, Sf9 cell infection, and PGRP-SD purification were performed as described before (Sumathipala and Jiang, 2010).

**Results**

We used the purified recombinant proteins to test for possible hydrolysis of peptidoglycan (Fig. 1). Incubation of buffer, BSA, and GNBP1 with the labeled substrate led to decreases in fluorescence intensity, likely due to photobleaching of the fluorophore. In the positive controls of chicken lysozyme and *M. sexta* PGRP4, an *N*-acetylmuramoyl-ʟ-alanine amidase that hydrolyzes the substrate (Hou et al., 2025), *M. luteus* peptidoglycan was hydrolyzed at 22131 ± 3337 and 26.9 ± 2.1 U/μg, respectively. While a minor signal increase was detected in the PGRP-SA reaction (2.0 ± 0.1 U/μg), adding GNBP1 did not cause additional increase. This confirms that GNBP1 does not participate in peptidoglycan processing, either independently or by enhancing PGRP-SA activity.

We then tested the role of PGRP-SD in the activation of ModSP using an in vitro approach. We incubated *E. coli* or *S. aureus* peptidoglycan with GNBP1 for up to 24−48 h, then PGRP-SA and PGRP-SD for 30 min, and finally proModSP for 90 min, there was no increase in the amount of ModSP catalytic domain between 0 and 24/48 h (Fig. 2A, lanes 2−4 and lanes 5−8), indicating that preincubation of the two peptidoglycans with GNBP1 had no effect. For *M. luteus* peptidoglycan,

PGRP-SD or PGRP-SA did not affect the proteolytic activation of proModSP (Fig. 2B, lanes 1−3 vs. lane 4). The 31 kDa catalytic domain of ModSP appeared in the presence of proModSP with or without PGRP-SD (Fig. 2B, lanes 5−8). This experiment also tested if the lack of PGRP-SD affects proModSP autoactivation caused by PGRP-SA and GNBP1 recognition of *M. luteus* peptidoglycan, and the necessity of GNBP1 preincubation (lanes 2−4, 20 h; lanes 5−8, 0 h). As long as proModSP was present, we did not detect any change in intensity of the 31 kDa band, which was present in proModSP only controls (Fig. 2A, lane 1). Addition of PGRP-SD did not affect proModSP activation (Fig. 2B, lane 2 vs lane 4; lane 6 vs lane 8). Again, neither did GNBP1 preincubation with the peptidoglycan for 20 h discernibly stimulate proModSP autoactivation (Fig. 2B, lane 2 vs lane 6; lane 4 vs lane 8).

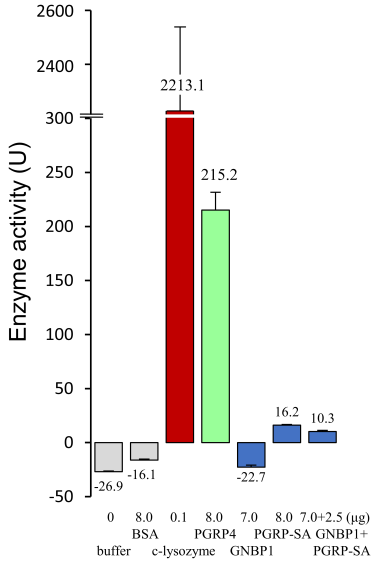

**Fig. 1. Hydrolysis of *M. luteus* peptidoglycan labeled with fluorescein.** Proteins at the indicated amounts were separately incubated with the substrate in 50 μl 0.1 M NaCl, 0.1 M sodium phosphate, pH 7.5, according to the manufacturer's protocol. Fluorescence intensity (FI) at 530 nm in each well was monitored at one-minute intervals on a microplate reader (Synergy H1, BioTek, $\lambda_{ex}$: 485 nm) to test the rates of signal change. One unit of activity is defined as the amount of enzyme causing increase in one FI unit per minute. The activities were plotted as mean ± Standard Deviation (SD) (n = 3). *Gray bars*, negative controls of buffer and BSA; *maroon* and *green bars*, positive controls of chicken lysozyme and *M. sexta* PGRP4; *cyan bars*, samples of GNBP1, PGRP-SA, and their mixture.

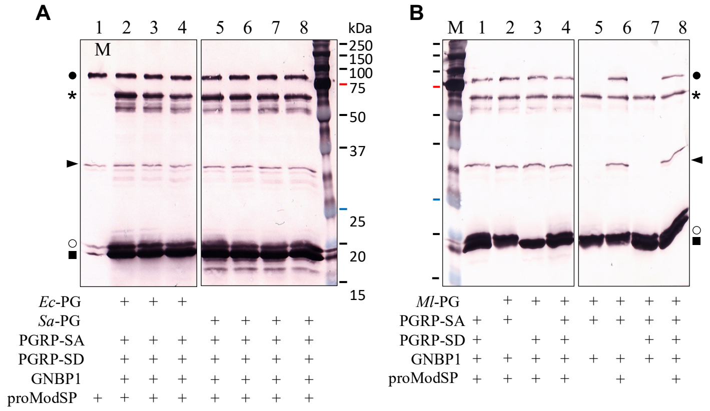

**Fig. 2. Assay of proModSP autoactivation.** (**A**) *E. coli* peptidoglycan (2 μg) and GNBP1 (800 ng) were incubated in 20 μL buffer A (20 mM Tris-HCl, pH 7.5, 20 mM NaCl, 5 mM $CaCl_2$, 0.001% Tween-20) at 16°C for 0 h (lane 2), 12 h (lane 3), and 24 h (lane 4). PGRP-SA (500 ng) and PGRP-SD (500 ng) were then added to the reactions and incubated at 37°C for 30 min, before proModSP (800 ng) was added to the reactions and incubated at 37°C for 1.5 h. Similarly, *S. aureus* peptidoglycan (2 μg) and GNBP1 (800 ng) were incubated in 20 μL buffer A at 16°C for 0 h (lane 5),

12 h (lane 6), 24 h (lane 7), and 48 h (lane 8). PGRP-SA (500 ng) and PGRP-SD (500 ng) were then added to the reactions and incubated at 37°C for 30 min, before proModSP (800 ng) was added to the reactions and incubated at 37°C for 1.5 h. (**B**) GNBP1 (400 ng) was incubated with *M. luteus* peptidoglycan (1.5 µg) in 15 µL buffer A at 25°C for 20 h (lanes 2–4) or 0 h (lanes 5–8), and then PGRP-SA (250 ng) and/or PGRP-SD (250 ng), and ModSP (400 ng) were added to the reactions and incubated at 37°C for 1.5 h. The reaction mixtures and controls were resolved by 12% SDS-PAGE under reducing condition and detected by immunoblotting using anti-6×His antibody. Sizes and positions of the $M_r$ markers are indicated, with the 75 kDa bar *highlighted red*. The GNBP1, PGRP-SA, PGRP-SD, proModSP, and ModSP catalytic domain are marked with an *asterisk*, *open circle*, *square*, *dot*, and *arrowhead*, respectively.

#### Discussion and conclusion

Using purified proteins, we could not detect any hydrolyzing activities of GNBP1 on peptidoglycan. We however detected low hydrolytic activity of PGRP-SA. As we reported before (Shan et al., 2023), a measurable level of proModSP activation during preparation positively and negatively impacted cascade elucidation. Recombinant proModSP and its orthologs in other insects are difficult to activate *in vitro*. While ModSP along with its precursor was conveniently used to activate its substrate, procSP48, the presence of 31 kDa catalytic domain made it difficult to detect minor band intensity increases caused by peptidoglycan recognition using immunoblot. In spite of this limitation, our data do not support GNBP1-mediated peptidoglycan hydrolysis or its enhancement of proModSP activation. Neither did PGRP-SD enhance this process in a noticeable way. These conclusions are consistent with the knowledge on ModSP activation in multiple insects (Kim et al., 2008; Takahashi et al., 2015; Wang et al., 2020; Wang et al., 2022), which do not seem to involve peptidoglycan hydrolysis by GNBP1 during preincubation or PGRP-SD stabilization of the interactions among PGRP-SA, GNBP1, and proModSP.

### Supplementary S2: Spirit is not required for Toll activation

**Abstract**

A loss-of-function *spirit*$^{DR32}$ mutation was used to investigate a previous result obtained using RNAi that the serine protease *spirit* is required for Toll activation and for survival to Gram-positive bacteria and fungi (Kambris et al., 2006). Contrary to previous results, we find that *spirit* is not required for induction of the Toll pathway and is not required to mount an effective immune response.

**Introduction**

(Kambris et al., 2006) found that RNAi knockdown of *spirit* (*CG2056*), a serine protease, produced an immune deficiency phenotype similar to Toll pathway mutants including high susceptibility to septic infection with *Enterococcus faecalis* and *Candida albicans* and natural infection with *Beauveria bassiana,* and failure to induce *Drosomycin* following infection with *Micrococcus luteus*. Notably, *E. faecalis* was used at an $OD_{600}$ of 200, which is higher than the concentration of this bacterium typically used for survivals, since an $OD_{600}$ of 5 is already significantly lethal to wild-type flies (Hanson et al., 2019). It is not clear from the methods at which temperature the survivals were performed, but we assume flies were maintained at either the rearing temperature of 25°C or at 29°C, a temperature more suited to bacterial growth. Although qPCR was performed using males only in this study, it is not clear whether male or female flies were used for survivals.

To determine whether *spirit* is required for Toll pathway activation, we tested Toll pathway activity and survival to Gram-positive pathogens in *spirit*$^{DR32}$ null mutants.

**Methods**

*Fly lines and bacterial strains.*

*Spirit*$^{DR32}$ mutants were generated by homologous recombination as previously described (Baena-Lopez et al., 2013). The insertion, which deletes 622bp including the start site of *spirit* and half of the protein (Figure 1), was verified by PCR. *w*$^{1118}$ flies (BDSC #3605) were used as a wild-type control; *Hayan-psh*$^{Def}$ (Dudzic et al., 2019) or *PPO1*$^{D}$*, PPO2*$^{D}$ (Binggeli et al., 2014) double mutants were used as melanization-deficient comparisons; and in survival experiments *spz*$^{rm7}$ (Neyen et al., 2014) or *Bom*$^{\Delta 55C}$ *Drosdel* (Clemmons et al., 2015; Hanson et al., 2019) flies were used as immune-deficient mutants. The strains *E. faecalis* (ATCC 29212) and *M. luteus* JL (Jean Lambert collection, Strasbourg) were used in this study.

*DNA extraction and PCR*

*Spirit*$^{DR32}$ flies were confirmed to bear the expected mutation using PCR. DNA was extracted from male and female flies by macerating single flies in 50uL squishing buffer (10 mM Tris--Cl pH 8.2, 1 mM EDTA, 25 mM NaCl and 200 ug/ml Proteinase K, with the enzyme added fresh) followed by incubation at 37°C for 30 minutes, and 2 minutes at 95°C. PCR was performed in a 25uL reaction volume using QuantaBio AccuStart II PCR SuperMix using the conditions specified below. Bands were separated on a 1% agarose gel.

- *Spirit*$^{DR32}$ primer set: amplifies junction between inserted Hsp70 gene and interrupted Spirit sequence.
    - Forward: 5'- TGT CAG GGA GTG AGT TTG CTT AA -3'
    - Reverse: 5'- CAA AGT TGT TGG CGA TAA GGG C -3'
    - Annealing temperature: 58°C
    - Extension time: 30s
    - No band amplifies in flies not bearing the *Spirit*$^{DR32}$ mutation.
- Wild type *spirit* primer set: amplifies region replaced by the plasmid in the *spirit*$^{DR32}$ mutant.
    - Forward: 5'- TCA TTT GGC CCT AGT TTC GAG C -3'
    - Reverse: 5'- CAA AGT TGT TGG CGA TAA GGG C -3'
    - Annealing temperature: 58°C
    - Extension time: 30s
    - No band amplifies in flies bearing the *Spirit*$^{DR32}$ mutation.

*Melanization in adult flies*

5 to 7-day old male and female flies reared on cornmeal fly medium (6% cornmeal, 6% yeast, 0.6% agar, 0.1% fruit juice, 10.6 g/L moldex and 4.9 mL/L propionic acid) at 25°C were anaesthetized with $CO_2$ and pricked on the left side of the thorax anterior and ventral to the wing joint with a 0.2 mm tungsten needle. The needle was sterilized with ethanol prior to clean injury or dipped in bacterial pellets concentrated from cultures of *M. luteus* and diluted to $OD_{600}$=200 with PBS prior to septic injury. Bacteria were grown in overnight shaking cultures at 29°C in LB medium. *M. luteus* was heat-killed prior to injury by heating the pellet to 67°C for 1 hour. Flies were incubated at rearing conditions for 18h after pricking and melanization of the injury site was scored. Melanization was scored as 'Strong' if injury produced a dark brown or black spot similar to that typically produced in wild-type flies, 'Weak' if only a light yellow or brown spot was visible, and 'None' if no melanization was apparent. 10 flies per sex per genotype were scored in two independent experiments; male and female data were combined as they yielded similar results.

*Melanization in larvae*

In clean injury experiments larvae were pricked at the posterior end with a 0.1mm needle and placed on grape juice agar in a humidification chamber at 25°C for 2h before assessing melanization.

*Survivals*

5 to 7-day old male and female flies reared on cornmeal fly medium (6% cornmeal, 6% yeast, 0.6% agar, 0.1% fruit juice, 10.6 g/L moldex and 4.9 mL/L propionic acid) at 25°C were anaesthetized with $CO_2$ and pricked on the side of the thorax anterior and ventral to the wing joint with a 0.2 mm tungsten needle dipped in a concentrated bacterial pellet. Survivals were incubated at 25°C and flies were transferred to fresh vials every 2-3 days. ~20 flies were used per genotype per experiment; two independent experiments were performed for each treatment.

*qPCR for Toll pathway induction*

Male flies were pricked with *E. faecalis* (overnight, 37°C, shaking, BHI) or *M. luteus* (overnight, 29°C, shaking, LB) and maintained on cornmeal fly medium at 25°C. For each treatment, 6-8 flies were frozen at -80°C at 6h, 24h, and 48h post infection. Two independent experiments were performed for each treatment. Total RNA was extracted from samples using TRIzol reagent (Invitrogen), and re-suspended in Milli-Q water. DNase treatment was performed using TURBO DNA-free kit (Invitrogen). Reverse transcription was performed using 0.5 mg total RNA in 10 ul reactions using Prime-Script RT (Takara) with random hexamer and oligo dT primers. Quantitative PCR was performed on a LightCycler 480 (Roche) in 96-well plates using PowerUp SYBR Green Master Mix (Applied Biosystems). *Drosomycin* expression was normalized to expression of the ribosomal protein *rp49* (*rpl32*). For graphical display, *Drosomycin* expression levels were normalized to the unchallenged (UC) condition in wild type $w^{1118}$ flies.

- *Rp49* Forward: 5' – GCC GCT TCA AGG GAC AGT ATC TG – 3'
- *Rp49* Reverse: 5' – AAA CGC GGT TCT GCA TGA G – 3'
- *Drosomycin* Forward: 5' – CGT GAG AAC CTT TTC CAA TAT GAT – 3'
- *Drosomycin* Reverse: 5' – TCC CAG GAC CAC CAG CAT – 3'

**Results and discussion**

*Spirit*$^{DR32}$ null mutant flies were fully viable and showed no obvious morphological defects. Wild type and *spirit* mutant flies and larvae had comparable melanization in response to all treatments (Figure 2A,B). Female *spirit*$^{DR32}$ flies showed a slight susceptibility to *E. faecalis* at high doses ($OD_{600}$=5, $OD_{600}$=200), but males did not (Figure 3). This susceptibility was significant ($P<0.05$, Cox Proportional Hazards test) only at $OD_{600}$=5 in females. At low concentrations of *E. faecalis* ($OD_{600}$=0.5) both males and females survived as wild type (data not shown). *Spirit*$^{DR32}$ mutant flies did not have reduced induction of *Drosomycin* compared to wild-type flies in response to either *E. faecalis* or *M. luteus* (Figure 4A,B).

Contrary to (Kambris et al., 2006), these results show that *spirit* is not required for induction of the Toll pathway and is not required to mount an effective immune response. *Spirit* is not required for *Drosomycin* induction, melanization, or survival to Gram-positive bacterial challenge. Although females showed slight susceptibility to *E. faecalis* at high concentrations, the strong susceptibility found by (Kambris et al., 2006) using RNAi to silence *spirit* was not reproduced using the *spirit*$^{DR32}$ null mutant line. Since Toll induction is normal in *spirit*$^{DR32}$ mutants, it is unlikely that susceptibility in females is due to reduced Toll activity. High throughput expression data show that *spirit* is highly expressed in the larval fat body but has very low expression in adult flies, indicating that it may play a more important role in larvae than adult flies (FlyAtlas2).

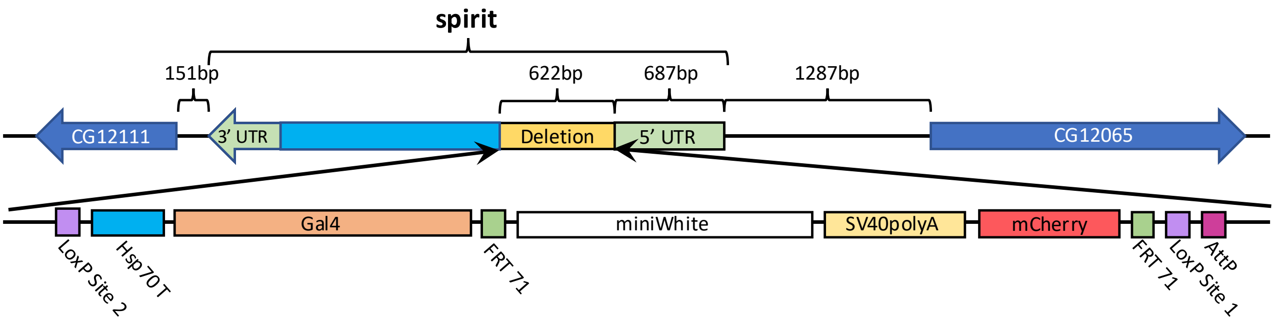

**Figure 1.** Schematic of the *spirit*$^{DR32}$ mutation generated by homologous recombination. 622bp of Spirit beginning at the translational start site was deleted and replaced with a 7918bp cassette including mCherry, miniWhite and a Gal4 driver. The 5' UTR sequence and sequence of adjacent genes are unaffected.

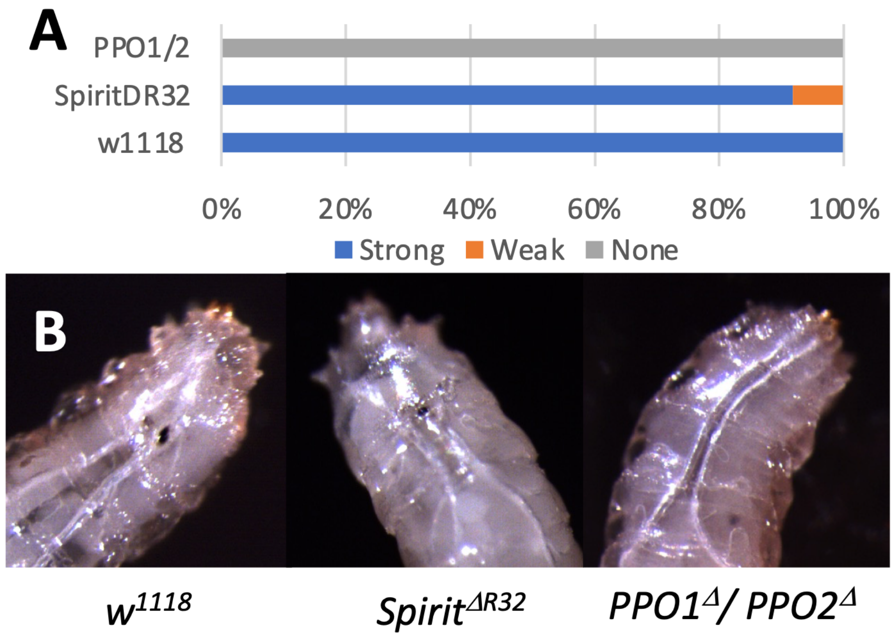

**Figure 2. A)** Proportion of adult flies of each genotype displaying strong, weak and no melanization 18h following injury with heat-killed *M. luteus*. Two independent experiments with 10 males and 10 females per genotype per experiment. **(B).** Melanization in larvae following clean injury. Representative images are shown.

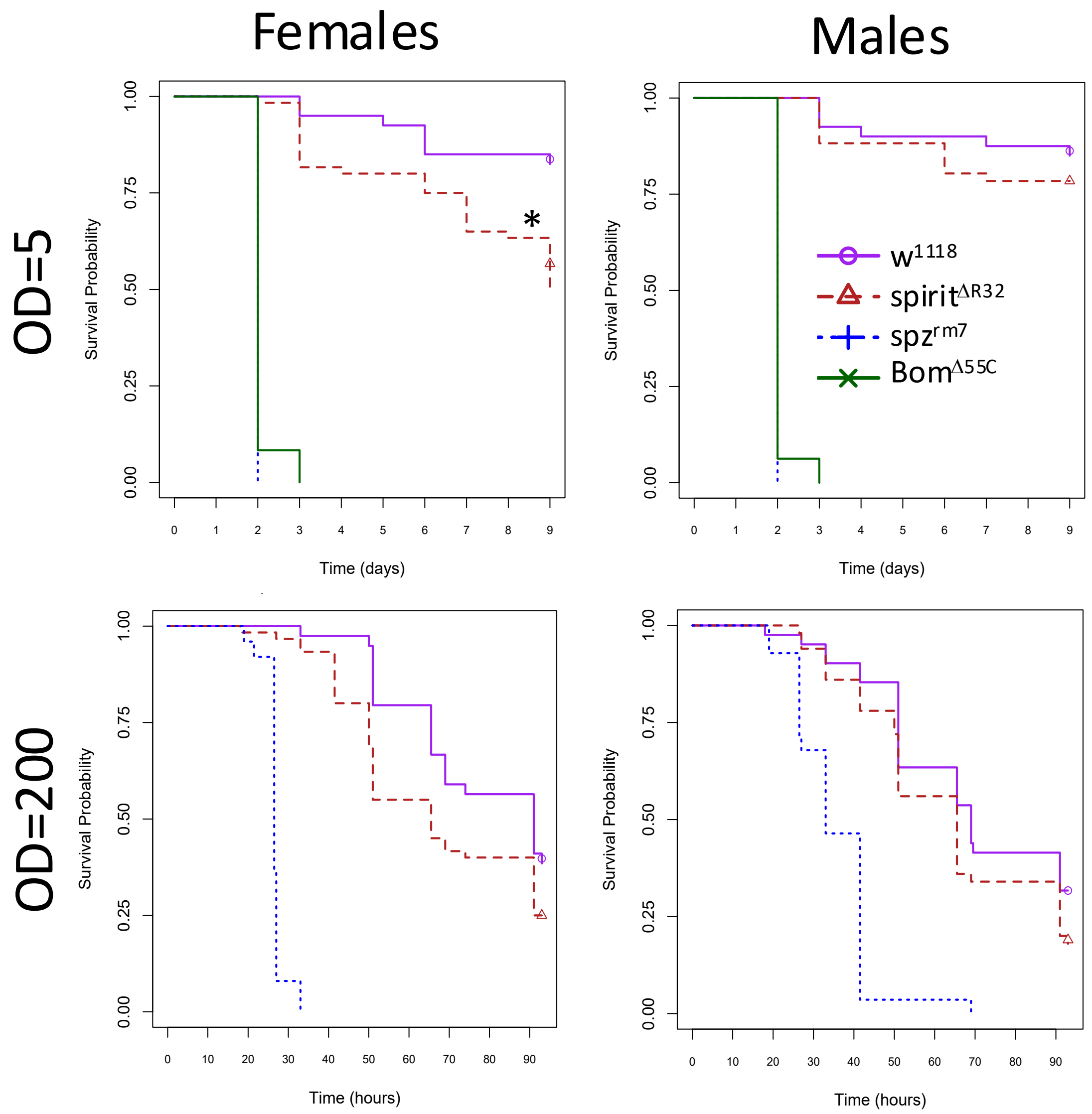

**Figure 3.** Survival of male and female flies following septic infection with $OD_{600}$=5 and $OD_{600}$=200 *E. faecalis.* ~20 flies were used per genotype per experiment; n=2 independent experiments for all survivals shown. Pooled data are shown. *, P < 0.05

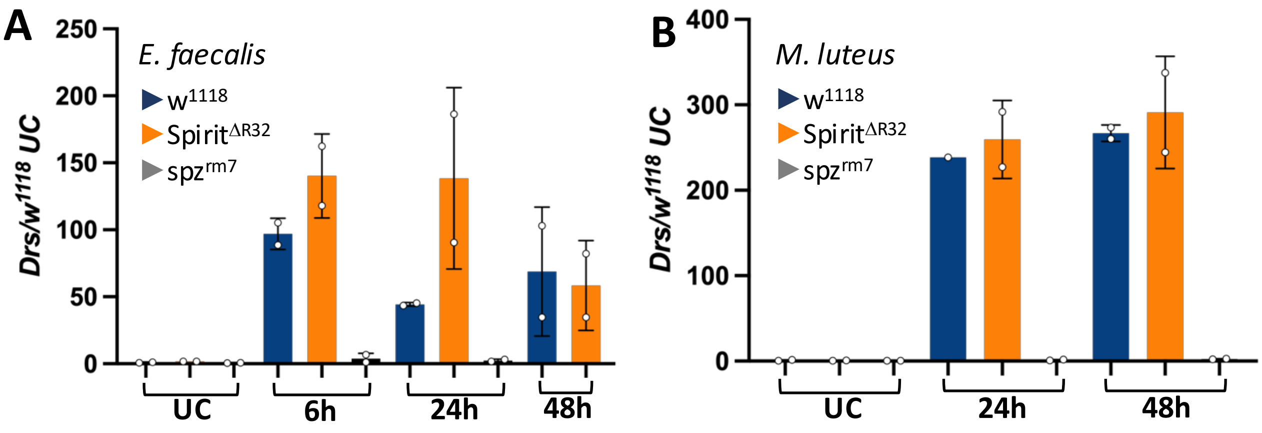

**Figure 4.** Induction of *Drosomycin* following septic injury with *E. faecalis* **(F)** and *M. luteus* **(G)**. UC=Unchallenged. n=2 experiments, 6-8 flies per treatment per experiment. Error bars represent the standard deviation.

#### Conclusion

Our data indicate that *spirit* is not required for induction of the Toll pathway. Although these results contradict (Kambris et al., 2006), they do not exclude the possibility that Spirit acts redundantly with another serine protease(s) in Toll activation or melanization.

### Supplementary S3: Spheroide is not required for Toll activation

**Abstract**

A previous result obtained using RNAi suggests that the serine protease *spheroide* (*sphe*) is required for Toll activation and for survival to Gram-positive bacteria and fungi (Kambris et al., 2006). We used a loss-of-function *spheroide*$^{D104}$ mutation to investigate this claim. Contrary to previous results, we find that *sphe* is not required for induction of the Toll pathway and is not required to mount an effective immune response against the Gram-positive pathogen *E. faecalis*.

**Introduction**

(Kambris et al., 2006) found that RNAi knockdown of *spheroide* (*sphe*)*,* a serine protease homologue, produced an immune deficiency phenotype similar to Toll pathway mutants including high susceptibility to septic infection with *Enterococcus faecalis* and *Candida albicans* and natural infection with *Beauveria bassiana,* and failure to induce *Drosomycin* following infection with *Micrococcus luteus*.

(Patrnogic and Leclerc, 2017) subsequently generated a *sphe* null mutant through imprecise excision of a Minos transposable element. These authors found that *sphe* mutants had reduced *Drosomycin* induction (70% reduction) and increased susceptibility (intermediate between wild-type and *spz*$^{rm7}$ mutant) following challenge with pathogenic Gram-positive bacteria (*E. faecalis*) but not fungi (*B. bassiana*) or non-pathogenic *M. luteus*. Precise excision of the element or overexpression of *sphe* in the mutant rescued these phenotypes. Overexpression of *sphe* further enhanced survival to *E. faecalis.* Epistasis suggested that *sphe* is part of the danger signal cascade, and protease-deficient *E. faecalis* were avirulent against *sphe* mutants and induced *Drs* as in wild type flies. *Sphe* mutant flies were also susceptible to *Staphylococcus aureus* but did not have reduced *Drs* induction in response to this bacterium. Double mutants for *sphe* and *grass* had increased susceptibility to *S. aureus*, further indicating that *sphe* acts in the danger signal pathway in parallel to the PRR pathway in which *grass* acts. Epistasis indicated *sphe* acts downstream or at the level of *persephone.* These data broadly agree with the findings of (Kambris et al., 2006).

Note that *sphe* is overwhelmingly expressed in the adult midgut (and to a lesser extent the larval midgut) and has very low expression in other tissues and life stages, suggesting a local function for this gene. Expression of *sphe* decreases following septic infection and recovers at late time points (FlySick). This protein is not present in an adult hemolymph proteome (Rommelaere et al., 2025) and is therefore unlikely to participate in the canonical Toll cascade. (De Gregorio et al., 2001) also confirm that this gene is not upregulated following infection.

In order to verify a role for *sphe* in Toll activation, we performed similar survival, expression and melanization studies in *sphe*$^{D104}$ null mutants with a modified genetic background.

**Methods**

*Fly lines*

*Spheroide*$^{D104}$ mutants generated by imprecise excision of the *Mi{ET1}spheroide*$^{MB11555}$ Minos element were obtained from (Patrnogic and Leclerc, 2017). The line carried balancers on the second chromosome, and so the 2$^{nd}$ chromosome was swapped with the *w*$^{iso}$ *Drosdel* background, and the 3$^{rd}$ chromosome was partially recombined with the *w*$^{iso}$ *Drosdel* background; the X chromosome bearing the *Spheroide*$^{D104}$ mutation was not recombined during this process. Several wild-type lines were used as comparison: *w*$^{1118}$ (BDSC #3605)*, w*$^{iso}$ *Drosdel* (Ryder et al., 2004) and *OR-R.* In survival experiments *spz*$^{rm7}$ (Neyen et al., 2012) or *Bom*$^{\Delta 55C}$ in the *w*$^{iso}$ *Drosdel* background (Clemmons et al., 2015; Hanson et al., 2019) flies were used as Toll immune deficient lines, and *Rel*$^{E20}$ mutants in the *w*$^{iso}$ *Drosdel* background were used as an Imd-deficient comparison (Ferreira et al., 2014).

*DNA extraction and PCR*

Absence of the wild-type sequence in *sphe*$^{D104}$ mutants was verified by PCR. DNA was extracted from male and female flies by macerating single flies in 50uL squishing buffer (10 mM Tris--Cl pH 8.2, 1 mM EDTA, 25 mM NaCl and 200 ug/ml Proteinase K, with the enzyme added fresh) followed by incubation at 37°C for 30 minutes, and 2 minutes at 95°C. PCR was performed in a 25µL reaction volume using QuantaBio AccuStart II PCR SuperMix using the conditions specified below. Bands were separated on a 1% agarose gel.

- *Spheroide* primer set:
  - Forward: 5'- CGA CGA GAA TAT AAG AGA CTA GC -3'
  - Reverse: 5'- ACT GGT CGT TTG GTT TAT TTG G -3'
  - Annealing temperature: 58°C
  - Extension time: 30s
  - Amplifies wild-type sequence but not $sphe^{D104}$ or $sphe^{D49}$

*Survivals*

5 to 7-day old male and female flies reared on cornmeal fly medium (6% cornmeal, 6% yeast, 0.6% agar, 0.1% fruit juice, 10.6 g/L moldex and 4.9 mL/L propionic acid) at 25°C were anaesthetized with $CO_2$ and pricked on the side of the thorax anterior and ventral to the wing joint with a 0.2 mm tungsten needle dipped in a concentrated bacterial pellet. Survivals were incubated at the temperature indicated in figures and flies were transferred to fresh vials every 2-3 days. Natural infections with *B. bassiana* were performed by shaking anesthetized flies in a vial with 200mg of spores (BB-PROTEC *B. bassiana* strain R444 from Andermatt Biocontrol). Flies were flipped into fresh vials the day after fungal inoculation. ~20 flies were used per genotype per experiment; 2-3 independent experiments were performed for each treatment. Data were analyzed in R using a Cox Proportional Hazard test. Male and female survival data were combined as they did not show significantly different trends in survival, except in survivals to *B. bassiana* where effect size differed but trends were similar between sexes.

*Bacterial culture conditions*

*Enterococcus faecalis* (ATCC 29212) and *Bacillus subtilis* 168 were cultured overnight at 37°C shaking in BHI and diluted to the desired OD in PBS. *Micrococcus luteus* JL (Jean Lambert collection, Strasbourg) was cultured overnight at 29°C shaking in LB and diluted to the desired OD in PBS.

*Quantitative-PCR*

Male flies were pricked with *E. faecalis* or *M. luteus* and maintained on cornmeal fly medium at 25°C. For each treatment, 6-10 flies were frozen at -80°C at 24h and 48h post infection. Total RNA was extracted using TRIzol reagent (Invitrogen), and re-suspended in MilliQ water. DNase treatment was performed using TURBO DNA-free kit (Invitrogen). Reverse transcription was performed using 0.5 mg total RNA in 10 µl reactions using Prime-Script RT (Takara) with random hexamer and oligo dT primers. Quantitative PCR was performed on a LightCycler 480 (Roche) in 96-well plates using PowerUp SYBR Green Master Mix (Applied Biosystems). *Drosomycin* expression was normalized to expression of the ribosomal protein *rp49* (*Rpl32*). For graphical display, *Drosomycin* expression levels were normalized to the unchallenged (UC) condition in the wild type $w^{1118}$ flies.

- *rp49* Forward: 5' – GCC GCT TCA AGG GAC AGT ATC TG – 3'
- *rp49* Reverse: 5' – AAA CGC GGT TCT GCA TGA G – 3'
- *Drosomycin* Forward: 5' – CGT GAG AAC CTT TTC CAA TAT GAT – 3'
- *Drosomycin* Reverse: 5' – TCC CAG GAC CAC CAG CAT – 3'

*Melanization in adult flies*

5 to 7-day old male and female flies reared on cornmeal fly medium at 25°C were anaesthetized with $CO_2$ and pricked on the left side of the thorax anterior and ventral to the wing joint with a 0.2 mm tungsten needle. The needle was sterilized with ethanol, dried, and dipped in bacterial pellets concentrated from cultures *M. luteus* and diluted to $OD_{600}$=200 with PBS prior to septic injury. Bacteria were grown in overnight shaking cultures at 37°C in LB medium. Flies were incubated at rearing conditions for 16h after pricking and melanization of the injury site was scored with a dark brown or black spot indicating "strong" melanization; light brown or yellow spot indicating "weak" melanization; and no apparent spot indicating no melanization. At least 20 flies

per genotype per sex were scored; male and female data were combined as they yielded similar results.

#### Results and discussion

Recombined *spheroide*$^{D104}$ mutants had similar survival to *E. faecalis* septic infection as *w*$^{iso}$ and *OR-R* wild-type flies (Figure 1A, B). These data are somewhat in contrast to (Kambris et al., 2006; Patrnogic and Leclerc, 2017) who found that *sphe*-deficient flies had slightly increased susceptibility to virulent (wild-type) but not avirulent *E. faecalis* infection. However, the susceptibility shown by (Patrnogic and Leclerc, 2017) was subtle and may be contingent on having a perfect background control. Survival of *sphe*$^{D104}$ mutants to natural infection with *B. bassiana* (Figure 1C) was also similar to wild-type. Wild-type resistance of *sphe*-deficient flies to natural infection with *B. bassiana* agrees with (Patrnogic and Leclerc, 2017) but contrasts with (Kambris et al., 2006). In response to *B, subtilis* infection, *sphe*$^{D104}$ mutants had similar survival to the *w*$^{1118}$ control and higher survival than *OR-R* and *w*$^{iso}$ (Figure 1D).

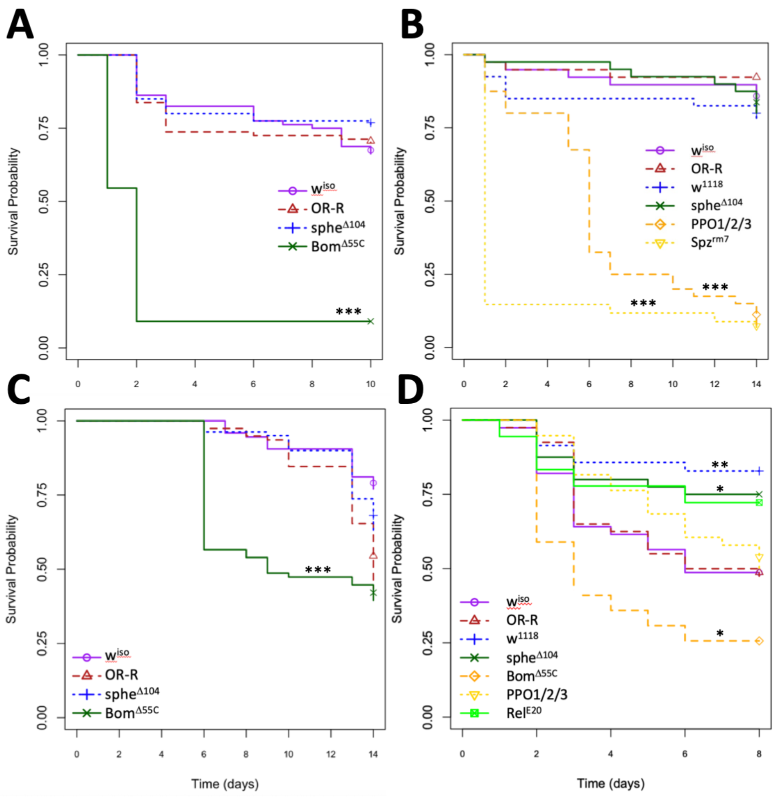

**Figure 1.** Survival of *sphe*$^{D104}$ mutant flies to infections with a variety of pathogens. No significant difference was found for *sphe*$^{D104}$ survival to *E. faecalis* $OD_{600}$=0.5 at (**A**) 25°C or (**B**) 29°C, or to natural infection with 200mg *B. bassiana* spores at 29°C (**C**). **D)** In response to *B. subtilis* $OD_{600}$=5, 25°C, *sphe*$^{D104}$ flies had somewhat higher survival than the *w*$^{iso}$ control, but similar survival to the alternate *w*$^{1118}$ line. 20 flies were used per genotype per experiment; 2-3 independent experiments were performed for each treatment.

Contrary to both (Kambris et al., 2006) and (Patrnogic and Leclerc, 2017) we found no marked difference in basal or induced *Drosomycin* expression in *sphe*$^{D104}$ mutant flies upon infection with *E. faecalis* (Figure 2A, B). (Patrnogic and Leclerc, 2017) found a ~50% in *Drs* expression in *sphe* mutants at 24hpi with *E. faecalis.* Our data shows similar *Drs* expression across all genotypes at the 24hpi time point. However, our data do indicate reduced *Drs* expression in *sphe* mutants compared to the *w*$^{iso}$ control at the 48h time point, suggesting a possible difference in the kinetics of our experiment. Further replicates are required to verify whether this is a consistent effect. Melanization in *sphe*$^{D104}$ adult flies showed only a slight qualitative reduction compared to controls (Figure 2C).

Based on these data, we conclude that loss of *sphe* does not result in a consistent reduction in Toll pathway activity upon infection with *M. luteus* or *E. faecalis.*

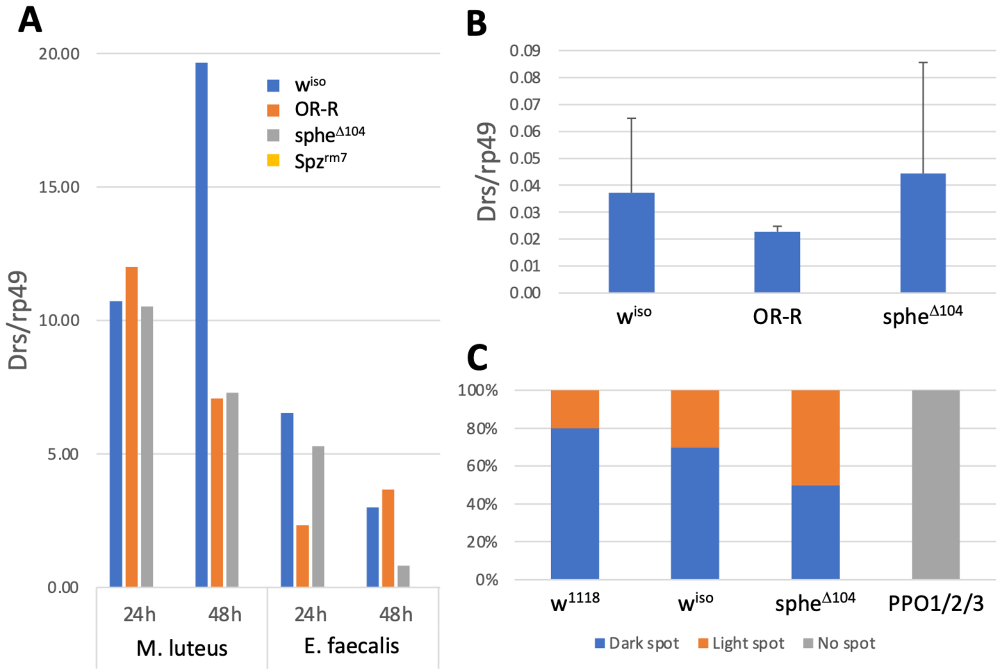

**Figure 2.** *Drosomycin* expression and melanization in response to septic injury. *Drosomycin* expression following infection with *M. luteus* and *E. faecalis* at **A)** 24h and 48h time points after infection (single replicate) and **B)** in the absence of infection (two replicates). **C)** Melanization in adult *Drosophila* at 16 hours following septic injury with live *M. luteus* (single replicate, n=20 males and 20 females for each genotype).

#### Conclusion

We find that contrary to (Kambris et al., 2006; Patrnogic and Leclerc, 2017) loss of *sphe* has no strong or consistent effect on Toll pathway gene expression, survival to *B. bassiana* or *E. faecalis* infection, or melanization. However, results (Patrnogic and Leclerc, 2017) suggest that *sphe* could have a role in survival in certain contexts, and may have a subtle role in Toll pathway activity.

### Supplementary S4: Sphinx1/2 are not required for Toll activation

**Abstract**

A previous result obtained using RNAi suggested that the serine proteases *sphinx1* and *sphinx2* are required for Toll activation and for survival to Gram-positive bacteria and fungi (Kambris et al., 2006). Using a newly CRISPR/Cas9 mediated deletion that removes both *sphinx1* and *sphinx2*, we showed that both genes are not required for Toll pathway activity. We conclude that *sphinx1/2* do not have a strong role in regulation of the Toll pathway, or in systemic immunity.

**Introduction**

*sphinx1* and *sphinx2* are two serine protease homolog genes clustered at the position 66A1 on the chromosome 3L. (Kambris et al., 2006) found that ubiquitous knock down of the serine proteases *sphinx1* and *sphinx2* suppressed Toll pathway activity and increased susceptibility to Gram-positive bacteria and fungi, including susceptibility to septic infection with *Enterococcus faecalis* and *Candida albicans* and natural infection with *Beauveria bassiana.* They also reported failure of the knocked down flies to induce *Drosomycin* following infection with *Micrococcus luteus*. No further studies have been published on the function of these genes. Furthermore, high-throughput expression data (FlyAtlas2, FlyBase modENCODE) show that *sphinx1* and *sphinx2* are expressed exclusively in the adult male accessory gland, which argues against a significant role in general immunity on an organismal level.

We generated by CRISPR/Cas9 a short deletion that remove both *sphinx1* and *sphinx2* to test their role in Toll-mediated immunity.

**Methods**

*Fly lines and Survival analysis*

We generated two fly lines carrying a small deficiency that removed both *sphinx1* and *sphinx2* (*Δ(sphinx1/2)*$^{\Delta 6}$ and (*Δ(sphinx1/2)*$^{\Delta 7}$) *in the w* iso background (Figure 1). The deletion was verified by PCR. *w*$^{1118}$ *Drosdel* (*w*$^{iso}$ ) were used as the wild-type comparison (Ryder et al., 2004). Flies were kept at 25°C on cornmeal fly medium (6% cornmeal, 6% yeast, 0.6% agar, 0.1% fruit juice, 10.6 g/L moldex and 4.9 mL/L propionic acid). 5-10 days old flies were used for survival experiments with *Enterococcus faecalis* (*E. faecalis;* $OD_{600}$ 0.1) performed at 25°C. Two independent survival experiments with 20 flies were conducted. The strains *E. faecalis* (ATCC 29212) and *M. luteus* JL (Jean Lambert collection, Strasbourg) were used in this study

*qPCR for Toll pathway induction*

Male flies were pricked with *Micrococcus Luteus* (OD600= 200, overnight, 29°C, shaking, LB) and maintained on cornmeal fly medium at 25°C. For each treatment, 6-8 flies were frozen at -80°C at 6h, 24h, and 40h post-infection. Two independent experiments were performed for each treatment. Total RNA was extracted from samples using TRIzol reagent (Invitrogen), and re-suspended in Milli-Q water. DNase treatment was performed using TURBO DNA-free kit (Invitrogen). Reverse transcription was performed using 0.5 mg total RNA in 10 ul reactions using Prime-Script RT (Takara) with random hexamer and oligo dT primers. Quantitative PCR was performed on a LightCycler 480 (Roche) in 96-well plates using PowerUp SYBR Green Master Mix (Applied Biosystems). *Drosomycin* expression was normalized to expression of the ribosomal protein *rp49* (*rpl32*). For graphical display, *Drosomycin* expression levels were normalized to the unchallenged (UC) condition in wild type *w*$^{1118}$ flies.

- *Rp49* Forward: 5' – GCC GCT TCA AGG GAC AGT ATC TG – 3'
- *Rp49* Reverse: 5' – AAA CGC GGT TCT GCA TGA G – 3'
- *Drosomycin* Forward: 5' – CGT GAG AAC CTT TTC CAA TAT GAT – 3'
- *Drosomycin* Reverse: 5' – TCC CAG GAC CAC CAG CAT – 3'
- *BomBc3*_Forward_5'_CTGATCGGCGCTCATCCCAG
- BomBc3_Reverse_5'_GGGATGAGGAGAAGCTGCGG

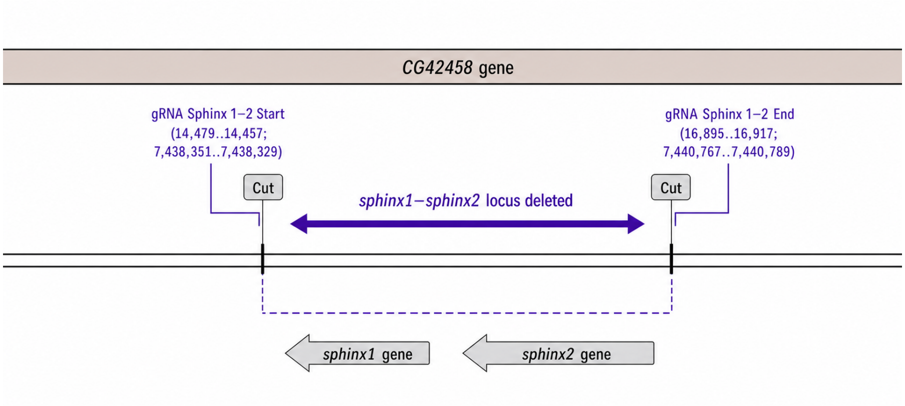

**Figure 1:** Description of the *Δ(sphinx1/2)* mutants. *Δ(sphinx1/2)*$^{\Delta 6}$ and *Δ(sphinx1/2)*$^{\Delta 7}$ flies were obtained by deleting the *sphinx1* and *sphinx2* genomic region by CRISPR/Cas9.

#### Results

*Δ(sphinx1/2)*$^{\Delta 6}$ and *Δ(sphinx1/2)*$^{\Delta 7}$ flies deleted for *sphinx1* and *sphinx2* were viable. They did not shown any marked susceptibility to *E. faecalis* in contrast to *Grass*$^{Herrade}$ flies with impaired Toll pathway activation to Gram-positive bacteria (Kambris *et al.,* 2009; El Chamy *et al.,* 2008) or Bom55C flies that lack 10 Bomanin genes (Clemmons et al.,2015) (Figure 2). *Δ(sphinx1/2)*$^{\Delta 6}$ flies express wild-type level of the Toll target genes *Drosomycin* and BomC3 upon septic injury with *M. luteus* (Figure 3).

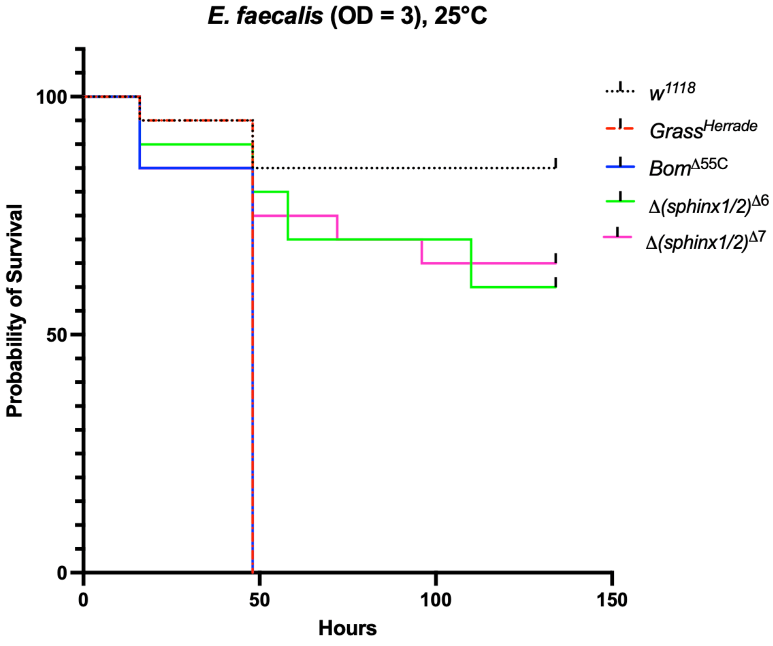

**Figure 2.** Survival *of Δ(sphinx1/2* male flies following septic infection with $OD_{600}$=5 *E. faecalis.* 20 flies were used per genotype per experiment; two independent experiments were performed for each treatment. Pooled data are shown.

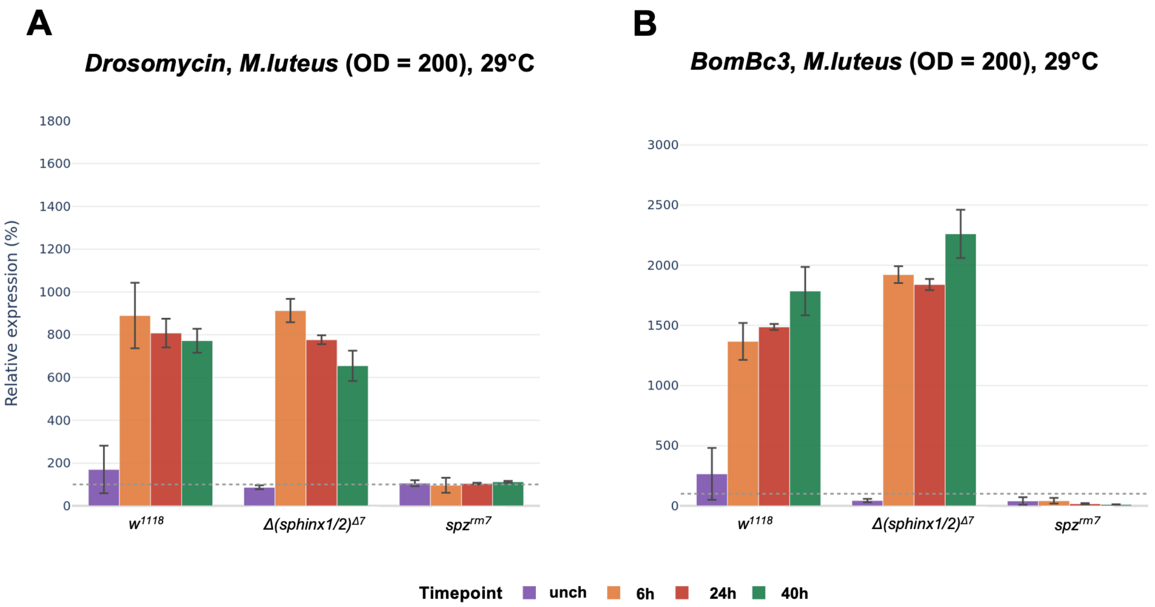

**Figure 3.** Expression of *Drosomycin* and *BomBc3* following septic injury with *M. luteus in* unchallenged (unch) male flies collected 6, 24 and 48 hours post-infection. Error bars represent the standard deviation of three independent experiments.

#### Conclusion

Based on our data and contrary to (Kambris et al., 2006), we conclude that *sphinx1/2* do not have a strong role in regulation of the Toll pathway.

### Supplementary S5: Role of wntD in Dorsal-mediated immunity

**Abstract**

(Gordon et al., 2005) found that *wntD* acts as a feedback inhibitor of *dorsal* but not *Dif* during the innate immune response, and that *wntD* mutants had elevated expression of *Diptericin* (*Dpt)* but not *Drosomycin* (*Drs)* in response to septic infection with *Micrococcus luteus*. However, *wntD* is only expressed at the embryonic stage and not at the adult stage at which the experiments were performed by (Gordon et al., 2005). Using an independent null mutant of *wntD (wntD(Gal4))*, we did not observe any consistent effect of *wntD* on expression of *Diptericin* and *Drosomycin*. Moreover, *wntD* null flies were not more sensitive to *Listeria monocytogenes* infection. Thus, we conclude that *wntD* is not a negative regulator of the Toll pathway during the immune response. Results published by (Gordon et al., 2005) may be explained by a genetic background effect independent of *wntD*.

**Introduction**

(Gordon et al., 2005) found that *wntD* acts as a feedback inhibitor of the Toll transcription factor *dorsal* but not of *dif* during the innate immune response, and that *wntD* mutants had elevated expression of the Imd target *Diptericin (Dpt)* but not the Toll target *Drosomycin (Drs)* in response to septic infection with *M. luteus*. They explain this pattern by citing the fact that constitutive Toll signaling upregulates *Dpt* (Lemaitre et al., 1996), and that *dorsal* can activate the promoter of *Dpt* but not *Drs* in cell culture (Reichhart et al., 1993). In addition, increased susceptibility to *Listeria monocytogenes* in *wntD* mutants was rescued in *wntD, dorsal* double mutants (Gordon et al., 2005), suggesting that mortality in *wntD* mutants is due to overactivation of *dorsal.*

The observation that *wntD* negatively impacts *Dpt* expression but not *Drs* upon systemic infection by the Gram-positive bacterium *Micrococcus luteus* is perplexing: *Dpt* acts as a read-out of Imd pathway activation, whereas *Drs* is a well-established read-out of the *M. luteus*-elicited Toll pathway (Lemaitre et al., 1995, 1996). The role of *wntD* in the immune response has never been further assessed since the initial 2005 results.

Here we have used an independent null mutant of *wntD* (*wntD(Gal4))* in an alternate genetic background to assess the role of this secreted protein in the activation of Toll and Imd pathways.

**Methods**

*Fly lines and bacterial strains*

*wntD(Gal4)* (TI{GAL4}wntD[GAL4], BDSC #94895) and *yw; wntD(KO1)* ($y^1$ $w^*$; TI{TI}$wntD^{KO1}$, BDSC #38407) were obtained from Bloomington. For survival experiments and assessment of transcript levels in response to systemic infections, *yw* and $w^{1118}$ (BDSC #3605) flies were used as wild-type controls. $Spz^{rm7}$ (Neyen et al., 2014) or $Bom^{\Delta 55C}$ *Drosdel* (Clemmons et al., 2015; Hanson et al., 2019) flies were used as immune deficient mutants to control for susceptibility. The strains *L. monocytogenesis* (CIP 107776, EGDe), *Pectobacterium carotovorum Ecc15* and *M. luteus* JL (Jean Lambert collection, Strasbourg) were used in this study

*Survivals*

5 to 7-day old male flies reared on cornmeal fly medium (6% cornmeal, 6% yeast, 0.6% agar, 0.1% fruit juice, 10.6 g/L moldex and 4.9 mL/L propionic acid) at 25°C were anaesthetized with $CO_2$ and pricked on the side of the thorax anterior and ventral to the wing joint with a 0.2 mm tungsten needle dipped in a $OD_{600}$=0.5 solution of *L. monocytogenes*. Bacteria were grown overnight in BHI, shaking at 29°C, and diluted to the desired OD in PBS. Survivals were incubated at 25°C and flies were transferred to fresh vials every 2-3 days. ~20 flies were used per genotype per experiment; three independent experiments were performed (n=3). Survival was assessed daily.

*qPCR for assessment of Toll and Imd pathway activation*

5-7-day old male flies were pricked with *L. monocytogenes* ($OD_{600}$=0.1), *M. luteus* ($OD_{600}$=200) or *Ecc15* ($OD_{600}$=200) and were maintained on cornmeal fly medium at 25°C before freezing 5-7 flies per treatment at -80°C at 4h, 8h, and 24h post infection (hpi). *M. luteus* and *Ecc15* were grown overnight in LB, shaking at 29°C, and diluted to the desired OD in PBS. *L. monocytogenes* was grown overnight shaking at 37°C in BHI. Uninfected flies were frozen at timepoint 0h as unchallenged control (UC). Two independent experiments were performed for *L. monocytogenes* (n=2). Experiments with *M. luteus* (n=1) and *Ecc15* (n=1) were only performed once. Total RNA was extracted from samples using TRIzol reagent (Invitrogen) and re-suspended in MilliQ water. Reverse transcription was performed using 0.5 mg total RNA in 10 µl reactions using Prime-Script RT (Takara) with random hexamer and oligo dT primers and diluted in MiliQ water to a final concentration of 2,5 ng/µl. Quantitative PCR was performed on a LightCycler 480 (Roche) in 96-well plates using PowerUp SYBR Green Master Mix (Applied Biosystems). Gene expression levels were normalized to the expression of the ribosomal protein *rp49* (*Rpl32*) and set relative to unchallenged (UC) controls.

*WntD* expression in *wntD* mutants and controls was assessed after RNA extraction from 4-8h old embryos. Embryos were collected on fruit juice agar plates and per sample 1/3 of the embryos on the plate were pooled and manually crushed in a tube before continuing with RNA extraction using TRIzol as described above.

- *Rp49* Forward: 5' – GCC GCT TCA AGG GAC AGT ATC TG – 3'
- *Rp49* Reverse: 5' – AAA CGC GGT TCT GCA TGA G – 3'
- *Drosomycin* Forward: 5' – CGT GAG AAC CTT TTC CAA TAT GAT G – 3'
- *Drosomycin* Reverse: 5' – TCC CAG GAC CAC CAG CAT – 3'
- *Diptericin A* Forward: 5' – GCT GCG CAA TCG CTT CTA CT – 3'
- *Diptericin A* Reverse: 5' – TGG TGG AGT GGG CTT CAT G – 3'
- *Bom55C* Forward: 5' – CTG ATC GGC GCT CAT CCC AG – 3'
- *Bom55C* Reverse: 5' – GGG ATG AGG AGA AGC TGC GG – 3'
- *wntD* Forward: 5' – TAT GTG GCG GCC ATT TCC AT – 3'
- *wntD* Reverse: 5' – GTG GGC ACA GGG TAC ATT CA – 3'

**Results**

*WntD is not expressed in adult flies*

We analyzed the gene profile of *wntD* using public databases. High-throughput expression data (modENCODE, FlyAtlas) data reveal that *wntD* is expressed only at the embryonic stage. Using RT-qPCR, we confirmed that *wntD* is not or only very weakly expressed in L3 larvae and adult stage (Ct values not detectable or > Ct 30). We confirm that *wntD* transcript levels are easily detectable in 4-8h old embryos, consistent with information from FlyBase (Figure 1A). RT-qPCR using RNA extracted from 4-8h old embryos confirmed that *wntD* transcripts were not detectable in the *wntD(Gal4)* null mutant, but surprisingly *wntD* transcript levels in the *wntD(KO1)* line used by (Gordon et al., 2005) were not reduced compared to wild-type (Figure 1A). This may indicate that the original mutation was lost during passaging of the fly line, as the *wntD(KO1)* line was originally confirmed to be a protein null by Southern and Western blot (Gordon et al., 2005).

*WntD flies do not show increased susceptibility to infection with* Listeria monocytogenes*.*

We then analyzed the survival rate at 25°C of *wntD* mutants and wild type flies upon systemic infection with *L. monocytogenes* ($OD_{600}$=0.1). In this experiment we included *spz*$^{rm7}$ and *Bom*$^{\Delta 55C}$ mutants as susceptible controls, and two different wild type genotypes: *w*$^{1118}$and *yw* reflecting the backgrounds of the two *wntD* mutant lines. As expected, *Bom* and *spz* mutants were significantly more susceptible than wild type (Figure 1B). However, the two *wntD* mutants did not have similar survival phenotypes. A susceptibility of *wntD* mutants compared to *yw* controls similar to that of (Gordon et al., 2005) could only be obtained when comparing *yw* wild type flies with *yw; wntD(KO1)*, which suggests this susceptibility is due to a background effect independent of *wntD,* particularly given that the expected mutation seems to have been lost. In contrast, *w*$^{1118}$ wild type flies and the independent *wntD(Gal4)* line were not as susceptible to this infection (Figure 1C). This reveals that the susceptibility of the *yw; wntD(KO1)* line as found by (Gordon et al., 2005) is likely due to genetic background.

*wntD flies show wild-type Imd and Toll pathway activity.*

We then assessed the functionality of Toll and Imd pathway activation in *wntD* flies. For this, we monitored the expression levels of *DptA*, a read-out of the Imd pathway, upon infection with *Pectobacterium carotovorum Ecc15* ($OD_{600}$=200) and *Drs* expression, a read-out of the Toll pathway, upon systemic infection with *M. luteus* ($OD_{600}$=200) and *L. monocytogenes* ($OD_{600}$=0.1) (Figure 2 A, B and C respectively). *WntD* flies did not show consistent changes in activation of Toll or IMD pathways compared to wild type flies in response to any of these systemic infections. An exception to this is the 24h time point after infection with *M. luteus,* where both *wntD* alleles had increased expression of *Drs* compared to wild-type. However, a similar effect did not occur with *L. monocytogenes*. As found by (Gordon et al., 2005), we found that *DptA* transcript levels were elevated in *yw; wntD(KO1)* flies in response to both *M. luteus* and *L. monocytogenes.* However, the independent *wntD(Gal4)* line did not show a similar effect. Thus, it is unlikely that *wntD* acts as a feedback inhibitor of *dorsal* (even given the (Gordon et al., 2005) interpretation that *dorsal* regulates *Dpt* which is typically seen as an Imd target, rather than *Drs* which is a classical Toll target), and instead elevated *Dpt* in the *yw; wntD(KO1)* line is likely due to genetic background. Although *Drs* was upregulated in both *wntD* lines at the 24h time point following *M. luteus* infection, a role of *wntD* as a feedback inhibitor of the Toll response would imply elevated *Drs* expression across all time points and in response to all infections. We therefore conclude that *wntD* does not act as a feedback inhibitor of *dorsal* or Toll during the immune response, and that similar to effects on survival, the effects on immune gene expression observed by (Gordon et al., 2005) were due to genetic background.

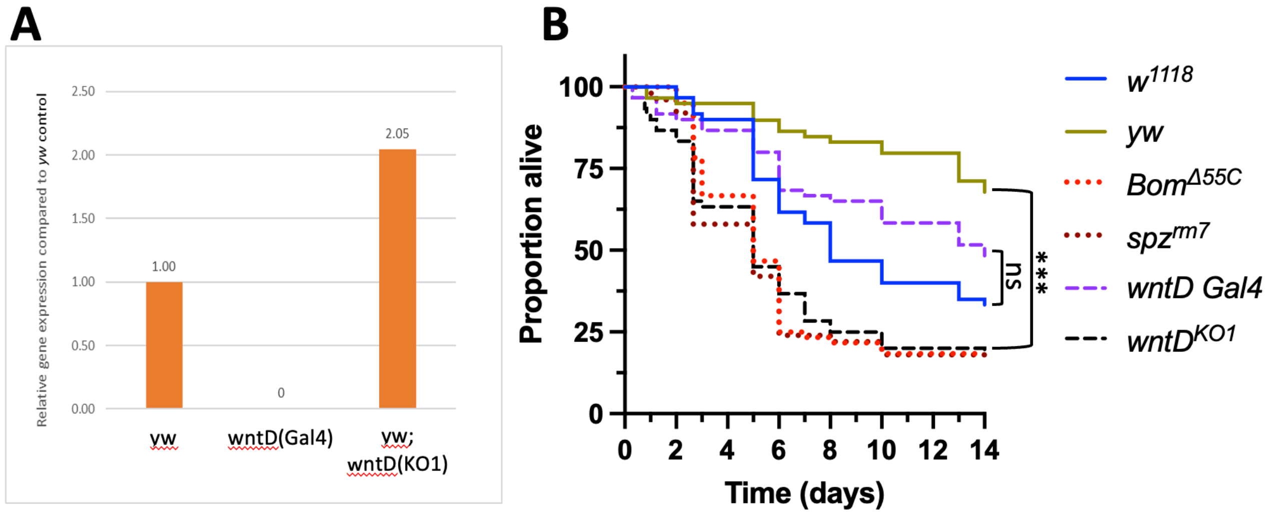

**Figure 1. (A**) Relative gene expression of *wntD* in 4-8h embryos confirms that *wntD(Gal4*) is a null mutant, but suggest *wntD(KO1)* is not (n=1). **(B)** Survival plot displaying survival probability over time (days) after systemic infection with *Listeria monocytogenes* (n=3). Significance calculated using a Cox proportional hazard test. ns, not significant; ***, P < 0.001. 20 flies were used per genotype per experiment; three independent experiments were performed.

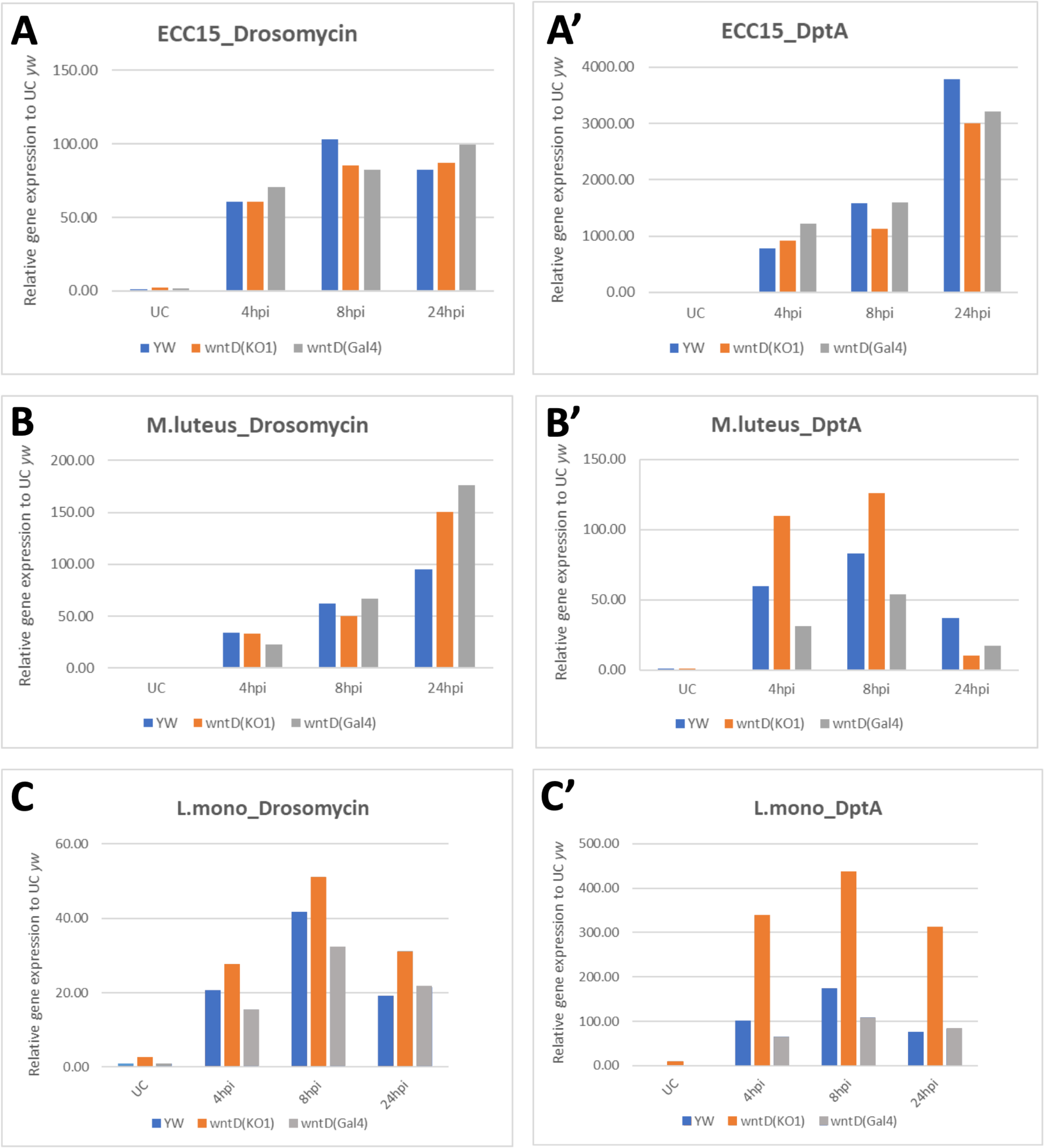

**Figure 2.** Induction of *Drosomycin* and *Diptericin* as read-outs of Toll and Imd pathway activation, respectively. Transcript levels were measured at different time points after systemic infection with *Ecc15* (**A**, $n^{exp}$=1), *M. luteus* (**B**, $n^{exp}$=1) and *L. monocytogenes* (**C**, $n^{exp}$=1,). Increased transcript levels of *Diptericin A* (DptA) in *yw,wntD(KO1)* do not occur in the second *wntD* mutant *wntD(Gal4)*. Relative gene expression is normalized to *rp49* and set relative to respective unchallenged (UC) control. hpi=hours post infection

#### Conclusion

We were unable to reproduce the key findings of (Gordon et al., 2005) using an independent *wntD(Gal4)* null allele. Given that we reproduced susceptibility using the original *yw; wntD(KO1)* line even despite apparent loss of the expected mutation of *wntD*, we conclude that immune effects observed by (Gordon et al., 2005) were due to genetic background. Systemic infection with *Ecc15*, *M. luteus* and *L. monocytogenes* did not reveal any consistent effects of an independent *wntD(Gal4)* mutation on the expression of *Diptericin* and *Drosomycin*, or on survival to systemic *L. monocytogenes* infection. Collectively, we conclude that *wntD* is not a negative regulator of the Toll pathway during the immune response. This is consistent with data showing that *wntD* is only expressed at the embryonic stage, and not at the adult stage where immune effects were tested by (Gordon et al., 2005). The role of *wntD* in the regulation of the Toll pathway during development has been confirmed by other studies (Rahimi et al., 2016), but in the absence of additional evidence, we conclude that *wntD* does not have a role in the inducible immune response.

### Supplementary S6: Activation of Toll by overexpression of GNBP3

**Abstract**

(Gottar et al., 2006) found that overexpression of *GNBP3* using an *hsp-Gal4* driver induced *Drosomycin* to a similar level as *Micrococcus luteus* infection. Experiments using two *UAS-GNBP3* constructs and a variety of drivers suggests that contrary to this report, overexpression of *GNBP3* is not an effective method to activate Toll signaling.

**Introduction**

Ectopic activation of immune pathways is frequently achieved through overexpression of core immune proteins. This technique is commonly used in epistasis studies. However, many proteins including most serine proteases and receptors require activation by cleavage or by ligand binding, respectively, and therefore overexpression of these is not an effective method of stimulating immune signaling.

GNBP3 (Gram Negative Binding Protein 3) is a core immune protein required for activation of Toll signaling by fungi, but not bacteria (Buchon et al., 2009; Gottar et al., 2006). GNBP3 is considered to be a pattern-recognition receptor that binds long, linear fungal beta-1,3-glucans (such as curdlan and laminaran/laminarin) but not branched beta-1,3-glucans or beta-1,6-glucans (Mishima et al., 2009), and is required for both expression of antimicrobial peptides and activation of melanization (Gottar et al., 2006; Matskevich et al., 2010). It is primarily secreted by the fat body (Gottar et al., 2006; Mishima et al., 2009) but also by epithelia including the tracheae upon natural infection of larvae (Wagner et al., 2009, 2008). In contrast, GNBP3 is not expressed in hemocytes (Tattikota et al., 2020), and is not upregulated upon infection of adult flies (Kim et al., 2000) or only at late time points (De Gregorio et al., 2001).

(Gottar et al., 2006) found that overexpression of *GNBP3* using an *hsp-Gal4* driver induced *Drosomycin* to a similar level as *Micrococcus luteus* infection at 24hpi, about 100x the baseline *Drs* expression of unchallenged wild-type flies. In contrast, (Buchon et al., 2009) were not able to reproducibly induce *Drs* by overexpression of *GNBP3* using a *c564-Gal4* driver. In only one experiment of several was *Drs* induced upon *GNBP3* overexpression to 10% of the level reached in wild-type flies collected 16h after infection with *M. luteus* (see Buchon et al., 2009 - Figure 4). *GNBP3* is not a strongly immune-responsive gene, and is not induced by septic bacterial or natural fungal infection until mild upregulation late in infection (48h) (De Gregorio et al., 2001). GNBP3 is strongly expressed in the fat body compared to other tissues (FlyAtlas2), so use of the *c564-Gal4* driver by (Buchon et al., 2009) in an attempt to replicate the phenotype achieved by (Gottar et al., 2006) using a ubiquitous driver was reasonable. (El Chamy et al., 2008; Fullaondo et al., 2011) also report activation of Toll through overexpression of *GNBP3* using the same UAS construct as (Gottar et al., 2006). (El Chamy et al., 2008) characterize this activation as ‘weak but detectable’ using Northern blot (see El Chamy et al., 2008 - Figure 3), and found that mutation of *grass* eliminated Toll activation by *GNBP3* overexpression (*hsp-Gal4,* ubiquitous). (Fullaondo et al., 2011) found that overexpression of this construct (*YP-Gal4* driver, fat body) activated a Drs-GFP reporter in adult flies, and that this activation was reduced by concurrent overexpression of the serpin *Spn1* (*Spn42Dd*).

Given these inconsistencies, we aimed to determine whether overexpression of *GNBP3* is a reliable method of strongly inducing Toll signaling by overexpressing two *UAS-GNBP3* constructs using a variety of drivers.

**Methods**

*Fly lines and bacterial strains*

Flies were reared at 25°C on standard cornmeal fly medium (6% cornmeal, 6% yeast, 0.6% agar, 0.1% fruit juice, 10.6 g/L moldex and 4.9 mL/L propionic acid), and flipped to new medium every 2-3 days.

- *$w^{1118}$* wild type (Bloomington #3605)
- *c564-Gal4* (II, homozygous)
- *hs-Gal4/CyO* (BDSC #2077)

- *UAS-GNBP3* Lemaitre (III, homozygous, from Dominique Ferrandon), referred to below as *UAS-GNBP3(L)*
- *UAS-GNBP3* (Gottar et al., 2006) (II, homozygous), referred to below as *UAS-GNBP3(G)*

The strains *M. luteus* JL (Jean Lambert collection, Strasbourg) was used in this study.

*Crosses and gene induction*

Drivers were checked for efficacy by crossing to a *UAS-GFP/CyO* line and verifying expression in appropriate tissues using larvae and adults of both sexes. Stocks were maintained on cornmeal fly medium at 25°C; crosses were maintained at 29°C for optimal expression of the UAS constructs. Flies were aged to 3-6 days. Adult emergents bearing the *hs-Gal4* driver as well as appropriate controls were heat shocked as follows: 20 min at 37°C, 20 min at 18°C, 20 min at 37°C, and then 24 h incubation at 29°C. Concurrent with heat shocks, wild type flies were pricked in the thorax with a 0.2 mm tungsten needle dipped in a concentrated overnight culture of *M. luteus* grown shaking at 29°C in LB and diluted to $OD_{600}$=200 in PBS. 24h after heat shock, groups of 8-10 male flies per treatment were frozen at -80°C.

*RNA extraction and qPCR*

Total RNA was extracted from pooled samples using TRIzol reagent (Invitrogen), and re-suspended in MilliQ water. DNase treatment was performed using TURBO DNA-free kit (Invitrogen). Reverse transcription was performed using 0.5 mg total RNA in 10 µl reactions using Prime-Script RT (Takara) with random hexamer and oligo dT primers. Quantitative PCR was performed on a LightCycler 480 (Roche) in 96-well plates using PowerUp SYBR Green Master Mix (Applied Biosystems). Expression of *Drosomycin* and *GNBP3* was normalized to expression of the ribosomal protein *rp49* (*Rpl32*). At least two biological replicates were performed per sample.

- *Rp49* Forward: 5' – GCC GCT TCA AGG GAC AGT ATC TG – 3'
- *Rp49* Reverse: 5' – AAA CGC GGT TCT GCA TGA G – 3'
- *Drosomycin* Forward: 5' – CGT GAG AAC CTT TTC CAA TAT GAT G – 3'
- *Drosomycin* Reverse: 5' – TCC CAG GAC CAC CAG CAT – 3'

#### Results and Discussion

As shown in Figure 1, the *hs-Gal4* driver similar to that used by (Gottar et al., 2006) induced *GNBP3* to a higher level than the *c564-Gal4* used by (Buchon et al., 2009) even in the absence of heat shock. The *UAS-GNBP3(L)* construct produced a higher level of overexpression regardless of driver. In our hands and similar to the findings of (Buchon et al., 2009), no combination of driver and UAS construct appreciably upregulated *Drosomycin* expression. The *hs-Gal4>UAS-GNBP3* treatment produced extremely high levels of *GNBP3* (~130x over unchallenged flies, ~50x septic injury) when heat shocked, but this treatment only induced *Drosomycin* to 10% of the level achieved by *M. luteus* septic injury. This is contrary to (Gottar et al., 2006) who found that this treatment induced *Drosomycin* to levels comparable to those achieved with *M. luteus.*

A possible explanation for the upregulation observed by (Gottar et al., 2006) upon overexpression of *GNBP3* is that these authors is a difference in baseline expression of *Drs* between lines used in the comparison, which were in different genetic backgrounds. Alternatively, since the endogenous immune response is not accompanied by upregulation of *GNBP3* (De Gregorio et al., 2001), activation of *Drs* due to overexpression of this gene is unlikely to mimic the conditions of a natural immune response. Toll activation in overexpression lines may be an indirect effect due to damage caused by overabundance or mislocalization of a protein normally expressed at low levels. In support of this last interpretation, we also attempted to cross both constructs to a *Da-gal4* driver to obtain information using an independent ubiquitous driver, and found that *Da>UAS-GNBP3* escapers died very quickly after eclosion and before cuticle hardening.

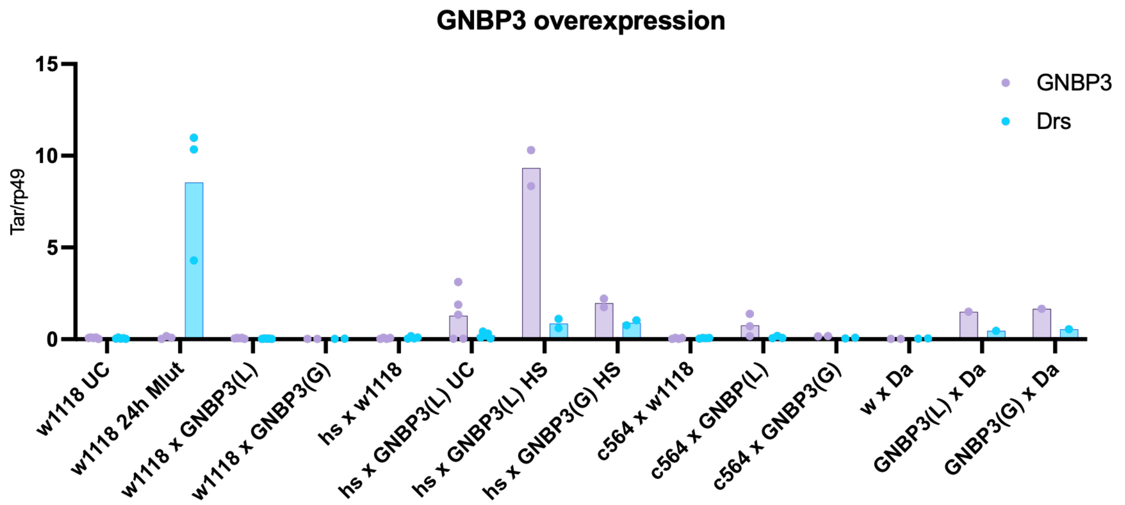

**Figure 1.** Data show average of two biological replicates of GNBP3(L) overexpression, plus one biological replicate including two technical replicates for treatments using the GNBP3(G) construct. Expression of the target gene (*GNBP3, Drs*) is normalized to expression of the housekeeping gene *rp49* (*rpL32*) for respective treatments. Extractions were performed 24h after treatment. All treatments are unchallenged unless otherwise noted: HS=24h after heat shock, Mlut=24h after septic infection with *M. luteus.*

#### Conclusion

Overexpression of *GNBP3* either ubiquitously or in the fat body does not effectively activate the Toll pathway. At most, we find similar to (El Chamy et al., 2008) that overexpression may produce 'weak but detectable' activation of Toll. Use of two *UAS-GNBP3* constructs yielded similar results. It would be interesting to determine whether the weak induction of *Drs* upon over-expression of *GNBP3* also occurs in germ-free flies, or whether it is dependent on the presence of environmental microbes.

### Supplementary S7: Activation of Toll by overexpression of *grass*.

**Abstract**

(El Chamy et al., 2008) found that overexpression of *grass* using *hsp-Gal4* induced the Toll pathway in adults. Using an independent construct, we find that contrary to (El Chamy et al., 2008), overexpression of *grass* in the absence of immune challenge does not strongly activate the Toll pathway.

**Introduction**

Ectopic activation of immune pathways is frequently achieved through overexpression of core immune proteins. This technique is commonly used in epistasis studies. However, many proteins such as serine proteases and receptors require activation by cleavage or a secondary factor, and therefore overexpression of these is not an effective method of stimulating immune signaling. Grass is a serine protease in the Toll pathway that requires cleavage to take its active form. Using Northern blots, (El Chamy et al., 2008) found that overexpression of *grass* using a UAS construct under control of the ubiquitous temperature-activated *hsp-Gal4* driver induced the Toll pathway. The authors have confirmed that the construct used in this study was a full-length *grass* construct (Laure El Chamy, personal communication). To confirm whether overexpression of uncleaved Grass is an effective method of activating the Toll pathway, we repeated these experiments using an independent full-length UAS-Grass construct.

**Methods**

*Fly lines and husbandry*

A full-length *UAS-Grass/CyO* (II) construct from Strasbourg was backcrossed to the $w^{iso}$ background to remove balancers on the third chromosome. The *UAS-Grass* was then crossed to a ubiquitous driver (*Da-Gal4/TM6C,* Lemaitre lab collection), a heat-shock inducible ubiquitous driver (*hs-Gal4,* BDSC #2077) and fat body driver (*c564-Gal4,* BDSC #6982). Drivers were checked for efficacy by simultaneously crossing to a *UAS-GFP/CyO* line, and expression in appropriate tissues was verified using larvae and adults of both sexes. Stocks were maintained on cornmeal fly medium (6% cornmeal, 6% yeast, 0.6% agar, 0.1% fruit juice, 10.6 g/L moldex and 4.9 mL/L propionic acid) at 25°C; crosses were maintained at 29°C for optimal expression of UAS constructs. Flies were aged to 3-6 days. Adults bearing the *hs-Gal4* driver as well as appropriate controls were heat shocked as follows: 20 min at 37°C, 20 min at 18°C, 20 min at 37°C, and then 24 hr incubation at 29°C. 24h after heat shock, groups of 8-10 male flies per treatment were frozen at -80°C. Flies carrying the heat shock driver were also left unshocked for comparison. $w^{1118}$ flies (Bloomington #3605) and appropriate crosses of this wild-type to the driver and UAS construct were used as controls. 24h after heat shock, groups of 8-10 male flies per treatment were frozen at -80°C. For septic infection treatments, flies were pricked in the thorax with a 0.2 mm tungsten needle dipped in a concentrated overnight culture of *Micrococcus luteus* JL (Jean Lambert collection, Strasbourg) grown at 29°C overnight in LB and diluted to $OD_{600}$=200 in PBS. If applicable, heat shock was performed concurrent with infection.

*qPCR*

Total RNA was extracted from samples using TRIzol reagent (Invitrogen), and re-suspended in MilliQ water. DNase treatment was performed using TURBO DNA-free kit (Invitrogen). Reverse transcription was performed using 0.5 mg total RNA in 10 µl reactions using Prime-Script RT (Takara) with random hexamer and oligo dT primers. Quantitative PCR was performed on a LightCycler 480 (Roche) in 96-well plates using PowerUp SYBR Green Master Mix (Applied Biosystems). *Drosomycin* and *grass* expression was normalized to expression of the ribosomal protein *rp49.* For graphical display, *Drosomycin* and *Grass* expression levels were normalized to the unchallenged (UC) condition in the wild type $w^{1118}$ flies. Two independent experiments were performed with similar results.

- *Drosomycin* Forward: 5' – CGT GAG AAC CTT TTC CAA TAT GAT – 3'

- *Drosomycin* Reverse: 5’ – TCC CAG GAC CAC CAG CAT – 3’
- *Grass* Forward: 5’ – GCC AGT TGT GAA TGT GGG CAT TG – 3’
- *Grass* Reverse: 5’ – GGC AAA TGG GCT TGA TGT GCT TC – 3’
- *Rp49* Forward: 5’ – GCC GCT TCA AGG GAC AGT ATC TG – 3’
- *Rp49* Reverse: 5’ – AAA CGC GGT TCT GCA TGA G – 3’

#### Results

*Grass* expression was low in wild type flies. We successfully overexpressed *grass* to ~20x wild-type levels using the *hs-Gal4* driver with or without heat shock treatment (Figure 1). The *c564-Gal4* fat body driver and *Da-Gal4* drivers were also successful in overexpressing *grass*. However, in our hands there was no correlation between high *Grass* expression levels and Toll activation as determined by *Drosomycin* expression. The *Da-Gal4>UAS-grass* lines with *grass* expression over 100x higher than wild-type expression had lower *Drosomycin* expression than wild type ($w^{1118}$) flies where *grass* expression was barely detectable. Furthermore, even when the *hs-Gal4>UAS-grass* flies were challenged with *M. luteus* to activate Toll signaling, *Drosomycin* expression was lower than in challenged wild-type flies.

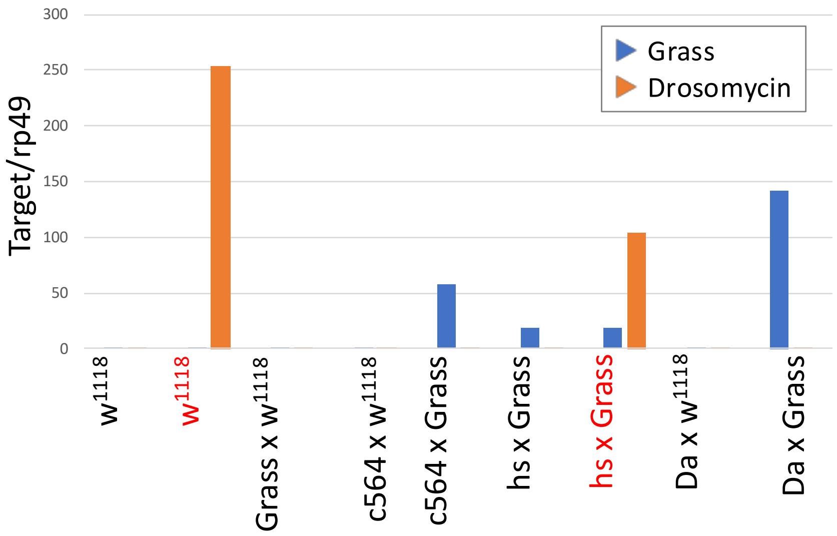

**Figure 1.** Levels of *grass* and *Drosomycin* expression (*target/rp49*) normalized to expression in the unchallenged wild-type line. Treatments in red are challenged with *M. luteus* (24h); all other treatments are unchallenged.

#### Conclusion

Our results suggest that an excess of Grass is not sufficient to strongly activate Toll signaling, even in the presence of an immune stimulus. A lack of Toll activation is consistent with data showing that *grass* is only mildly upregulated following infection (De Gregorio et al., 2001). This is also consistent with its structure as a CLIP domain serine protease, which typically requires cleavage to become active. We conclude that contrary to (El Chamy et al., 2008), overexpression of full-length *grass* does not activate the Toll pathway.

### Supplementary S8: PGRP-LE is not secreted in the hemolymph

**Abstract**

A previously published result from an overexpression study indicated that a truncated form of the intracellular PGRP-LE protein is secreted to the larval hemolymph and can act in a non-cell autonomous manner (Takehana et al., 2004). However, in the absence of overexpression, no studies have found an extracellular role for PGRP-LE. In order to determine whether PGRP-LE is endogenously present in the hemolymph of larvae, we performed Western blots of larvae expressing a PGRP-LE::GFP fusion protein under its native promoter sequences. Our results suggest that in larvae, PGRP-LE is exclusively intracellular in the absence of overexpression.

**Introduction**

PGRP-LE is a pattern-recognition receptor upstream of the Imd pathway that acts in parallel to PGRP-LC. PGRP-LE lacks a signal peptide and is the only known soluble intracellular pattern-recognition receptor in *Drosophila*. It activates the Imd pathway in response to monomeric peptidoglycan fragments (Bosco-Drayon et al., 2012; Kaneko et al., 2006). (Takehana et al., 2004) characterized features of PGRP-LE in wild-type *Drosophila* and a *PGRP-LE*$^{112}$ mutant line that removes the start codon of PGRP-LE. Endogenous *PGRP-LE* produces two transcripts but a single polypeptide; one transcript has a longer 3' UTR and is expressed only in adult neural tissue (see FlyAtlas2 and FlyBase).

Based on Western blots from wild-type and *PGRP-LE*$^{112}$ mutant larvae, (Takehana et al., 2004) concluded that in addition to a long (45 kDa) intracellular form of PGRP-LE (in the tracheae, hemocytes and fat body), the larval hemolymph contained small amounts of additional short (21.5 and 30 kDa) secreted forms of PGRP-LE consisting of the PGRP domain. They surmised these shorter products were proteolysis products of the long form. Immunostaining further indicated that PGRP-LE was present at the lumenal surface of the tracheae, which would require secretion (Takehana et al., 2004). They also found that overexpression of *PGRP-LE* in larval tracheae or fat body cells had non-cell autonomous effects on activation of antimicrobial peptides in adjacent cells.

In a follow-up paper, (Kaneko et al., 2006) found that when overexpressed in S2 cells, full-length PGRP-LE was not present in the culture medium. Staining indicated that both endogenous PGRP-LE and tagged V5-PGRP-LE were exclusively intracellular and cell-autonomous in the Malpighian tubules and hemocytes (Kaneko et al., 2006). However, when a truncated construct of PGRP-LE was expressed in S2 cells (corresponding to the 30 kDa form found in the hemolymph by Takehana et al., 2004), it was present in the culture medium and enhanced the *Dpt* response to TCT (Kaneko et al., 2006). PGRP-LE lacks a signal peptide, making it unclear how a truncated form could be secreted in cell culture medium and, most importantly, in hemolymph *in vivo*. Although alternative mechanisms exist for protein secretion (e.g. (Shibata et al., 2017)), such a mechanism is counterintuitive for PGRP-LE as the many other PGRP homologs present in the hemolymph (PGRPs SB1, SB2, SC1a/b, SD, and LB) all retain a signal peptide (Werner et al., 2000). It is unclear whether a short isoform of PGRP-LE exists in the larval hemolymph endogenously.

A methodological concern with (Takehana et al., 2004) is worth noting: the antibody used to detect PGRP-LE cross-reacted with at least three other PGRPs. These were excluded by pre-absorbing the antibody in *PGRP-LE*$^{112}$ mutants. However, pre-absorption may not have been fully effective at eliminating cross-reactivity to other PGRPs in the unspecified wild-type line. This is evidenced by the fact that a nonspecific 21.5 kDa protein was tagged by the pre-absorbed PGRP-LE antibody even in *PGRP-LE*$^{112}$ larvae. Many secreted PGRPs known to be present in the hemolymph have a predicted molecular weight close to 20-30 kDa, including PGRPs SB1, SB2, SC1a/b, SD, and the cleaved extracellular domain of PGRP-LC (23.5 kDa).

(Abdelsadik and Roeder, 2010) also activated the Imd pathway using overexpression of *PGRP-LE* in the larval salivary glands. They similarly found that overexpression of *PGRP-LE* had non-cell autonomous effects throughout the digestive system. Note that the salivary glands are a specialized secretory tissue designed for atypical export of large amounts of intracellular cargo, including immune molecules (see (Beňová-Liszeková et al., 2021)). (Fink et al., 2016) show a similar non-cell autonomous effect of PGRP-LE upon overexpression in enterocytes in the midgut of adult flies (NP1-Gal4).

Although many studies indicate that endogenous PGRP-LE is active in the cytoplasm of larval and adult *Drosophila* (Libert et al., 2006; Neyen et al., 2012; Yano et al., 2008), non-cell autonomous effects have never been reported for PGRP-LE in a context that excludes overexpression of the protein in specialized secretory tissues. An adult hemolymph proteome (Rommelaere et al., 2025) shows that PGRP-LE is either absent in hemolymph (immune challenged or not) or so weakly expressed as to be functionally absent. (Libert et al., 2006) used the same *UAS-PGRP-LE* construct as (Takehana et al., 2004) to overexpress this protein in the fat body but found that effects were cell-autonomous in adult flies. (Libert et al., 2006) qualify that their results may contradict those of (Kaneko et al., 2006; Takehana et al., 2004) due to a difference between adult and larval Drosophila. A process unique to the larval stage is that some tissues dissociate during the late L3 larval stage in preparation for metamorphosis, which may release normally intracellular proteins to the hemolymph. One of these tissues is the salivary gland, a region where PGRP-LE is strongly expressed in the larva (FlyAtlas). This could in theory result in a transient period in larval development where PGRP-LE is present in the hemolymph due to tissue dissociation.

To clarify whether PGRP-LE is present in the hemolymph of larvae in the absence of overexpression, we performed Western blots on protein extracts from larvae expressing a PGRP-LE::GFP fusion protein under its native promoter sequences.

**Methods**

*Fly lines and husbandry*

The *PGRP-LE::GFP* line (Bosco-Drayon et al., 2012) was a gift from Bernard Charroux. Balancers and additional mutations present in the original line were removed by recombination with the $w^{iso}$ *Drosdel* background. The $w^{1118}$ line (Bloomington #3605) was used as a wild-type control. Controls expressing a GFP tag in the hemocytes (*w;*$hml^{D}$*-Gal4, UAS-GFP,* courtesy of B. Mattey-Prevot) and GFP secreted from the larval fat body ($TubGal80^{TS}$*, UAS-GFP; Lpp-Gal4/TM3* (Brankatschk et al., 2018) crossed to *y,w*;;P[UASp4sec::GFP]/TM3ser* (Fabrowski et al., 2013)). Flies were maintained at 25°C on cornmeal fly medium (6% cornmeal, 6% yeast, 0.6% agar, 0.1% fruit juice, 10.6 g/L moldex and 4.9 mL/L propionic acid), and flipped to new vials every 2-3 days. After egg laying, vials for all genotypes were transferred to 29°C to promote Gal4-UAS expression.

*DNA extraction and PCR*

Active transcription of the *PGRP-LE::GFP* fusion construct was verified by PCR. DNA was extracted by macerating single flies in 50 µL squishing buffer (10 mM Tris-HCl pH 8.2, 1 mM EDTA, 25 mM NaCl and 200 µg/ml Proteinase K) followed by incubation at 37°C for 30 minutes, and 2 minutes at 95°C. PCR was performed in a 25 µL reaction volume using AccuStart II PCR SuperMix (QuantaBio) using the conditions specified below. Bands were separated on a 1% agarose gel.

- *PGRP-LE::GFP* primer set:
  - Forward: 5'- CAG TGC AAT TCG ACG GAG AG -3'
  - Reverse: 5'- GAA CTC CAG CAG GAC CAT GT G -3'
  - Annealing temperature: 56°C
  - Extension time: 30s
  - The forward primer binds to the final exon of PGRP-LE; the reverse primer is contained within the eGFP tag.

*Larval oral infections*

For *PGRP-LE::GFP* oral infections, ~60-100 synchronized early L3 larvae were washed out of the food and stirred into 200 µL of $OD_{600}$=400 *Ecc15* grown in an overnight shaking culture (LB, 29°C) mixed with 400 µL of banana puree in an Eppendorf. The Eppendorf was plugged with foam to allow oxygen diffusion and left at room temperature for 30 minutes, then the contents were transferred to a vial containing cornmeal fly medium and incubated at 29°C.

*Extraction of protein samples*

L3 larvae just beginning to wander were selected for each genotype. 13-14 larvae were opened in 100 µL dissection buffer (sterile PBS, phenoloxidase inhibitor phenylthiourea (PTU) on ice three times, for a total of ~40 larvae per genotype. Carcasses (all remaining tissues including epithelium, cuticle, gut and fat body) were removed to the side. The buffer plus hemolymph and hemocytes was transferred to Lo-bind Eppendorfs on ice. Carcasses were placed in 300 µL RIPA (50mM Tris-HCl pH 7.5, 150 mM NaCl, 1% NP-40, 0.1% sodium deoxycholate, 0.1 % de SDS) buffer in a tube with glass beads and homogenized in a TissueLyzer at 10000 rpm for 30s at 4°C, and then placed on ice and agitated every 10 minutes for 30 minutes. Hemolymph samples were centrifuged 10 minutes at 300 g (4°C) to pellet the hemocytes, and 250 µL supernatant was removed to a new Lo-bind tube, taking care not to disturb the pellet. This purified hemolymph fraction was frozen at -20°C. 300 µL RIPA buffer was then added to the hemocyte pellet, and this sample was left on ice and agitated every 10 minutes for 30 minutes. At the end of the incubation, the carcass and hemocyte fractions were centrifuged at 12000 rpm for 10 minutes at 4°C. 250 µL supernatant was removed to a new Lo-bind tube from the hemocyte fraction to exclude the cell debris, and frozen at -20°C. 300 µL of homogenate was removed from the centrifuged carcass samples and placed in a new Lo-bind tube. This was centrifuged at 12000 rpm for 10 minutes at 4°C, and supernatant was removed to a new Lo-bind tube and frozen at -20°C for the carcass fraction.

*BCA protein quantification and Western blot*

Protein samples were thawed, vortexed briefly, spun down, and then kept on ice. Protein concentration in the samples was determined using a Pierce™ BCA Protein Assay Kit (ThermoFisher catalog #23225) according to the manufacturer's instructions. Samples were diluted to a concentration of 1µg/µL in dH2O and Blue buffer with 20mM DTT, heated for 10 minutes at 90°C, and cooled to 4°C. Samples were run on a precast 10% Tris-Glycine Novex gel (Invitrogen) at 90V with a PageRuler™ Prestained Protein Ladder (Thermo Scientific). Following protein transfer to a nitrocellulose membrane (Invitrogen iBlot), the membrane was blocked in 5% milk in PBST (PBS + 0.1% Tween), stained overnight with Roche™ mouse-anti-GFP primary antibody (1:1000) in milk + PBST, rinsed 3x in PBST, incubated 45 minutes in anti-mouse-HRP secondary antibody (1:10000, Abcam), and rinsed 3x in PBST. The signal was developed with Amersham ECL Western Blotting Detection Reagent and digitally imaged with Fusion Solo (Vilber Lourmat) and the presented images are inverted.

#### Results and Discussion

We attempted to determine whether PGRP-LE was detectable in larval hemolymph in the absence of overexpression. The *PGRP-LE::GFP* construct is tagged at the C-terminus following the PGRP domain, so even if the protein is cleaved, the truncated PGRP domain, the region reported to be present in the hemolymph, should retain the GFP tag. Microscopic inspection of the dissected gut of *PGRP-LE::GFP* larvae before and after oral infection with *Ecc15* revealed puncta of GFP fluorescence only after infection, as noted by (Bosco-Drayon et al., 2012) using the same protein construct. Preliminary Western blots of hemolymph and hemocyte extracts from unchallenged *PGRP-LE::GFP* L3 larvae failed to detect the fusion protein, so *PGRP-LE::GFP* larvae orally infected with *Ecc15* were included, and the remaining hemolymph- and hemocyte-free carcass (including fat body, gut, tracheae, etc.) was also assayed for expression of the fusion protein.

Full-length PGRP-LE was previously reported to have a molecular weight of ~45 kDa *in vivo* (Kaneko et al., 2006; Takehana et al., 2004). Based on sequence, *PGRP-LE* transcripts generate identical polypeptides with a predicted molecular weight of 39.4 kDa (FlyBase), predicted to have 5 N-linked glycosylations with predicted molecular weights of 1.4 to 2.7 kDa (Wang et al., 2017), resulting in a predicted total apparent weight of 46.4-52.9 kDa. As the GFP tag has a molecular weight of 26.8 kDa, the full-length fusion protein is predicted to have a weight of 73.8 kDa. Genotypes expressing secreted GFP from the fat body (*Lpp-Gal4>UAS-secGFP,* referred to below as *secGFP*) and intracellular GFP in the hemocytes (*hml-Gal4>UAS-GFP,* referred to below as *hmlGFP*) were also included as controls.

A ~25kDa protein corresponding to the GFP tag alone was detected in the control *secGFP* and *hmlGFP* samples. As expected, this band was stronger in the intracellular fraction from *hmlGFP* samples, and stronger in the hemolymph fraction from *secGFP* samples, indicating that the blot effectively showed the major location of the GFP tag (Figure 1). These bands were also detected in the *secGFP* and *hmlGFP* carcass samples, which retain some hemolymph and hemocytes. A slightly larger secondary band (between 25-35kDa) was detected primarily in the carcass extracts of the *secGFP* larvae, likely corresponding to the intracellular uncleaved secGFP protein retaining the signal peptide.

No bands were apparent in $w^{1118}$ or *PGRP-LE::GFP* (challenged or unchallenged) samples at moderate exposure times (30s). At a long exposure time (10 minutes), a faint band at ~55 kDa was visible only in the *PGRP-LE::GFP* challenged whole body sample (Figure 1, lower panels). This band likely represents the PGRP-LE::GFP fusion protein, whose expression increases following immune challenge. The fact that the size is smaller than the expected ~75 kDa may be due to degradation of the full-length protein construct (e.g. 26.8 kDa GFP tag + 30 kDa PGRP-LE fragment noted by (Takehana et al., 2004)). Small bands similar to the predicted size of the cleaved GFP tag are also detectable in the whole body *PGRP-LE::GFP* sample only at ~25 kDa.

These results show that the fusion protein was only detectable at a very low level in the whole-body sample following immune challenge, and was undetectable in both the hemolymph and hemocyte fractions of L3 larvae regardless of challenge. This indicates that PGRP-LE is not present at detectable levels in the larval hemolymph when under the control of its own promoter, and that the endogenous protein is likely also exclusively intracellular.

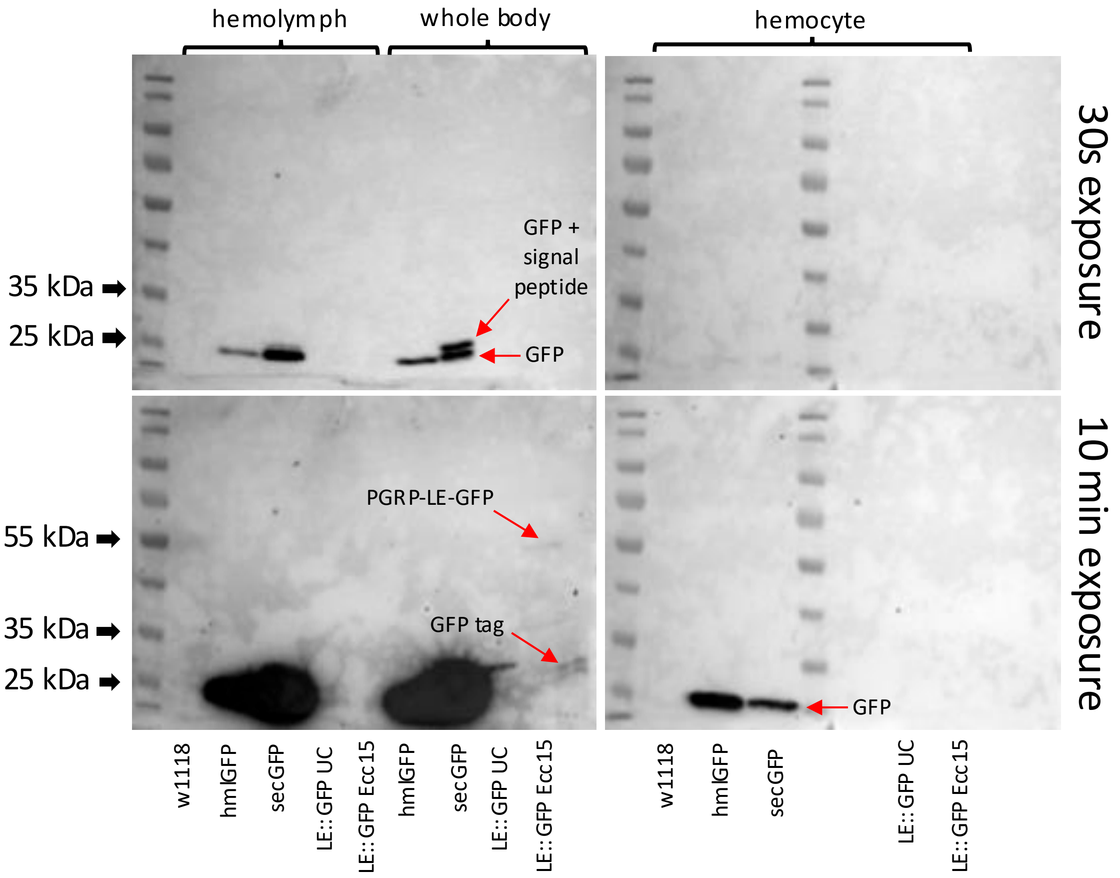

**Figure 1.** Western blot of hemolymph, hemocyte, and carcass fractions of L3 larvae. The same gel is shown at short (30s) and long (10 minute) exposures. The GFP tag is undetectable in all *PGRP-LE::GFP* samples except the *Pectobacterium carotovorum Ecc15* oral infection whole body sample, where very faint bands are visible as indicated by labels.

**Conclusion**

Data from other papers (Rommelaere et al, 2025, Libert et al., 2006) and our own data indicate that in both larvae and adult flies, endogenous PGRP-LE is not present in the hemolymph at detectable levels. In the absence of overexpression, it likely acts only in a cell-autonomous manner to promote immune activation in response to diffusible TCT (Bosco-Drayon et al., 2012). Reports of an extracellular version of PGRP-LE may be due to overexpression of the protein in specialized secretory tissues, or assaying of larvae at a stage in development where tissues have begun dissociating prior to metamorphosis.

### Supplementary S9: PGRP-LE is not required for infection-dependent melanization at the wound site

**Abstract**

Two studies performed within a single laboratory found involvement of the Imd pathway receptor PGRP-LE in melanization (Takehana et al., 2004, 2002). These results are counterintuitive as the Imd pathway is not typically involved in *Drosophila* melanization. Here, we used two independent loss-of-function mutants to assess the role of PGRP-LE in melanization. Our results indicate that PGRP-LE is not required for infection-dependent cuticular melanization at the wound. The melanization observed in response to PGRP-LE overexpression, as reported by (Takehana et al. 2004, 2002), may instead result from tissue damage caused by excessive immune activation.

**Introduction**

PGRP-LE is an intracellular pattern-recognition receptor of monomeric peptidoglycan (TCT) in the *Drosophila* Imd pathway, activation of which culminates in a local or systemic immune response including transcription of antimicrobial peptides in epithelia or the fat body. (Takehana et al., 2002) found that overexpression of *PGRP-LE* ubiquitously or in the fat body and hemocytes of larvae induced melanization. In contrast, (Libert et al., 2006) found that long or short-term overexpression of *PGRP-LE* in the adult fat body did not result in melanotic phenotypes. Activation of melanization in response to strong overexpression of *PGRP-LE* is consistent with tissue damage due to overactivation of the Imd pathway and activation of JNK/apoptotic programs, leading to necrosis and melanization (see e.g., Fink et al., 2016; Schmidt et al., 2011; Tsuda et al., 2005), but does not necessarily support a direct role of PGRP-LE in the melanization cascade.

(Takehana et al., 2004) generated a *PGRP-LE*$^{112}$ loss-of-function mutant in the *yw* background through P-element mobilization and found that melanization following septic injury with live *E. coli* was reduced, suggesting that PGRP-LE is required for melanization. Importantly however, the authors do not specify which genetic background was used as a wild-type control. This is particularly significant since the *yw* background that the *PGRP-LE*$^{112}$ mutation was generated in has reduced cuticle melanization. Here we tested whether PGRP-LE is required for infection-dependent melanization in *Drosophila* using two independent loss-of-function mutations in *PGRP-LE.*

**Methods**

*Fly lines and bacterial strains*

*PGRP-LE*$^{112}$ and *PGRP-LE*$^{\Delta 53}$ (both in the *yw* background) loss-of-function mutants were used. *PGRP-LE*$^{112}$ is the original mutant generated by Takehana et al. (2004). *PGRP-LE*$^{\Delta 53}$ was generated by François Rouyer (personal communication, nature of mutation unknown). Mutations were verified by PCR (*PGRP-LE*$^{112}$) and qPCR (*PGRP-LE*$^{112}$ and *PGRP-LE*$^{\Delta 53}$). In addition, we recombined the *PGRP-LE*$^{112}$ mutation away from the *yellow* marker by backcrossing to the *w*$^{iso}$ background to obtain *PGRP-LE*$^{112}$ flies with wild-type cuticle pigmentation. Chromosomes II and III were swapped to the *Drosdel iso* (Ryder et al., 2004) background, and the X chromosome was recombined to the *Drosdel iso* background to remove the *yw* mutation. This line is referred to as *PGRP-LE*$^{112}$*R. OR-R, yw*, and *w*$^{iso}$ *Drosdel* flies were used as wild-type in their respective backgrounds, and either *Hayan-psh*$^{Def}$ (Dudzic et al., 2019) or *PPO1*$^{\Delta}$*, PPO2*$^{\Delta}$ (Binggeli et al., 2014) double mutants were used as controls lacking melanization. The strains *E. coli 1106* and *M. luteus* JL (Jean Lambert collection, Strasbourg) were used in this study.

*Melanization in adult flies*

5 to 7-day old male and female flies reared on cornmeal fly medium (6% cornmeal, 6% yeast, 0.6% agar, 0.1% fruit juice, 10.6 g/L moldex and 4.9 mL/L propionic acid) at 25°C were anaesthetized with $CO_2$ and pricked on the left side of the thorax anterior and ventral to the wing joint with a 0.2 mm tungsten needle. The needle was sterilized with ethanol prior to clean injury or dipped in bacterial pellets (OD$_{600}$=200) concentrated from overnight cultures of *Escherichia coli*

(37°C, shaking, LB) or *Micrococcus luteus* (29°C, shaking, LB) prior to septic injury. Bacteria were diluted to the desired concentration in PBS. *M. luteus* was heat-killed prior to injury by heating the pellet to 67°C for 1 hour. Flies were incubated at rearing conditions for 18h after pricking and melanization of the injury site was scored. A dark brown or black spot was scored as "strong" melanization; a light brown or yellow spot as "weak" melanization; and no apparent spot as no melanization. At least 20 flies per genotype were scored in at least two independent experiments; data for male and female flies were combined as they yielded similar results.

*Melanization in larvae*

10 third instar larvae of each genotype were heated for 20 minutes at 37°C in 1mL PBS and imaged to assess melanization of crystal cells. To assess injury-dependent melanization, 20 larvae were pricked at the posterior end with a 0.1mm needle and placed on grape juice agar in a humidification chamber at 25°C for 2h. Melanization was scored similar to adults. Experiments were performed twice and yielded similar results.

#### Results

Contrary to (Takehana et al., 2004) we find using two independent *PGRP-LE* loss-of-function mutations that PGRP-LE is not required for infection-dependent melanization at the wound site, or for any melanization process tested in larvae or adults (Figure 1). Notably, we found that injury with live *E. coli* as used by (Takehana et al., 2004) produced more variable melanization than either clean injury or septic injury with heat killed *M. luteus* in adult flies of all genotypes (Figure 1C), which may have contributed to the original results. In addition to our results, (Bidla et al., 2007) found that hemolymph clots from *PGRP-LE*$^{112}$ larvae melanize normally.

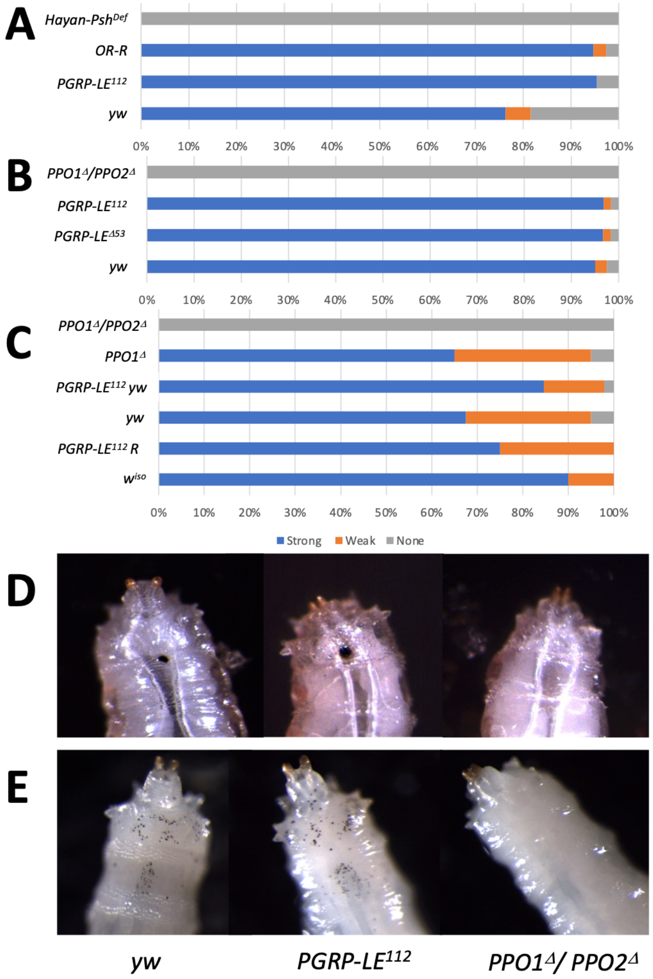

**Figure 1.** Melanization in PGRP-LE loss-of function mutants compared to wild-type. **A-C:** Percentage of adult flies displaying strong, weak or no melanization in response to **A)** clean injury,

**B)** septic injury with heat-killed *M. luteus*, or **C)** septic injury with live *E. coli*. **D)** Melanization of larvae in response to clean injury. **E)** Melanization of crystal cells in larvae in response to heating.

#### Conclusion

Using the PGRP-LE$^{112}$ mutant recombined away from the *yw* marker and an independent *PGRP-LE*$^{\Delta 53}$ mutant compared to appropriate background controls, we find that loss of *PGRP-LE* does not result in cuticular melanization defects.The original results may have been a result of comparison to a wild-type with a different genetic background than the mutant, or use of a bacterial elicitor that our results suggest produces variable melanization. We cannot however exclude a subtler role of PGRP-LE in other forms of melanization such as hemolymph melanization.

### Supplementary S10: PGRP-LE is not required for survival to *Bacillus* infection

**Abstract**

The *Drosophila* Imd pathway constitutes the major defense against Gram-negative bacteria, while the Toll pathway constitutes the major defense against Gram-positive bacteria and fungi. Although *Bacillus* bacteria are Gram-positive, they have DAP-type cell wall peptidoglycans that weakly activate both pathways. An early report found that flies deficient for the intracellular Imd receptor *PGRP-LE* were susceptible to infection with *Bacillus* species (Takehana et al., 2004). In contrast, experiments performed for ReproSci find no significant role for PGRP-LE alone in defense against *Bacillus* septic infection, and instead suggest that the initial result was due to comparison to a robust wild-type line.

**Introduction**

PGRP-LE is an intracellular pattern-recognition receptor of monomeric DAP-type peptidoglycan (TCT) in the *Drosophila* Imd pathway, culminating in a local or systemic immune response including transcription of antimicrobial peptides in epithelia or the fat body. (Takehana et al., 2004) found that a loss-of-function $PGRP\text{-}LE^{112}$ mutant had increased susceptibility to *Bacillus subtilis* and *Bacillus megaterium* infections. Susceptibility of these flies was intermediate between wild type (*OR-R*) and *PGRP-LE;;PGRP-LC* double mutant flies. However, the $PGRP\text{-}LE^{112}$ mutant was generated in the *yw* background and *OR-R* was used as the wild type comparison, making it possible that the observed difference in survival is due to background.

*Bacillus subtilis* and some other *Bacillus* species have amidated DAP-type peptidoglycans (Atrih et al., 1999; Schleifer and Kandler, 1972). Because of this, despite being Gram-positive (thus expected to elicit the Toll pathway), *Bacillus* species typically are weak inducers of both pathways (Chang et al., 2004; Leone et al., 2008; Leulier et al., 2003; Mellroth et al., 2005; Stenbak et al., 2004; Yamamoto-Hino and Goto, 2016). *Bacillus* species produce very low levels of TCT (Stenbak et al., 2004), the major ligand for the Imd pathway receptors PGRP-LE and PGRP-LCa (Kaneko et al., 2006). Recent studies have confirmed that the Toll pathway has a significant role in survival to *B. subtilis* infections (Hanson et al., 2019), but the role of the Imd pathway and particularly PGRP-LE in survival to infection by *Bacillus* species has not been independently assessed since (Takehana et al., 2004). To address this, we tested survival to infection with *Bacillus subtilis* of flies bearing a null mutation of *PGRP-LE,* and compared to appropriate background controls.

**Methods**

*Fly lines* and bacterial strain

The $PGRP\text{-}LE^{112}$ (*yw* background) loss-of-function mutant was generated by Takehana et al. (2004). Loss of *PGRP-LE* expression in the mutant was verified by PCR and qPCR. In addition, $PGRP\text{-}LE^{112}$ was backcrossed to $w^{iso}$ *Drosdel* (Ryder et al., 2004) to obtain the mutation in a different genetic background. Chromosomes II and III were swapped to the *Drosdel iso* background, and the X chromosome was recombined to the *Drosdel iso* background to remove the *yw* mutation. This line is referred to as $PGRP\text{-}LE^{112}R$. *OR-R, yw*, $w^{1118}$ (BDSC #3605) and $w^{iso}$ *Drosdel* flies were used to provide a range of wild-type comparisons. $Relish^{E20}$ *iso* (Ferreira et al., 2014) flies were used as controls lacking Imd activity. $Spz^{rm7}$ (Neyen et al., 2012) and $Bom^{\Delta 55C}$ *Drosdel* (Clemmons et al., 2015; Hanson et al., 2019) mutants were used as susceptible controls with reduced Toll pathway or Toll-dependent effector activity. *B. subtilis* FS strain was obtained from J.Millet and A. Klier, (Pasteur Institute of Paris).

*DNA extraction and PCR*

DNA was extracted from male and female flies by macerating single flies in 50 µL squishing buffer (10 mM Tris--Cl pH 8.2, 1 mM EDTA, 25 mM NaCl and 200 µg/ml Proteinase K) followed by incubation at 37°C for 30 minutes, and 2 minutes at 95°C. PCR was performed in a 25 µL reaction volume using AccuStart II PCR SuperMix (QuantaBio) at an annealing temperature of 54°C with a 60 s extension. Primers are specified in the supplementary of (Takehana et al., 2004) and amplify the $PGRP\text{-}LE^{112}$ mutation.

- *PGRP-LE*$^{112}$ Forward: 5'- CTG GCC CAA CTC GCT TCA GT -3'
- *PGRP-LE*$^{112}$ Reverse: 5'- GAG AGA CGT CTG CGA GCT CT -3'

*RNA extraction and qPCR*

Total RNA was extracted from pooled samples of 7-10 flies (5-7 days old) each using TRIzol reagent (Invitrogen), and re-suspended in MilliQ water. DNase treatment was performed using TURBO DNA-free kit (Invitrogen). Reverse transcription was performed using 0.5 mg total RNA in 10 µl reactions using Prime-Script RT (Takara) with random hexamer and oligo dT primers. Quantitative PCR was performed on a LightCycler 480 (Roche) in 96-well plates using PowerUp SYBR Green Master Mix (Applied Biosystems). *PGRP-LE* expression was normalized to expression of the ribosomal protein *rp49* (*Rpl32*).

- *Rp49* Forward: 5' – GCC GCT TCA AGG GAC AGT ATC TG – 3'
- *Rp49* Reverse: 5' – AAA CGC GGT TCT GCA TGA G – 3'
- *PGRP-LE* Forward: 5' - TGA CAC TAG GCA CTC AAA TCG C – 3'
- *PGRP-LE* Reverse: 5' - ACG TTG CCC AAA GTC TGA ATG C – 3'

*Survival assays*

5 to 7-day old male and female flies reared on cornmeal fly medium (6% cornmeal, 6% yeast, 0.6% agar, 0.1% fruit juice, 10.6 g/L moldex and 4.9 mL/L propionic acid) at 25°C were anaesthetized with $CO_2$ and pricked on the thorax anterior and ventral to the wing joint with a 0.2 mm tungsten needle dipped in a concentrated overnight (shaking, BHI, 29°C) culture of *Bacillus subtilis* at $OD_{600}$=5 diluted in PBS. Flies were incubated at rearing conditions and scored daily for survival; flies were flipped to new medium every 2-3 days. At least 20 flies per genotype and sex were scored in 2-3 independent replicates per genotype; data for male and female flies were combined as they were not significantly different. Data were analyzed in R using a Cox Proportional Hazard test.

#### Results

The fly sex and bacterial infection dose was not specified in the original paper. However, a high dose must have been used as survival was monitored for only ~72h at 25°C and significant death was observed although *Bacillus* species are mild pathogens for *Drosophila.* Here, we show that *PGRP-LE*$^{112}$ mutant flies do not have significantly increased susceptibility to *Bacillus subtilis* at a dose widely used for standard challenge. In both the *yw* and partially isogenised (*w*$^{iso}$ *Drosdel*) backgrounds, *PGRP-LE*$^{112}$ mutant flies had nearly identical survival to their respective background controls, and within the distribution of a variety of wild type lines (Figure 1). In contrast, *Relish*$^{E20}$ mutants lacking Imd activity had intermediate susceptibility to *B. subtilis,* while Toll-deficient genotypes were strongly susceptible, similar to (Hanson et al., 2019). This is consistent with previous results showing that the Toll pathway has a greater role in defense against *Bacillus* species.

#### Conclusion

Contrary to (Takehana et al., 2004), our results suggest that PGRP-LE does not significantly contribute to survival to *Bacillus subtilis* infection. Although we have only tested *Bacillus subtilis*, (Takehana et al., 2004) state that loss of PGRP-LE more strongly reduced survival to *B. subtilis* than *B. megaterium,* suggesting that the effect, if present, should have been visible using *B. subtilis.* Our results are consistent with *Bacillus* signalling through PGRP-LCx but not PGRP-LCa, likely because *Bacillus* species produce very small amounts of TCT (Stenbak et al., 2004), which is the major activating ligand for both PGRP-LCa and PGRP-LE (Kaneko et al., 2006).

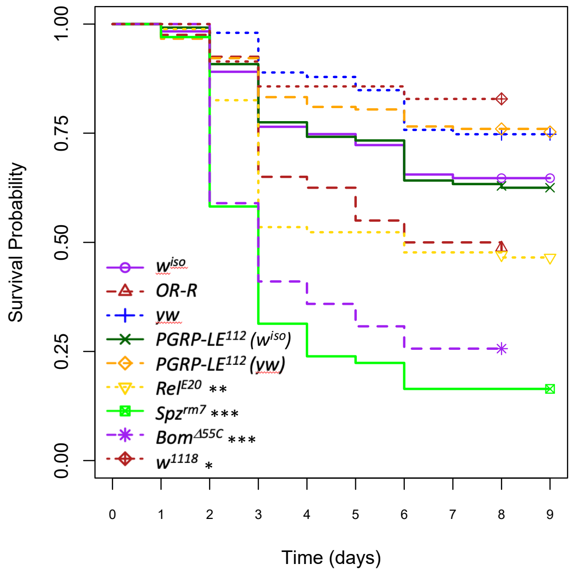

**Figure 1.** Survival to $OD_{600}$=5 *Bacillus subtilis* at 25°C. At least 20 flies per genotype and sex were scored in 2 independent experiments per genotype; pooled data for male and female flies were combined as they were not significantly different. Asterisks in legend indicate genotypes that were significantly different from the $w^{iso}$ wild type: * P<0.05, ** P<0.01, *** P<0.001.

### Supplementary S11: Impact of DNaseII on the regulation of antimicrobial peptide genes

**Abstract**

An initial study suggested that flies deficient in *DNaseII* undergo spontaneous immune activation of the Imd pathway due to the presence of undegraded DNA (Mukae et al., 2002), while another study showed that *DNaseII* deficient flies are sensitized to bacterial infection, and have reduced hemocyte numbers (Seong et al., 2006). No following studies have addressed the role of DNaseII in immunity. Using an independent *DNaseII*$^{sk4}$ null mutant and backcrossing of the hypomorphic *DNaseII*$^{lo}$ line used to obtain the original results, we find that loss of DNaseII does not reproducibly activate the immune response in the absence of infection, and that naked DNA is not immunogenic in *Drosophila melanogaster.* Contrary to (Seong et al., 2006), we found that *DNaseII* mutants did not have reduced hemocyte numbers. However, consistent with their findings, we find that *DNaseII* deficient flies are sensitized to bacterial infections through an indirect mechanism.

**Introduction**

Undegraded DNA can trigger sterile inflammation by acting as a DAMP (damage-associated molecular pattern) in mammals. Pioneer studies showed that accumulation of undegraded DNA inside cells stimulates the production of IFN-ß in mice, a factor that activates pathways homologous to NF-κB pathways of *Drosophila* (Okabe et al., 2005; Yoshida et al., 2005). Chromosomal DNA is typically degraded by deoxyribonucleases (DNases) as part of the engulfment and degradation of apoptotic cells. DNases also degrade extra-cellular DNA and prevent “sterile” activation of innate immunity in mammals.

The Drosophila gene *DNaseII* is induced upon septic injury, pointing to a role in the immune response (De Gregorio et al., 2001). (Mukae et al., 2002) found that a hypomorphic mutation of *DNaseII* (*DNaseII*$^{lo}$) resulted in accumulation of undegraded chromosomal DNA during oogenesis and activation of the innate immune response, including constitutive expression of antimicrobial peptide genes *Diptericin* and *Attacin*. These authors inferred that endogenous *Drosophila* DNA, when not degraded by DNase enzymes, activates innate immunity. Subsequent work confirmed a role for *DNaseII* during embryogenesis: (Tarayrah-Ibraheim et al., 2021) showed that DNaseII is required for cell death of primordial germ cells, and (Bass et al., 2009) produced similar results as (Mukae et al., 2002) showing impaired clearance of chromosomal DNA during embryogenesis using the same *DNaseII*$^{lo}$ mutants. (Seong et al., 2006) built on the immune findings by showing that flies knocked-down for *DNaseII* had increased susceptibility to Gram-negative and -positive bacteria, and decreased hemocyte numbers compared to wild-type flies.

Since these studies, the immune phenotypes of *DNaseII*-deficient *Drosophila* have not been further investigated. The results above suggest that susceptibility of infected *DNaseII* mutants is due to ineffective degradation of debris generated by pathogen clearance, rather than pathogen virulence. Although Imd pathway-dependent antimicrobial peptides were upregulated in *DNaseII* hypomorphs, there was no genetic demonstration that the Imd pathway was specifically activated. As extensive crosstalk occurs between immune pathways, it is essential to identify which intracellular and extracellular sensors are activated by DNA, and which downstream pathway(s) is/are activated. To address these questions, we generated an independent CRISPR null mutant of *DNaseII* and explored its immune characteristics.

**Methods**

*Drosophila stocks and rearing conditions*

All fly stocks were maintained on cornmeal fly medium (6% cornmeal, 6% yeast, 0.6% agar, 0.1% fruit juice, 10.6 g/L moldex and 4.9 mL/L propionic acid) at 25°C. Experiments were performed on 5-7 days old adults, or third instar wandering larvae (L3) selected 110-120 hours after egg laying. Isogenic $w^{1118}$ Drosdel flies ($w^{iso}$) (Ryder et al., 2004) were used as control. *DNaseII*$^{sk4}$ mutants were generated using CRISPR-mediated mutagenesis as previously described (Kondo and Ueda, 2013). There is one nucleotide deletion causing a frameshift with a stop codon at amino-acid 90. This line was backcrossed to the $w^{iso}$ isogenic background to replace chromosome II. *DNaseII$^{lo}$* hypomorphic mutants were obtained from Bloomington (Stock #1042). The *Bom$^{\Delta 55C}$* and *Relish$^{E20}$* flies were the same as used in (Carboni et al., 2022). The Sting$^{JLI}$ line is described in (Cai et al., 2020).

*Survivals*

Bacteria were cultured on a shaking plate at 180 rpm. Overnight cultures were pelleted by centrifugation (4000 rpm at 4°C) and resuspended to the desired optical density at 600 nm ($OD_{600}$). *Pectobacterium carotovorum carotovorum 15* (*Ecc15*) and *Micrococcus luteus* JL (Jean Lambert collection, Strasbourg) were grown in shaking LB media at 29°C. *Enterococcus faecalis* was grown shaking in BHI medium at 37°C. All experiments were done on 3- to 5-day old adult female flies. Flies were pricked in the thorax with a 0.1 mm thick needle previously dipped into the concentrated bacterial pellet. Infected flies were maintained at 25°C (*E. faecalis ATCC 29212*) or 29°C (*Ecc15* and *M. luteus*) and survivals were recorded daily. Flies were flipped into fresh vials every 2-3 days. 5-10 flies were used per genotype per replicate, and at least three replicates were performed for each genotype.

*DNA injections*

DNA was extracted from an *E. coli* pellet (overnight, shaking, LB, 37°C) or from 5 flies using the High-quality DNA extraction for RT-PCR and Sequencing kit (Qiagen) following manufacturers' instructions. Briefly, tissues were lysed on ice, heated to 65°C for 15 minutes, and 100 µL of protein precipitation solution was added. The samples were centrifuged and the supernatants containing DNA were harvested. The DNA was precipitated using isopropanol and washed two times with ethanol. DNA was re-hydrated with MilliQ water. The samples were adjusted to the desired concentration (200 ng/ul), and 23,3nl were injected using a nanoinjector Nanoject II (Drummond) and glass capillary needles.

*RT-qPCR*

Gene expression measurements were performed by RT-qPCR as previously described (Carboni et al., 2022). Briefly, 5 whole flies/larvae were homogenized and their RNA was extracted using TRIzol (Invitrogen) and resuspended in RNase-free water. Reverse transcription was performed using PrimeScript RT kit (Takara) with random hexamers and oligo dTs. Quantitative PCRs were performed on a LightCycler 480 (Roche) using PowerUp SYBR Green Master Mix (Applied Biosystems). Expression levels were normalized to that of *rp49* (*Rpl32*).

- *Rp49* Forward: 5' – GCC GCT TCA AGG GAC AGT ATC TG – 3'
- *Rp49* Reverse: 5' – AAA CGC GGT TCT GCA TGA G – 3'
- *Diptericin A* Forward: 5' – GCT GCG CAA TCG CTT CTA CT – 3'
- *Diptericin A* Reverse: 5' – TGG TGG AGT GGG CTT CAT G – 3'

*Statistical analysis*

Survival experiments were performed using a Cox proportional hazards (CoxPH) regression model in R 1.4.1103. Each experiment was repeated independently three times unless otherwise indicated. Quantitative PCR data were compared by one-way ANOVA with Holm-Šidak multiple test correction in Prism (GraphPad). Hemocyte counts were performed using FACS counting of larval hemocytes as described in (Melcarne et al., 2019). Hemocyte count data was analyzed by T-test. Error bars represent the standard error of the mean of replicate experiments (SEM). P values are represented in the figures by the following symbols: ns for $p \geq 0.05$, * for *P* between 0.01 and 0.05; ** for *P* between 0.001 and 0.01, *** for *P* between 0.0001 and 0.001, **** for $p \leq 0.0001$.

#### Results and Discussion

*Immune gene expression is not constitutively active in DNaseII mutant flies*

(Mukae et al., 2002) previously found that accumulation of undegraded chromosomal DNA from apoptotic cells in phagocytes of *DNaseII$^{lo}$* mutant flies resulted in constitutive elevated expression of *Diptericin* and *Attacin,* but not *Drosomycin.* This suggests that endogenous

*Drosophila* DNA, when not degraded by the DNase II enzyme, activates innate immunity (Mukae et al., 2002). This finding is consistent with data showing that undegraded DNA activates immunity in other animal models (Liu et al., 2008; Yu et al., 2015), but has not been further investigated in *Drosophila*. Furthermore, the original results did not clarify which specific immune pathways were activated, or how this activation occurred.

In unchallenged conditions, we confirmed that the expression level of *Diptericin A* (a readout for the Imd pathway) was significantly higher in hypomorphic *DNaseII*$^{lo}$ adult flies than wild type flies, although much lower than the level observed upon septic injury. However, the independent *DNaseII*$^{sk4}$ null mutant and *DNaseII*$^{lo}$/*DNaseII*$^{sk4}$ transheterozygous crosses did not display similar antimicrobial peptide gene upregulation (Figure 1A). We therefore backcrossed the *DNaseII*$^{lo}$ line to the isogenic Drosdel *w*$^{1118}$ (*w*$^{iso}$) background to replace chromosomes I and II (referred to as *DNaseII*$^{lo}$ *(iso)*) and compared this to a similarly backcrossed wild type line (+;+;*w*$^{1118}$) (Figure 1A). Backcrossing eliminated constitutive activation of *DptA* in the *DNaseII*$^{lo}$ line and produced results similar to those obtained with *DNaseII*$^{sk4}$, indicating that this activation is an artifact associated with genetic background. Contrary to (Mukae et al., 2002), we conclude that loss of DNaseII does not constitutively activate immune gene expression in *Drosophila.*

*Naked DNA is not immunogenic in* Drosophila

To determine whether DNA is generally immunogenic in wild-type and *DNaseII* mutant flies, we injected flies with endogenous DNA (*i.e.*, from *Drosophila*) or exogenous bacterial DNA (from *E. coli*). Backcrossed *DNaseII*$^{lo}$ *(iso)* and *DNaseII*$^{sk4}$ mutant flies were not susceptible to DNA injections (Figure 1B, C). Moreover, *Diptericin A* gene expression levels did not differ between wild-type and *DNaseII*$^{lo}$ *(iso)* mutants injected with PBS or DNA (Figure 1D). These results suggest that loss of DNaseII has no effect on basal or DNA-induced immune activation in unchallenged adult flies. Furthermore, injection of DNA into the hemolymph is not immunogenic in flies regardless of DNaseII (Figure 1D). Notably, our experiments were performed using naked DNA (protein-free DNA). Further experiments are needed to see whether proteins such as histones associated with DNA are immunogenic rather than the DNA strands themselves.

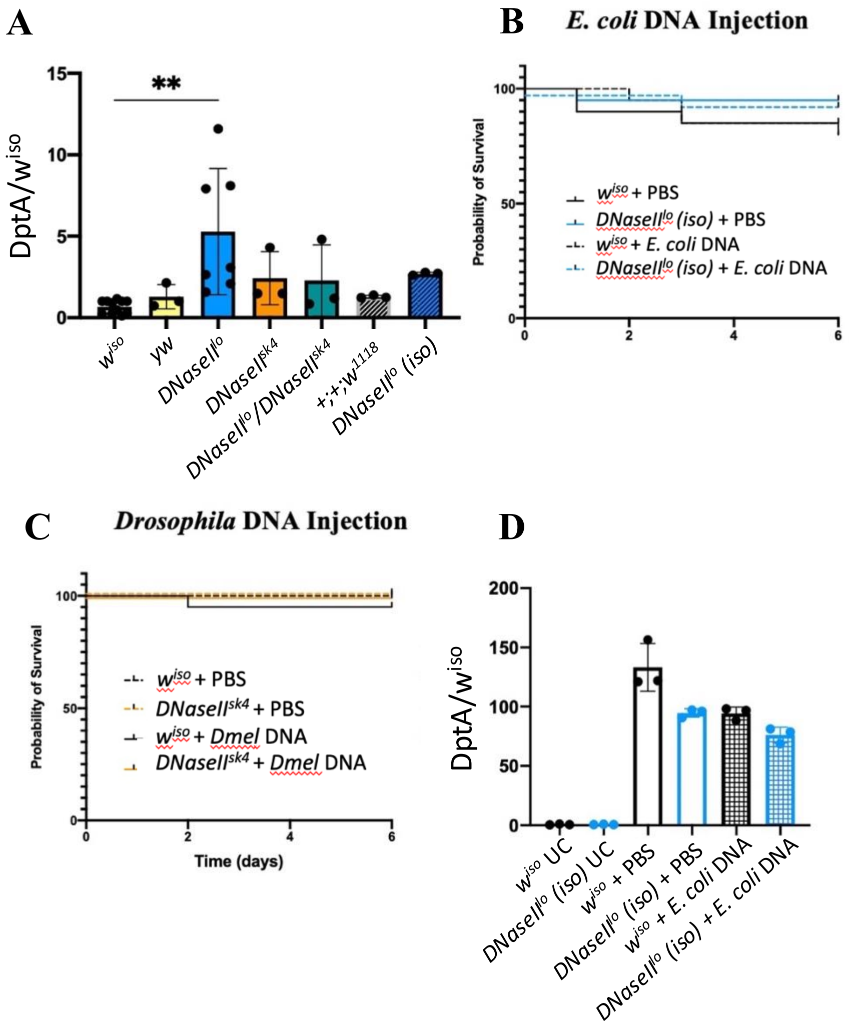

**Figure 1: Naked DNA is not immunogenic in *Drosophila.* A:** qRT-PCR of *DptA* expression in unchallenged adult flies, including *DNaseII$^{lo}$* mutants in the original genetic background and backcrossed to the isogenic Drosdel *w$^{iso}$* background to replace chromosomes I and II (*DNaseII$^{lo}$ (iso)*). Error bars represent the standard deviation. **B:** Survival of wild type (*w$^{iso}$*) and backcrossed *DNaseII$^{lo}$* hypomorphic mutant adult flies after PBS injection, and after injection of DNA extracted from *E. coli*. **C:** Survival of wild type (*w$^{iso}$*) and backcrossed *DNaseII$^{lo}$* hypomorphic mutant adult flies after PBS injection, and after injection of DNA extracted from flies. **D:** qRT-PCR of *DptA* expression in wild-type (*w$^{iso}$*) and backcrossed *DNaseII$^{lo}$* flies 6h after injection with PBS (control) or *E. coli* DNA.

DNaseII *deficient flies are susceptible to bacterial infections*

(Seong et al., 2006) found that *DNaseII* knockdown in the fat body (*Cg-Gal4>DNAseII-RNAi* or *c564-Gal4>DNaseII-RNAi*) strongly increased fly susceptibility to infection with Gram-negative (*E. coli*) and Gram-positive (*M. luteus*) bacteria. Furthermore, the level of susceptibility to infection correlated with DNaseII activity levels measured by a radial diffusion assay. The authors concluded that DNaseII is required to maintain proper immune function in *Drosophila.* In our hands, *DNaseII$^{sk4}$* flies were not susceptible to clean injury, but were susceptible to Gram-negative (*Ecc15*) and Gram-positive (*E. faecalis*) bacterial infections compared to wild-type flies (Figure 2A, B). These results agree with the findings of (Seong et al., 2006).

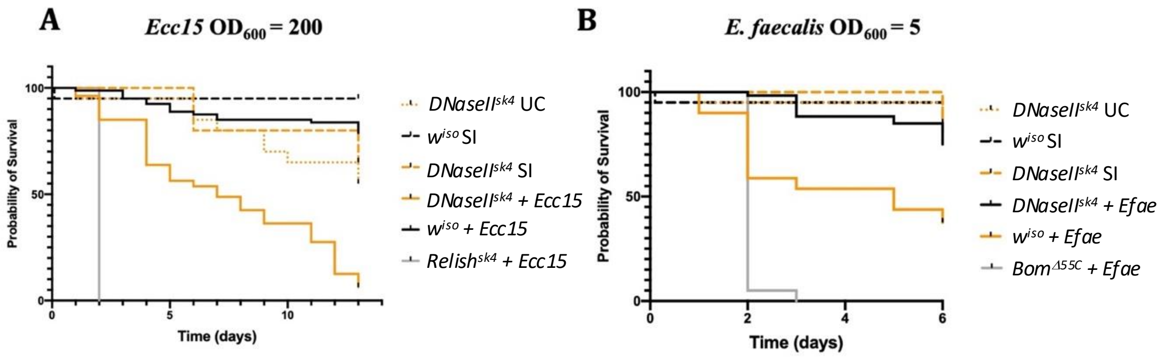

**Figure 2: DNaseII deficient flies are susceptible to bacterial infection. A, B:** Survival of wild type ($w^{iso}$) and *DNaseII*$^{sk4}$ flies upon infection with *Pectobacterium carotovorum Ecc15* (**A**) and *E. faecalis* (*Efae*) (**B**). UC, unchallenged; SI, sterile injury.

Hemocyte numbers are slightly higher in *DNaseII* deficient larvae

(Seong et al., 2006) found that hemocyte numbers were significantly reduced in flies by knockdown of *DNaseII* in the fat body (*Cg-Gal4>DNAseII-RNAi* or *c564-Gal4>DNaseII-RNAi*). Contrary to (Seong et al., 2006), hemocyte counts in wild type ($w^{iso}$) and *DNaseII*$^{sk4}$ mutant larvae show that hemocyte numbers are slightly higher in *DNaseII* deficient larvae (Figure 3, p=0.03). The lower hemocyte counts found by (Seong et al., 2006) may be attributable to genetic background of the RNAi lines used in this study, similar to the elevated antimicrobial peptide gene expression found in *DNaseII*$^{lo}$ larvae by (Mukae et al., 2002) (Figure 1A). Alternatively, this could be due to a change in proportion between sessile and circulating hemocytes rather than a change in absolute hemocyte number; results have shown that hemocyte extraction without vortexing larvae recovers only a proportion of hemocytes, as many are sequestered in sessile patches under the cuticle, information that was not available at the time of the initial study (Makhijani et al., 2011).

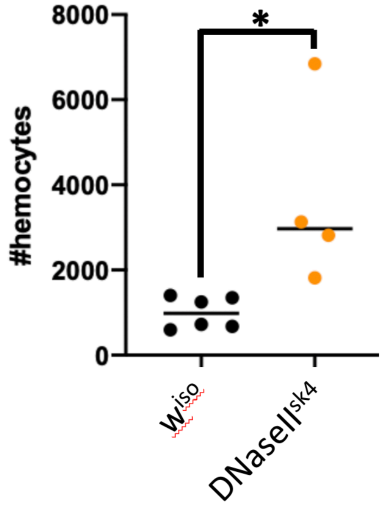

**Figure 3: Hemocyte count in *DnaseII*$^{sk4}$ larval hemocytes.** Hemocyte counts in *DNaseII*$^{sk4}$ L3 wandering larvae are higher than in wild type ($w^{iso}$), p=0.03.

**Conclusion**

In contrast to (Mukae et al., 2002), we find that loss of DNaseII does not cause constitutive activation of antimicrobial peptide gene expression, and that upregulation seen in hypomorphic *DNaseII*$^{lo}$ mutants was due to genetic background. Furthermore, we find that naked DNA is not immunogenic in *Drosophila*. In support of the notion that excess of DNA in *DnaseII* mutant does not activate Imd signaling, germ-free conditions eliminate high Dpt expression in *DnaseII*$^{lo}$ flies (West, 2018).

In agreement with previous RNAi-based results (Seong et al., 2006) we find that *DNaseII* deficient flies are susceptible to systemic bacterial infections. However, contrary to classical immune-deficient mutants, *DNaseII$^{sk4}$* mutants show no hemocyte number decrease and we even found a slight increase in hemocyte number. In agreement with previous results (Seong et al., 2006), we found that *DNaseII$^{sk4}$* mutant flies have a delay in development, reduced lifespan, and a strong locomotor defect. This general reduction in fitness could explain the susceptibility to infections, although this remains to be tested experimentally.

### Supplementary S12: The JNK pathway does not directly regulate antimicrobial peptide gene expression

**Abstract**

The JNK pathway has been suspected to regulate antimicrobial peptide (AMP) gene expression in *Drosophila*, but studies have yielded contradictory results. This hypothesis is supported by results from (Delaney et al., 2006; Kallio et al., 2005; Kleino et al., 2005) but contradicted by (Boutros et al., 2002; Kenmoku et al., 2017; Park et al., 2004; Silverman et al., 2003). Experiments performed for ReproSci and review of the literature suggest that JNK is unlikely to have a strong direct role on *Drosophila* AMP regulation *in vivo,* and is not required in the adult fat body for AMP expression in response to infection. Previous results finding a direct role for JNK in AMP production are likely due to a combination of **i)** use of cell culture systems that do not fully reflect *in vivo* systems; **ii)** use of kinase-dead rather than null mutants of TAK1; and **iii)** underappreciation for the overarching role that JNK plays in cytoskeletal remodeling, pathway activation, and cell differentiation, which is not specific to immunity.

**Introduction**

(Delaney et al., 2006) found that AMP expression was impaired in a *dTAK1*$^{179}$ kinase dead mutant despite functional Relish processing and nuclear translocation. (Kallio et al., 2005; Kleino et al., 2005) similarly found that RNAi of TAK1 or other JNK intermediates in S2 cells impaired signal-dependent AMP reporter production, without reducing Relish cleavage or translocation. (Delaney et al., 2006) further found that overexpression of activated *hep* (a JNKK) rescued signal-dependent *Att* and *Dpt* expression in *dTAK1*$^{179}$ but not *IKKγ*$^{key}$ mutants in response to infection. Furthermore, overexpression of *puc*, a JNK inhibitor, suppressed expression of AMPs, and JNK mutant mosaic clones in the larval fat body failed to express (*Dpt* and *Att*) or had reduced expression (*Mtk* and *Drs*) of AMPs. Based on this evidence, they concluded that JNK signaling is required for AMP gene expression in the fat body of adult flies and larvae.

In contrast to this, several other groups found that RNAi against JNK pathway intermediates in S2 cells did not impair signal-dependent AMP expression (Boutros et al., 2002; Silverman et al., 2003; Park et al., 2004). More recently, (Kenmoku et al., 2017) overexpressed dominant-negative *bsk* in the larval fat body and found no role for JNK in induction of AMPs in larvae following sterile injury. (Paquette, 2009) note that the long-term ectopic JNK pathway activation employed by (Delaney et al., 2006) for *in vivo* experiments may have developmental effects, and does not reflect the transient activation of JNK that occurs in response to septic injury or immunostimulatory peptidoglycan fragments.

One of the major conclusions of (Delaney et al., 2006; Kallio et al., 2005), that TAK1 is not required for Relish cleavage and nuclear translocation ('activation'), is technically correct as stated. However, later studies refined this by showing that TAK1 is required for Relish phosphorylation and full transcriptional activity, even though it is not required for cleavage (Ertürk-Hasdemir et al., 2009). Therefore, the finding that AMP expression was reduced in *dTAK1*$^{179}$ kinase dead mutants despite successful Relish cleavage is not strong evidence that JNK is required for AMP expression, but does show that the kinase activity of Tak1 is required for full Relish functionality. The (Delaney et al., 2006) finding that overexpression of activated *hep* rescued AMP expression in *dTAK1*$^{179}$ but not *IKKγ*$^{key}$ mutants may indicate that *hep* is capable of phosphorylating Relish when present in excess.

A major limitation of the analyses discussed above is that JNK is required for expression of many genes involved in cytoskeletal remodeling (Boutros et al., 2002; Homsy et al., 2006; Irving et al., 2005) and hemocyte functions requiring cell shape change, including motility, differentiation and proliferation (Williams et al., 2006; Irving et al., 2005; Tokusumi et al., 2009; Zettervall et al., 2004; Ming et al., 2014; Ramesh et al., 2021; Stofanko et al., 2008). Disruption of cytoskeletal function or vesicular trafficking (which contributes to processes including endosomal recycling) can have strong effects on immune pathway activation and attenuation (e.g. (Akbar et al., 2016; Neyen et al., 2016; Schmid et al., 2016)). In a cell culture system such as the S2 cells used in the analyses discussed above, inhibition of JNK may have a number of effects that result in secondary reduction of AMP expression or translation.

This is not to say there is no direct cross-regulation between immune pathways and JNK signaling, as many studies have found evidence of this. Expression of *Jra* (*jun*) and *puc* appear to be partially regulated by Toll/Imd (De Gregorio et al., 2002), while *Relish* is a target of JNK (Soory and Ratnaparkhi, 2021). Imd and JNK activity are potentially interrelated in a number of additional ways: **(i)** they share the TAK1/TAB2 complex, allowing JNK to be activated by peptidoglycan upstream of Imd (Hua et al., 2022); **(ii)** they share the ubiquitin ligase POSH, although only the scaffolding activity of POSH is required for JNK, whereas Imd requires its ligase activity (Tsuda et al., 2005; Zhang et al., 2010); **(iii)** Relish activity can upregulate *DIAP1* to suppress JNK, and can negatively affect developmental processes (Tavignot et al., 2017) **(iv)** JNK activity can suppress DIAP2 activity to attenuate Imd (Kietz et al., 2022); and **(v)** overactivation of JNK can result in tissue damage that activates Imd signaling.

To clarify whether JNK activation is required for AMP expression *in vivo,* we tested *Diptericin* expression in response to septic infection in adult flies with reduced JNK activity in the fat body.

**Methods**

*Fly stocks*

Flies were developed at 18°C to suppress driver activity and overcome developmental lethality associated with JNK depletion. The Bloomington food recipe (39 L water, 625 g yeast, 390 g soy flour, 2850 g yellow cornmeal, 225 g agar, 3 L light corn syrup, 188 mL propionic acid (>99% pure, 13.4M)) was used for rearing flies. After eclosion, flies were transferred to 29°C for 7-10 days, then infected by pricking with 0.2 mm tungsten needle in the thorax dipped in $OD_{600}$=200 *Ecc15* from an overnight culture (29°C, shaking, LB). Three independent replicates were performed with 7-10 flies per replicate per genotype.

| **Genotype** | **Stock number** |
|---|---|
| *c564-Gal4* | BDSC6982 |
| *UAS-Bsk$^{DN}$* | BDSC6409 |
| *UAS-Hep-RNAi* | BDSC35210 |
| *w$^{1118}$* | VDRC |

*RNA extraction and qRT-PCR*

Flies were collected, sorted, and transferred to TRIzol reagent, homogenized with a motorized pestle, and snap-frozen in dry ice. Total RNA was extracted using the Direct-zol RNA Microprep kit following the manufacturer’s protocol. RNA (2mg) was reverse transcribed using the High-capacity Script cDNA Synthesis kit, and the synthesized cDNA was amplified by QPCR using the SYBR-Green PCR master mix using a QuantStudio 6 Flex qPCR machine (Applied Biosystems, Thermo Fisher Scientific). Samples were normalized to *Rpl23* gene expression, the relative quantification methods, and primer efficiency. Primer sequences used in real-time qPCR are: Bsk-F 5’-CACCAACACTACACCGTCGA; Bsk-R 5’ AAGCGGCGCATACTATTCCT; hep-F 5’ CCCCGCCGACAACTAGAGTG; hep-R 5’ CACCACCGGGACCACTAGAAA; Rpl23-F 5’ GACAACACCGGAGCCAAGAACC; Rpl23-R 5’ GTTTGCGCTGCCGAATAACCAC.

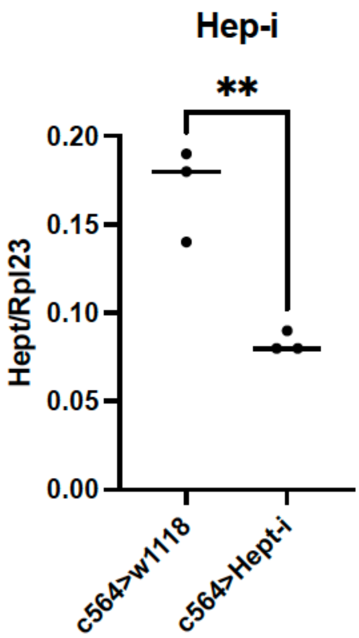

**Figure 1.** Validation of the *Hep* RNAi construct. The expression of *Hep* in *C564>HepRNAi* flies is decreased. For control treatments, the fat body *c564-Gal4* driver was crossed to the $w^{1118}$ wild-type flies. Results are from three independent experiments

**Results**

We knocked down JNK activation in the fat body of adult flies and tested *Diptericin* expression in response to septic infection with the Gram-negative bacterium *Ecc15*. To silence the JNK pathway, we over-expressed a dominant negative form of the JNK Basket (*UAS-Bsk$^{DN}$*) or RNAi against the JN2K Hemipterous (*UAS-Hep-RNAi*) using the fat body *c564-Gal4* driver. These treatments did not reduce expression of the Imd pathway readout *Diptericin* in either male or female flies compared to the wild-type control (Figure 2). Instead, *Diptericin* expression was slightly increased compared to wild-type at 8h following septic infection with *Ecc15* in the Hep knockdown flies. *Diptericin* expression in unchallenged flies was similar between JNK-deficient and wild-type flies (Figure 3).

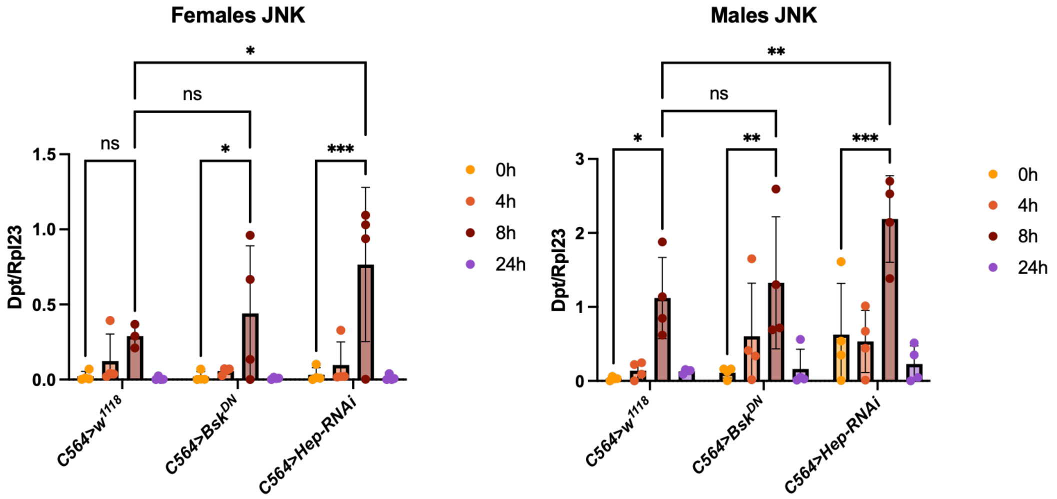

**Figure 2.** *Diptericin* expression in female (left) and male (right) control and JNK-deficient flies in response to *Pectobacterium carotovorum Ecc15* septic infection ($OD_{600}$ = 200). For control treatments, the fat body *c564-Gal4* driver was crossed to the $w^{1118}$ wild-type flies; for active treatments, *c564-Gal4* drove overexpression of a dominant negative allele of the JNK kinase Basket (*UAS-Bsk$^{DN}$*) or RNAi against the Basket kinase Hemipterous (*UAS-Hep-RNAi*). Within-timepoint comparisons showed no significant reduction in *Diptericin* expression between control and JNK-suppressed flies in response to infection. Instead, at the 8h time point only, male and

female Hep-RNAi flies showed a mild increase in *Diptericin* expression. Data were analyzed in Prism using a two-way ANOVA followed by Šídák's multiple comparisons test. ***, p<0.001; ** p<0.01; *p<0.05; ns, not significant. Results are from four independent experiments.

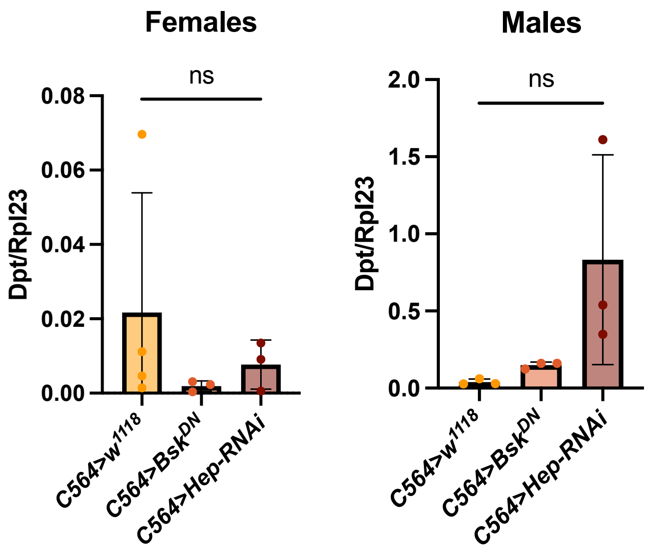

**Figure 3.** *Diptericin* expression in female (left) and male (right) experimental genotypes did not differ significantly from the control in unchallenged conditions. Data were analyzed in Prism using an ordinary one-way ANOVA followed by Šídák's multiple comparisons test. ns, not significant. Results derived from three independent measurements

#### Conclusion

Experiments performed for ReproSci show that inhibition of JNK (RNAi against *hep* or overexpression of $bsk^{DN}$) in the adult fat body does not significantly suppress *Diptericin* expression in response to infection with *Ecc15.* This contradicts conclusions of (Delaney et al., 2006; Kallio et al., 2005; Kleino et al., 2005) and supports those of (Boutros et al., 2002; Park et al., 2004; Silverman et al., 2003). This is in agreement with the more recent *in vivo* results of (Kenmoku et al., 2017). In the absence of further evidence, we conclude that JNK does not have a direct role in regulating AMP expression *in vivo*.

### Supplementary S13: Role of Caspar in negative regulation of Imd signaling

**Abstract**

(Kim et al., 2006) found using a *caspar*$^{P1}$ null allele that *caspar* is a negative regulator of the Imd pathway that acts by inhibiting DREDD-dependent cleavage and nuclear translocation of Relish. Contrary to this, using an alternative allele (*caspar*$^{lof}$) we found that *caspar*-deficient flies and larvae did not display constitutive upregulation of *Diptericin* in absence of infection. We did observe a stronger expression of *Diptericin* upon infection in *caspar*$^{lof}$ compared to wild-type in larvae but not adults. We also did not find increased resistance to *Ecc15* infection in *caspar*-deficient flies. In our hands, overexpression of *caspar* mildly suppressed *Dpt* expression at the 6hpi time point only. We conclude that *caspar* is not a canonic negative regulator of the Imd pathway on the whole-body level in absence of infection. Caspar may still regulate the Imd pathway in specific contexts.

**Introduction**

(Kim et al., 2006) found that a *caspar*$^{P1}$ mutant had constitutive melanization of the gut and fat body, constitutive induction of *Diptericin* to 60% of the level caused by infection, and strongly increased survival to Gram-negative *Escherichia coli* or *Ecc15* infection compared to wild-type flies. Transgenic expression of *caspar* rescued constitutive overexpression of *Dpt* in the mutant and inhibited *Dpt* induction in wild-type flies in a dose-dependent manner. Overexpression of *caspar* in the fat body (*yolk-Gal4*) decreased survival and increased bacterial load similar to *key*$^{1}$ and *relish*$^{E20}$ alleles. Immunostaining showed that overexpression of *caspar* blocked nuclear translocation of Relish in the fat body following immune challenge, while Relish was constitutively nuclear in uninfected *caspar*$^{P1}$ flies. Epistasis showed that Caspar acted at the level of DREDD. Toll activation and *Drs* expression was unaffected in *Caspar*-deficient flies. Based on these lines of evidence (Kim et al., 2006) concluded that Caspar acts as a negative regulator of the Imd immune signaling pathway.

Relatively little subsequent work has been done on the role of *caspar* in the immune response. (Kounatidis et al., 2017) found that *caspar* mutants did not share phenotypes found for negative regulators of Imd including elevated antimicrobial peptide (AMP) gene expression in the brain, increased neurodegeneration and decreased lifespan, but they note that this may be a tissue specific effect. (Handu et al., 2015) found that overexpression of *caspar* in S2 cells did not modulate AMP expression, but that LPS (containing immunostimulatory peptidoglycan fragments) stimulation strongly increased SUMOylation of Caspar (Sumo/Ubc9-dependent) similar to some other proteins involved in the immune response (e.g. IKKß (Fukuyama et al., 2013)). (Han et al., 2020) found that overexpression of *caspar* in the brain suppressed translocation of cleaved Relish to the nucleus, and suppressed neuronal phenotypes associated with overactivation of Imd signaling (similar to the phenotype produced by RNAi of DREDD).

The available evidence suggests that Caspar may negatively regulate the Imd pathway in certain contexts. However, further investigation is needed to confirm that Caspar acts as a negative regulator of Imd *in vivo* on a whole-body level both in absence of infection and upon infection. To test this, we looked at Imd pathway activation and survival in *caspar* loss-of-function and overexpressing flies.

**Methods**

Flies were raised on standard cornmeal agar at 25°C unless otherwise stated. *Casp*$^{lof}$ (BDSC #11373), *Casp*$^{Df}$ (BDSC #23691) and the adult fat body Gal4 driver (BDSC #6983, c729-Gal4) were obtained from the Bloomington *Drosophila* Stock Center. The *pUASp casp*$^{wt}$ transgenic line is described in (Das et al., 2024). Crosses and experiments were performed at 25°C except for *caspar* overexpression lines (*Fat Body Gal4*> *UASp Casp*$^{wt}$), which were maintained at 29°C. Flies were subjected to septic thoracic injury using a 0.2mm tungsten needle dipped in concentrated cultures of *Pectobacterium carotovorum Ecc15* grown overnight in LB at 29°C and diluted to $OD_{600}$=200 in PBS.

#### Results

Contrary to (Kim et al., 2006), experiments performed for ReproSci showed no enhanced resistance of *caspar* null flies to infection with *Ecc15* (Figure 1A), and showed no striking elevation of *Diptericin* expression in unchallenged or infected conditions in these flies compared to wild-type (Figure 1B, C, D). The level of expression of Dpt in unchallenged *$caspar^{lof}$/Df(Caspar)* was in the range of unchallenged flies and low compared to bacteria challenged flies (roughly 2% of wild-type challenged flies; Figure 1B). Furthermore, overexpression of *caspar* in adult flies did not decrease *Diptericin* expression in unchallenged conditions (Figure 1B), although this treatment did partially suppress *Dpt* expression at 6 hours post-infection (hpi) (Figure 1C). In *caspar* null larvae, *Diptericin* expression was elevated at 6 hpi but not in unchallenged conditions (Figure 1E).

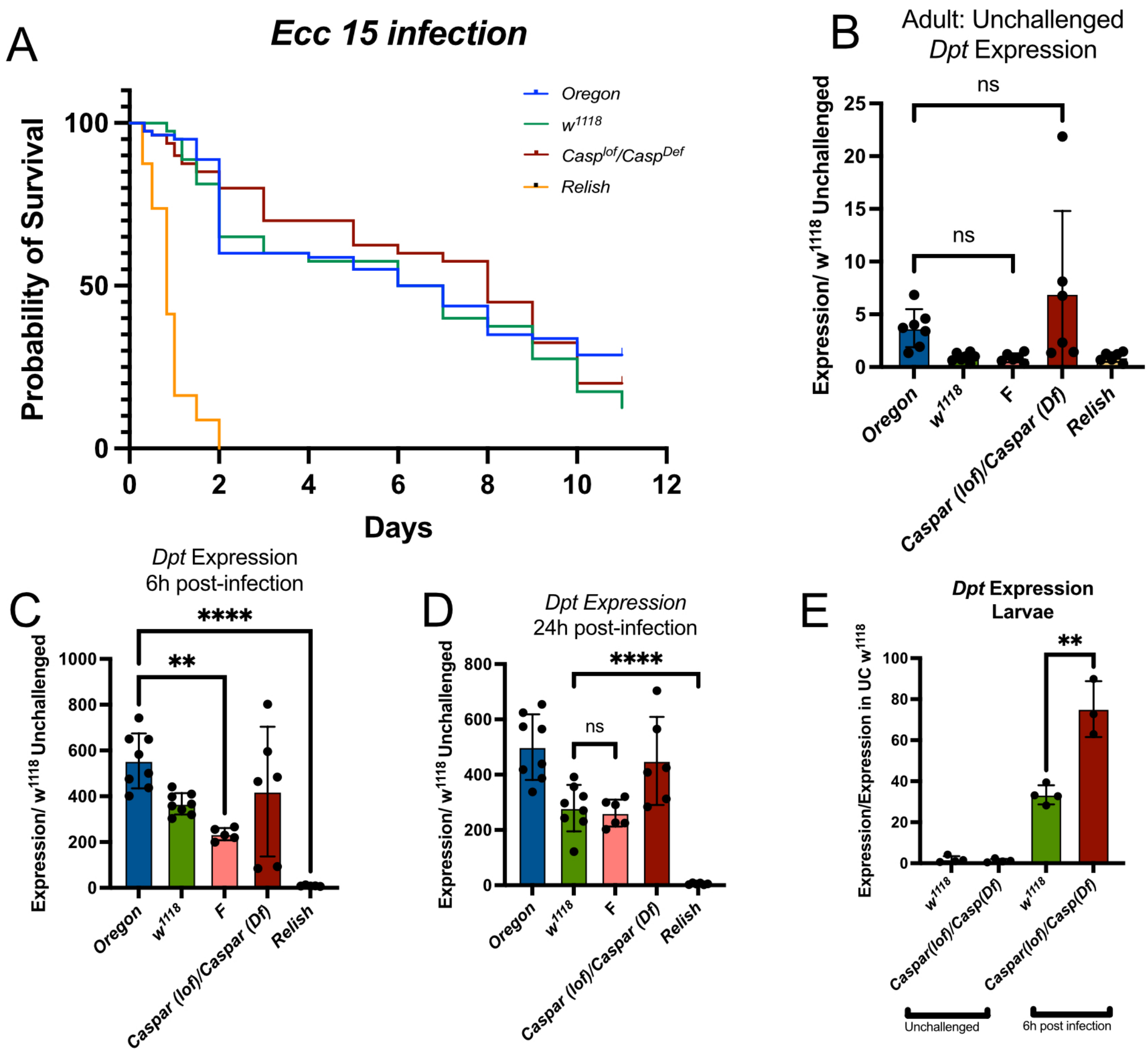

**FIGURE 1 A)** Survival of *caspar* null (*$Casp^{lof}/Casp^{Df}$*) adult flies to septic infection with *Ecc15* ($OD_{600}$=200) with 80 flies per genotype at 25°C. '*Relish*' indicates the *$Rel^{E20}$* allele. **B-D)** Expression of *Diptericin* in *caspar*-null (*$Casp^{lof}/Casp^{Df}$* ) and caspar-overexpressing (**F**, Fat Body gal4>UAS Caspar) adult flies in unchallenged conditions **(B)** and 6h **(C)** and 24h **(D)** after septic infection with Ecc15 ($OD_{600}$=200) as shown by qPCR. **E)** Expression of *Diptericin* in *caspar*-null (*$Casp^{lof}/Casp^{Df}$*) larvae. Diptericin expression is normalized to the housekeeping gene *RpL32*. Individual measurements are shown. ns, not significant; *, P<0.05; ** P<0.01; *** P<0.001. Data were visualized in GraphPad Prism 8.4.3. Statistics: One-way ANOVA with multiple comparison analysis was used to detect significance of results in a singular condition. Each dot represents independent experiments.

#### Conclusion

Our results indicate contrary to (Kim et al., 2006) that Caspar is likely not a canonical negative regulator of the Imd pathway on a whole-body level in absence of infection. We still observe a reduced activation of the Imd pathway upon caspar over-expression at an early time point as well as a higher Dpt expression in *caspar* deficient larvae upon infection, findings that require further investigation. Initial results of (Kim et al., 2006) may be due to effects of genetic background in the *caspar*$^{P1}$ allele. Caspar may impact the Imd pathway in certain contexts but is likely not a canonical negative regulator of the Imd pathway.

### Supplementary S14: Role of Edin in Immunity

**Abstract**

(Gordon et al., 2008) identified *edin,* an antimicrobial peptide (AMP)-like gene highly expressed upon *Listeria monocytogenes* infection. They found that overexpression of *edin* reduced lifespan and fly fitness whereas RNAi knockdown of *edin* in the fat body increased susceptibility to *L. monocytogenes* infection. Contrary to (Gordon et al., 2008), experiments performed for ReproSci using an *edin* null mutant indicate that loss of *edin* does not increase susceptibility to *L. monocytogenes* infection compared to appropriate controls. Moreover, overexpression of *edin* did not cause a major reduction in lifespan.

**Introduction**

In a paper identifying *edin* as a novel Imd-regulated AMP-like gene, (Gordon et al., 2008) claimed that increased susceptibility of *wntD* mutants to *Listeria monocytogenes* infection was due to reduced tolerance caused by elevated expression of the AMP-like gene *edin*. This inference was based on experiments showing that ubiquitous overexpression of *edin* caused high developmental and age-dependent mortality with only 20% of *Act5C-GAL4>UAS-edin* flies surviving at 6 days post-eclosion. In contrast, RNAi knockdown of *edin* in the fat body increased susceptibility of flies to *L. monocytogenes* similar to susceptibility observed in the *wntD* mutant.

Overactivation of immune signaling and excessive AMP production has been associated with shortened lifespan and reduced fitness due to AMP damage to self-tissues (Cao et al., 2013; Chinchore et al., 2012; Fink et al., 2016; Lhocine et al., 2008; Libert et al., 2006; Petersen et al., 2013; Rommelaere et al., 2023; Schmidt et al., 2011; Tsuda et al., 2005; Badinloo et al., 2018). A genetic background with elevated AMP expression such as the *wntD* mutant employed in this study may be expected to have a shortened lifespan due to this effect. Independently, we have found that increased AMP expression and susceptibility to *L. monocytogenes* in the *wntD* mutant used in the (Gordon et al., 2008) was due to genetic background (see supplementary S5).

A follow-up study confirmed that Edin is an Attacin paralog regulated by Imd signalling (Vanha-Aho et al., 2012). Multiple studies also show that *edin* is immune responsive to septic and oral infections with Gram-negative bacteria in both larvae and adults (Buchon et al., 2013; Douglas et al., 2011; Kleino et al., 2008; Ramond et al., 2020; Schlamp et al., 2021; Short and Lazzaro, 2013; Troha et al., 2018; Vanha-Aho et al., 2012; Verleyen et al., 2006), although none of these specifically look at immune responsiveness to *L. monocytogenes.*

Subsequent studies have not identified a role for Edin in defense against Gram-negative bacteria similar to other Attacins, instead suggesting mild susceptibility to Gram-positive bacterial infection (Vanha-Aho et al., 2012) and wasp infestation (Vanha-Aho et al., 2015). However, these *edin* studies and (Gordon et al., 2008) only used overexpression or RNAi tools, which were problematically associated with mortality after eclosion (Gordon et al., 2008) or variation in hemocyte differentiation (Vanha-Aho et al., 2015), which can affect immune function. (Vanha-Aho et al., 2012) dismissed Edin antibacterial activity on the basis of lack of activity in S2 cells. However, the pMT-edin-V5 construct used in this study failed to exhibit furin cleavage of the propeptide (Vanha-Aho et al. (2012 - Fig. 1B). Furin cleavage is expected to be required for Edin function based on comparison with similar Attacin/Diptericin-like genes across insects (Hanson and Lemaitre, 2020), although cleavage of the Edin precursor has not yet been confirmed *in vivo* using proteomics approaches. Therefore, this test of the unmaturated precursor protein likely does not reflect its *in vivo* activity

In order to clarify the role of *edin* in *L. monocytogenes* infection, we generated an *edin* knockout mutant and tested survival to septic infection in these flies. We also overexpressed *edin* using a ubiquitous driver and analyzed its impact on longevity.

**Methods**

*Fly lines and husbandry*

We generated a null mutant of Edin (*edin*$^{KO}$) in the isogenic $w^{1118}$ Drosdel background by inserting a DsRed cassette into the *edin* gene coding region using CRISPR-Cas9-mediated homology-directed repair (Figure 1). A pScarless vector containing a 3xP3-DsRed insert flanked by two 1kb homology arms was used as a template for repair. A *Relish*$^{E20}$ (*Rel*$^{E20}$) line was used as an Imd-deficient positive control. Flies were reared at 25°C on standard cornmeal and molasses medium, and flipped to new medium every 2-3 days. The *UAS-edin* construct was made by Vanha-Aho et al., (2015); the Act5C-Gal4 driver is from Bloomington.

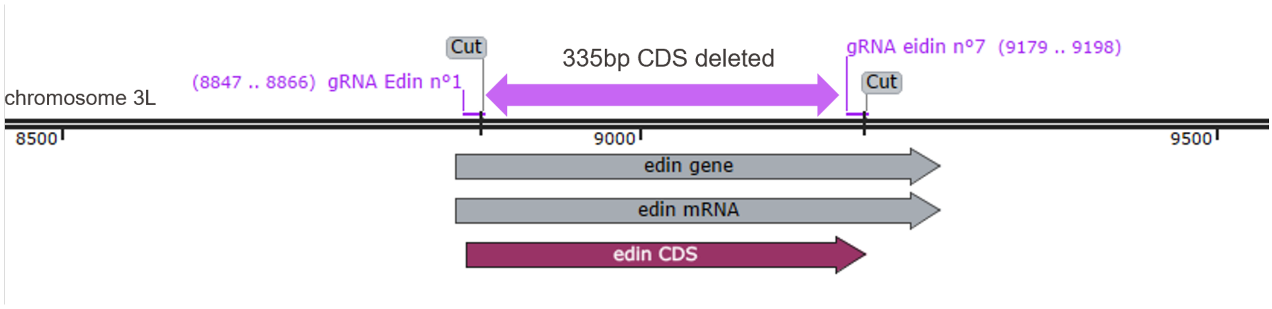

**Figure 1. Description of the *edin*$^{KO}$ mutant**

*Survival and lifespan assays*

5 to 7-day old female flies were anaesthetized with $CO_2$ and pricked on the thorax anterior and ventral to the wing joint with a 0.2 mm tungsten needle dipped in a concentrated overnight (BHI, shaking, 29°C) culture of *L. monocytogenes* (CIP 107776, EGDe) diluted to $OD_{600}$=0.5 in PBS. Flies were incubated at 29°C and scored daily for survival; flies were flipped to new medium during the survival every 2-3 days. ~20 flies per genotype were scored in 2 independent experiments. Lifespans were monitored at 25°C, and flies were flipped every two days.

*RNA extraction and qPCR*

Challenged flies were collected 6h after septic injury with the Gram-negative bacterium *Pectobacterium carotovorum (Ecc15,*OD600=200*),* with incubation at 25°C*.* Total RNA was extracted from pooled samples of 7-10 flies (5-7 days old) each using TRIzol reagent (Invitrogen), and re-suspended in MilliQ water. DNase treatment was performed using TURBO DNA-free kit (Invitrogen). Reverse transcription was performed using 0.5 mg total RNA in 10 µl reactions using Prime-Script RT (Takara) with random hexamer and oligo dT primers. Quantitative PCR was performed on a LightCycler 480 (Roche) in 96-well plates using PowerUp SYBR Green Master Mix (Applied Biosystems). *Edin* expression was normalized to *rp49* (*RpL32*), with the unchallenged condition set to 1.

- *Rp49* Forward: 5' – GCC GCT TCA AGG GAC AGT ATC TG – 3'
- *Rp49* Reverse: 5' – AAA CGC GGT TCT GCA TGA G – 3'
- *Edin* Forward: 5' GTGGCCTCCTATCGTCAACCATATC -3'
- *Edin* Reverse: 5' – ATCGAGCACGGCGTACCAAG -3'

**Results**

Survivals did not reveal any increased susceptibility of *edin*$^{KO}$ flies to infection with *L. monocytogenes* compared to wild-type. As expected, *Relish*$^{E20}$ flies were markedly susceptible to this infection (Figure 2). RT-qPCR confirmed that overexpression of *UAS-edin* with the *Act5C-Gal4* driver leads to a constitutive expression of *edin* (Figure 3A). Lifespan analysis at 25°C of flies overexpressing *edin* with the ubiquitous *Act5C-Gal4* driver was similar to control flies (Figure 3B).

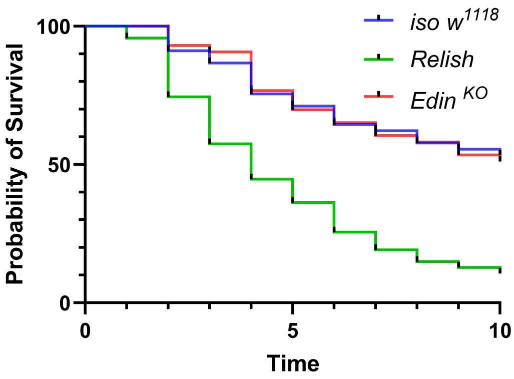

**Figure 2.** Female *edin* null mutant flies are not susceptible to *Listeria monocytogenes* infection ($OD_{600}$=0.5, 29°C) compared to wild type (*iso $w^{1118}$*). n=2 independent experiments, ~20 flies per genotype per replicate.

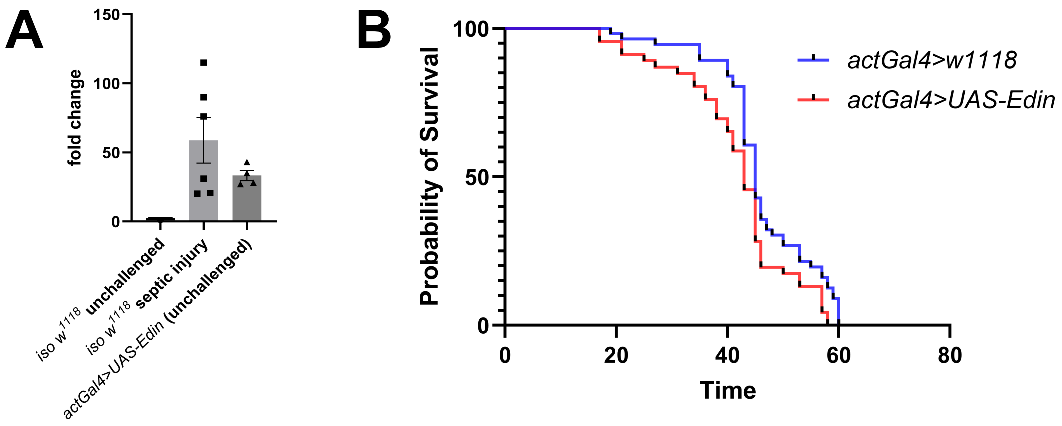

**Figure 3. Over-expression of *edin* does not increase lifespan. A)** Validation of the *UAS-edin* construct with the *Act5C-Gal4* driver at 25°C. Overexpression of *edin* with the *Act5C-Gal4* driver leads to strong expression of *edin* compared to unchallenged flies. Error bars represent the standard deviation. **B)** Lifespan of female flies overexpressing *edin* under the control of the ubiquitous *Act5C-Gal4* driver. The lifespan of female flies carrying the *Act5C-Gal4* driver alone or in combination with a *UAS-edin* construct was analyzed at 25°C. One experiment, n=56 *Act5C-Gal4>$w^{1118}$*, n=43 *Act5C-Gal4>UAS-edin* flies.

#### Conclusion

Experiments performed for ReproSci indicate that contrary to (Gordon et al., 2008), loss of *edin* does not increase susceptibility to *L. monocytogenes* infection, and therefore likely has no appreciable effect on disease tolerance to this bacterium. Since the (Gordon et al., 2008) study was performed using RNAi, the previous result is likely due to an off-target or genetic background effect, although the authors did include appropriate RNAi controls. Furthermore, experiments performed for ReproSci indicate that contrary to (Gordon et al., 2008), overexpression of *edin* does not strongly reduce lifespan.

### Supplementary S15: Listericin in immunity

**Abstract**

(Goto et al., 2010) found using transfection and RNAi in S2 cells that the *Listericin* gene encodes an antibacterial peptide-like protein whose induction is cooperatively regulated by PGRP-LE and the JAK-STAT pathway. The (De Gregorio et al., 2002) microarray indicates that this gene is regulated by the Imd pathway with a minor impact of Toll. Very little subsequent work has examined the role or regulation of *Listericin.*

Experiments performed for ReproSci suggest that *Listericin* expression is largely controlled by the Imd pathway, and does not have a specific dependence on PGRP-LE but rather is likely induced through both PGRP-LC and -LE as other Imd-regulated antimicrobial peptide (AMP)-like genes are. RNAi against *STAT92E* in the fat body did not impair *Listericin* induction *in vivo*, suggesting that contrary to (Goto et al., 2010), the JAK-STAT pathway does not have a strong role in induction of this gene. Finally, we find that overexpression of *Listericin* in the fat body does not affect survival to septic infection with *L. monocytogenes.* We also generated a *Listericin* null mutant and did not find major susceptibility to *Listeria monocytogenesis.*

**Introduction**

(Goto et al., 2010) identified *Listericin* as upregulated in response to *Listeria monocytogenes* in transfected cells expressing YFP-tagged PGRP-LE but not in cells lacking PGRP-LE, in cell culture. RT-PCR verified that *Listericin* was induced by wild-type but not cell-entry deficient *L. monocytogenes*. (De Gregorio et al., 2002) show that expression of *Listericin* is dependent on both Toll and Imd pathways in response to other bacteria, with a greater contribution of the Imd pathway. The dependence on PGRP-LE found by (Goto et al., 2010) is consistent with very low levels of both PGRP-LE and PGRP-LC expression in S2 cells (FlyBase), which act partially redundantly in activation of the Imd pathway (Kaneko et al., 2006; Neyen et al., 2012). Transfection of S2 cells with PGRP-LE is expected to make the response to *L. monocytogenes* PGRP-LE dependent, whereas transfection of the cells with PGRP-LC would likely have made the *Listericin* response PGRP-LC dependent.

(Goto et al., 2010) further found that RNAi of *Stat92E* (key transcription factor of the JAK-STAT pathway) but not of *key* (component of the Imd pathway) or *Dif* and *Dorsal* (transcription factors of the Toll pathway) suppressed induction of *Listericin* in response to *L. monocytogenes,* whereas RNAi of either *key* or *Stat92E* suppressed induction of *Listericin* in response to heat-killed *Escherichia coli*. They concluded that regulation of *Listericin* is Imd-independent specifically in response to *L. monocytogenes*. Furthermore, activation of the JAK-STAT pathway by overexpression of the ligand *upd* (most likely *upd1,* although this is not made clear in the paper) induced *Listericin* expression in PGRP-LE expressing cells but not in PGRP-LE deficient cells.

It is unclear if Listericin displays antimicrobial activity but, in general, AMPs are not regulated by JAK-STAT signaling (except Drosomycin-like peptides in the gut (Buchon et al., 2009)). This apparent dependency on JAK-STAT may be due to the use of a hemocyte-like S2 cell culture system, as the JAK-STAT pathway has strong effects on hemocyte differentiation (see (Banerjee et al., 2019)). Blocking this pathway may be expected to change the nature of the cell culture, for example causing differentiation into lamellocyte-like cells, which have a unique expression profile for AMPs and many other genes (Tattikota et al., 2020). Lack of suppression by either *key* or *Dif/Dorsal* knockdown is likely due to the fact that both Toll and Imd pathways contribute to expression of *Listericin* (De Gregorio et al., 2002). Significantly, the hemocyte RNAseq data of (Tattikota et al., 2020) show that *Listericin* is not expressed in any of the hemocyte populations regardless of immune challenge, suggesting that S2 cells may be a particularly poor model for study of this gene. In contrast to most other AMPs that are strongly expressed in the fat body and hemocytes, *Listericin* is primarily expressed in the adult head (FlyAtlas2).

To determine whether *PGRP-LE* and JAK-STAT are required for expression of *Listericin in vivo* as it (Goto et al., 2010) found in an S2 cell culture system, we tested basal and induced expression of *Listericin* in $w^{1118}$ wild-type flies, in *PGRP-LE* mutants*,* and in flies expressing a STAT 92E RNAi in the fat body*.* We also examined whether *Listericin* confers protection against *Listeria monocytogenes* infection *in vivo* using both overexpression and null mutant flies.

#### Methods

*Fly lines* and *bacterial strains*

All flies were reared on cornmeal fly medium (6% cornmeal, 6% yeast, 0.6% agar, 0.1% fruit juice, 10.6 g/L moldex and 4.9 mL/L propionic acid) at 25°C. *PGRP-LE*$^{112}$ (*yw* background) loss-of-function mutants were used (Takehana et al., 2004). qPCR confirmed lack of expression of *PGRP-LE* in the *PGRP-LE*$^{112}$ mutant. The *PGRP-LE*$^{112}$ mutant was validated in Figure 2C. *Listericin*$^{1}$ and *Listericin*$^{2}$ were generated by replacing a 340-bp coding sequence of the *Listericin* gene with a 1.7-kb *dsRed* cassette using CRISPR/Cas9 (Figure 1, Figure 4D). For RNAi experiments, *Lpp-Gal4*$^{ts}$ (Brankatschk et al., 2018), *c564-Gal4* (BDSC #6982), or *w*$^{1118}$ (BDSC #3605) flies were crossed to *STAT92E-RNAi* or *UAS-Listericin* (BDSC #33056, II and #33058, III) flies at 25°C. *Lpp-Gal4*$^{ts}$ crosses were moved to 29°C after egg laying to promote driver activity. The strains *Listeria monocytogenes* (CIP 107776, EGDe) and *Pectobacterium carotovorum Ecc15* were used in this study.

*qPCR for Listericin induction*

Male flies were pricked with concentrated overnight cultures of *Ecc15* ($OD_{600}$=200) or *L. monocytogenes* ($OD_{600}$=0.1) diluted to the desired concentration in PBS and maintained on cornmeal fly medium at 25°C. 7-10 flies per treatment were frozen at -80°C at 8h post infection. Two independent experiments were performed. Total RNA was extracted from samples using TRIzol (Invitrogen), and re-suspended in MilliQ water. DNase treatment was performed using TURBO DNA-free kit (Invitrogen). Reverse transcription was performed using 0.5 mg total RNA in 10 ul reactions using Prime-Script RT (Takara) with random hexamer and oligo dT primers. Quantitative PCR was performed on a LightCycler 480 (Roche) in 96-well plates using PowerUp SYBR Green Master Mix (Applied Biosystems). *Listericin* expression was normalized to expression of the ribosomal protein *rp49.* Data were analyzed in Prism 10 (GraphPad) by t-test.

- *Rp49* Forward: 5' – GCC GCT TCA AGG GAC AGT ATC TG – 3'
- *Rp49* Reverse: 5' – AAA CGC GGT TCT GCA TGA G – 3'
- *Listericin* Forward: 5' – CAG TAC CTG GTG CTT GCC CTC – 3'
- *Listericin* Reverse: 5' – CTT CAG CAT CTT CCA GCG TGA TCT – 3'
- *PGRP-LE* Forward: 5' – AAG CTC CTC AGT TTT ATA CGT CA – 3'
- *PGRP-LE* Reverse: 5' – CGC TGT TCT TCC CTC CAA GT – 3'
- *Turandot A* Forward: 5' – TCT TCA ACT GCT CTT ATG TGC TTT GC – 3'
- *Turandot A* Reverse: 5' – TCT AAG GTT GTC AGC CTC ACG ATC – 3'

*Survivals*

3 to 5-day old male flies were anaesthetized with $CO_2$ and pricked on the thorax anterior and ventral to the wing joint with a 0.2 mm tungsten needle dipped in a concentrated overnight culture of *P. carotovorum Ecc15* (shaking, LB, 29°C, $OD_{600}$=200) or *L. monocytogenes* (shaking, BHI, 37°C, $OD_{600}$=0.1) diluted to the desired concentration in PBS. Flies were incubated at 29°C and scored daily for survival; flies were flipped to new medium during the survival every 2-3 days. At least 20 flies per genotype were scored in 2-3 independent replicates per genotype. Data were analyzed in Prism 10 (GraphPad) using a Log-rank test.

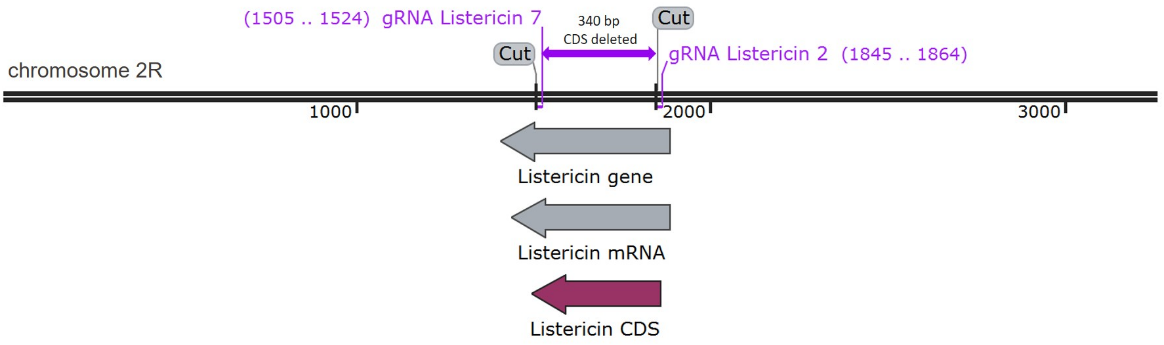

**Figure 1 Description of the *Listericin*$^{\Delta1}$ and *Listericin*$^{\Delta2}$ mutants.** The two mutations were obtained by deleting the Listericin coding region with two gRNA as indicated, which was replace by 1.7-kb 3xP3-*dsRed* cassette.

#### Results and Discussion

*Listericin expression is not dependent on PGRP-LE or JAK-STAT*

Contrary to (Goto et al., 2010), we did not find that loss of PGRP-LE strongly impaired expression of *Listericin in vivo* in adult flies upon systemic infection with the Gram-negative bacterium *Ecc15* (Figure 2A). We were also unable to find any differences in induction of *Listericin* by *L. monocytogenes* as found by (Goto et al., 2010) (Figure 2A-B). This is consistent with a model where *Listericin* expression is Imd dependent and PGRP-LE and PGRP-LC function as partially redundant pattern recognition receptors (PRRs) in expression of this peptide as they do for other Imd-dependent AMPs. Microarrays have also shown that both Toll and Imd pathways contribute to expression of *Listericin* (De Gregorio et al., 2002). The results seen by (Goto et al., 2010) may have been due to use of an S2 cell culture system with non-representative expression of immune proteins including the major Imd pattern recognition receptors PGRP-LE and PGRP-LC.

Contrary to (Goto et al., 2010), in our hands reduction of JAK-STAT activity through RNAi of the JAK-STAT transcription factor STAT92E did not impair expression of *Listericin* in adult flies upon systemic infection with *Ecc15* compared to an appropriate control (Figure 3A,B). The STAT92 RNAi construct was validated by revealing its impact on Turandot A gene expression (Figure 3C-D) as previously shown (Agaisse et al., 2003)

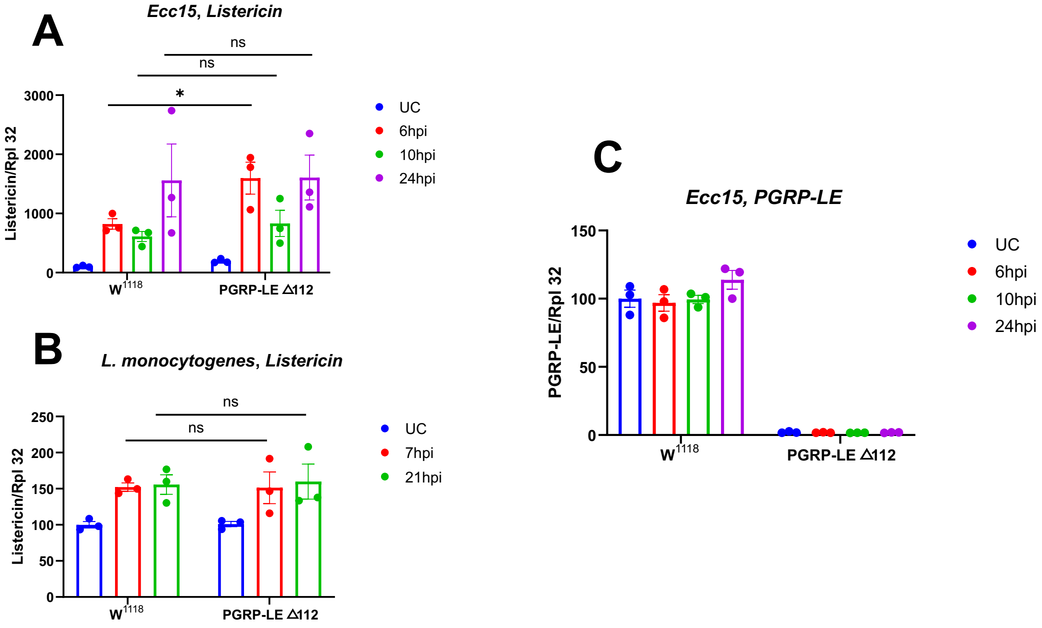

**Figure 2. The expression of *Listericin is not affected by PGRP-LE.*** Expression of *Listericin* in a *PGRP-LE*$^{112}$ mutant compared to wild-type following septic infection with **(A)** *Ecc15* ($OD_{600}$=200) at

25°C or **(B)** *L. monocytogenes* ($OD_{600}$=0.5) at 29°C. **(C)** Validation of *PGRP-LE*$^{112}$ mutant. 10 flies per replicate, 3 experimental repeats. Error bars represent s.e.m. Statistical analysis: Tukey's multiple comparisons test, not significant = ns, p<0.05 = *. Results are from three independent experiments.

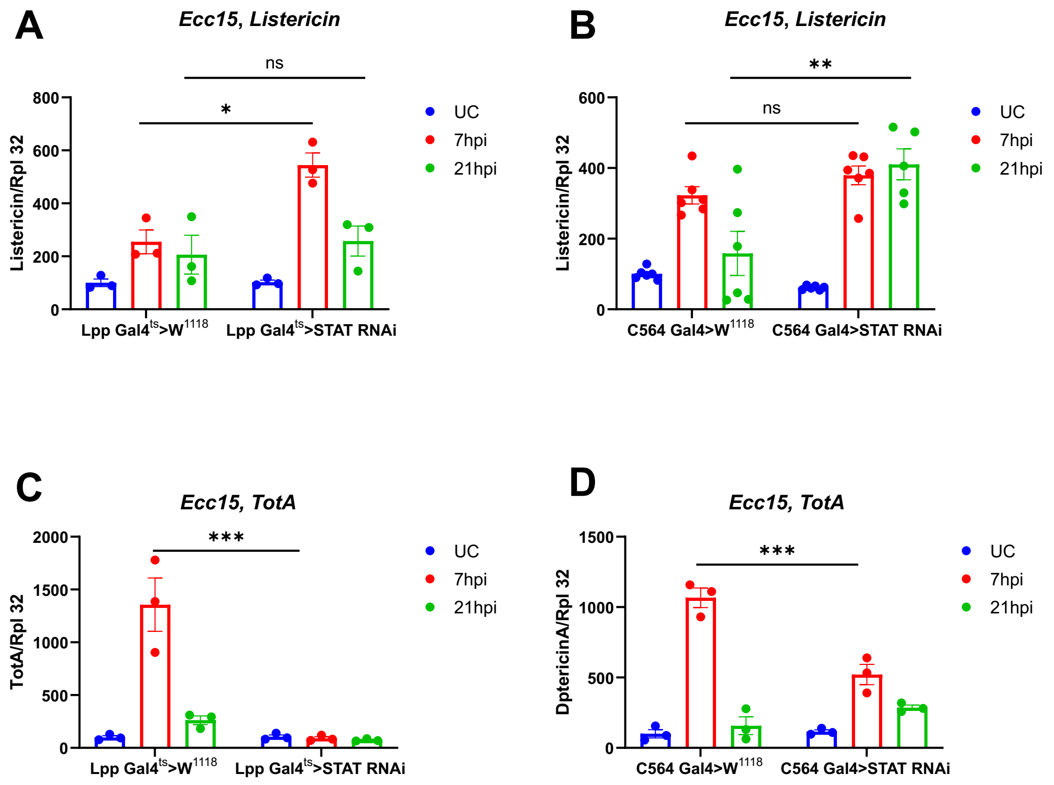

**Figure 3. Listericin expression is not affected by the JAK-STAT pathway.** Expression of *Listericin* in flies with knockdown of JAK-STAT activity in the fat body compared to driver-only controls, following septic infection of 3-5 day-old male flies with *Ecc15* ($OD_{600}$=200). Two independent fat body drivers were used. *Lpp-gal4*$^{ts}$ crosses were moved to 29°C after egg-laying to activate the temperature-sensitive driver **(A)**, while *c564-Gal4* crosses were maintained at 25°C **(B)**. **(C and D)** *TotA* (*Turandot A*) expression was used as a readout of JAK/STAT pathway activity to validate RNAi efficiency. 10 flies per replicate, 3-6 experimental repeats. Statistical analysis: Error bars represent s.e.m. Tukey's multiple comparisons test, not significant = ns, p<0.05 = *, p<0.01 = **, and p<0.001 = ***.

Listericin *does not have antibacterial effects against Gram-negative bacteria* in vivo

(Goto et al., 2010) found that medium from *Listericin* overexpressing cells suppressed growth of *L. monocytogenes* and *E. coli* but not *Staphylococcus saprophyticus.* Overexpression of *Listericin* in the hemocytes and fat body (using *Cg-Gal4*) restored survival to *L. monocytogenes* in a *PGRP-LE*$^{112}$ mutant and in wild-type flies. Overexpression of *Listericin* reduced bacterial loads of Gram-negative bacteria (*L. monocytogenes, E. coli*) but not Gram-positive bacteria (*Staphylococcus aureus, S. saprophyticus*) in the hemolymph of infected wild-type flies.

Experiments performed for ReproSci show no effect of *Listericin* overexpression on survival to *L. monocytogenes* (Figure 4A, B). Moreover, flies carrying a null mutation in *Listericin* did not exhibit increased susceptibility to *L. monocytogenes* infection (Figure 4C) suggesting that this AMP-like gene does not have a prominent role in survival to this bacterium *in vivo.*

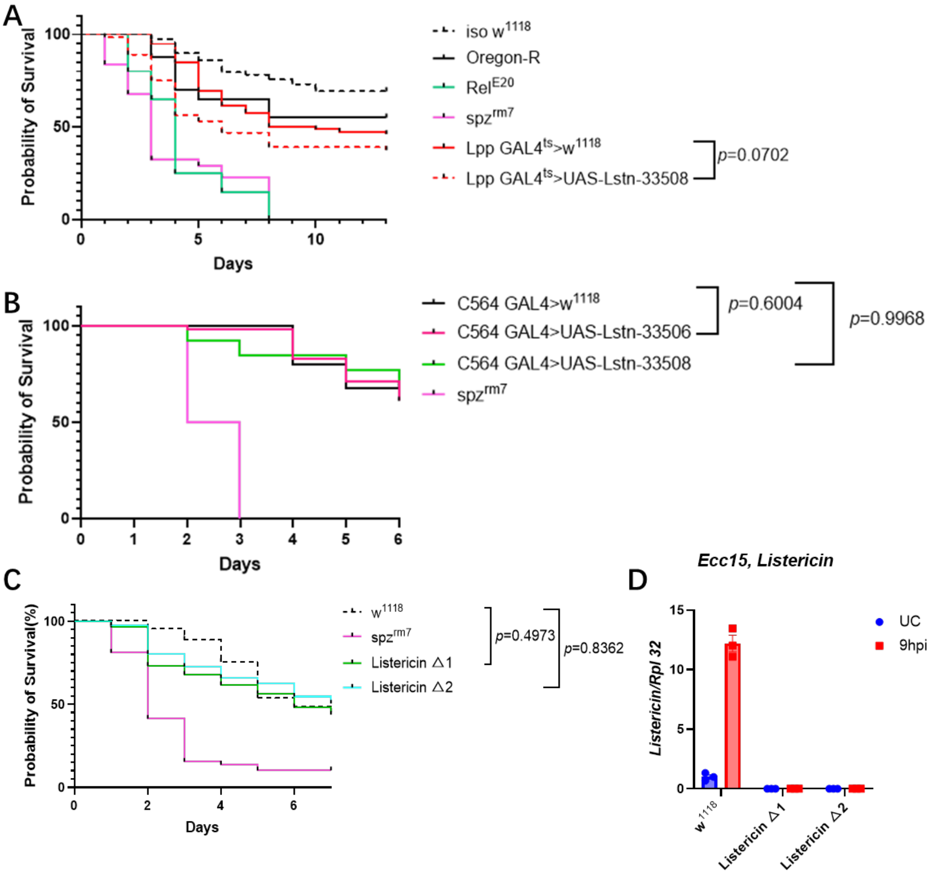

**Figure 4. Survival of flies overexpressing *Listericin* in the fat body compared to driver-only controls (A, B)** and *Listericin* null mutants **(C)** following septic infection of 3-5 day-old male flies with *L. monocytogenes* ($OD_{600}$=0.1) at 29°C. Two independent fat body drivers and two independent *UAS-Listericin* constructs were used. **(D)** Validation of *Listericin* mutants by qPCR. 20 flies per replicate, 2 experimental repeats. Statistical analysis: Log-rank test.

#### Conclusion

Contrary to (Goto et al., 2010), we find that *Listericin* expression is not specifically dependent on PGRP-LE, and is not dependent on JAK-STAT signaling in flies, although this could be the case in S2 cells where the initial experiments were done. Evidence indicates that that this gene is regulated by the Imd pathway with a minor impact of Toll (De Gregorio et al., 2002). Furthermore, survival experiments show that overexpression of *Listericin* is not protective against *L. monocytogenes* infection as initially found by (Goto et al., 2010). These conclusions may have been made due to a combination of use of an S2 cell culture system that does not fully reflect *in vivo* immune function, and genetic background effects that were not fully controlled for.

### Supplementary S16: Role of Dscam1 in Drosophila immunity

**Abstract**

Previous results suggested that secreted Dscam1 plays an active role in immunity by binding bacteria in the hemolymph and enhancing phagocytosis (Watson et al., 2005). Contrary to this, results from ReproSci and more recent literature do not find evidence for secreted isoforms of Dscam1 in *Drosophila*. ReproSci does find impaired phagocytic uptake upon *Dscam1* depletion. This impaired phagocytosis may be due to reduced hemocyte numbers, as found in other studies.

**Introduction**

Although there are four *Drosophila* '*Dscam*' genes, only *Dscam1* is considered a "hypervariable Dscam" (Ng and Kurtz, 2020) that has the potential to produce thousands of different isoforms through alternative exon splicing (Armitage et al., 2014; Celotto and Graveley, 2001). (Watson et al., 2005) found using Western Blots and LC-MS/MS that a ~160kDa isoform of Dscam1 corresponding to a truncated ectodomain was present in extracts of larval hemolymph and S2 cell culture media, and that dozens of endogenous isoforms of Dscam1 were present in both intracellular and secreted forms. Using FACS, (Watson et al., 2005) found that Dscam1 isoforms with specific exon combinations were able to bind heat-killed *Escherichia coli* and that RNAi knock down of *Dscam* impaired phagocytic uptake of *E. coli* in larvae. Based on these results (Watson et al., 2005) concluded that binding of hemolymph isoforms of Dscam1 to *E. coli* enhances phagocytosis through opsonization. A similar role for Dscam proteins has been studied in crustaceans, where heterophilic binding of hemolymph Dscam to pathogens can enhance binding and uptake by immune cells (Ng and Kurtz, 2020).

The majority of research on *Drosophila Dscam* genes has focused on their roles in neurons. Homophilic binding of Dscam1 isoforms mediates neurite repulsion in the CNS (Millard et al., 2010) and (Ouyang et al., 2020) found that Dscam1 is required for development and maintenance of neuronal microenvironments that promote haematopoiesis and hemocyte maintenance at the larval stage. *Dscam1* is not upregulated following infection in *Drosophila* (Armitage et al., 2014; Ramond et al., 2020), and infection does not induce any change in the expression pattern of *Dscam1* isoforms (Armitage et al., 2014), contrary to what has been shown in crustaceans (e.g. (Li et al., 2019)). No published results have followed up on roles for *Dscam1* in *Drosophila* immunity.

Soluble isoforms of Dscam have been detected in shrimps (Chou et al., 2009) and crabs (Wang et al., 2013) where a role for Dscam in opsonizing hemolymph pathogens has been better studied (Ng and Kurtz, 2020). In *Drosophila*, Dscam1 is better studied in a neuronal context, and is typically discussed as a transmembrane protein (Matthews et al., 2007; Millard et al., 2010; Schmucker et al., 2000; Yu et al., 2009). The exons encoding the transmembrane domain are constitutive parts of all *Dscam1* transcripts, while the extracellular domains are variably spliced to produce thousands of isoforms that are developmentally and spatially regulated (Celotto and Graveley, 2001). (Armitage et al., 2014) similarly found that many isoforms of *Dscam1* are expressed in *Drosophila,* although they did not examine tissue-specific expression. High throughput expression data show that *Dscam1* is overwhelmingly expressed in the CNS, head and ganglia, but is expressed at low but detectable levels in other tissues, including the fat body (FlyAtlas2). Recent RNAseq data show that *Dscam1* mRNA is slightly enriched in the larval hemocytes compared to whole body expression, and that expression of *Dscam1* isoforms in the hemocytes is reduced by infection (Ramond et al., 2020). However in contrast to (Watson et al., 2005), soluble isoforms of Dscam1 were not detected in a *Drosophila* adult hemolymph proteome upon challenge (Rommelaere et al. 2025). These data suggest that Dscam1 isoforms primarily exist in a transmembrane configuration in *Drosophila*. ReproSci concludes that there is no strong evidence for secreted isoforms of Dscam1 in *Drosophila,* and this argues against an opsonizing role for *Dscam1* in *Drosophila* despite research showing that Dscam acts in this capacity in crustaceans.

To determine whether the effect of *Dscam1* depletion on phagocytosis is replicable, we repeated phagocytosis experiments in *Dscam1*-depleted larvae using an alternate RNAi construct.

#### Methods

Flies bearing a hemocyte driver (*hmlD-Gal4*, BDSC #30139) were crossed to a *Dscam1* RNAi construct line (BDSC #38945) or $w^{1118}$ wild-type flies (BDSC #3605) at 29°C for optimal driver activity. Phagocytic index was measured and calculated in larval hemocytes as previously described (Melcarne et al., 2019). Six larvae per genotype were bled per genotype per experiment, and three biological replicates were performed. Statistical differences were calculated by t-test.

#### Results and Discussion

*Dscam1 deficiency has a mild effect on phagocytosis of* E. coli

(Watson et al., 2005) found that a heteroallelic mutation of *Dscam1*, RNAi against *Dscam1* in hemocytes, and blocking of Dscam1 with antibodies in S2 cells all impaired uptake of heat-killed *E. coli* by ~40%, and that certain labelled isoforms of Dscam1 were able to bind *E. coli* in cell culture. Based on these results (Watson et al., 2005) concluded that binding of soluble isoforms of Dscam1 to *E. coli* enhances phagocytosis. Experiments performed as part of ReproSci similarly show that knockdown of *Dscam1* in the hemocytes (*hml>Dscam1-RNAi*) reduces phagocytosis of *E. coli* (~50%, p<0.5) but not *S. aureus* bioparticles (~30% reduction, nonsignificant) (Figure 1). While these data support the initial observations of (Watson et al., 2005), it is not sufficient to demonstrate a direct or opsonizing role of Dscam1 in phagocytosis (see Introduction).

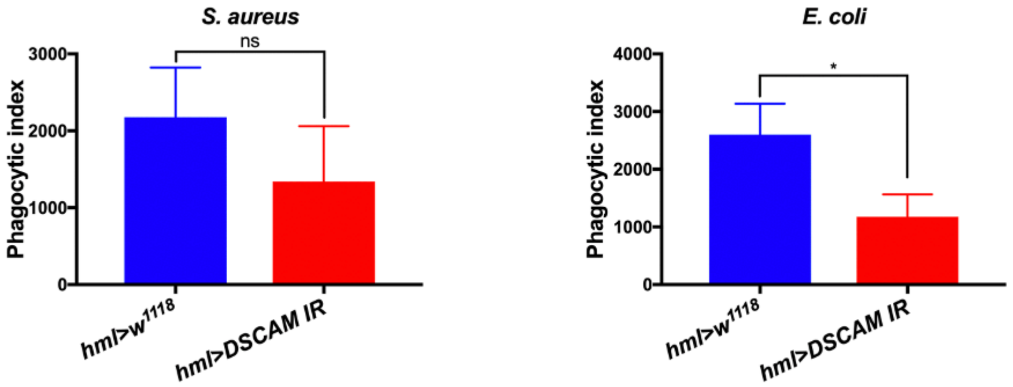

**Figure 1.** Knockdown of *Dscam1* in larval hemocytes (*hml>Dscam1-RNAi*) reduces phagocytosis of *E. coli* (~50%, p<0.5) but not *S. aureus* bioparticles (~30% reduction, nonsignificant). Error bars represent the standard deviation. Results were analyzed by T-test.

#### Conclusion

Experiments done for ReproSci confirm that depletion of *Dscam1* slightly impairs phagocytosis of *E. coli* as suggested by (Watson et al., 2005). (Ouyang et al., 2020) found that Dscam1 may be involved in development of neuronal microenvironments required for hemocyte maintenance at the larval stage (Gold and Brückner, 2015; Makhijani et al., 2017, 2011). (Vlisidou et al., 2009) found that at the embryonic stage, *Dscam1* null hemocytes recognize and phagocytose *E. coli* similar to wild-type hemocytes, which similarly argues against a direct role of Dscam1 in phagocytosis. Furthermore, the predicted presence of a transmembrane domain in all *Dscam1* isoforms (Celotto and Graveley, 2001) and the lack of secreted isoforms of Dscam1 in hemolymph of adult *Drosophila* (Rommelaere et al., 2025) argue against an opsonizing role for Dscam1 isoforms in *Drosophila.* Given these lines of evidence, we believe that the impact of Dscam1 on phagocytosis is indirect, and likely explained by a defect in hematopoiesis as found by (Ouyang et al., 2020).

### Supplementary S17: Role of Croquemort in phagocytosis of apoptotic cells

**Abstract**

*Croquemort* (*crq*) encodes one of many transmembrane CD36 homologs present in the *Drosophila* genome (class B scavenger receptors) that typically have functions related to recognition and transport. Croquemort has long been implicated in both phagocytosis of apoptotic cells and bacteria (Franc et al., 1999, 1996; Stuart et al., 2005), but the specific functions of this protein are unclear due to variable results and imprecise descriptions of study outcomes. Experiments performed for ReproSci find no clear role for *croquemort* in phagocytosis of apoptotic cells at the embryonic stage. These results and a review of the literature suggest that Croquemort has a negligible role in phagocytosis of apoptotic cells and bacteria at all life stages, but promotes degradation of lysosomal cargo including nonapoptotic cell debris and bacteria in both macrophages and phagocytic epithelial cells.

**Introduction:**

Results suggesting a role for *croquemort* in phagocytosis of apoptotic cells and bacteria have varied greatly in methodology. Because of this, it is difficult to compare data across studies. These issues include ecdysone status of the model used (S2 cells, mbn-2 cells, larvae), which affects expression of *croquemort* and other receptors that may influence phagocytosis (Manaka et al., 2004; Rus et al., 2013). All but one study that presented evidence for reduced phagocytosis of 'apoptotic cells' in *croquemort*-deficient cells used stains or markers that more generally label fragmented cells or debris, as opposed to markers that specifically label apoptotic cells such as Caspase 3/Dcp-1 or Acridine Orange.

(Franc et al., 1999) found that loss of *croquemort* in embryos resulted in a strong 90% reduction in engulfment of 7AAD-stained bodies. 7AAD binds non-viable cells including strong labelling of dendrite debris (Hilpert et al., 2005), and is not exclusive to apoptotic cells. Later, results of (Han et al., 2014) revealed that *croquemort* is required for removal of dendrite debris and CNS development. (Han et al., 2014) visualized engulfed material using dendrite-specific cell surface markers, rather than apoptotic markers such as Acridine Orange, which does not stain dendrite debris (Mai et al., 1984). The results of (Han et al., 2014) suggest that the 'apoptotic cells' measured by (Franc et al., 1999) largely represented dendritic fragments rather than classical apoptotic cells. Similarly, (Meehan et al., 2016) found no role for *croquemort* in phagocytosis of Dcp1-stained apoptotic cells in Croquemort-enriched follicle cells of adult ovaries. Using RNAi in ecdysone-treated l(2)mbn cells, (Manaka et al., 2004) found no role for *croquemort* in phagocytosis of apoptotic bodies generated by treating S2 cells with caspase inhibitor and cycloheximide. Importantly, the results of (Zheng et al., 2021) regarding phagocytosis of apoptotic cells might be discarded due to methodological issues (see additional discussion on ReproSci website). The sole study that finds a role for *croquemort* in phagocytosis of true apoptotic cells is (Cuttell et al., 2008), whose data indicate that ecdysone-activated S2 cells treated with RNAi against *croquemort* showed a relatively mild ~20% reduction in engulfment of actinomycin D-treated S2 cells after a lengthy 5h incubation period (Cuttell et al., 2008 - Figure S3).

Several studies have shown that loss of *croquemort* causes a mild reduction in uptake of *Staphylococcus aureus* (Cuttell et al., 2008; Franc et al., 1999; Guillou et al., 2016; Han et al., 2014 ; Melcarne et al., 2019; Stuart et al., 2005). A reduction in uptake of *Escherichia coli* upon loss of *croquemort* is less consistent in published data, with only (Guillou et al., 2016) showing a strong reduction in phagocytosis of this bacterium. However, (Guillou et al., 2016; Han et al., 2014; Petrignani et al., 2021) show that defects in internalization of bacteria and debris only appear at late time points as a result of impaired lysosomal degradation, and that initial uptake of bacteria is not impaired. This bloating and impairment of phagocytosis at late time points due to defects likely explains some of the observed variability in phagocytic uptake of bacteria in *croquemort*-deficient cells across the literature.

In light of these issues, we undertook experiments to verify whether loss of *croquemort* results in defects in phagocytosis of apoptotic cells.

#### Methods

*Fly lines and husbandry*

The *crq*$^{KO}$ line generated by (Han et al., 2014) was verified by PCR using the primers below. The *w*$^{1118}$ line (BDSC #3605) was used as a wild-type comparison. Flies were maintained at 25°C on cornmeal and molasses medium and flipped every 2-3 days to fresh medium.

*Apoptotic cell engulfment*

Macrophages of stage 16 (13-16 hours) embryos were labeled with *srpHemo-3xmCherry* and apoptotic particles immunostained with anti-cleaved Caspase 3 (Dcp-1) antibodies. Ten random well-separated macrophages in each embryo were selected, and the volume of apoptotic particles inside the macrophages was determined using Imaris software and a Fluorescent Apotome Zeiss. The volume of apoptotic cells outside the macrophages in a selected area of every embryo (n=7), not including the particles inside the macrophages, was also measured.

#### Results and Discussion

*Loss of* Croquemort *does not impair apoptotic cell engulfment in embryos*

Experiments performed for ReproSci show that uptake of apoptotic cells stained with Dcp-1 was not impaired in hemocytes of *crq*$^{KO}$ embryos, and that apoptotic cells did not accumulate in these hemocytes (Figures 1, 2). This result is consistent with previously published results when methodology is accounted for, and contradicts the original claim that *croquemort* is implicated in phagocytosis of apoptotic cells.

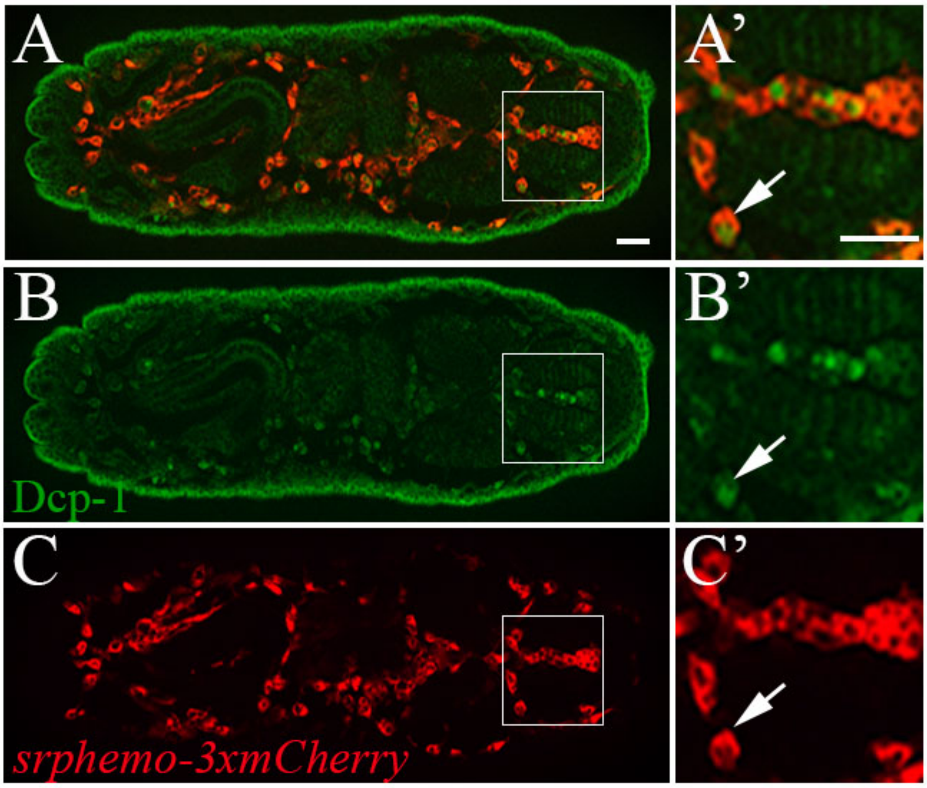

**Figure 1. Evaluation of apoptotic cell clearance in the *crq*$^{KO}$ embryo.** (A,B,C) Dorsal view of stage 16 embryo. Apoptotic cells are labeled with anti-Dcp-1 (green) and macrophages with srphemo-3xmCherry (red). (A',B',C') Close up of the rectangle area, arrows point at engulfed apoptotic cell inside macrophage. Bar 20 µm.

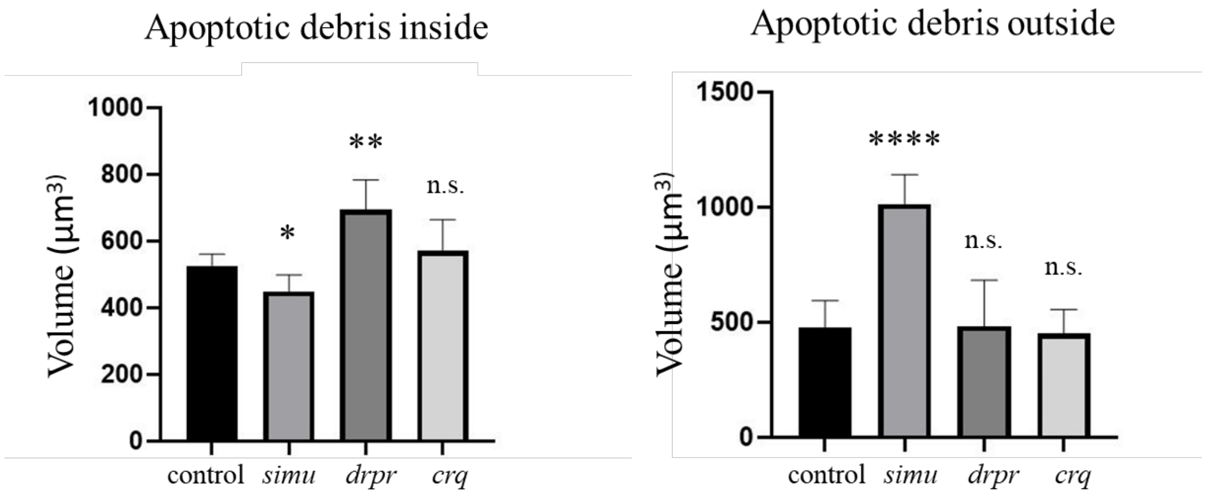

**Figure 2.** Lack of *croquemort* does not affect the volume of apoptotic cells inside (**left panel**) or outside (**right panel**) embryonic macrophages. Columns represent mean volume of apoptotic cells inside or outside of macrophages ± SEM of control, *simu* (*NimC4*$^{KO}$), *drpr* (*drpr*$^{\Delta 5}$) and *crq* (*crq*$^{KO}$) mutants. N = 7 stage 16 embryos of each genotype. Statistical significance versus control, as determined by one-way ANOVA followed by Bonferroni post hoc test, **$p < 0.01$, *$p < 0.05$, n.s. $p > 0.05$.

#### Conclusion

Overall, our results show that contrary to (Franc et al., 1999, 1996; Stuart et al., 2005), loss of *croquemort* does not affect uptake of apoptotic cells by hemocytes. More recent literature suggests that *croquemort* is involved in debris clearance and sustained uptake of cargo including bacteria through a role in lysosomal maturation (Guillou et al., 2016; Han et al., 2014) Melcarne et al., 2019; Petrignani et al., 2021) but has a nonessential or minor role in phagocytosis of apoptotic cells and bacteria. Because initial uptake is not affected, Croquemort does not act as a phagocytic receptor for apoptotic cells or bacteria as suggested by (Franc et al., 1999, 1996; Stuart et al., 2005). Interestingly, (Meehan et al., 2016) found that *croquemort*-deficient ovaries exhibited some defects in cell death, suggesting that *croquemort* may function in nonapoptotic ‘pruning’ of cells in the adult ovaries similar to its role in dendrite pruning during development (embryo/larva/pupa).

When carefully considered with attention to methodology, published results form a consensus supporting an essential role for *croquemort* in sustained phagocytosis of cell debris and specifically dendrite debris at the embryonic, larval and pupal stages. The data indicate that *croquemort* is required in this capacity at multiple life stages, and in at least two cell types: phagocytic epithelial cells in embryos and ecdysone-activated macrophages in larvae and pupae.

### Supplementary S18: Role of the scavenger receptor SR-CI in in hemocyte binding to bacteria and phagocytosis

**Abstract**

Early cell culture studies using RNAi suggested a role for the scavenger receptor SR-CI in hemocyte binding to Gram-negative and Gram-positive bacteria (Rämet et al., 2001). Using an *SR-CI*$^{SK6}$ null mutant and a $\Delta$*(Sr-CI,Sr-CIII)* double mutants, we could observe a reduced ability to bind *E. coli* and *S. aureus* consistent with the main claim of the article. Sr-CI has often been considered as a phagocytic receptor (Melcarne et al., 2019; Ulvila et al., 2011), we were unable to find any consistent effect of this mutation on phagocytosis of *E. coli* and *S. aureus* by larval hemocytes. Collectively, our study conclude that SR-CI contribute to hemocyte binding to bacteria but also revealt that it should not be considered as a major phagocytic receptor for bacteria in *Drosophila.*

**Introduction**

Early cell culture studies suggested a role for *SR-CI* as a pattern recognition receptor involved in the binding to bacteria. (Rämet et al., 2001) in a series of experiments in *Drosophila* S2 and mammalian cells found that expression of *SR-CI* conferred binding of *E. coli* and *S. aureus*. Furthermore, dsRNA knockdown of *SR-CI* in S2 cells caused reduced binding of both *Staphylococcus aureus* and *Escherichia coli* (20-30% reduction). These authors concluded that SR-CI is a macrophage pattern recognition receptor required for optimal binding and raised the hypothesis that it could play a role in phagocytosis. (Saleh et al., 2006) in a similar dsRNA-based study found that knockdown of *SR-CI* strongly decreased uptake of fluorescent *E. coli* by S2 cells. (Ulvila et al., 2006) in another RNAi-based in cell culture study found that *SR-CI* and *eater* together accounted for 90% of dsRNA endocytosis in S2 cells, and that transfection of *SR-CI* in mammalian cells was sufficient to confer endocytosis of dsRNA fragments. Following these initial characterizations, this receptor has received little attention and its role in bacterial binding and phagocytosis was not investigated with the use of null mutations.

At all life stages, *SR-CI* is strongly expressed in hemocytes, but not in non-hemocyte cell populations (Pearson et al., 1995; Tattikota et al., 2020; FlyAtlas2). Several cell culture studies have shown that SR-CI has a role in binding a variety of ligands, and particularly acetylated low-density lipoproteins (Pearson et al., 1995; Rämet et al., 2001; Ulvila et al., 2006). The fact that SR-CI can bind a broad range of ligands led to speculation that it may participate in a variety of processes such as pathogen recognition and phagocytosis, cell-cell or cell-matrix adhesion, wound healing, and apoptotic cell clearance.

Recent expression data show that *SR-CI* is somewhat expressed in all hemocyte types (plasmatocyte, crystal cell, lamellocyte), but is expressed in nearly all cells of the PM12 (cytochrome c-oxidase *cox8*-expressing), PM9 and PM10 plasmatocyte populations (which express the protooncogenes *Abl*, *Sno oncogene* (*Snoo*), and the transcriptional regulator *Ubx*) that appear upon wounding (Tattikota et al., 2020). *SR-CI* transcripts were least enriched in the PM6 population (35.8%), which highly expresses genes coding for the Cecropin family.

*SR-CI* belongs to a family of four scavenger receptor genes, of which *SR-CIII* and *SR-CIV* are also expressed in hemocytes (Tattikota et al., 2020). *SR-CI* and *SR-CIII* are clustered on the chromosome suggesting that they emerged from a recent duplication. These genes may have overlapping functions, which may complicate functional characterization.

Here, we generated a *SR-CI* null mutant (*SR-CI*$^{SK6}$ ) and a small deficiency removing both *SR-CI* and *SR-CIII* ($\Delta$*(Sr-CI,Sr-CIII)*) using a CRISPR-Cas9 approach to investigate the role of SR-CI in bacterial binding, phagocytosis and immunity.

**Methods**

*Fly lines*

The *SR-CI*$^{SK6}$ allele was generated as previously described (Kondo and Ueda, 2013) in the *yw* background using a GTGGCGGGGTCCCGTTCTTA gRNA, deleting base pairs 623-638 of the gene region and resulting in a frameshift (see Figure 1A). The mutation was subsequently isogenized into the $w^{iso}$ *Drosdel* background through 8 generations of backcrossing to generate the *w iso SR-CI*$^{SK6}$ line. Experiments were performed by comparing *w iso SR-CI*$^{SK6}$ to $w^{iso}$ background with appropriate controls. We also generated a small deficiency that removed both *SR-CI and its flanking gene Sr-CIII (Δ(Sr-CI,Sr-CIII)) in the w* iso background (Figure 1B).

Flies were maintained at 25°C on cornmeal fly medium (6% cornmeal, 6% yeast, 0.6% agar, 0.1% fruit juice, 10.6 g/L moldex and 4.9 mL/L propionic acid) and flipped every 2-3 days to fresh medium.

- *SR-CI*$^{SK6}$ mutant primers:
  - Forward Primer: 5'- CGA TAT TCA CCC CGC CA -3'
  - Reverse Primer: 5'- CGG ACA CCA CCA GAT TAT CC -3' (same as wild type)
  - Annealing temperature: 66°C
  - Extension time: 30s
  - No amplification in wild type at this temperature. Cross-amplifies wild-type gene at lower temperatures
- *SR-CI* wild type primers:
  - Forward Primer: 5'- ATA TTC ACT CCC TAA GAA CG -3'
  - Reverse Primer: 5'- CGG ACA CCA CCA GAT TAT CC -3' (same as mutant)
  - Annealing temperature: 58°C
  - Extension time: 30s
  - No amplification in mutant at this temperature.

*Cold-binding assay*

To assess bacterial binding, hemocytes were isolated by bleeding wandering third-instar larvae into 130 µL cold Schneider's insect medium on pre-chilled glass slides. Approximately, $10^5$ fluorescent bacteria bioparticles(from Thermo Fisher Scientific) were added immediately after bleeding, and samples were incubated on ice for 60 min to allow binding while minimizing internalization. Hemocytes were fixed in 4% paraformaldehyde (PFA) for 15 min at room temperature. Following fixation, cells were washed three times with 1× PBS to remove unbound bacteria and debri. Hemocytes were stained with Phalloidin and imaged using an Olympus FV4000 confocal microscope with a 60× objective. Image analysis was performed in Fiji, and the number of bound bacteria per hemocyte was quantified by scoring 100 hemocytes per genotype in each experiment.

*Phagocytosis assays and microscopy*

For ex vivo phagocytosis assays, five third-instar larvae were surface-sterilized and bled into 130 µL of Schneider's medium within 2 minutes as in (Melcarne et al., 2019).. The hemolymph was collected in Protein Lo-Bind Eppendorf tubes to minimize sample loss. Approximately, $10^5$ bacterial bioparticles were added to the hemocyte suspension and incubated at room temperature for 1 hour. Following incubation, the samples were placed on ice, and hemocytes were immediately analyzed using a CytoFLEX flow cytometer (Beckman Coulter).
The phagocytic index (PI) was calculated using the following equations:The phagocytic index (PI) was calculated using the following equations:

$$f = \frac{\text{Number of hemocytes in fluorescence positive gate}}{\text{Total number of hemocytes}}$$

$$PI = \text{Mean fluorescence intensity of hemocytes in fluorescence positive gate} * \text{f}$$

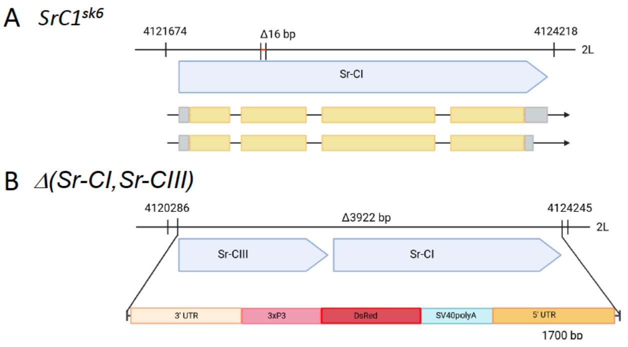

**Figure 1: Description of the *SR-CI*$^{SK6}$ and *Δ(Sr-CI,Sr-CIII)* mutants. A**. The *SR-CI*$^{SK6}$ mutation is a deletion of 16bp causing a frameshift.  was generated. **B**.  *Δ(Sr-CI,Sr-CIII)*  was obtained by deleting the *Sr-CI-Sr-CIII* genomic region which was replaced by 1.7-kb 3xP3-*dsRed* cassette.

#### Results

*Mutations in SR-CI reduced hemocyte binding to E. coli and S. aureus*

We first examined the role of SR-CI in the ability of hemocyte to bind to *S. aureus* (Gram-positive bacterium) and *E. coli* (Gram-negative bacterium) bioparticles using a cold-binding assay, which allows particle attachment while minimizing internalization. We used *NimC1*$^{1}$*, eater*$^{1}$ hemocytes as a negative control for a phagocytic deficient mutant.  Under these conditions, both *SR-CI*$^{SK6}$ and (*Δ(Sr-CI,Sr-CIII)*) hemocytes showed reduced binding to *S. aureus* and E. coli bioparticles compared with wild type hemocytes, as shown in Figure 2, confirming the main observation of (Rämet et al., 2001). The binding defect was however weaker than the ones observed with *NimC1, eater* deficient hemocytes (Figure 2).

*SR-CI is not required for phagocytosis of Gram-positive and -negative bacteria*

Although this was not initially claimed by (Rämet et al., 2001), SR-CI has often been considered as a phagocytic receptor (Melcarne et al., 2019; Ulvila et al., 2011). Experiments performed for ReproSci show no consistent reduction in phagocytosis of *S. aureus* or *E. coli* bioparticles by larval hemocytes in a *SR-CI*$^{SK6}$ or *Δ(Sr-CI,Sr-CIII)* mutants (Figure 3). In contrast, hemocytes from
*NimC1, eater* deficient larvae display strong phagocytosis defect as previously reported by (Melcarne et al ., 2019).

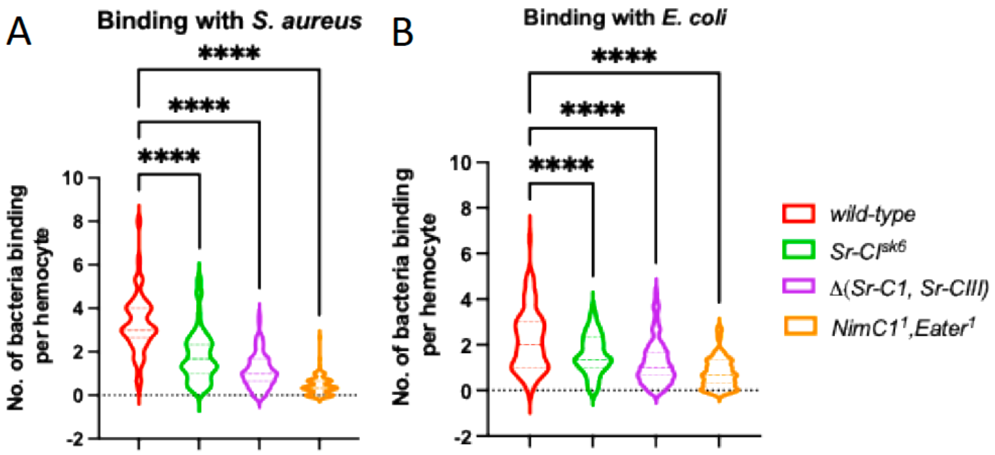

**Figure 2. SR-CI promotes the binding of *S. aureus and E. coli* bioparticles to hemocytes.** Quantification of cold-binding assayof hemocytes from hemocytes mutants and S. aureus (A) and E. coli (B) bioparticules. Violin plot from three independent experiments is represented. ****: $p<0.0001$ by one-Anova test.

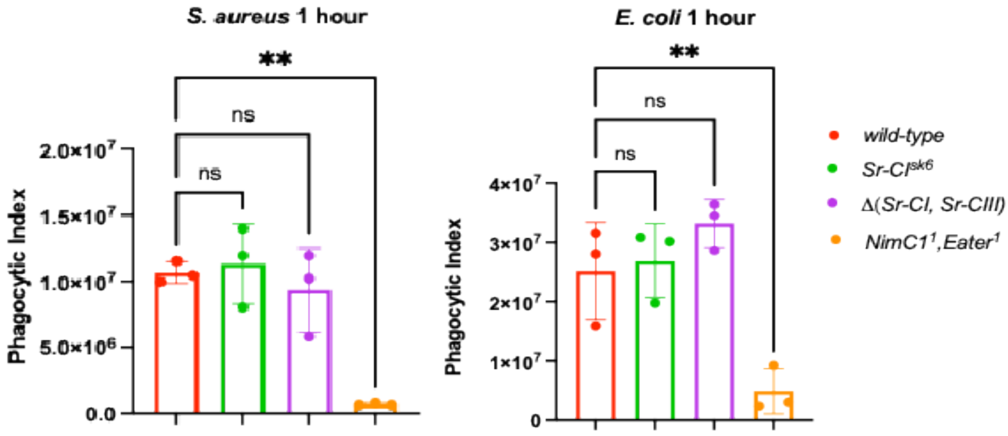

**Figure 3. Phagocytosis of *S. aureus* (A) and *E. coli* (B) bioparticles by hemocytes from *SR-CI*$^{SK6}$ , *Δ(Sr-CI,Sr-CIII), NimC1, eater* and wild-type larvae.** All the mutations were in the iso-drosdel background. No reduction in phagocytosis of *S. aureus or E. coli* bioparticles by *SR-CI* null or *Sr-CI,Sr-CIII* double mutant larval hemocytes was observed. A strong reduction in the phagocytosis of *E. coli and S. aureus bioparticules* was observed in *NimC1*$^{1}$*, eater*$^{1}$ as previously reported (Melcarne et al., 2019). Error bars represent the standard deviation from three different experiments. **: p<0.01 by one-Anova test. NS: not significant.

#### Conclusion

Consistent to (Rämet et al., 2001), we find that SR-CI contributes to hemocyte ability to bind to bacteria. SR-CI is however unlikely to be a major *Drosophila* phagocytic receptor for bacteria *in vivo.* Although SR-CI was defined as a pattern recognition, precise effector function in the *Drosophila* immune response remains to be identified .

### Supplementary S19: Role of Undertaker in phagocytosis of apoptotic cells and bacteria

**Abstract**

(Cuttell et al., 2008) concluded that Undertaker (also named UTA, Retinophilin, rtp) functions in phagocytosis of apoptotic cells and bacteria. Analysis of the literature and experiments performed for ReproSci instead suggest that these results were due to RNAi off-target effects, and that Undertaker is unlikely to have a role in phagocytosis.

**Introduction**

(Cuttell et al., 2008) found that Undertaker (UTA, Retinophilin, rtp), a Junctophilin, functions in phagocytosis of apoptotic cells and bacteria and links Draper-mediated phagocytosis to Ca2+ homeostasis. They found that Draper, its adaptor drCed-6, Undertaker, the Ryanodine receptor Rya-r44F, the ER Ca2+ sensor dSTIM, and the Ca2+ channel dOrai all act in the same pathway that promotes calcium homeostasis and phagocytosis. These results were based on RNAi against *undertaker* and a deficiency line (*Df(3R)3-4*) where reduced phagocytosis was rescued through transgenic re-expression of *undertaker.* Independent studies have shown that Retinophilin/Undertaker is a light-sensitive peripheral membrane protein in photoreceptor cells (Mecklenburg et al., 2010) and the primary phenotype of $rtp^1$ null mutants is age-dependent impairment in termination of the photoresponse (Venkatachalam et al., 2010).

Independent data support the notion that calcium flux and Store-Operated Calcium Entry (SOCE), including all the other components discussed in (Cuttell et al., 2008), are required for phagocytosis, and more generally for blood cell activation and cell shape change in a variety of animal models including *Drosophila.* A requirement for calcium flux in efferocytosis is conserved in other animals (Gronski et al., 2009; Moon et al., 2020); a requirement in phagocytosis of bacteria has not been further studied but is consistent with a requirement for calcium flux in cell shape changes during engulfment. (Mortimer et al., 2013) found that *Rya-r44F* and SOCE are required for encapsulation of parasitoid wasp eggs, suggesting a more general requirement for SOCE in activation of cellular immunity. They found that *Ganaspis sp.* wasp venom contained a SERCA protein that inhibited the calcium burst associated with hemocyte activation, and suppressed hemocyte migration towards the parasitoid egg. The results of (Etchegaray et al., 2012) suggest that some functions attributed to Draper may instead be due to a requirement of Draper in activating JNK, and that $Ca^{2+}$ entry and associated cell remodeling is JNK-dependent since overexpression of JNK components are sufficient to rescue phagocytic defects in *draper*-deficient cells. This is consistent with many results showing that JNK is generally required for cytoskeletal remodeling, activation of signaling pathways, and phagocytosis. (Weavers et al., 2016) similarly show that engulfment of apoptotic cells by *Drosophila* hemocytes triggers calcium flux that induces JNK signaling, and that this results in upregulation of phagocytic receptors including *Draper*; in the absence of this priming by apoptotic cells, hemocytes fail to migrate to wound sites and engulf apoptotic cells.

Although the (Cuttell et al., 2008) results regarding SOCE are well supported, those regarding the role of *undertaker* in this process are not. The same authors pursued the role of *undertaker* in a follow-up paper, but did not use specific mutants which were available at this time and instead used deficiencies (Goethem et al., 2012). Furthermore, the claim that *retinophilin* (*undertaker*) is involved in phagocytosis is inconsistent with several lines of evidence. First, it is not expressed or minimally expressed at the embryonic stage in which (Cuttell et al., 2008) performed the experiments: both protein and mRNA are only detectable in the adult head from the pupal stage onwards (FlyAtlas2, (Lee et al., 2012)). Furthermore, Retinophilin is entirely absent at the mRNA and protein level in fly mutants lacking eyes (Mecklenburg, 2007), which argues against a role in embryonic macrophages. Second, many of the experiments of (Cuttell et al., 2008) were performed in S2 cells, in which *retinophilin* is not expressed at detectable levels (FlyBase modENCODE). *retinophilin* expression is detected in only a very small subset of hemocytes by single cell RNAseq (Tattikota et al., 2020), arguing against a significant role in hemocyte function. Third, it is relevant to note that *undertaker/retinophilin* contains a repetitive motif (MORN repeat) that is found in several other proteins largely implicated in cellular trafficking in *Drosophila* (encoded by *CG3222, CG3306, junctophilin, CG30429, Rsph1, Als2*).

These data collectively suggest that results obtained using RNAi of *undertaker* may be due to off-target effects on a gene with similar motifs. A possible off-target effect of RNAi is consistent with the results of (Bhattacharya et al., 2012), who used KK RNAI to find a role for *retinophilin* in promoting axonal degeneration in larvae, another stage in which this gene is ostensibly not expressed. Off-target effects would be expected to impair cytoskeletal function, which is essential in phagocytosis. Furthermore, (Cuttell et al., 2008) concluded that Undertaker is present in hemocytes based on staining with a polyclonal antibody generated against *retinophilin* cDNA; this antibody was generated by (Mecklenburg, 2007) and used to stain *Drosophila* Retinophilin in the eye, where *retinophilin* is known to be expressed (FlyAtlas2, (Lee et al., 2012; Mecklenburg, 2007; Mecklenburg et al., 2010; Venkatachalam et al., 2010)). However, neither these authors nor (Cuttell et al., 2008) verified that this antibody specifically binds a single protein in *Drosophila,* and (Mecklenburg, 2007) did not look for antibody binding in tissues other than the eye. This lack of control combined with data showing that *retinophilin* is not expressed in the hemocytes suggests that this antibody bound a different protein bearing similar motifs (e.g. MORN repeats) that is present in the hemocytes, such as Junctophilin. Lastly, *Drosophila* Undertaker/Retinophilin is not expected to perform a junctophilin-like function, as it lacks the predicted ER/SR spanning transmembrane domain required for this function found in junctophilins (Mecklenburg, 2007).

Given these lines of evidence, *retinophilin* is unlikely to have a role in phagocytosis. To more conclusively determine whether *retinophilin* has a role in phagocytosis *in vivo,* we undertook experiments using the $rtp^1$ null allele.

#### Methods

The $rtp^1$ null allele (BDSC #35530) was generated by (Venkatachalam et al., 2010). $w^{1118}$ (BDSC #3605) flies were used as a wild-type control. *NimC1, eater*$^1$ and *draper*$^{D5}$ flies are described in (Melcarne et al., 2019) and (MacDonald et al., 2006), respectively. All fly stocks were raised on standard cornmeal–molasses–yeast agar at 25°C in a 12-hour light/dark cycle.

Phagocytic index was measured and calculated as previously described (Melcarne et al., 2019). Hemocytes from wandering third instar larvae were incubated with $2 \times 10^7$ *Escherichia coli* (K-12 strain) bioparticles Alexa Fluor 488 conjugate (Invitrogen™), or Alexa Fluor 488 conjugate from *Staphylococcus aureus* (Wood strain without protein A) bioparticles (Invitrogen™) for 60min. For the phagocytosis assay including the apoptotic cells, the S2 *Drosophila* cells were cultured in Schneider's insect medium (Sigma-Aldrich) containing 10% FBS (Gibco™), penicillin (Sigma-Aldrich) and streptomycin (Sigma-Aldrich) at a concentration of 100 U/ml. To induce apoptosis, cycloheximide (CHX, Sigma-Aldrich) was added at a final concentration of 50μg/ml. 20 hours after the cycloheximide treatment, the apoptotic cells were isolated and removed by pelleting with centrifugation at 400g for 5 min at 4°C. In order to stain the apoptotic bodies, CellTrace™ CFSE Cell Proliferation Kit (Invitrogen™) was added to the supernatant harvested from the cells at a final concentration of 5 μM and incubated 15 min at room temperature in the dark.

The statistical analysis was performed using ordinary one-way ANOVA followed by Dunnett's multiple comparisons test with a single pooled variance.

#### Results

Experiments performed for ReproSci using the $rtp^1$ null allele show no effect of this mutation on phagocytosis of bacteria or apoptotic cells by larval hemocytes in an *ex vivo* assay (Figure 1).

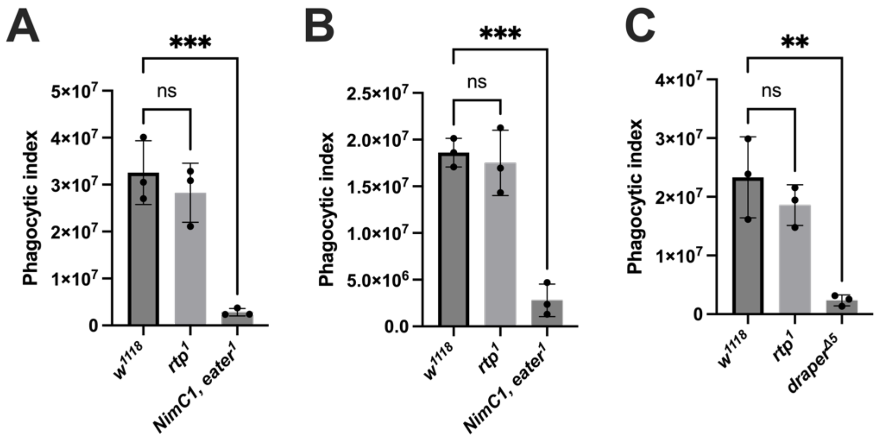

**Figure 1.** Phagocytosis of **A)** *E. coli* bioparticles, **B)** *S. aureus* bioparticles and **C)** apoptotic cells by *rtp*$^1$ wandering third instar larval hemocytes compared to wild-type (*w*$^{11118}$) and phagocytosis-deficient (*NimC1, eater*$^1$ and *draper*$^{\Delta 5}$) controls. Phagocytosis was measured by FACS after 1h incubation. Each data point represents the mean of two technical replicates of 5 larvae each from one independent experiment. n=3 experiments. Error bars represent the standard deviation.

#### Conclusion

Based on these lines of evidence we conclude that *undertaker* is unlikely to have a major role in phagocytosis.The results of (Cuttell et al., 2008) are most consistent with a scenario in which RNAi generated off-target knockdown of a protein related to *retinophilin/undertaker* (most likely *junctophilin,* which is expressed in hemocytes). We cannot however exclude that Undertaker play a role at a specific stage of development that we did not assess.

### Supplementary S20: Role of psidin in the regulation of Defensin

**Abstract**

(Brennan et al., 2007) found that Psidin acts nonautonomously in the hemocytes to control induction of *Defensin* in the fat body, and suggested that a cytokine produced by hemocytes is required to activate systemic *Defensin* expression. In contrast to this, experiments performed for ReproSci using two independent *psidin* mutants show that *Defensin* expression is similar to wild-type in these lines. Furthermore, evidence from multiple papers suggests that this result and other instances where various mutations have been found to specifically eliminate *Defensin* expression is likely due to segregating polymorphisms in *Defensin* that disrupt primer or probe binding in some genetic backgrounds and lead to a false negative result.

(Brennan et al., 2007) further found that Psidin is required for phagolysosomal maturation. Experiments performed for ReproSci confirm that phagolysosomal maturation is impaired in independent *psidin* mutants, and furthermore that *psidin* mutant hemocytes are abnormally large and multinucleate.

**Introduction**

*psidin* (*phagocyte signaling impaired, ird16, NAA25*) encodes a component of the widely conserved NatB complex, an N-terminal acetyltransferase complex that acetylates proteins following translation. N-terminal acetylation may affect protein stability, localization, interactions, and cytoskeletal function; mis-regulation of this process is frequently associated with tumor development (Rathore et al., 2016). Consistent with this, homozygous and trans-heterozygous *psidin* mutants are lethal at the late L3 larval stage and die with progressive formation of melanotic tumors (Brennan et al., 2007; Wu et al., 2001).

In *Drosophila,* Psidin binds actin, and is present in puncta in the cytoplasm and lamellipodia. Notably, lamellipodia of *psidin* mutant neurons and neurites are abnormally elongated and many other cell migration and morphological defects are present (Brennan et al., 2007; Kim et al., 2011; Stephan et al., 2012). *Drosophila* Psidin and the NatB complex have been found to promote EGFR/ERK activity by affecting stability and post-translational levels of positive and negative pathway regulators (Sheng and Du, 2020). EGFR signaling is thought to restrict immune activation in *Drosophila* through upregulation of the negative regulator *pirk* (Ragab et al., 2011). High throughput expression data show that *psidin* is expressed in all tissues and at all developmental stages, and is not appreciably regulated by infection (FlyAtlas2).

*Results showing specific elimination of* Defensin *expression are likely due to segregating polymorphisms*

Brennan et al. (2007) infected trans-heterozygous mutant larvae (psidin$^1$ EMS/Df(3R)H15) and examined antimicrobial peptide (AMP) gene expression at 3 hours post-infection with *Escherichia coli, Micrococcus luteus* and *Bacillus subtilis* using Northern blot. They found that *psidin*-deficient larvae showed no systemic induction of *Defensin*, while the expression of other AMPs was not strongly affected. *Defensin* expression in larvae was rescued through transgenic re-expression of *psidin* ubiquitously or in the hemocytes, but not the fat body. Furthermore, (Brennan et al., 2007) found that *domino* mutant larvae lacking hemocytes had impaired *Defensin* expression similar to *psidin* mutants. Epistasis suggested that a cytokine other than Spz or Upd3 or alternatively a product of bacterial degradation was responsible for this effect.

(Brennan et al., 2007) concluded that Psidin is required nonautonomously in the hemocytes to control induction of specifically *Defensin* in the fat body. This is inconsistent with results showing that hemocytes are generally dispensable in larvae for systemic (fat body) AMP expression in response to septic infection with *Ecc15* (Basset et al., 2000), although hemocytes are required for full expression of AMPs in response to *Ecc15* oral infection (Basset et al., 2000; Charroux and Royet, 2009). In contrast to these results, (Shia et al., 2009) found that AMP induction in hemocyte-ablated larvae following septic infection was impaired more strongly than would be expected if only hemocyte expression of AMPs was lost (the hemocytes account for only 15-25%

of total *Diptericin* and *Drosomycin* expression). They concluded that hemocytes are required for AMP induction in the fat body following both septic and oral infection, but more strongly following oral infection. However, (Charroux and Royet, 2009) and (Defaye et al., 2009) both found that *Defensin* expression was not reduced upon septic bacterial infections in flies with a genetic ablation of hemocytes, in contrast to the (Brennan et al., 2007) conclusion. Interestingly, while many AMPs are strongly expressed in the hemocytes, single cell RNAseq shows that *Defensin* is expressed at a near-zero level in hemocytes (Cho et al., 2020; Tattikota et al., 2020), and is not upregulated in hemocytes in response to bacterial infections (Ramond et al., 2020).

*Defensin* has repeatedly been singled out as an AMP whose expression is eliminated by a variety of mutations, while other AMPs are mildly affected or unaffected (e.g., (Brennan et al., 2007; Dijkers and O'Farrell, 2007; Jin et al., 2008)). Serendipitously, we found in the course of other studies that the *Defensin* qPCR primers listed by (Neyen et al., 2014) in "Methods to Study *Drosophila* immunity" do not successfully amplify *Defensin* in all wild type lines, leading us to suspect that a segregating polymorphism may interfere with quantification of this gene across genetic backgrounds. There are multiple polymorphisms segregating in *Defensin* in wild flies (Lazzaro and Clark 2003). Analysis of DGRP sequences showed two polymorphisms in the binding site of the (Neyen et al., 2014) forward *Defensin* primer that are expected to affect binding and amplification in qPCR. Note that these polymorphisms would not be expected to affect measurement of *Defensin* expression in high-throughput RNAseq studies due to the way in which gene expression is measured. In light of this, we analyzed the primer sequences used in studies that find 'ablation' of specifically *Defensin* expression.

Using qPCR, (Jin et al., 2008) found that in adult flies, several mutations impairing hemocyte development also caused a specific defect in *Defensin* expression more than other AMPs 6h following *Beauveria bassiana* injection. However, analysis of the DGRP showed that the (Jin et al., 2008) reverse primer binds over a polymorphism that occurs in ~20% of DGRP lines, causing a 3' mismatch to the alternate allele, which would greatly interfere with amplification by qPCR.

Northern Blot is a technique where a gel containing separated RNA is bathed with DNA probes corresponding to specific gene sequences, produced through PCR using forward and reverse primers and a genomic DNA template. This technique should be less sensitive to mismatches than qPCR, but polymorphisms and binding conditions used for Northern blot may still affect probe binding and sensitivity. The reverse primer used by (Brennan et al., 2007) to generate DNA probes for Northern blot binds over two polymorphisms annotated in the DGRP that appear to be in linkage (frequencies of ~20% for both alternate alleles). The multiple nucleotide mismatches present in the region covered by the DNA probe used in (Brennan et al., 2007) may interfere with the signal recovered by Northern blot by in unclear ways.

These findings suggest that studies showing a specific 'elimination' of *Defensin* expression may be due to poor primer binding, generating a false negative result in some genetic backgrounds. Presence of multiple alleles of *Defensin* in lines used for genetic crosses in studies are expected to result in a distribution of homozygotes with the reference allele that show 'strong *Defensin* expression' due to successful primer binding, non-binding homozygotes with alternate alleles that show 'no *Defensin* expression', and half-binding heterozygotes with one reference allele and one alternate allele that show '50% *Defensin* expression'. This may explain the pattern observed in the Northern blot results of (Brennan et al., 2007) where they show a rescue of *Defensin* expression through transgenic re-expression of *psidin*. In crossing the *psidin* mutant to the UAS and driver lines, *Defensin* expression was 50% restored or fully restored as alternate *Defensin* alleles in the *psidin* line were combined with lines bearing reference alleles.

*Psidin in phagolysosomal degradation*

(Brennan et al., 2007) found that hemocytes of *psidin* mutant larvae developed large multinucleate cells, failed to degrade engulfed bacteria, and did not clear infections as wild-type hemocytes did, resulting in persistent elevated bacterial counts following infection. Immunostaining of larval hemocytes and S2 cells showed punctate labelling of Psidin in the cytoplasm that colocalized with a lysosome marker. (Kim et al., 2011) similarly show punctate cytoplasmic staining of Psidin.

To clarify the role of *psidin* in *Defensin* expression and phagolysosomal degradation, we tested induction of *Defensin* and bacterial engulfment in independent *psidin* null mutants.

**Methods**

*Fly lines and infection*

The $psidin^{85D1}$ ($psidin^{85D1}$ $e^{s}/TM3, Sb^{1}$) and $psidin^{55D4}$ ($w^{1118}$; $psidin^{55D4}$ $e^{s}/TM3, Sb^{1}$) alleles were obtained from Bloomington (stocks #41122 and #41123). These are EMS mutations that generate stop codons (Kim et al., 2011). $w^{1118}$ flies were used as wild type (Bloomington #3605); $NimC1^{1}$, $eater^{1}$ animals were used as a phagocytosis-deficient control (Melcarne et al., 2019).

All stocks were reared on cornmeal fly medium comprising 6% cornmeal, 6% yeast, 0.62% agar, 0.1% fruit juice, supplemented with 10.6g/L Moldex and 4.9ml/L propionic acid. Fly stocks were maintained at 25°C on a 12h light/12h dark cycle and flipped every 2-3 days to fresh medium. For experiments, *psidin* homozygous L3 wandering larvae were selected from $psidin^{85D1}/TM6B_{tb}$ and $psidin^{55D4}/TM6B_{tb}$ lines, respectively. The mutations were verified through Sanger sequencing of ~1100bp of the *psidin* gene region. The primers used for the sequencing are the following:

| Name | Sequence |
|---|---|
| Psidin_2834_F | CGGAAATGGGACCATCGGATGC |
| Psidin_3957_R | TAAACTGGCTGCCACTTTCTGCTG |

*Infection experiments*

The posterior part of the larva near the spiracles of L3 larvae was pierced using a 0.1 mm tungsten needle dipped in $OD_{600}$=200 *E. coli*-GFP from an overnight culture in LB medium (37°C, shaking).

*Phagocytosis assays and microscopy*

For microscopy experiments on hemocytes obtained from wandering third instar larvae after *E. coli*-GFP infection, larvae were bled into 150 µl Schneider medium, and hemocytes were allowed to adhere on a glass slide for 40 min and fixed for 10 min in PBS containing 4% paraformaldehyde. After fixation the cells were subsequently rinsed in 1x PBS pH 7.4 + 0.1% Triton X-100 (PBS-T), permeabilized, and blocked in PBS-T + 2% bovine serum albumin (BSA) + 10% goat serum for 1 h and incubated overnight with anti-GFP (chicken, Abcam, 1:1000) primary antibody solution in the same blocking solution at 4°C. After 3 5-minute washes, secondary antibody (anti-chicken labelled with AlexaFluor488, 1:2000) was applied at room temperature for 2 h. Finally, cells were stained with 1/15000 dilution of DAPI (Sigma-Aldrich) and mounted in Dako fluorescence media. Cells were imaged under a Leica UP2 Confocal microscope with a 69X oil objective and analyzed using the Fiji software.

For cell area measurements, hemocytes were collected from L3 larvae, fixed for 10 minutes in 4% PFA, and stained with 1:1000 phalloidin in PBS-T for 1 hour at room temperature. Mounting medium containing DAPI was applied and hemocytes were imaged using appropriate fluorescence filters. All settings were designed by visual inspection. Approximately 50 cells per genotype were analyzed.

*Gene expression levels by RT-qPCR*

Gene expression measurements were performed by RT-qPCR as previously described (Carboni et al., 2022). 15 whole larvae were homogenized and their RNA was extracted using TRIzol reagent (Invitrogen) and resuspended in RNase-free water. Reverse transcription was performed using PrimeScript RT kit (Takara) with random hexamers and oligo dTs. Quantitative PCRs were performed on a LightCycler 480 (Roche) using PowerUp SYBR Green Master Mix (Applied Biosystems). Expression levels were normalized to *rp49* (*rpl32*).

| Gene | Source | Name | Primer sequence |
|---|---|---|---|

| | | | |
|---|---|---|---|
| *DptA* | Hanson et al. (2016) | DptA-HanF | ATG CCC GAC GAC ATG ACC AT |
| | | DptA-HanR | TTG TCG GTG GTC CAC ACC TT |
| *Drs* | Bruno Lemaitre | Drom-F | CGTGAGAACCTTTTCCAATATGAT |
| | | Drom-R | TCCCAGGACCACCAGCAT |
| *Def* | Mark Hanson | Def_77F | TTCTTCGTTCTGTGGCTATCG |
| | | Def_185R | CATCCTCATGCACCAGGACAT |
| *Rp49/RpL32* | Bruno Lemaitre | RpL32 F | GCC GCT TCA AGG GAC AGT ATC TG |
| | | RpL32 R | AAA CGC GGT TCT GCA TGA G |

*Statistical analysis*

Quantitative PCR statistical data analysis was performed using GraphPad Prism software. A two-way ANOVA was carried out followed by a post-hoc Dunnett's multiple comparisons test with a single pooled variance to compare mutants to the controls.

**Results and Discussion**

*Loss of psidin does not impair* Defensin *expression*

Contrary to (Brennan et al., 2007), experiments performed for ReproSci using two independent null EMS mutations of *psidin* show that *Defensin* expression in larvae in response to septic infection is not impaired as a result of *psidin* deficiency at 6h after infection with *E. coli*. Expression of *Diptericin* and *Drosomycin* was also similar to that of wild-type flies (Figure 1). Furthermore, AMP expression was normal in *NimC1, eater[1]* larvae whose phagocytes fail to phagocytose bacteria, indicating that degradation products of phagocytosis are not necessary for systemic AMP induction in larvae.

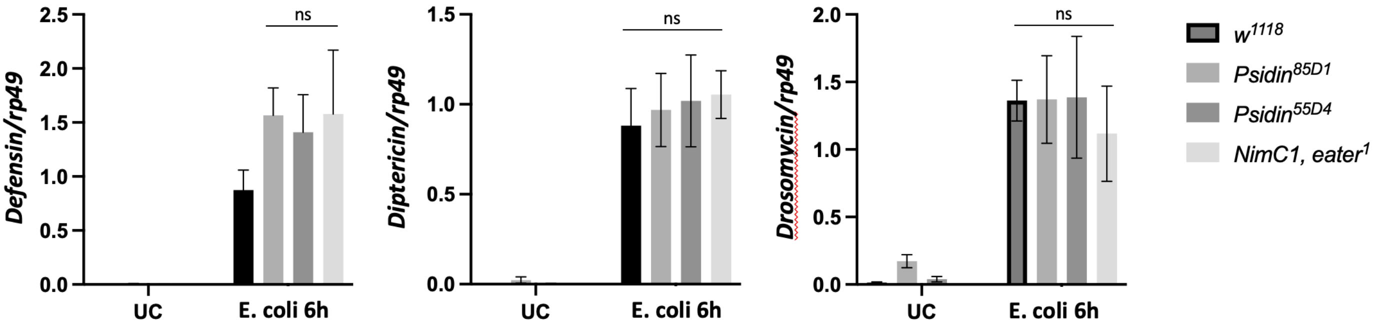

**Figure 1.** AMP expression in wild-type, *psidin* mutant and *NimC1, eater[1]* mutant larvae in unchallenged conditions (UC) and 6h following septic infection with *E. coli (1106).* Error bars represent standard error of the mean. ns, not significant.

*Loss of psidin impairs hemocyte phagolysosomal degradation*

Experiments performed for ReproSci show that hemocytes of larvae from independent *psidin*-deficient lines were significantly larger than those of wild-type larvae (Figure 2A, B). Furthermore, a greater proportion of *psidin*-deficient hemocytes had internalized bacteria compared to wild-type hemocytes (Figure 2B, C). This indicates that *psidin*-deficient larval hemocytes accumulate bacteria, consistent with impaired phagolysosomal degradation. This is consistent with studies showing that Psidin and the *Drosophila* NatB complex interact with and aid in control of the actin cytoskeleton (Kim et al., 2011; Stephan et al., 2012), which is essential for fusion events leading to lysosomal maturation.

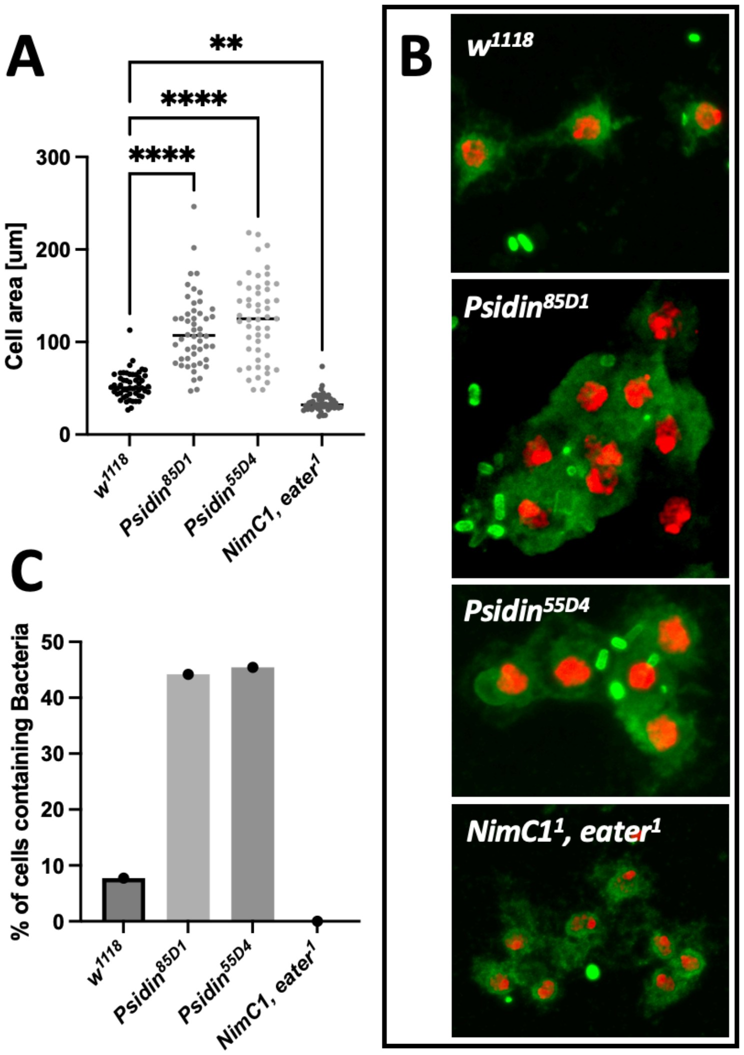

**Figure 2. A,** Hemocytes of *psidin* mutant larvae were significantly larger than those of wild-type larvae. **B,** Representative images showing undegraded GFP-bacteria (bright green spots) within multinucleate *psidin* mutant hemocytes. Phalloidin is in green and DAPI is in red. **C,** A larger proportion of *psidin* mutant hemocytes were observed with internalized bacteria following 2h incubation with *E. coli*. ****, p<0.0001; ***, p<0.001; ** p<0.01; *p<0.05; ns, not significant.

#### Conclusion

Contrary to (Brennan et al., 2007), we find that *psidin* is not required for systemic *Defensin* expression following septic infection of larvae. It is likely that this conclusion was due to segregating polymorphisms in *Defensin* that interfered with primer binding in a subset of genetic backgrounds. However, similar to (Brennan et al., 2007), experiments performed for ReproSci indicate that *psidin* is required for phagolysosomal maturation and degradation of bacteria internalized by hemocytes. Furthermore, our data confirm that *psidin* mutants develop large, multinucleate hemocytes.

### Supplementary S21: Role of Hemese in lamellocyte differentiation and encapsulation

**Abstract**

(Kurucz et al., 2003) found that loss of *Hemese*, a hemocyte-specific transmembrane protein, causes overproduction of lamellocytes in permissive conditions. In contrast to this, using *Hemese*$^{JP187}$ and *Hemese*$^{JP828}$ mutants, we find that loss of *Hemese* does not strongly affect lamellocyte number or encapsulation.

**Introduction**

(Kurucz et al., 2003) found that Hemese, a hemocyte-specific transmembrane protein, modulates activation or recruitment of hemocytes in response to wasp parasitization, and may be involved in transformation of blood cells into lamellocytes. (Kurucz et al., 2003) found that *Hemese* knockdown (dsRNA) did not have noticeable effects in unchallenged larvae, but upon wasp parasitization by *Leptopilina boulardi*, the number of lamellocytes produced was greatly increased compared to wild type.

Single cell RNAseq of hemocytes and tissues (Li et al., 2022; Ramond et al., 2020; Tattikota et al., 2020) suggests that *Hemese* is overwhelmingly expressed in hemocytes and not in other cell types. High throughput expression data show that *Hemese* expression is negligible in all adult tissues, and in larvae is primarily found in tissues likely to be contaminated by adherent hemocyte clusters (FlyAtlas2). The function of Hemese has not been further investigated.

To clarify the role of Hemese, we tested lamellocyte differentiation and wasp egg encapsulation in *Hemese*-deficient larvae.

**Methods**

*Fly stocks*

We generated two small deletions of *Hemese*, *Hemese*$^{JP187}$ and *HemeseJP*$^{828}$, by replacing a 370-bp coding sequence of the *Hemese* coding sequence with a 1.7-kb *dsRed* cassette using CRISPR/Cas9 (Figure 1). *w*$^{1118}$ (BDSC #3605) flies were used as a wild-type comparison.

*Hemocyte Counts by Flow Cytometry*

Hemocytes from *Hml-GAL4 > UAS-GFP* larvae were initially used to define the hemocyte population for flow cytometry analysis. GFP-positive hemocytes were identified and gated using the B525 fluorescence channel, allowing exclusion of debris and non-hemocytic cells. This gating strategy was subsequently applied to all genotypes to quantify total hemocyte numbers. For each genotype, five wandering third-instar (L3) larvae were bled into 150 μL of Schneider's insect medium. Prior to bleeding, larvae were vortexed in 1× PBS for 1 min to dislodge sessile hemocytes. Approximately 60 μL of the resulting hemocyte suspension was analyzed by flow cytometry. For each biological replicate, flow cytometry measurements were performed in three technical replicates.

*Hemocyte Staining and Quantification*

Before and 96 hours after wasp infection, five third-instar larvae were washed and dissected in 150 μL of Schneider's medium while keeping vital organs intact. Samples were incubated in a humidified chamber for 45 min to allow hemocytes to settle and adhere to the slide surface. Cells were then fixed with 4% paraformaldehyde (PFA) and stained with phalloidin (1:200) for 1 h at room temperature, followed by three washes with 1× PBS. Subsequently, DAPI (1:20,000) was added for 10 min at room temperature, and samples were washed three additional times with 1× PBS. Slides were mounted using mounting medium and sealed with nail polish. Hemocytes were imaged using an inverted Evident confocal microscope. Maximum-intensity Z projections were generated for analysis, and image quantification was performed using FIJI/ImageJ software. For each biological replicate, 10 fields of view were analyzed. Lamellocyte abundance was quantified as the ratio of lamellocytes to plasmatocytes per field of view, and the mean value from the 10 analyzed fields was used for statistical analysis.

*Wasp infestation and quantification of encapsulation*: For wasp infestation experiments, 30 synchronized second instar (L2) larvae were placed on a pea-sized mound of fly food within a custom-built wasp trap in the presence of three female *Leptopilina boulardi* G486 parasitic wasps (kindly provided by Michèle Crozatier, Toulouse). Infested third instar larvae were bled 96 hours later and the presence of lamellocytes was checked using a phalloidin staining. Pictures of melanised eggs were taken with a Leica DFC300FX camera and Leica Application Suite.

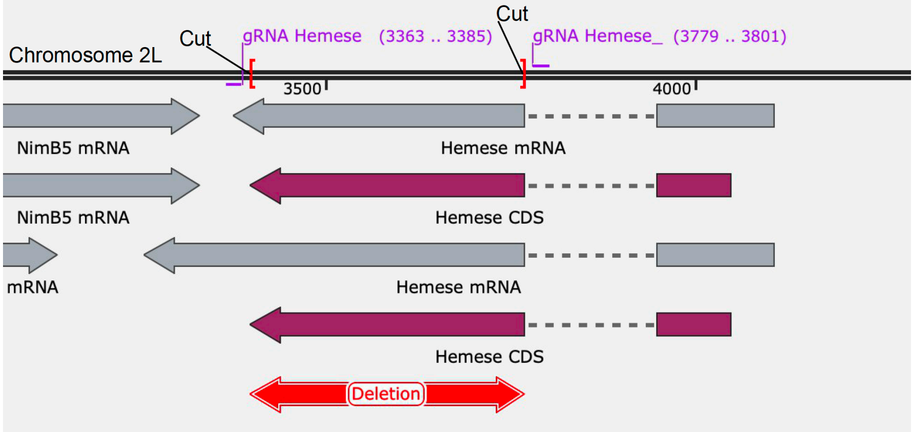

**Figure 1 Generation of two small deletions of *Hemese*.** The *Hemese*$^{JP187}$ and *HemeseJP*$^{828}$ mutants were produced by replacing a 370-bp coding sequence of the *Hemese* coding sequence with a 1.7-kb *dsRed* cassette using CRISPR/Cas9. Guide RNAs used to generated the deletion are indicated.

**Results**

We generated two fly lines deleted for the *Hemese* gene, *Hemese*$^{JP187}$ and *HemeseJP*$^{828}$ in the iso *w*$^{1118}$ background by replacing a 370-bp coding sequence of the *Hemese* gene with a 1.7-kb *dsRed* cassette using CRISPR/Cas9. These mutations did not affect fly viability. We however observed a higher hemocyte counts in *Hemese*$^{JP187}$ and *Hemese*$^{JP828}$ third instar larvae compared to the wild-type (Figure 2A). Encapsulation of wasp eggs, measured by the presence of black spots was similar in *Hemese*$^{JP187}$ and *Hemese*$^{JP828}$ and wild type larvae (Figure 3A). Furthermore, *Hemese*$^{JP187}$ and *Hemese*$^{JP828}$ larvae did not shown enhanced lamellocyte differentiation upon wasp infection compared to wild-type larvae (Figure 3B and 2C).

**Conclusion**

Our results do not support the conclusion of Kurucz et al. (2003) that Hemese negatively modulates lamellocyte numbers in response to L. boulardi infestation. While our data suggest that Hemese may influence overall hemocyte abundance, they do not rule out a more subtle role in lamellocyte differentiation or in the encapsulation process.

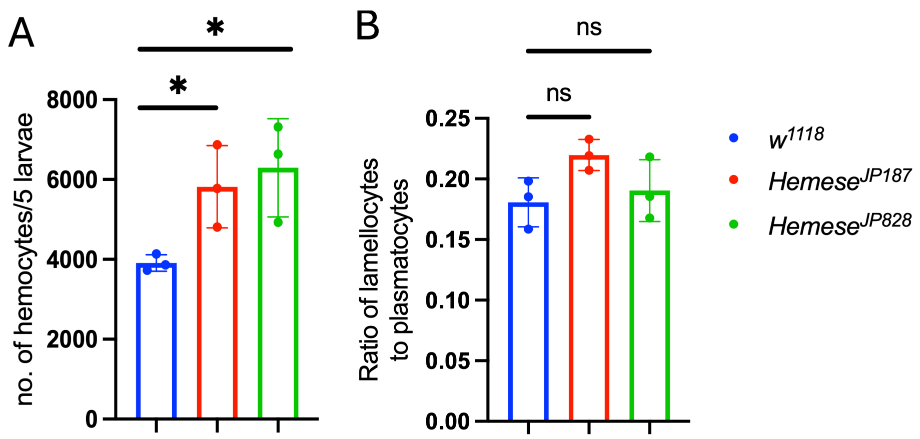

**Figure 2: Hemocyte number and lamellocyte differentiation in wild-type and *Hemese* mutant larvae.**
**A.** Number of hemocytes in $w^{1118}$ (wild-type) and $Hemese^{JP187}$ and $Hemese^{JP828}$ third instar larvae (each point is biological replicate of 3 samples of 5 larvae). **B.** Ratio of lamellocytes to plasmatocytes in third instar larvae of w1118, HemeseJP187, and HemeseJP828 following L. boulardi infection. Lamellocyte abundance was quantified as the ratio of lamellocytes to plasmatocytes per field of view (Figure 3B). Each data point represents a biological replicate, calculated as the mean of ten fields from hemocyte preparations obtained from five larvae. Statistical significance was assessed using a one-way ANOVA.

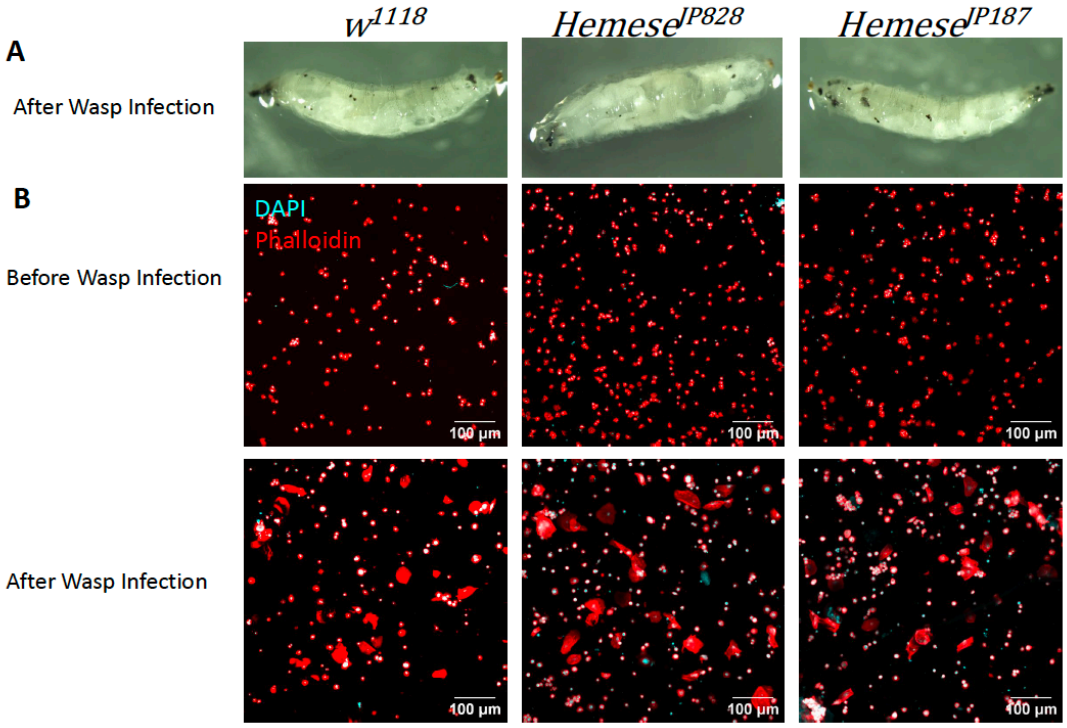

**Figure 3: Encapsulation of *Leptopilina boulardi* wasp eggs in third instar larvae of wild-type (*$w^{1118}$*), *$Hemese^{JP187}$*, and *$Hemese^{JP828}$*.**
(**A**) Following L. boulardi infestation, wild-type and *Hemese* mutant larvae exhibit a similar number of melanized black spots.
(**B**) Phalloidin staining of hemocytes from wasp-infested $w^{1118}$, $Hemese^{JP187}$, and $Hemese^{JP828}$ larvae. Plasmatocytes and lamellocytes are shown; lamellocytes weree identified as large, flattened cells. Quantification is provided in Figure 2B.

Li H, Janssens J, De Waegeneer M, Kolluru SS, Davie K, Gardeux V, Saelens W, David FPA, Brbić M, Spanier K, Leskovec J, McLaughlin CN, Xie Q, Jones RC, Brueckner K, Shim J, Tattikota SG, Schnorrer F, Rust K, Nystul TG, Carvalho-Santos Z, Ribeiro C, Pal S, Mahadevaraju S, Przytycka TM, Allen AM, Goodwin SF, Berry CW, Fuller MT, White-Cooper H, Matunis EL, DiNardo S, Galenza A, O'Brien LE, Dow JAT, FCA Consortium§, Jasper H, Oliver B, Perrimon N, Deplancke B, Quake SR, Luo L, Aerts S, Agarwal D, Ahmed-Braimah Y, Arbeitman M, Ariss MM, Augsburger J, Ayush K, Baker CC, Banisch T, Birker K, Bodmer R, Bolival B, Brantley SE, Brill JA, Brown NC, Buehner NA, Cai XT, Cardoso-Figueiredo R, Casares F, Chang A, Clandinin TR, Crasta S, Desplan C, Detweiler AM, Dhakan DB, Donà E, Engert S, Floc'hlay S, George N, González-Segarra AJ, Groves AK, Gumbin S, Guo Y, Harris DE, Heifetz Y, Holtz SL, Horns F, Hudry B, Hung R-J, Jan YN, Jaszczak JS, Jefferis GSXE, Karkanias J, Karr TL, Katheder NS, Kezos J, Kim AA, Kim SK, Kockel L, Konstantinides N, Kornberg TB, Krause HM, Labott AT, Laturney M, Lehmann R, Leinwand S, Li J, Li JSS, Li Kai, Li Ke, Li L, Li T, Litovchenko M, Liu H-H, Liu Y, Lu T-C, Manning J, Mase A, Matera-Vatnick M, Matias NR, McDonough-Goldstein CE, McGeever A, McLachlan AD, Moreno-Roman P, Neff N, Neville M, Ngo S, Nielsen T, O'Brien CE, Osumi-Sutherland D, Özel MN, Papatheodorou I, Petkovic M, Pilgrim C, Pisco AO, Reisenman C, Sanders EN, Dos Santos G, Scott K, Sherlekar A, Shiu P, Sims D, Sit RV, Slaidina M, Smith HE, Sterne G, Su Y-H, Sutton D, Tamayo M, Tan M, Tastekin I, Treiber C, Vacek D, Vogler G, Waddell S, Wang W, Wilson RI, Wolfner MF, Wong Y-CE, Xie A, Xu J, Yamamoto S, Yan J, Yao Z, Yoda K, Zhu R, Zinzen RP. 2022. Fly Cell Atlas: A single-nucleus transcriptomic atlas of the adult fruit fly. *Science* **375**:eabk2432. doi:10.1126/science.abk2432

Ramond E, Dudzic JP, Lemaitre B. 2020. Comparative RNA-Seq analyses of Drosophila plasmatocytes reveal gene specific signatures in response to clean injury and septic injury. *PLOS ONE* **15**:e0235294. doi:10.1371/journal.pone.0235294

Tattikota SG, Cho B, Liu Y, Hu Y, Barrera V, Steinbaugh MJ, Yoon S-H, Comjean A, Li F, Dervis F, Hung R-J, Nam J-W, Ho Sui S, Shim J, Perrimon N. 2020. A single-cell survey of Drosophila blood. *eLife* **9**:e54818. doi:10.7554/eLife.54818

### Supplementary S22: Intestinal IRC in not required for survival to benign oral infection

**Abstract**

An early report found that RNAi-mediated knockdown of *immune regulated catalase* (*Irc*) produced broad susceptibility to feeding with a wide range of pathogenic and non-pathogenic microbes (Ha et al., 2005). The authors concluded that *Irc*-mediated regulation of ROS (reactive oxygen species) levels in the gut was required to prevent lethal damage by ROS in response to enteric infections. To test these findings, we characterized an $Irc^{MB11278}$ null mutant line. Although we found some susceptibility of $Irc^{MB11278}$ null flies to oral infection with *Ecc15,* our observations indicate that this is due to locomotor defects found in these flies. Further studies are needed to clarify the role of *Irc* in immunity.

**Introduction**

(Ha et al., 2005) found that depletion of the extracellular *immune regulated catalase* (*Irc*) by ubiquitous or gut-specific RNAi caused high mortality in adult flies upon natural (oral) infection with a variety of weak pathogens (*Escherichia coli, Ecc15, Saccharomyces cerevisiae, Salmonella typhimurium, Micrococcus luteus*), although NF-κB mutants were entirely resistant to these benign infections. *Irc*-knocked-down flies had high basal and induced levels of ROS (reactive oxygen species) in the hemolymph compared to wild-type flies. They concluded that IRC is required to reduce ROS levels and prevent self-inflicted damage to the host in response to natural infections. Susceptibility of IRC-depleted flies was partially rescued by feeding flies with the antioxidant NAC, or by increasing JNK activity to promote expression of alternate antioxidant genes. The authors inferred that susceptibility of IRC-depleted flies was due to insufficient elimination of high ROS levels and subsequent lethal damage to the host. (Ha et al., 2005) conclude that *Irc*-mediated redox balance in the gut epithelium is the most important factor influencing host survival to ingestion of pathogens.

Although *Irc* was initially thought to be an extracellular catalase, homolog search show that IRC should not be considered as a bona fide catalase, but rather as a heme peroxidase, enzymes that uses hydrogen peroxide as a substrate to mediate some chemicals reactions such as protein cross-linking by forming dityrosine bonds (Kumar et al., 2010). It is upregulated upon septic infection in both the midgut and fat body (Chakrabarti and Visweswariah, 2020; De Gregorio et al., 2001; Ha et al., 2005) and less so upon natural infection (De Gregorio et al., 2001). Proteomic data confirm that IRC protein increases in the hemolymph in response to septic infection with either *Ecc15* or *M. luteus* (Rommelaere et al., 2025). Overexpression of *Irc* has been used in several studies as an effective method of reducing extracellular ROS in various contexts (Buchon et al., 2009; Chakrabarti and Visweswariah, 2020; Nam et al., 2012; Wang et al., 2013), confirming that IRC acts as a secreted catalase. However, few follow-up studies have been published on the role of IRC in survival to ingested pathogens. (Wang et al., 2013) found that overexpression of *Irc* had no effect on fly survival to ingestion of highly pathogenic *Vibrio cholerae.* (Prakash et al., 2022) found that flies bearing the $Irc^{MB11278}$ Minos insertion allele backcrossed to $w^{1118}$ had slightly increased susceptibility to oral infection with *Pseudomonas entomophila* ($OD_{600}$=25), but maintained lower intestinal bacterial loads than wild-type flies. The authors surmised that lower bacterial loads are maintained in *Irc* flies due to increased clearance by higher ROS.

To test whether *Irc* is required for wild-type resistance in response to enteric infections, we characterized the null $Irc^{MB11278}$ mutation.

**Methods**

*Fly lines*

The $Irc^{MB11278}$ allele (BDSC #29191) carries a Minos insertion element (7.5kb) with a Gal4 construct and EGFP marker that expresses in the *Drosophila* eye. It is inserted 1385bp into the gene region and 132bp downstream of the translational start site, upstream of the conserved Haem-peroxidase domain that is presumably responsible for the catalytic activity of IRC. This insertion also causes a frameshift (Figure 1). qPCR using primers targeted to the region downstream of the Minos insertion and upstream of the Haem-peroxidase domain showed ~6% of wild type expression in the *Irc*$^{MB11278}$ line. *w*$^{1118}$ (BDSC #3605), *w*$^{iso}$ *Drosdel* (Ryder et al., 2004) and *OR-R* flies were used as wild-types; in survival experiments *spz*$^{rm7}$ (Neyen et al., 2012), *Bom*$^{\Delta55C}$ *Drosdel* (Clemmons et al., 2015 ; Hanson et al., 2019) flies were used as *Toll*-pathway deficient comparisons, while *Relish*$^{E20}$ *Drosdel* (Ferreira et al., 2014) mutants were used as Imd-pathway deficient comparisons.

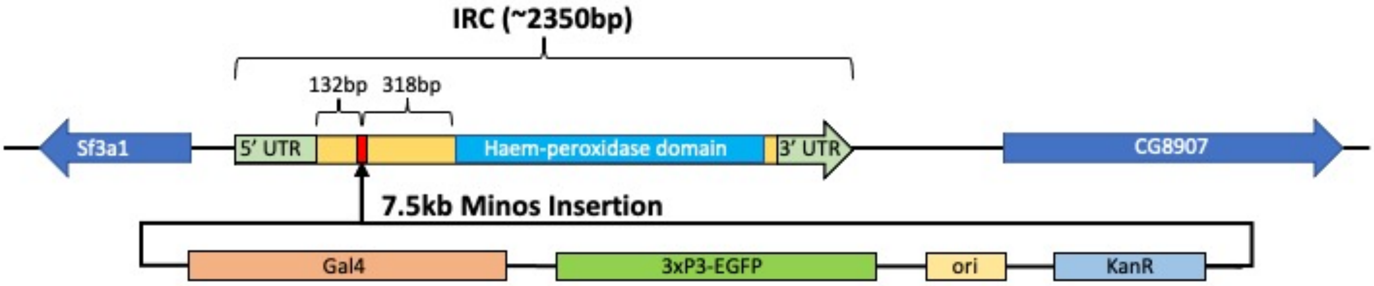

**Figure 1.** Diagrammatic representation of the *Irc*$^{MB11278}$ Minos insertion allele. The 7.5kb Minos element including a Gal4 construct and EGFP marker that expresses in the *Drosophila* eye is inserted 1385bp into the gene region and 132bp downstream of the translational start site, upstream of the conserved Haem-peroxidase domain that is presumably responsible for the catalytic activity of IRC. This insertion also generates a frameshift. Structure of Minos element from (Metaxakis et al., 2005).

*Survivals*

For natural (oral) infections, 5 to 7-day old male and female flies reared on cornmeal fly medium (6% cornmeal, 6% yeast, 0.6% agar, 0.1% fruit juice, 10.6 g/L moldex and 4.9 mL/L propionic acid) at 25°C were starved for 2h in empty vials at 29°C and flipped into vials in which a filter paper completely covering the cornmeal medium was soaked with 150 μL of 5% sucrose solution in either PBS (for controls) or a concentrated solution of the desired pathogen in PBS. *Ecc15* (*Pectobacterium carotovorum carotovorum*) was grown at 29°C in LB, shaking; *Candida albicans SC5314* was grown at 29°C in YPG, shaking. A final concentration of $OD_{600}$=400 was used for *C. albicans* and *Ecc15* oral infections, and the flies were flipped to new vials with fresh filter papers and microbial cultures every 2 days. To test whether survival differed in flies supplemented with *Saccharomyces cerevisiae,* bakers' yeast was sprinkled directly on the surface of the medium. Natural infection survivals were incubated at 29°C.

As *Irc* flies quickly became stuck to the medium and died as a result of being unable to free themselves, all survivals were performed with vial racks on their sides to reduce the number of flies falling into the medium.

#### Results

Initial characterization of Irc$^{MB11278}$ line was hampered by very low homozygous viability of the stock, particularly in females (5% -/- males, <1% -/- females). Crosses to a deficiency uncovering Irc (Df(3R)BSC792) increased the male homozygote rate to ~60% of the expected level, indicating that background/secondary mutations were partially responsible for lethality. However, no female flies were recovered from these crosses. Later crosses showed that some homozygous female flies eclosed, but immediately fell into the medium and died. Survival rate was improved by lying vials on their sides to allow eclosion, but females had very poor motor control and their wings were typically stuck to the dorsal surface along their length, preventing flight (Figure 2A). Flies were backcrossed eight times to the wiso line as described in (Ferreira et al., 2014) in an effort to improve viability. The isogenized *Irc*$^{MB11278}$ line had much better viability, although wing defects that were present in the line initially remained in both male and female flies (Figure 2). The line was therefore maintained over a balancer as eclosing homozygous females tended to fall into the medium and die. In female *Irc*$^{MB11278}$ iso flies, the wings were sometimes stuck to the dorsal abdomen (Figure 2A), preventing flight and impairing locomotion, but more often failed to completely extend, resulting in twisted wingtips (Figure 2D) or crumpled blades (Figure 2E). Male flies and less severely affected female flies had slightly curved or twisted wing blades (Figure 2C, F). However, in maintaining the line with no further backcrossing over a period of several months, the wing defects disappeared and the balancer was eventually lost, resulting in a fully homozygous viable line. The line was checked for contamination by verifying presence of the GFP marker expressed in the eyes encoded by the Minos insertion and by qPCR, and was found to be uncontaminated and have very low (6%) Irc expression.

A phenotype of reduced motor control in *Irc*$^{MB11278}$ flies, again particularly in females, was consistent across replicates in initial experiments and strongly enhanced by septic injury (Figure 3). However, this phenotype was also largely lost after maintaining the line for six months. These data may indicate some level of adaptation to loss of Irc in flies over time, perhaps through compensatory upregulation of other genes.

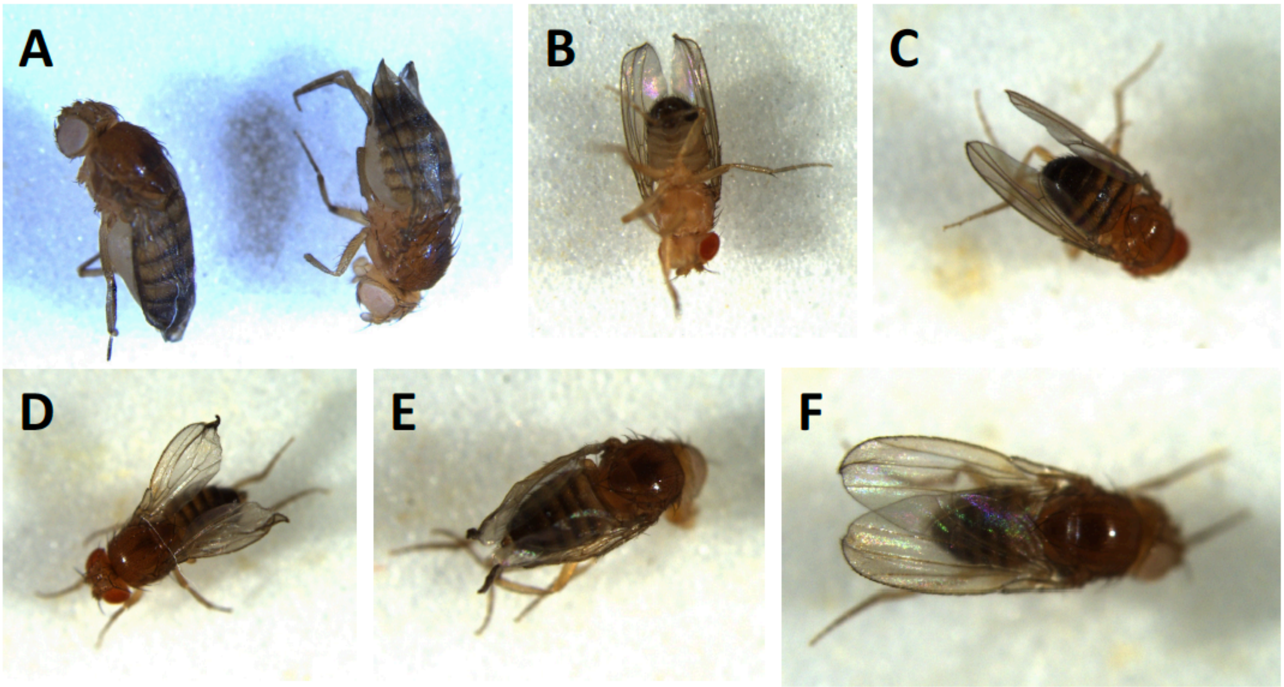

**Figure 2**. A: Wings of partially isogenized females were often completely glued to their backs despite not falling into food – these females were still alive after vial was left on its side to allow emergence. Fully isogenized *Irc*$^{MB11278}$ male with slightly bent wingtips (B) and male with nearly normal wings with slight crumpling of inside edges (C). At the end of the isogenization process, *Irc*$^{MB11278}$ females were much more viable but nearly all had wing defects. Most defects were similar to (D) with twisted wingtips but many had more severe detects, with wings that failed to inflate along their entire length (E). Some females had slight defects where only the tips of the wings bent and cupped as in some males (F). These defects were gradually lost with no further backcrossing as the line was maintained through several generations.

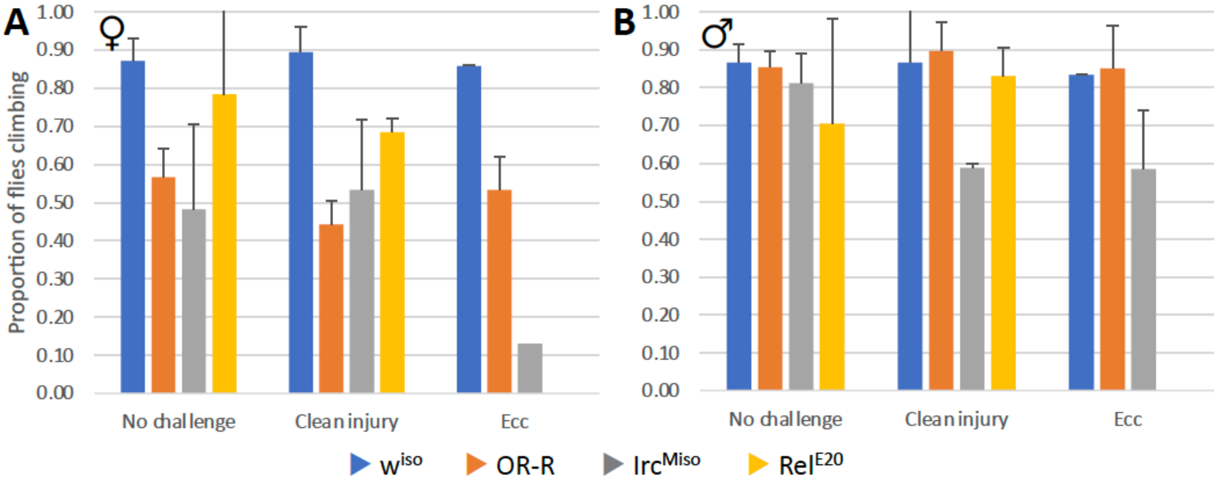

**Figure 3**. Two climbing pass rate assays performed 24h after injury or septic infection confirm qualitative observations that *Irc*$^{MB11278}$ iso mutant flies have locomotory impairment compared to wild-type flies, and that this is more severe in female flies than male flies, which require injury or infection to reveal a locomotory defect. This phenotype was gradually lost as the line was maintained with no further recombination. *Irc*$^{Miso}$: *Irc*$^{MB11278}$.

(Ha et al., 2005) found that IRC-RNAi flies (*Da>IRC-IR* or *Cad>IRC-IR*, where gut-specific RNAi had a weaker effect) had strongly reduced survival when fed with *E. coli, Ecc15, S. cerevisiae*, *S. typhimurium*, or *M. luteus*, with at least 60% of *Irc*-depleted flies dying by the fifth day of feeding, whereas wild-type flies and *relish* mutants were largely unaffected by this treatment. These results were seen with two insertions of an *Irc-RNAi* construct generated in-house. In their protocol, adults were starved 2h and transferred to a vial with filter paper hydrated with 5% sucrose solution and $10^{10}$ cfu/mL overnight bacterial culture, followed by incubation at 25°C. The filter paper was changed every day, and flies fed only sucrose were used as a control.

Similar to (Ha et al., 2005), our results showed that *Rel*$^{E20}$ flies were resistant as wild-types to short and long-term feeding with *Ecc15* (Figure 4A). We found that *Irc*$^{MB11278}$ (iso) flies were more susceptible to oral infection with *Ecc15* (Figure 4A). However, even with vials on their sides, *Irc*$^{MB11278}$ mutants in both *Ecc15* and sucrose treatments died after first becoming stuck to the medium by their wings, always surviving at least 24h prior to death. Similarly, *Irc*$^{MB11278}$ (iso) flies did not have increased susceptibility beyond that expected of their poor motor skills in response to feeding with *C. albicans* (Figure 4B), supplementation of standard medium with *S. cerevisiae* (data not shown), or feeding flies on medium containing 4% $H_2O_2$ (Figure 4C).

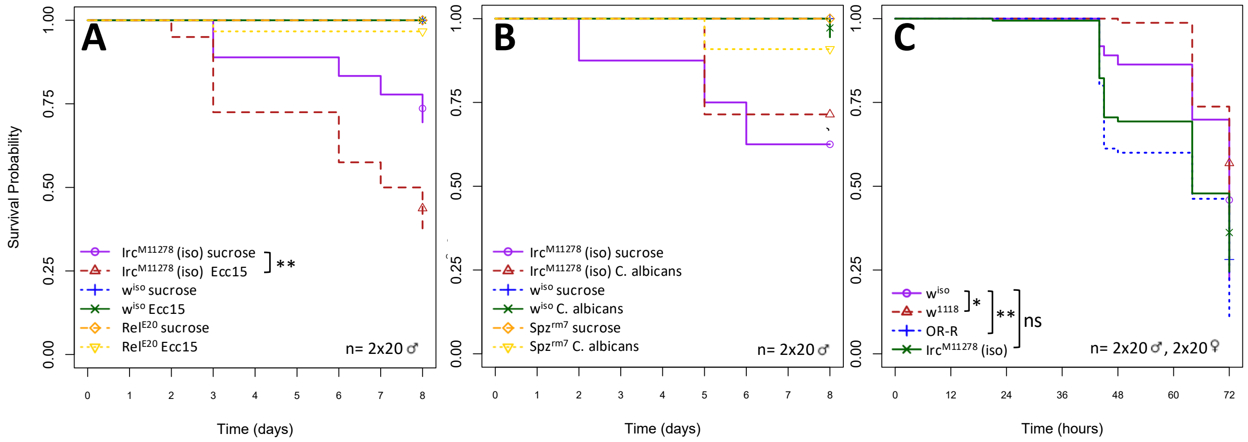

**Figure 4.** Survival of *Irc*$^{MB11278}$ (iso) mutant flies to oral treatment with two benign microbes and 4% $H_2O_2$. **A)** *Irc* flies fed *Ecc15* ($OD_{600}$=400 in 5% sucrose, 25°C) died significantly more than *Irc* flies fed sucrose. **B)** Feeding with *C. albicans* shows that both challenged and unchallenged *Irc* flies had a higher rate of death than wild-type (*w*$^{1118}$) and immune-deficient (*spz*$^{rm7}$) flies, but this

difference was not significant. **C)** Flies of all genotypes fed 4% $H_2O_2$ quickly developed severe locomotor defects, became stuck by the wings and died. Differences in survival calculated using a Cox Proportional Hazards test. * P<0.05, ** P<0.01, *** P<0.001.

#### Conclusion

Due to the fact that *Irc*$^{MB11278}$ flies always survived at least 24h prior to death after becoming stuck to the substrate by their wings, we do not attribute the increased mortality in *Ecc15*-fed *Irc*$^{MB11278}$ flies primarily to pathogen ingestion, but rather to locomotor defects. The difference in survival between sucrose-fed and *Ecc15*-fed *Irc*$^{MB11278}$ flies may be explained by the increased stickiness of the *Ecc15*-containing substrate compared to the sucrose-containing substrate. We conclude that contrary to (Ha et al., 2005), loss of *Irc* does not markedly increase susceptibility of flies to feeding with a wide range of non-pathogenic microbes.

### Supplementary S23: Duox and reactive oxygen species (ROS) in gut epithelial immunity

**Abstract**

The characterization of the role of Duox was undoubtedly an important landmark in the characterization of *Drosophila* gut immunity. Early experiments found that ingestion of pathogenic bacteria by adult flies resulted in rapid upregulation of *Duox* expression in the midgut and an associated *Duox*-dependent reactive oxygen species (ROS) burst as shown by ubiquitous knockdown of this gene; these authors inferred that ROS produced by *Duox,* particularly HOCl, had highly bactericidal effects (Ha et al., 2005). However, there is little independent corroboration of a direct role of *Duox* in gut immunity. *Duox* has other important roles which may indirectly affect immune function. Developmental and signaling roles of ROS may contribute to phenotypes currently assigned only to microbicidal activity of ROS in the gut, as many of the conclusions on the roles of *Duox* in the gut are exclusively based on ubiquitous knockdown affecting all tissues and life stages. Moreover, some studies highlight the role of the alternate *Drosophila* oxidase Nox in intestinal stem cell proliferation as well as the role of Duox in producing ROS that acts as a signaling molecule to promote gut homeostasis (Du et al., 2016; Iatsenko et al., 2018; Jones et al., 2013).

In the framework of ReproSci, we address the evidence on the roles of *Duox* and ROS in gut epithelial immunity, particularly: **i)** Is the ROS burst in the gut following oral infection dependent on *Duox* in the gut? **ii)** Are ROS in the gut bactericidal? **iii)** Are the effects of *Duox* knockdown on gut epithelial renewal direct, or due to roles of this gene in other tissues and processes?

Data from recent articles and experiments performed for ReproSci do not support a major role of Duox in the midgut in the production of microbicidal ROS. They also suggest that the effects of Duox on gut epithelial renewal are not caused by Duox activity directly in the midgut but rather indirectly due to Duox activity in the Malpighian tubules (and possibly other tissues). Thus, data on the essential role of *Duox* in the gut upon oral infection remain contradictory. More precise experiments are needed to clearly define the role of *Duox* in gut epithelial immunity, which may be more complex than currently thought.

**Introduction**

In a landmark paper, Ha et al., (2005a) found that Dual oxidase (Duox) generates an epithelial oxidative burst that limits microbial proliferation in the gut. This revealed that oxidant-mediated antimicrobial responses are not restricted to the phagocytes, but also function in mucosal barrier epithelia. Studies in other insects have confirmed this observation, pointing to a role of ROS in gut immunity and microbiota control. Recent studies have also highlighted additional roles of Duox, some of which could impact gut immunity, that are discussed in this section.

*Duox* gene expression appears to have complex regulation and participates in many processes. *Duox* is an essential gene for *Drosophila* development as null mutations are lethal at larval stages (Flybase, (Anh et al., 2011; Xie et al., 2010)). For this reason, ubiquitous or tissue-specific RNAi that leaves sufficient expression for essential developmental functions is used to study functions of this gene in adults. However, this introduces the complication that observed effects of *Duox* knockdown may be due to essential roles of this gene in developmental processes, rather than a direct role in gut immunity.

In addition to its initial proposed role in midgut immunity, evidence in *Drosophila* and other insects show that *Duox* has essential roles in cuticle cross-linking and sclerotization, including stabilization of the eggshell (Dias et al., 2013), exterior cuticle (Anh et al., 2011; Lee and Kim, 2014; Zuber et al., 2019), tracheae (Jang et al., 2021; Kizhedathu et al., 2021; Zuber et al., 2019), wings (Anh et al., 2011; Hurd et al., 2015), and peritrophic matrix (Kumar et al., 2010 : Bai et al. 2025). Studies in the stink bug *Riptortus pedestris* have shown that expression of *Duox* in the tracheae during development strongly influences gut microbiome composition in the adult due to altered oxygenation of the gut, and that knockdown causes dysbiosis (Jang et al., 2021). Mutation of *Duox* has been similarly shown to impair tracheal development in *Drosophila* (Kizhedathu et al., 2021; Zuber et al., 2019). Although studies have not yet examined whether tracheation and oxygenation of the gut influences gut microbiome composition in *Drosophila*, a similar connection would be logical as bacteria differ in oxygenation requirements as they do in their responses to antimicrobial peptides and ROS (Sekihara et al., 2016). Furthermore, recent studies in *Drosophila* have shown that tracheation affects epithelial cell division and renewal in the gut (Perochon et al., 2021; Tamamouna et al., 2021), which is known to alter the gut environment and affect composition of the microbiome (Buchon et al., 2009) (see also (Hochmuth et al., 2011)).

It is therefore possible that loss of *Duox* in *Drosophila* affects stabilization of the peritrophic matrix and increases permeability of the gut barrier, or reduces tracheation of the gut. This suggests that dysbiosis and mortality observed in Duox-RNAi flies may be due to indirect effects rather than a direct need for *Duox*-mediated production of bactericidal HOCl. Few studies have even tangentially addressed the role of peritrophic matrix crosslinking in *Drosophila* gut immunity (but see (Kuraishi et al., 2011; Shibata et al., 2017; Xiao et al., 2017)), highlighting the need for specific studies in *Drosophila* to determine whether loss of *Duox* impairs development of the peritrophic matrix or oxygenation of the gut. Moreover, we cannot exclude that some of the effects of Duox on the gut are due to its established roles in hemocytes, which are known to impact the gut (Ayyaz et al., 2015; Chakrabarti et al., 2016; Chakrabarti and Visweswariah, 2020).

Recent evidence also suggests lumenal ROS produced by Duox contributes to bacterial clearance in a signaling role, regardless of potential direct antimicrobial activity. HOCl produced by Duox in the enterocytes directly activates the TrpA1 receptor in enteroendocrine cells, promoting defecation and clearance of pathogenic bacteria by stimulating contractions of the visceral muscles (Benguettat et al., 2018; Du et al., 2016). This response promotes rapid early clearance of both Gram-positive *Bacillus thuringiensis* and Gram-negative *Ecc15* (Benguettat et al., 2018). A similar process involving TrpA1 also promotes intestinal stem cell (ISC) proliferation in the midgut (Xu et al., 2017). These processes likely contribute to preventing dysbiosis in both homeostatic and infective conditions, independent of direct antimicrobial roles of ROS and Duox. Finally, Rubinić et al. (2026) recently revealed a role for Duox in defecation following *P. entomophila* challenge. They also reported that Duox did not contribute to the strong ROS induction observed upon *P. entomophila* ingestion (Rubinić et al. 2026).

Furthermore, some roles initially attributed to Duox, notably control of the gut microbiota and stem cell proliferation, may instead be roles of the alternate oxidase Nox (Iatsenko et al., 2018; Jones et al., 2013; Patel et al., 2019). Using Hydro-Cy3 staining for intracellular ROS, Jones et al. (2013) showed that *Lactobacillus plantarum* but not *Ecc15* or *Bacillus cereus* induced a ROS burst in *Drosophila* larva and adult midgut upon enteric infection, and that Nox is responsible for this burst.

Finally, a recent article suggests that contrary to (Ha et al., 2005), **i)** Duox is not upregulated in the midgut upon oral bacterial infection but rather in the Malpighian tubules, and **ii)** that oral bacterial infection induces a strong ROS burst in the Malpighian tubules rather than the midgut (Liu et al., 2024). Moreover, this article shows that increased epithelial renewal in response to oral infection with *Ecc15* is not attributable to Duox-mediated production of ROS in the midgut as previously suggested using a ubiquitous silencing of Duox (Buchon et al., 2009). Instead, (Liu et al., 2024) find that Duox in the Malpighian tubules produces ROS that acts as a signal leading to the release of Upd3, which then stimulates epithelial renewal in the gut through a countercurrent flow mechanism.

To summarize, in addition to the bactericidal role proposed by Ha et al., (2005), Duox has the potential to influence gut homeostasis, mucosal immunity and microbiota composition in multiple ways, including through roles in **i)** crosslinking of the peritrophic matrix that promotes impermeability of the gut and prevents inappropriate immune activation in response to gut microbes; **ii)** tracheal development important for oxygenating the gut, enabling regeneration of the gut epithelium in adult flies (see (Tamamouna et al., 2021)) and maintaining an environment favorable to commensals; **iii)** production of HOCl that has a signaling role in promoting bacterial clearance (Du et al., 2016); and **iv)** production of ROS that stimulate epithelial renewal in the gut (Buchon et al., 2009; Du et al., 2016). Finally, the impact of Duox in the midgut could be due to its roles in tissues other than the gut (crop, Malpighian tubules, hemocytes) which may indirectly influence gut activity (Liu et al., 2024). These mechanisms are at present poorly studied in *Drosophila,* but should be considered in future studies to clarify direct 'immune' roles of *Duox* and ROS. In the following sections, we present evidence that contradicts a major direct role of Duox in the midgut in the production of microbicidal ROS.

#### Methods

*Drosophila stocks*

*Drosophila* stocks were maintained at 25°C on standard fly medium made of 6% cornmeal, 6% yeast, 0.62% agar, 0.1% fruit juice, supplemented with 10.6 g/L moldex and 4.9 ml/L propionic acid. In all experiments, $w^{1118}$ DrosDel flies were used as wild-type control unless otherwise stated (Ryder et al., 2004). Crosses with *Gal4* and infection were performed at 29°C. Fly strains used in the study are: isogenic DrosDel $w^{1118}$ from Luis Teixeira, *uas-Duox IR* from Won Jae Lee (Ha et al., 2005), *da-Gal4* from Bruno Lemaitre lab, *uas-Duox IR* (BL38916, BL38907) from Bloomington stock center.

*Oral infections*

Oral infections with *Pectobacterium carotovorum 15* (*Ecc15*) were performed using 20 5-10 day-old adult female flies. The flies were starved for 2 h at 29°C in an empty vial and then transferred to a vial containing the infection solution and maintained at 29°C (Neyen et al., 2014). The infection solution was obtained by mixing an equal volume of a concentrated overnight culture of *Ecc15* (optical density $OD_{600}$ = 200) with a 5% sucrose solution (1:1) and deposited on a filter paper completely covering the surface of the fly medium. For experiments described in Figure 2, flies were fed continuously with *Ecc15-GFP* by flipping flies on a new vial with a filter soaked with the *Ecc15-GFP*/sucrose solution.

Survival assays

For survival analysis, flies were maintained in vials containing filter paper saturated with infectious solution, which was replaced daily throughout the experimental period. Mortality was monitored at regular intervals, with dead flies being counted and removed during each observation time point.

Bacterial persistence assay

*For the bacterial persistence assay, filter papers containing the infectious solution were replaced daily. Homogenates from five surface-sterilized intact fly intestines collected at different time points post-oral infection were serially diluted, cultured overnight at 29°C, and* colony forming units (CFUs) were calculated*. All experiments were performed using rifampicin-resistant bacterial strains on selective agar plates.*

Fly and gut imaging

*Flies collected at designated feeding time points were placed on ice for immobilization, followed by* microscopy imaging (Olympus, SZX7, Japan)*. Subsequently, adult guts were dissected and photographed using the same imaging protocol.*

*RNA extraction and quantitative PCR*

For RNA extraction, 10 whole insects, 20 guts, and 40 MpT were used. Total RNA (100 ng for MpT or 500 ng for whole insects and guts) was reverse-transcribed using PrimeScript RT reagent (TAKARA) with a mix of oligo-dT and random hexamer primers. Quantitative PCR (qPCR) was performed in SYBR Green Mix (Vazyme, China). Reactions were carried out on an FQD-96X system (Bioer, China): 95 °C for 3 min (initial denaturation), followed by 40 cycles of 95 °C for 15 s and 60 °C for 30 s. Relative gene expression was calculated using the $2^{-\Delta\Delta Ct}$ method, with **RpL49** as the reference gene. All experiments included at least three independent biological replicates.

*Data statistics and analysis*

The software GraphPad 9.0 was used for data analysis and the appropriate data method is shown in the legends. Each experiment was independently performed in at least two replicates.

#### Results

**i) Is *Duox* upregulated in the gut epithelium upon oral infection? Is the ROS burst in the gut following oral infection dependent on *Duox* expression in the gut?**

Many groups have demonstrated a ROS burst in the gut shortly following oral infection or gut damage of adult (Buchon et al., 2009; Chakrabarti et al., 2014, 2012; Ha et al., 2005; Hori et al., 2018; Xiao et al., 2017) or larval (Ramond et al., 2021) *Drosophila* and other insects (Huang et al., 2020; Sajjadian and Kim, 2020; Yao et al., 2016). Subsets of these have found that *Duox* is upregulated in the insect gut tissue following oral infection or intestinal damage (Buchon et al., 2009; Chakrabarti et al., 2014; Ha et al., 2005; Ramond et al., 2021; Sajjadian and Kim, 2020; Yao et al., 2016), and that *Duox* is required for the post-infection ROS burst (Ha et al., 2005; Sajjadian and Kim, 2020; Xiao et al., 2017). Moreover, suppression of *Duox* or gut ROS has been shown to reduce survival to some oral infections in *Drosophila* and other insects (Ha et al., 2005; Ramond et al., 2021; Sajjadian and Kim, 2020). However, upregulation of *Duox* following oral infection was very variable and weak and required many replicates to reach significance. In contrast, modulation of *Duox* in response to *Ecc15* oral infection was not found by (Buchon et al., 2009), but this study used later (4h, 16h) time points. High throughput expression data also conversely show downregulation of *Duox* in the gut following oral infection at early time points (FlyGut), and slight upregulation only at late time points (16h) in response to non-pathogenic *Ecc15* and *Pseudomonas entomophila gacA* (a non-pathogenic *P. entomophila* variant). Flygut-seq (Dutta et al., 2015) shows that oral infection of adult flies with *P. entomophila* results in upregulation of *Duox* at 48hpi in intestinal stem cells (ISCs), enteroblasts (EBs), and enteroendocrine cells (EEs), but downregulation in enterocytes.

A recent article suggests that enteric infection induces a strong ROS burst in Malpighian tubules (MpT) and not the midgut through Duox. A DHE dye specific for intracellular superoxide showed that ingestion of *Ecc15* induced a ROS burst at an early time point (1.5 hpi) in the MpT, but not in the gut (Liu et al., 2024 - Figure 2B,D). Furthermore, the ROS level was much lower in the midgut than in the MpT both before and after infection, suggesting the MpT are the primary tissue of ROS production ((Liu et al., 2024) Figure 2C). A fluorometric hydrogen peroxidase assay confirmed that the MpT produced more hydrogen peroxide than the midgut. Knocking down *Duox* but not *Nox* using the MpT principal cell-specific driver *uro-GAL4* significantly decreased ROS levels in the tubules after infection ((Liu et al., 2024) Figure 2E). Collectively, these data suggest that oral infection with *Ecc15* induces a ROS burst through Duox in the MpT rather than the gut. One possible explanation for this discrepancy might be that studies showing upregulation of *Duox* in the gut and Duox-mediated ROS production may not have excluded Malpighian tubules and crop tissues, which are attached to the gut. However, some authors who found upregulation of *Duox* in the larval midgut clearly considered and avoided this complication (Ramond et al., 2021).

Considering the conflicting results surrounding *Duox* upregulation upon oral infection, we decided to test upregulation of *Duox* in response to *Ecc15* oral infection in the gut and Malpighian tubules.

**Results**

Experiments performed for ReproSci show that enteric infection with *Pectobacterium carotovorum Ecc15* leads to upregulation of *Duox* expression in the MpT but not the midgut. RT-qPCR of extracts from midgut (without crop and MpT) show an increase in *Duox* gene expression in the MpT, but not the midgut (Figure 1) upon oral *Ecc15* infection of adult female flies.

**Conclusion**

There is mixed set of evidence for a Duox-mediated ROS burst in the midgut and corresponding upregulation of *Duox* in the gut upon ingestion of pathogens. The data summarized here suggest that contrary to (Ha et al., 2005), *Ecc15* infection upregulates *Duox* gene expression in the Malpighian tubules and not the midgut. This is consistent with the findings of (Liu et al., 2024), who show a *Duox*-dependent ROS burst in the Malpighian tubules rather than the gut following infection, which serves in a signaling role to stimulate epithelial renewal through Upd3.

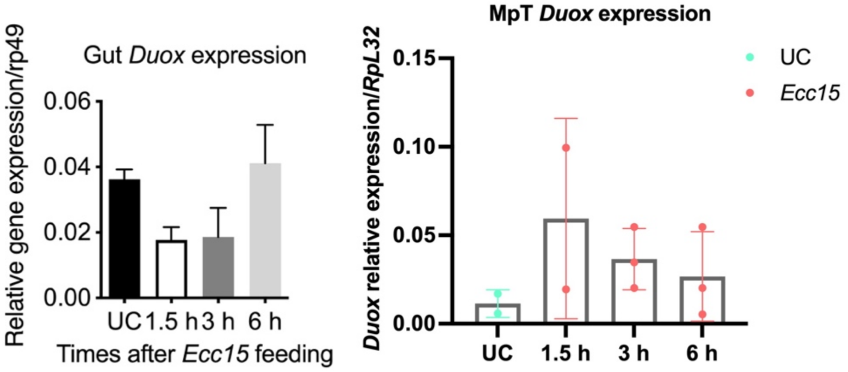

**Figure 1. Enteric infection induces upregulation of *Duox* in the MpT but not the gut.** qRT-PCR of *Duox* expression in the adult gut **(left)** and Malpighian tubules **(right)** at different time points after *Ecc15* oral infection. The crop was excluded from the samples. Error bars represent the standard deviation of three independent experiments.

**ii) Are ROS produced by Duox in the gut bactericidal?**

Ha et al. previously found that *Pectobacterium carotovorum* GFP (*Ecc15-GFP*) proliferated in the gut of *Drosophila* when *Duox* was silenced by RNAi (Ha et al., 2005). They concluded that Duox produces lumenal microbicidal ROS, particularly HOCl, that contribute to host defense in the gut. These results are consistent with data showing that reduced *Duox* expression causes dysbiosis (atypical proliferation of minor and pathogenic community members), and this dysbiosis results in immune activation in a variety of insect models (Chmiel et al., 2019; Diaz-Albiter et al., 2012; Ha et al., 2009, 2005; Henry et al., 2020; Huang et al., 2020; Kumar et al., 2010; Xiao et al., 2017; Yao et al., 2016). Dysbiosis also occurs when ROS is reduced using a scavenger (a chemical compound that neutralizes ROS) (e.g. (Diaz-Albiter et al., 2012; Ramond et al., 2021; Sajjadian and Kim, 2020)). Dysbiosis is typically characterized by an increase in total microbial load in the gut, an increase in microbial diversity, and expansion of pathobiont species that have immunogenic effects as they become dominant community members.

It should be noted that biological HOCl was initially thought to contribute to bacterial killing primarily through protein degradation and cell lysis. However, further investigation suggests that the role of HOCl may be primarily bacteriostatic at physiological concentrations and serves a complementary role to other defense systems (Dupré-Crochet et al., 2013; McKenna and Davies, 1988; Ulfig and Leichert, 2021). Differences in susceptibility of various species of bacteria (both pathogenic and commensal) to HOCl activity (e.g. (Sekihara et al., 2016)) is likely one of many factors that exerts pressure on the bacterial community and acts to maintain homeostasis, preventing dysbiosis and associated activation of immune programs. The available data suggest that regardless of source, HOCl in the *Drosophila* gut lumen has bacteriostatic if not bactericidal effects on bacteria in the gut. As discussed above, lumenal HOCl produced by Duox also contributes to bacterial clearance and epithelial renewal in a signaling role through TrpA1, regardless of its potential bacteriostatic activity (Benguettat et al., 2018; Du et al., 2016; Xu et al., 2017).

Here, we have examined whether Duox in the midgut affects bacterial persistence following oral infection, reproducing the experiments described by (Ha et al., 2005).

**Results**

(Ha et al., 2005) found that *Ecc15-GFP* bacteria over-proliferates in the gut of ubiquitous *Duox*-RNAi flies, and concluded that ROS produced by Duox are directly bactericidal. Since *Duox* null mutations are lethal, we similarly analyzed *Ecc15* persistence in the gut of *Da-Gal4>UAS-Duox* IR flies using two *Duox-IR* constructs, and used a continuous mode of infection as in (Ha et al., 2005). Overexpression of BDSC #38916 *Duox-IR* and BDSC #38907 *Duox-IR* with the ubiquitous *Da-Gal4* driver successfully reduced *Duox* gene expression in whole flies (Figure 2A). However, in our hands reduced *Duox* expression did not result in any striking increase in the proliferation of *Ecc15-GFP* in the midgut at 6h, 24, and 48h post-infection (Figure 2B, D and E), in sharp contrast to (Ha et al., 2005). When using the Mann-Whitney U test, we still observed a statistically significant difference in CFU counts at 6 hpi in Figure 2B, as well as at both 6 hpi and 24 hpi in Figure 2F. However, the magnitude of these differences is relatively small. Furthermore, silencing *Duox* did not affect fly survival to *Ecc15* (Figure 2C). Finally, we repeated assays for *Ecc15-GFP* persistence in *Duox-RNAi* flies using the same Duox-IR construct from (Ha et al., 2005) and did not observe any impact (Figure 2F).

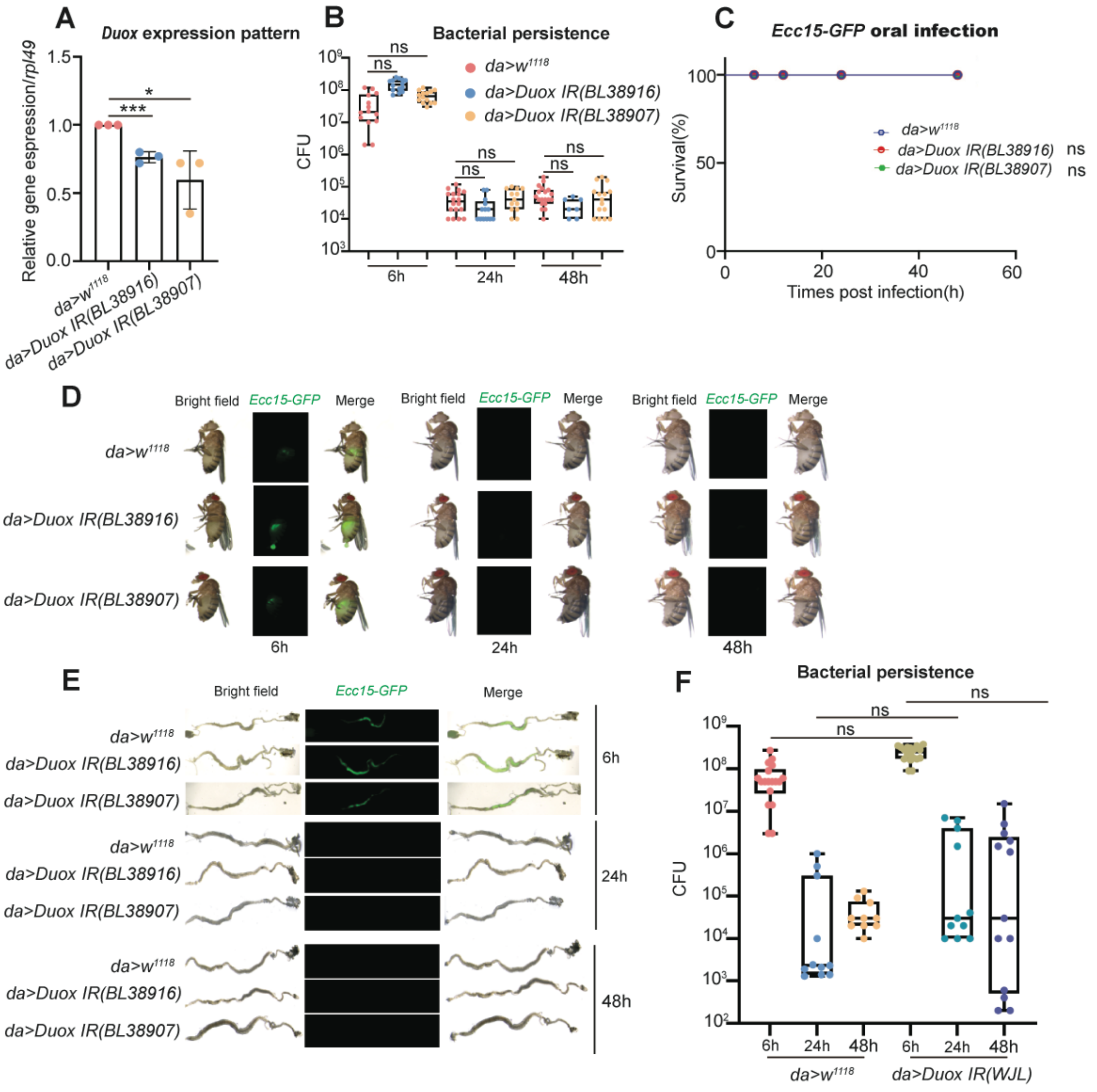

**Figure 2. Ubiquitous *Duox* knockdown does not affect *Ecc15* bacterial persistence in the midgut**

**A)** RNAi efficiency of two *Duox-IR* constructs (BDSC 38916 and BDSC38907) using a *da-GAL4* ubiquitous driver. *Duox* gene expression was monitored by RT-qPCR in whole flies raised at 29°C.

See also (Benguettat et al., 2018) for RNAi efficiency of BDSC38907, which is the most efficient (Armel Gallet, personal communication). **B)** CFU counts of midguts of *Duox RNAi* flies did not reveal any impact of *Duox* silencing on the persistence of *P. carotovorum* (*Ecc15-GFP*). **C)** Ubiquitous silencing of *Duox* did not affect the survival of flies upon oral infection with *Ecc15-GFP.* **D)** Representative images of wild-type and *Duox RNAi* females collected at 6, 24 and 48 post oral infection with *Ecc15-GFP*. No effect of *Duox* silencing was observed. **E)** Representative images of dissected midguts from wild-type and *Duox RNAi* females collected at 6, 24 and 48 post oral infection with *Ecc15-GFP*. No effect of *Duox* silencing was observed. **F)** CFU counts of midguts of *Duox RNAi* flies did not reveal any impact of *Duox* silencing on the persistence of *P. carotovorum* (*Ecc15-GFP*). This experiment was performed with the Duox-IR used by (Ha et al., 2005). A was analyzed using Unpaired Student's *t*-test. B and F were analyzed using Kruskal–Wallis test with post-hoc Dunn's test,. C was analyzed using Log-rank (Mantel-Cox) test. * $p<0.05$, *** $p<0.001$, ns non-significant.

**Conclusion**

Our investigations did not reveal a clear role of Duox in the control of *Ecc15,* suggesting that contrary to (Ha et al., 2005), ROS produced by Duox are not microbicidal against this bacterium. Note that Benguettat et al. (2018) did observe an effect of *Duox* on the persistence *Bacillus thuringiensis* and *Ecc15* but only between 2 and 6h post infection. These authors infected wild-type and *Duox RNAi* flies for 30 min after 2 hours of starvation. They attribute the impact of Duox on bacterial persistence not to the production of microbicidal ROS, but to reduced visceral muscle spasms and expulsion of bacteria upon silencing of *Duox*. Collectively, our data contradicts the claim that Duox produces microbicidal ROS in the gut to control bacterial infection. We cannot however exclude that the discrepancy between our results and those of (Ha et al., 2005) stems from differences in fly food, microbiota or other factors. Our study does not exclude that ROS produced by Duox has a short-term impact on bacterial persistence through stimulation of muscle contraction as suggested by Benguettat et al., (2018), or via a direct bacteriostatic effect (Sekihara et al., 2016).

**iii) Are the effects of *Duox* knockdown on gut epithelial renewal in response to bacterial infection direct, or due to roles of this gene in other tissues and processes?**

The role of Duox in epithelial renewal was established based on the observation that ubiquitous silencing of *Duox* by RNAi reduces stem cell proliferation upon bacterial infection (Buchon et al., 2009). Subsequently, several papers found a role for Nox but not Duox in ROS generation in the intestine and stimulation of intestinal stem cell proliferation (Iatsenko et al., 2018; Jones et al., 2013; Patel et al., 2019). This role for Nox does not rule out a role for Duox in response to gut bacteria, but does suggest that Duox in the gut is not the primary source of ROS in all cases. On this line, (Amcheslavsky et al., 2020) have proposed that enterocytes destined to die transiently enter an undead-like state through Myo1D-dependent membrane localization of Dronc, which leads to the production of ROS by Duox. HOwever, ROS produced by enterocytes have a signaling role attracting hemocytes to the gut.

Using tissue-specific RNAi (Liu et al., 2024) find that ROS-mediated intestinal stem cell proliferation upon oral infection does not require Duox in the gut. PH3 staining for cells undergoing mitosis showed that Duox was required only in the early phase of epithelial renewal at 16 hours post infection (hpi), as previously described by (Buchon et al., 2009). PH3 staining showed that epithelial renewal was unaffected by RNAi against *Duox* in gut cell types, whereas knocking down *Nox* either in progenitors or enterocytes reduced stem cell proliferation, consistent with previous studies (Jones et al., 2013; Patel et al., 2019). Furthermore, (Liu et al., 2024) show that Duox is necessary in the MpT for gut homeostasis after infection. The number of mitotic intestinal stem cells was significantly reduced when *Duox* was silenced in MpT using the *Uro-Gal4* driver ((Liu et al., 2024) Figure 3A) mimicking the results observed with the ubiquitous driver *Da-Gal4* ((Liu et al., 2024) Figure 1A). These data confirm the involvement of Duox in the early phase of infection-induced stem cell proliferation, but also suggest that, unlike Nox, Duox is not required in midgut epithelium homeostasis as initially proposed, but rather in the MpT.

These results suggest that Duox in the MpT impacts gut epithelium renewal upon oral bacterial infection, while in the midgut epithelium Nox contributes to proliferation. This contradicts the Buchon et al. (2009) interpretation that renewal is triggered by a ROS burst that damages the midgut epithelium, and instead show that ROS produced by Duox in the MpT lead to Upd3 production that stimulates mitosis in the gut. It is possible that ROS produced by Duox in the crop, where it is highly expressed (Leader et al., 2018), may also affect midgut epithelium renewal. *Duox* and ROS may also play a role in JNK-mediated cell turnover in other contexts, including cell death and proliferation in developmental, regenerative and tumorous contexts (Fogarty et al., 2016; Khan et al., 2017; Santabárbara-Ruiz et al., 2015).

**Final assessment from ReproSci**

Experiments performed in the framework of ReproSci and recent results by (Liu et al., 2024) contradict some key conclusions regarding the role of Duox in midgut ROS production and enteric infection. These data do not support a major direct role of Duox in the midgut in the production of microbicidal ROS. These data do not exclude bacteriostatic or signaling roles for Duox in the gut. One of the most attractive hypotheses that reconciles much of the literature and is supported by two independent groups is that enterocytes of the midgut produce a low level of ROS that regulates defecation by stimulating gut muscle contraction, which aids in clearing potentially pathogenic bacteria from the gut and preventing establishment of infection (Du et al., 2016; Benguettat et al., 2018). New data from (Liu et al., 2024) also suggests that the effects of Duox on gut epithelial renewal is not caused by damage caused by Duox-mediated ROS activity in the midgut but rather due to a signaling role in the Malpighian tubules. We conclude that data on a direct role of *Duox*-mediated ROS in the gut upon oral infection are contradictory, and that more precise experiments are needed to clearly define the role of *Duox* in gut epithelial immunity, which may be more complex than currently thought. We encourage authors to be careful in considering the literature on Duox, which requires more careful investigation given the many physiological roles of Duox.

### Supplementary S24: Contribution of NOS to gut-fat body signaling

**Abstract**

(Foley and O'Farrell, 2003) found that induction of both Dpt-lacZ and Drs-lacZ protein reporters (gene promoter sequences fused to a lacZ coding region) in larvae by natural (oral) *Ecc15* infection was suppressed by chemical inhibition of NOS activity using L-NG-Nitroarginine Methyl Ester (L-NAME). In contrast, (Chakrabarti et al., 2012) found no difference between Antimicrobial peptide (AMP) gene expression in wild-type and $NOS^{C}$ adults and larvae in response to oral and septic *Ecc15* infections. To clarify these apparently contradictory results, we tested survival of adult flies to a variety of pathogens and AMP induction by oral infection with *Ecc15* in larvae using $NOS^{Dall}$ null mutants. Use of a $NOS^{Dall}$ did not strongly affect expression of *Diptericin* upon natural infection with *Ecc15* and $NOS^{Dall}$ null flies did not show susceptibility to *Ecc15* or *Enterococcus faecalis.* We conclude that NOS is unlikely to be a key regulator of Imd pathway upon natural infection. It remains possible that NOS plays other roles in immunity in specific tissues or contexts.

**Introduction**

Several studies have found a link between NOS and the immune response in *Drosophila.* (Foley and O'Farrell, 2003) found that inhibiting endothelial NOS enzyme activity with L-NG-Nitroarginine Methyl Ester (L-NAME) in larvae and adults increased susceptibility to *Ecc15,* reduced activity of a *Dpt-lacZ* reporter, and prevented *spz* mutants from controlling bacterial growth. This is similar to the finding of (Glittenberg et al., 2011) that NO and the Toll pathway function in concert to activate systemic immunity in larvae following natural infection with *Candida albicans.* (Eleftherianos et al., 2014), who used two Bloomington Minos insertion alleles as mutants of NOS ($NOS^{24283}$ and $NOS^{53156}$) found slightly increased susceptibility to all bacteria tested (*Escherichia coli, Micrococcus luteus, Photorhabdus luminescens, Enterococcus faecalis*) in NOS mutants, and also in wild type strains with naturally lower NO levels. (McGettigan et al., 2005) found that fly survival in response to septic infection with *E. coli* was increased by overexpression of NOS in the Malpighian tubules, which also increased basal and induced levels of *Diptericin.*

Most of these results suggest that functional NOS activity promotes survival to a variety of microbial infections. Furthermore, they largely show that this is at least partially mediated by interaction of NOS with canonical immune pathways (Imd and Toll) and perhaps by modulation of AMP production. However, other studies failed to find a link between NOS and increased immune function. (Broderick et al., 2006) found that scavenging NO with combinamide or injection with low-dose L-NMMA (an inhibitor of all NOS enzyme activity) improved survival of flies to injection with high doses of *S. aureus* or *E. coli,* and similarly improved survival of *imd* mutant flies to bacterial injection. (Chakrabarti et al., 2012) found no difference between AMP gene expression in wild-type and $NOS^{C}$ adults and larvae in response to oral and natural *Ecc15* infections.

We set out to test the role of NOS in Imd pathway activation upon natural infection, as this finding is not consistent with the notion that natural infection is initiated by diffusion of peptidoglycan fragments from the gut to the hemolymph, which then activate Imd receptors in the fat body (Neyen et al., 2012; Zaidman-Rémy et al., 2006; Bosco-Drayon et *al.*, 2012; Charroux et al., 2017.). For this, we tested survival of $NOS^{Dall}$ null flies to *Ecc15* and *E. faecalis* and monitored AMP induction by oral bacterial infection in $NOS^{Dall}$ larvae.

**Methods**

*Fly lines and husbandry*

$NOS^{Dall}$ flies were obtained from (Rabinovich et al., 2016). $w^{1118}$ flies (BDSC #3605) were used as a control. Flies were raised on cornmeal fly medium (6% cornmeal, 6% yeast, 0.6% agar, 0.1% fruit juice, 10.6 g/L moldex and 4.9 mL/L propionic acid) and transferred to fresh medium every 2-3 days.

*Survivals*

3 to 5-day old male a flies reared on cornmeal fly medium at 25°C were anaesthetized with $CO_2$ and pricked on the thorax anterior and ventral to the wing joint with a 0.2 mm tungsten needle dipped in a concentrated overnight (shaking) culture of Pectobacterium carotovorum *Ecc15* (LB medium, 29°C) or *E. faecalis* (ATCC 29212, BHI medium, 37°C) diluted to the OD indicated in the figure in PBS. Flies were incubated at the temperature indicated in the figures and scored daily for survival; flies were flipped to new medium during the survival every 2-3 days. At least 20 flies per genotype were scored in 1-2 independent experiments; Data were analyzed using a Log-rank test.

*Larval oral infections*

For each genotype, ~60-100 synchronized early L3 larvae were washed out of the food and stirred into 200 µL of $OD_{600}$=400 *Ecc15* mixed with 400 µL of banana puree in an Eppendorf. The Eppendorf was plugged with foam to allow oxygen diffusion and left at room temperature for 30 minutes, then the contents were transferred to a vial containing cornmeal fly medium and incubated at 29°C. At the desired time points, 7-10 larvae per treatment were frozen and processed for RT-qPCR.

*qPCR for AMP induction*

Total RNA was extracted from samples using TRIzol reagent (Invitrogen) according to manufacturer instructions, and re-suspended in MilliQ water. DNase treatment was performed using TURBO DNA-free kit (Invitrogen). Reverse transcription was performed using 0.5 mg total RNA in 10 µl reactions using Prime-Script RT (Takara) with random hexamer and oligo dT primers. Quantitative PCR was performed on a LightCycler 480 (Roche) in 96-well plates using PowerUp SYBR Green Master Mix (Applied Biosystems). AMP expression was normalized to expression of the ribosomal protein *Rpl32* (*rp49*).

- *Rp49* Forward: 5' – GCC GCT TCA AGG GAC AGT ATC TG – 3'
- *Rp49* Reverse: 5' – AAA CGC GGT TCT GCA TGA G – 3'
- *Diptericin A* Forward: 5' - ATG CCC GAC GAC ATG ACC AT – 3'
- *Diptericin A* Reverse: 5' - TTG TCG GTG GTC CAC ACC TT – 3'
- *NOS* Forward: 5' - ACA ATG GCC GAG CAT GTG TA– 3'
- *NOS* Reverse: 5' - TGG CCC TTG ACT TTG TGT GT– 3'

#### Results

*Loss of NOS activity does not impair Diptericin gene expression upon natural infection with Ecc15*

To test the role of NOS in the activation of the Imd pathway, we used the viable null $NOS^{Dall}$ mutant described in (Rabinovich et al., 2016). We confirmed the absence of *NOS* transcript in $NOS^{Dall}$ larvae. We then monitored the expression of the *Diptericin* gene by RT-qPCR upon natural infection with *Ecc15* and found that *Dpt* was induced in both $NOS^{Dall}$ and wild-type larvae (Figure 1).

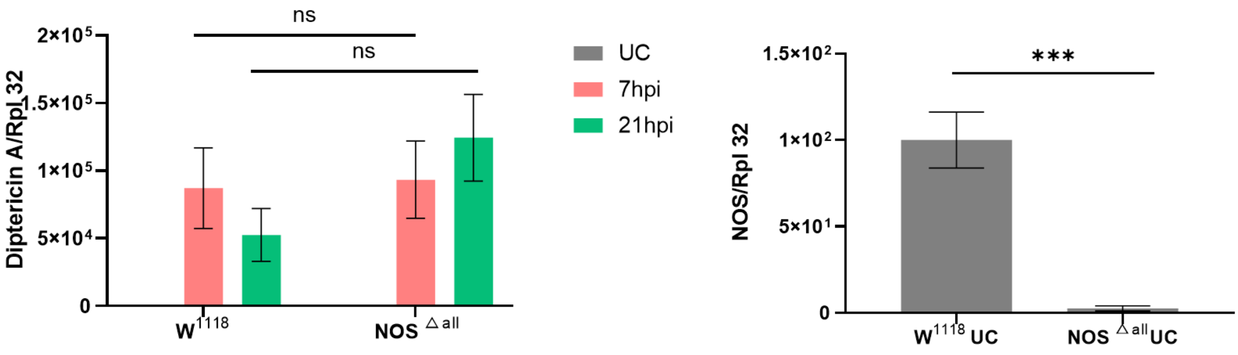

**Figure 1. (left)** Diptericin expression in wild-type ($w^{1118}$) and null $NOS^{Dall}$ mutant L3 larvae after oral infection with *Ecc15* and incubation at 29°C. **(right)** RT-qPCR confirms that $NOS^{Dall}$ larvae do not express the *NOS* gene. 10 flies per replicate, number of independent repeats=4. Statistical analysis: t-test, not significant = ns, p<0.001 = ***. Error bars represent the standard deviation.

*NOS-deficient flies do not show acute susceptibility to Ecc15 or E. faecalis*

We monitored the survival of wild-type and $NOS^{Dall}$ flies to *Ecc15* and *E. faecalis* and did not detect any significant difference (Figure 2). As expected, *Relish* deficient flies were very susceptible to Gram-negative *Ecc15* while $Bom^{55C}$ deficient flies had a marked susceptibility to *E. faecalis*.

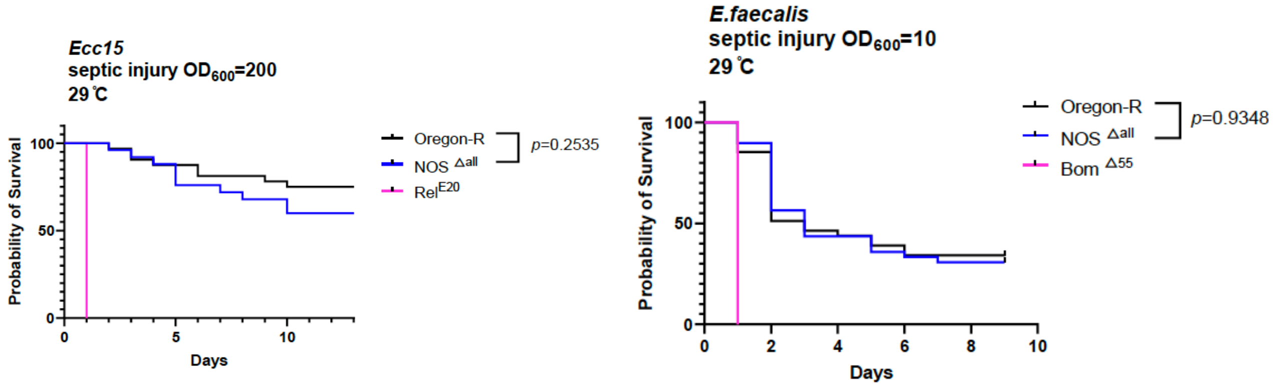

**Figure 2.** Survival of adult $NOS^{Dall}$ mutants to septic injury with the Gram-negative bacterium *Ecc15* ($OD_{600}$=200, 29°C) **(left)** and the Gram-positive bacterium *E. faecalis* ($OD_{600}$=10, 29°C) **(right)**. We did not observe any significant difference between $NOS^{Dall}$ and wild-type. *Relish* ($Rel^{E20}$) and $Bom^{55C}$ flies were used as positive control susceptible respectively to *Ecc15* and *E. faecalis*. 20 flies per replicate, number of repeats=2. Statistical analysis: Log-rank (Mantel-Cox).

#### Conclusion

Contrary to (Foley and O'Farrell, 2003) we do not find any evidence that NOS participates in systemic Imd pathway activation in response to oral infection using a $NOS^{Dall}$ null mutant. This contrast between results obtained by (Foley and O'Farrell, 2003) using AMP reporters and our results assaying *Diptericin* expression by qPCR may indicate that translation is impaired in NOS-deficient larvae. We conclude that NOS is unlikely to be involved in a signaling pathway linking oral *Ecc15* infection to activation of the Imd pathway in the fat body of larvae.

Furthermore, contrary to (Eleftherianos et al., 2014; Foley and O'Farrell, 2003), we find no evidence that fitness or resistance to systemic infection is impaired in adult flies with reduced NOS function. Further studies are needed to clarify the role of NOS, if any, in immunity.

### Supplementary S25: Contribution of *eiger* to survival to infection by various pathogenic bacteria

**Abstract**

*Eiger* (*egr*) is a TNF-related gene which several articles identify as an immune component based on mutant phenotypes. However, recent results showed that widely used *egr* mutants bore a secondary mutation of the Nimrod transmembrane receptor *NimC1*, which has roles in phagocytosis. This calls into question previous findings obtained using the $egr^1$ and $egr^3$ mutants, particularly results concerning immune effects. Survivals using clean $egr^1$ and $egr^3$ mutants with a wild-type *NimC1* locus suggest that both *eiger* and *NimC1* have independent effects on survival to several pathogens, and that the effect of one or both mutations may explain several survival effects previously attributed solely to *eiger*. Contrary to previous results, we find that mutation of *eiger* does not increase tolerance to *Salmonella typhimurium* infection and does not generally increase susceptibility to extracellular pathogens.

**Introduction**

A series of results implicating the TNF-related gene *eiger* in several aspects of *Drosophila* immune function have recently been called into question due to the discovery that widely used *eiger* mutants bore a secondary mutation of the phagocytic receptor *NimC1* (Kodra et al., 2020), which has independent roles in immunity (Melcarne et al., 2019). Because of this, and because most literature on the immune functions of *eiger* has not been independently verified, we set out to determine whether phenotypes previously assigned to *eiger* are reproducibly attributable to loss of either *eiger* or *NimC1*. For clarity throughout this discussion, *eiger* mutants bearing a secondary mutation of *NimC1* will be referred to as $egr^N$.

(Brandt et al., 2004) found that $egr^N$ mutants had prolonged survival but similar bacterial loads when infected with *Salmonella typhimurium*, indicating an increase in tolerance with no change in resistance. (Mabery and Schneider, 2010) found that RNAi of *eiger* in the fat body reproduced this effect. (Schneider et al., 2007) further reported that $egr^N$ mutants had decreased survival to extracellular pathogens *Beauveria bassiana, Enterococcus faecalis, Staphylococcus aureus, Burkholderia cepacia and Streptococcus pneumoniae.* For *B. cepacia* and *S. pneumoniae* but not *E. faecalis*, this was accompanied by increased bacterial growth. In contrast, $egr^N$ mutants were not more susceptible than wild type flies to facultative intracellular pathogens *S. typhimurium, Mycobacterium marinum* or *Listeria monocytogenes*, and had similar bacterial loads as wild-type flies. They concluded that loss of *eiger* increased resistance specifically to intracellular pathogens.

This conclusion was challenged by (Geuking et al., 2009), who did not find any effect of the $egr^N$ mutation on survival to *E. faecalis* and *S. aureus* when compared with the direct parental line. These authors suggested that susceptibility of these flies with respect to other wild-type lines was due to a background effect present in the parental line. This interpretation was subsequently supported by (Kodra et al., 2020), who showed that the $egr^1$ and $egr^3$ alleles used in these studies, and the parental line they were derived from (alternately referred to as *Regg1* or *Df(1)w*), carry a mutation of phagocytic receptor *NimC1* (Melcarne et al., 2019). Furthermore, the key studies showing survival differences did not use a wild-type control with a similar genetic background to the mutants (Brandt et al., 2004; Mabery and Schneider, 2010; Schneider et al., 2007). Background can strongly affect survival results, even among 'wild-type' strains (Apidianakis and Rahme, 2009; Bou Sleiman et al., 2015; Corby-Harris et al., 2007; Eleftherianos et al., 2014; Sasaki et al., 2021; Sorrentino et al., 2004). The secondary mutation of *NimC1* in the $egr^N$ lines and the lack of appropriate background controls complicates interpretation of the results obtained using these mutants, and requires a re-evaluation of the roles of *eiger* and *NimC1* in survival and resistance to bacterial infection.

(Brandt et al., 2004) found that impaired cell entry reduces the lethality of *Salmonella* infection, as *Salmonella* replicates intracellularly. In a follow-up paper, (Schneider et al., 2007) found that the $egr^N$ lines had increased resistance to two other intracellular pathogens, *L. monocytogenes* and *M. marinum,* but did not have increased resistance to a range of extracellular pathogens. *NimC1* has roles in phagocytosis of Gram-negative bacteria (Melcarne et al., 2019) such as *S. typhimurium*; therefore a secondary mutation of *NimC1* might be expected to slow progression of some intracellular pathogens by reducing invasion of hemocytes. However, (Mabery and Schneider, 2010) also found that RNAi of *eiger in vivo* increased tolerance to *S. typhimurium* in the same way as the $egr^{N}$ mutant lines, suggesting that increased resistance may indeed be a direct effect of *eiger* loss, as the RNAi and driver lines were generated in independent genetic backgrounds and are not expected to carry secondary mutations of *NimC1*.

(Mabery and Schneider, 2010) further found that loss of *eiger* resulted in a constellation of phenotypes including increased Imd-mediated antimicrobial peptide expression in the fat body, reduced melanization, and reduced feeding rates. All these factors can affect survival to infection. Our results show that *eiger* flies do not have reduced feeding compared to appropriate controls (See additional data reported on the ReproSci website) but do support reduced melanization upon loss of *eiger* (Supplementary S27). Other authors also support reduced melanization in the absence of *eiger,* and found that this was due to a failure of crystal cell rupture, which reduces the amount of phenoloxidase (PO) released at the melanization site (Bidla et al., 2007). Loss of PO activity can have variable effects on susceptibility to different pathogens, both alone and in combination with other immune defenses (Ayres and Schneider, 2008; Binggeli et al., 2014; Dudzic et al., 2019; Nam et al., 2012; Tang et al., 2006). Therefore, many of the survival effects of *eiger* mutation, if present, may be attributable to reduced melanization.

In light of these considerations, we tested $egr^{1}$ and $egr^{3}$ lines cleaned of the *NimC1* mutation through backcrossing to the *OR-R* wild-type line (Kodra et al., 2020) and independent *NimC1* mutants (Melcarne et al., 2019), in order to determine whether survival, feeding, antimicrobial peptide gene expression and melanization features ascribed to the $egr^{N}$ lines were in fact due to mutation of *eiger*, or alternatively due to loss of *NimC1* or other background effects.

#### Methods

*Fly lines*

$egr^{1}$ and $egr^{3}$ mutants cleaned by backcrossing to *OR-R* were obtained from (Kodra et al., 2020). The original P-element excision mutants are no longer available from Bloomington in the *Df(1)w* background bearing the secondary *NimC1* mutation. The $NimC1^{1}$ mutant was generated in-house in the $w^{1118}$ background (Melcarne et al., 2019). $w^{1118}$ (Bloomington #3605), $w^{iso}$ *Drosdel* (Ryder et al., 2004) and *OR-R* flies were used as wild-types; in survival experiments $spz^{rm7}$ (Neyen et al., 2012), $Bom^{\Delta 55C}$ *Drosdel* (Clemmons et al., 2015; Hanson et al., 2019) or $Dif^{1}$ (BDSC #36559) lines were used as Toll-deficient comparisons. $Relish^{E20}$ ($Rel^{E20}$) flies were used as Imd-pathway deficient mutants. *PPO1/2/3* ($PPO1^{D}$, $PPO2^{D}$, $PPO3^{1}$) triple mutants (Dudzic et al., 2015) were used as a melanization-deficient strain. Flies were reared at 25°C on cornmeal fly medium (6% cornmeal, 6% yeast, 0.6% agar, 0.1% fruit juice, 10.6 g/L moldex and 4.9 mL/L propionic acid), and flipped to new medium every 2-3 days.

*Microbial cultures and survivals*

5 to 7-day old male and female flies reared on cornmeal fly medium at 25°C were anaesthetized with $CO_2$ and pricked on the side of the thorax anterior and ventral to the wing joint with a 0.2 mm tungsten needle dipped in a concentrated bacterial pellet. Survivals were incubated at the temperature indicated in figures and flies were transferred to fresh vials every 2-3 days. ~20 flies were used per genotype per experiment and sex; two or three independent experiments were performed for each treatment. *S. typhimurium SL1344* was cultured as in (Schneider et al., 2007), and diluted in LB medium prior to injection. *S. aureus NCTC 8325, E. faecalis* ATCC 29212*,* and *L. monocytogenes* (CIP 107776, EGDe) were cultured overnight at 37°C shaking in BHI. Cultures were concentrated by centrifugation and diluted to the desired $OD_{600}$ in PBS prior to septic injury. The strain *Beauveria bassiana KN802* was used for infection

#### Results and Discussion

We set out to test survival of $egr^{1}$ and $egr^{3}$ lines and independent *NimC1* mutants to a variety of pathogens. In addition to the *OR-R* background control for the *eiger* lines and $w^{1118}$ control for the *NimC1* line, we also included the $w^{iso}$ line to determine a greater range of wild-type responses to infections.

*Salmonella typhimurium*

Male *egr*$^{1}$ and *egr*$^{3}$ mutants had significantly lower survival to injection of *S. typhimurium* than the appropriate *OR-R* control (P<0.001), but had similar survival to the $w^{1118}$ and $w^{iso}$ lines (Figure 1A). Both male and female *OR-R* flies had significantly higher survival to *S. typhimurium* than other wild-type lines (P<0.001). Male *NimC1* mutants had low survival to *S. typhimurium* in comparison to both $w^{1118}$ and *OR-R* (P<0.001), but comparable survival to the $w^{iso}$ line. Although we did not perform CFUs as (Brandt et al., 2004; Mabery and Schneider, 2010) did, in our hands male *eiger* mutants do not have prolonged survival to *S. typhimurium*, and therefore do not have reproducibly increased tolerance to this bacterium. Interestingly, male and female flies had strong differences in survival to this infection. Female *egr*$^{1}$ and *NimC1* mutants had significantly higher survival to *S. typhimurium* (p<0.001) than the $w^{iso}$ and $w^{1118}$ wild types. However, this increased survival was comparable to the *OR-R* wild-type, and furthermore, survival of female *egr*$^{3}$ mutants was low and similar to $w^{iso}$ and $w^{1118}$ wild types (Figure 1B). Only *NimC1* females had consistently higher survival to *S. typhimurium* infection compared to all wild-types tested. None of the genotypes used had a significant reduction in survival in response to injection of LB medium alone (data not shown). Our methodology differs as (Brandt et al., 2004) injected flies in the abdomen, whereas we injected in the thorax. We found that abdominal injection caused excessive damage to internal tissues and high fly mortality regardless of genotype.

It is worth noting that *egr*$^{3}$ mutants were more susceptible to *S. typhimurium* infection than *egr*$^{1}$ for both males and females (Figure 1A, B), suggesting that a factor other than loss of *eiger* function is contributing to survival in these lines despite backcrossing to the same wild-type. This may be attributable to residual genetic background effects. Alternatively, the specific deletions generated by excision of the P-element in each line may have different effects on the gene. For example, one or both may not be true null mutations but rather produce a truncated protein that generates non-equivalent effects between the two lines. (Brandt et al., 2004) used only male *egr*$^{N}$ flies and compared to an *OR-R* wild type, while (Schneider et al., 2007) compared to the $w^{1118}$ line. In our hands, males of all three mutant lines (*egr*$^{1}$*, egr*$^{3}$*, NimC1*) had lower survival than the *OR-R* wild type and similar or lower survival compared to the $w^{1118}$ wild type, rather than prolonged survival. These results suggest that contrary to (Brandt et al., 2004; Mabery and Schneider, 2010; Schneider et al., 2007) loss of *eiger* or *NimC1* does not reproducibly increase tolerance to *S. typhimurium* infection.

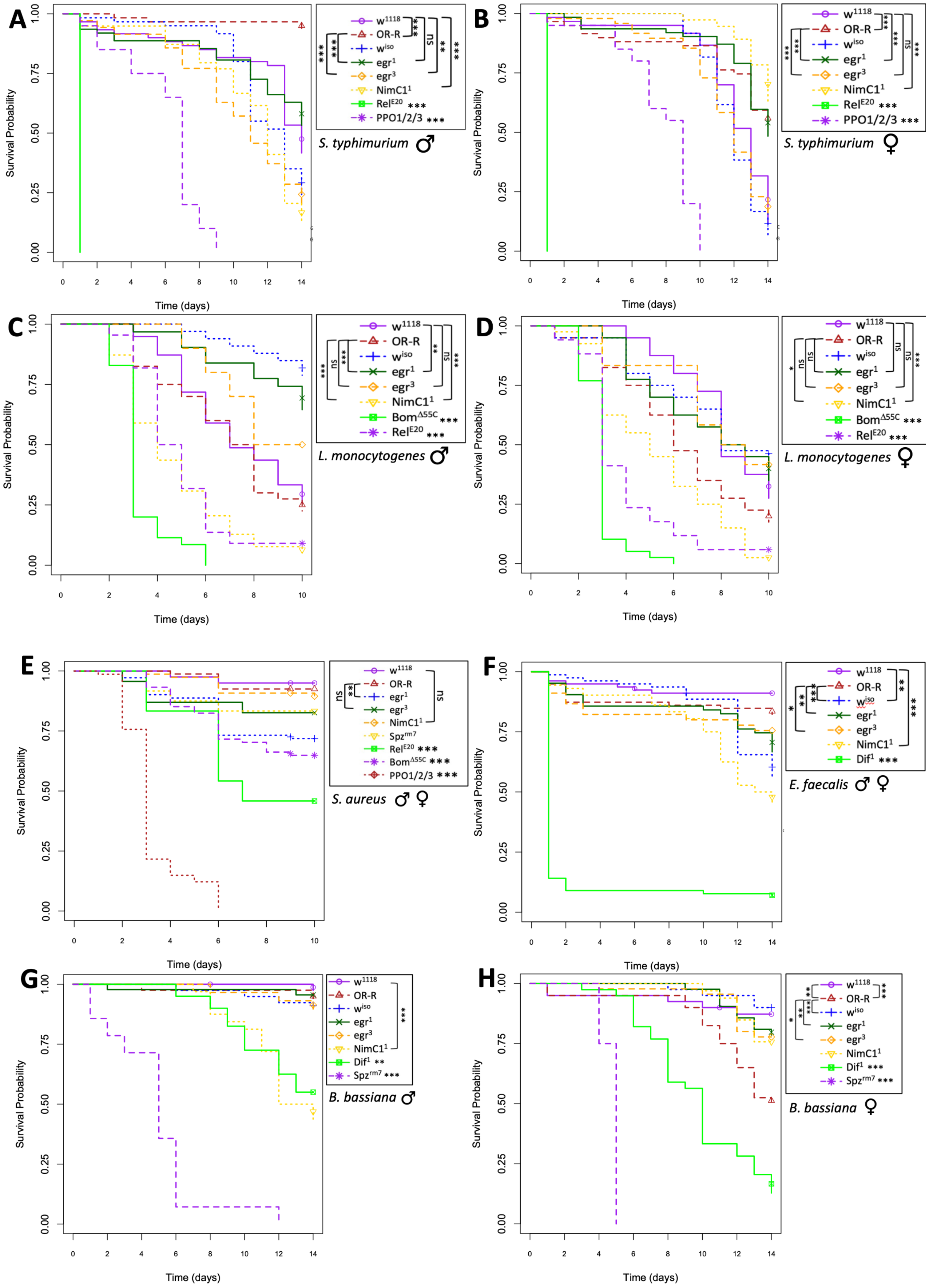

**Figure 1.** Survival of *eiger* and *NimC1* mutants in comparison to a range of wild type lines and susceptible controls to a range of infections: **A, B)** injection of $OD_{600}$=0.01 *S. typhimurium* at 29°C, n=3x20; **C, D)** septic infection with $OD_{600}$=0.5 *L. monocytogenes* at 25°C, n=2x20; **E)** septic

infection with $OD_{600}$=0.5 *S. aureus* at 25C, n=2x20 males and 2x20 females; **F)** septic infection with $OD_{600}$=0.5 *E. faecalis* at 29°C, n=2x20 males and 2x20 females; **G, H)** natural infection with 200mg *B. bassiana* spores at 29°C, n=2x20 males and 2x20 females. Male and female data was combined only where males and females did not have significantly different survival patterns to a given pathogen. Statistical differences calculated using a Cox Proportional Hazards test. Legend shows comparisons as indicated by brackets; where no bracket is given, genotype was significantly different from both $w^{1118}$ and *OR-R*. Significance code: ns = not significant, * = P<0.05, ** = P<0.01, ***P<0.001.

*Listeria monocytogenes*

(Schneider et al., 2007) found that $egr^N$ flies had similar survival to *L. monocytogenes* as $w^{1118}$ flies but higher bacterial loads, and concluded that the presence of *eiger* increases pathology to *L. monocytogenes.* Our results showed that male $egr^1$ mutants were resistant to this infection (P<0.001), while $egr^3$ males showed a similar but non-significant trend. However, both male (**Figure 1C**) and female (**Figure 1D**) *NimC1* mutants were strongly susceptible *L. monocytogenes* infection (P<0.01) compared to the appropriate $w^{1118}$ control. Females of both *eiger* lines were not significantly different from OR-R controls, but showed a similar trend of increased resistance (**Figure 1D**). Overall, our preliminary results suggest that mutation of *eiger* may confer resistance while mutation of *NimC1* increases susceptibility to *L. monocytogenes,* and that these effects were stronger in males. In the original $egr^N$ lines, these effects may have balanced each other, resulting in the net wild-type survival observed by (Schneider et al., 2007). Although we did not measure bacterial load, our results support the (Schneider et al., 2007) conclusion that loss of *eiger* improves tolerance to *L. monocytogenes.*

*Staphylococcus aureus*

In our hands, both *eiger* lines were slightly susceptible to *S. aureus* infection compared to the *OR-R* control, similar to $Dif^1$ or $spz^{rm7}$ flies (**Figure 1E**). This is a mild effect compared to the strong susceptibility found by (Schneider et al., 2007), which was more similar to survival of *PPO1/2/3* flies shown in **Figure 1E**. Stronger susceptibility as found by (Schneider et al., 2007) is not directly attributable to the secondary *NimC1* mutation, as in our hands *NimC1* flies had wild-type survival to *S. aureus*. It is possible that the *NimC1* and *eiger* mutations have a synergistic effect on survival to *S. aureus* when combined; alternatively, the additional susceptibility of the original $egr^N$ lines found by (Schneider et al., 2007) may be due to a separate genetic background effect. Similar to our results, (Geuking et al., 2009) found that the $egr^N$ lines had a mild effect on survival to *S. aureus* when compared with the parental line, suggesting that mild susceptibility to *S. aureus* is directly attributable to loss of *eiger*. An independent $eiger^{66}$ mutant with a full excision of the gene did not show this susceptibility ((Geuking et al., 2009) Figure 4C), but an appropriate background control was not included for this mutant (*w; Iso2A/Iso2A; Iso3A/Iso3A* background (Bellen et al., 2004; Wang et al., 2006)).

Overall, our results and those of (Geuking et al., 2009) suggest that mutation of *eiger* significantly and reproducibly causes mild susceptibility to *S. aureus,* while the stronger susceptibility found by (Schneider et al., 2007) was due to comparison to a robust wild-type. This is also consistent with results showing that melanization is reduced in *eiger* flies ((Bidla et al., 2007), Supplementary S27), and that melanization is an essential defense against *S. aureus* infection (Dudzic et al., 2019). In conclusion, our data support the (Schneider et al., 2007) conclusion that loss of *eiger* increases suceptibility to *S. aureus,* and further support the interpretation that this susceptibility is due to reduced melanization as a result of the *eiger* mutation.

*Enterococcus faecalis*

In our hands, *eiger* mutants had a slight but significant reduction in survival to *E. faecalis* compared to either $w^{1118}$ or *OR-R* wild types ($egr^1$ (P<0.01), $egr^3$ (P<0.05))) (**Figure 1F**). *NimC1* mutant flies were more strongly susceptible to *E. faecalis* (P<0.001). The susceptibility to *E. faecalis* attributed to *eiger* by (Schneider et al., 2007) was likely partially due to both mutations present in the original $egr^N$ lines. The data of (Geuking et al., 2009) support this conclusion, as the parental line mutated for *NimC1* but with intact *eiger* expression had reduced survival to *E. faecalis* compared to an independent *yw* wild-type. Both our results and (Geuking et al., 2009) also show a similar magnitude in reduction of survival by both the *eiger* and *NimC1* mutations to *E. faecalis*. Although (Geuking et al., 2009) show that the independent *egr*$^{66}$ mutant had high survival to *E. faecalis* similar to the *yw* wild type*,* an appropriate background control was not used. Their data suggest that the *egr*$^{66}$ mutant would have had lower survival than the range of wild types used in our survival, supporting the notion that loss of *eiger* consistently reduces survival to *E. faecalis* when compared to an appropriate background control. Our data support the (Schneider et al., 2007) conclusion that loss of *eiger* increases susceptibility to *E. faecalis,* but further find that this susceptibility was exaggerated by the secondary *NimC1* mutation.

*Beauvaria bassiana*

Contrary to (Schneider et al., 2007) our results suggest that loss of *eiger* does not increase susceptibility to *B. bassiana* natural infection (**Figure 1G)**. Male and female *eiger* mutants had similar or higher survival than the appropriate *OR-R* wild type. In contrast, male *NimC1* mutants were strongly susceptible to *B. bassiana* natural infection similar to Toll-deficient lines compared to all wild types (P<0.001) (**Figure 1G**). This suggests that susceptibility to *B. bassiana* in male flies attributed to the *eiger* mutation by (Schneider et al., 2007) was due to the secondary *NimC1* mutation. Interestingly, loss of *NimC1* has no effect on survival of females to the same treatment (**Figure 1H**). This is consistent with results showing that survival to *B. bassiana* natural infection is strongly sexually dimorphic (Belmonte et al., 2020). Our data contradict the (Schneider et al., 2007) conclusion that loss of *eiger* increases susceptibility to *B. bassiana* natural infection, and instead show that loss of *NimC1* confers strong susceptibility to this treatment on male flies only.

#### Conclusion

Our experiments suggest that patterns of susceptibility attributed to the *eiger* mutation by (Brandt et al., 2004; Mabery and Schneider, 2010; Schneider et al., 2007) are partially attributable to *eiger* itself, and partially due to genetic background or secondary mutation of *NimC1*. Contrary to (Schneider et al., 2007), we were unable to reproduce increased tolerance of male *eiger* or *NimC1* flies to *S. typhimurium* infection, and infer that this conclusion was reached due to comparison to a wild type with a different genetic background. Similar to (Schneider et al., 2007), we find that the *eiger* mutation increases survival to *L. monocytogenes,* although the *NimC1* mutation increases susceptibility. Similar to (Schneider et al., 2007), we find that loss of *eiger* increases susceptibility to *S. aureus,* but this is a much milder effect than originally stated. Finally, we find that susceptibility of male *egr*$^{N}$ flies to *B. bassiana* natural infection is instead attributable to the secondary mutation of *NimC1* in these lines.

### Supplementary S26: Role of *eiger* in melanization

**Abstract**

Previous publications indicated that melanization defects occurred in flies upon depletion of *eiger* by RNAi and in *eiger* mutants. However, these widely-used *eiger* mutants were later found to bear a secondary mutation of *NimC1,* a gene with independent roles in immunity. Our results using *eiger* mutants backcrossed to remove the *NimC1* mutation support a role for *eiger* in melanization in of both larvae and adult flies. However, in contrast to (Mabery and Schneider 2010), our results based on fat-body specific knock-down do not support a role of *eiger* in this tissue for melanization.

**Introduction**

A series of results implicating the TNF related gene *eiger* in several aspects of *Drosophila* immune function have recently been called into question due to the discovery that widely used *eiger* mutants bore a secondary mutation of the phagocytic receptor *NimC1* (Kodra et al., 2020), which has independent roles in immunity (Melcarne et al., 2019). Because of this, and because most literature on the immune functions of *eiger* has not been independently verified, we set out to determine whether phenotypes previously assigned to *eiger* are reproducibly attributable to loss of either *eiger* or *NimC1*.

(Bidla et al., 2007) found that *eiger* was required in hemocytes but not the fat body for JNK-mediated crystal cell rupture and melanization upon infection. Independently, (Mabery and Schneider, 2010) found that 'disseminated melanization' was reduced in *eiger* RNAi flies compared to controls, more strongly with a fat body driver than a hemocyte driver. 'Disseminated melanization' is dependent on Sp7, see (Ayres and Schneider, 2008), and was assayed by injecting 50 nL of *Salmonella typhimurium* into the ventrolateral abdomen of male flies, incubating flies 6 days at 29°C, and then imaging and counting melanized spots visible through the cuticle of the abdomen. Percentages of flies displaying melanized spots were compared between genotypes. These authors found that melanization at the actual wound site was not impaired. (Mabery and Schneider, 2010) hypothesized that starvation reduces phenoloxidase (PO) activity, as both *eiger* and *Gr28b* 'anorexic' flies had reduced feeding and capacity to melanize. Although independent results have found a role for *eiger* in insulin signaling and appetite stimulation (see (Agrawal et al., 2016)), our results show that reduced melanization in the *Gr28b* line is due to genetic background (Supplementary S28, Figure 4), and neither *Gr28b* (Supplementary S28, Figure 1) nor *eiger* (See complementary data reported in the ReproSci website) flies have reduced feeding compared to appropriate controls. Furthermore, (McCormack et al., 2016) instead found that starvation increases melanization and PO activity. Thus, if *eiger* has a role in melanization, it is likely independent to possible impacts on feeding.

To test for a role of *eiger* in melanization, we employed *eiger* mutants backcrossed to the *OR-R* wild-type line, and fat body specific RNAi.

**Methods**

*Fly lines and bacterial strains*

$egr^1$ ($egr^1$/*CyO-GFP*) and $egr^3$ ($egr^3$/*CyO-YFP*) mutants cleaned by backcrossing to *OR-R* were obtained from (Kodra et al., 2020). $w^{1118}$ (Bloomington #3605) $w^{iso}$ *Drosdel* (Ryder et al., 2004) and *OR-R* flies were used to provide a range of wild-type comparisons. *PPO1/2/3* ($PPO1^D$, $PPO2^D$, $PPO3^1$) triple mutants or single and double mutant subsets of these were used as melanization-deficient strains (Binggeli et al., 2014; Dudzic et al., 2019). RNAi lines (*UAS-eiger-RNAi* II #108814, *UAS-eiger-RNAi* III #45253) and an *eiger-GFP* construct (#318615, 2XTY1-SGFP-V5-preTEV-BLRP-3XFLAG, 3rd chromosome (Mirzoyan et al., 2023)) were obtained from VDRC. Drivers used are $Lpp\text{-}Gal4^{ts}$/*TM3* (Brankatschk et al., 2018), $w^{1118}$*; c564-Gal4* (Bloomington #6982), and *yw; Act5C-Gal4/CyO-GFP* (Bloomington #25374). Flies were reared at 25°C on cornmeal fly medium (6% cornmeal, 6% yeast, 0.6% agar, 0.1% fruit juice, 10.6 g/L moldex and 4.9 mL/L propionic acid), and flipped to new medium every 2-3 days. RNAi crosses were shifted to 29°C following egg laying to promote driver activity. The strains *S. typhimurium* SL1344 and *M. luteus* JL (Jean Lambert collection, Strasbourg) were used in this study

*Assessment of Melanization*

5 to 7-day old male and female flies reared on cornmeal fly medium were anaesthetized with $CO_2$ and pricked on the left side of the thorax anterior and ventral to the wing joint with a 0.2 mm tungsten needle dipped in concentrated *Micrococcus luteus* ($OD_{600}$=200) from overnight cultures (29°C, shaking, LB) or *S. typhimurium* SL1344 cultured as in (Schneider et al., 2007), and diluted to $OD_{600}$ = 200 in LB medium prior to septic infection. Flies were incubated at rearing conditions for 16h or 24h after pricking and melanization of the injury site was scored with dark brown or black spot indicating “strong” melanization; light brown or yellow spot indicating “weak” melanization; and no apparent spot indicating no melanization. Data for male and female flies were combined as they yielded similar results. To assess melanization in larvae, third instar wandering larvae were pricked at the posterior end with a 0.1 mm needle dipped in concentrated *M. luteus* and placed on grape juice agar in a humidification chamber at 25°C for 2h.

#### Results and Discussion

Initially, we attempted to assay ‘disseminated melanization’ in clean *egr*$^1$ and *egr*$^3$ mutants as described by (Mabery and Schneider, 2010). However, we found that the abdominal injection procedure caused excessive damage to internal tissues and high fly mortality regardless of genotype. We therefore used thoracic injection. We found that no flies from either the *eiger* mutant or wild-type lines produced melanized spots in the body cavity following 6d incubation at 29°C. Melanization in the body cavity also did not occur at later time points, even when flies began to die from bacterial infection. We did however find that the few flies that survived abdominal injection produced melanized spots as described by (Mabery and Schneider, 2010) regardless of genotype. That thoracic injection did not produce this effect suggests that this melanization is due to tissue damage from the abdominal injection procedure and subsequent melanization of that damaged tissue within the body cavity, rather than a response to the bacterial infection.

It is well established that disruption of basement membranes and tissue damage results in melanization (Kim and Choe, 2014; Mortimer et al., 2021; Scherfer et al., 2006), and that free-floating pieces of disrupted tissue in the body cavity become melanized (and in larvae, encapsulated by lamellocytes) (Minakhina and Steward, 2006; Mortimer et al., 2021; Paddibhatla et al., 2010; Rizki and Rizki, 1979). It is logical that injection of certain bacteria that strongly stimulate the melanization response could enhance melanization of disrupted tissue. Injection with some bacteria could also cause melanization by colonizing tissue and causing necrosis late in infection. However, in our hands, *S. typhimurium* injection did not promote melanotic spot formation in *Drosophila* in the absence of high levels of tissue damage inflicted by abdominal injection. We therefore sought alternate ways to assess melanization in *eiger* mutants compared to wild-type flies.

We assessed melanization at the wound site in larvae and flies in response to septic injury with *M. luteus. Eiger* mutants were compared both to wild-type animals and to *eiger* heterozygous siblings (bearing the *CyO* balancer). Homozygous *egr*$^1$ and *egr*$^3$ flies and larvae had reduced melanization compared to wild-type animals (Figure 1). Interestingly, larvae heterozygous for the *egr* mutations also had reduced melanization compared to wild-type larvae, although the effect was less marked compared to homozygous *eiger* mutants (Figure 1). This may indicate that *eiger* is haploinsufficient for wild-type melanization. However, this should be further tested by crossing *eiger* flies to the *OR-R* wild type, as balancer chromosomes can have odd phenotypic effects (see e.g. (Ekengren et al., 2001)). In adult flies, cuticle melanization at the wound site was not strongly impaired in either the *egr*$^1$ or *egr*$^3$ lines, although the degree of blackening was qualitatively decreased (Figure 1). These results are consistent with (Mabery and Schneider, 2010) who found that melanization at the wound site in adult *eiger* flies was not strikingly different from wild-type flies, but hemolymph melanization was reduced.

Melanization is known to be important in survival to certain infections, and particularly to the Gram-positive bacterium *Staphylococcus aureus* (Binggeli et al., 2014; Dudzic et al., 2019). Furthermore, the hemolymph blackening apparent in larvae, which is strongly reduced in *egr*$^1$ and *egr*$^3$ animals, is more directly associated with combatting systemic infections than the cuticle blackening observed in adults (Dudzic et al., 2019). Consistent with qualitative observations of decreased melanization in *egr*$^1$ and *egr*$^3$ larvae and adults, we found that survival *S. aureus* was reduced in *eiger* mutants (Supplementary S25). The susceptibility of *eiger* mutants was milder than that conferred by loss of all three hemolymph phenoloxidases, but similar to flies lacking *PPO2* alone (Binggeli et al., 2014). This is consistent with results showing that *eiger* is required for rupture of crystal cells, which contain high concentrations of PPO2, upon infection (Bidla et al., 2007).

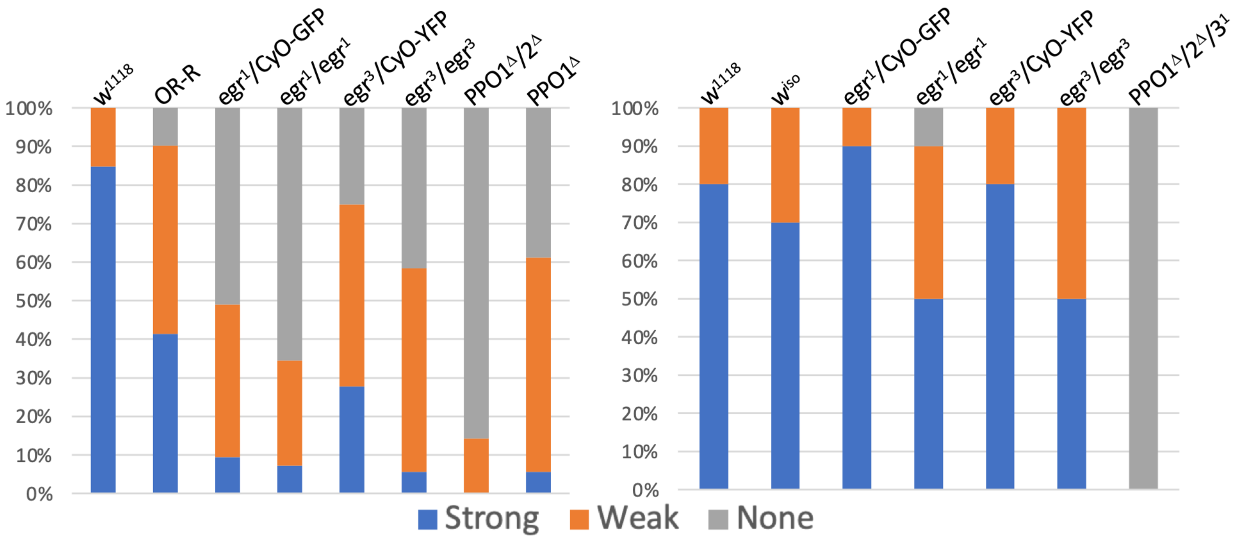

**Figure 1.** Melanization in larvae and adults in response to septic injury with *M. luteus.* **Left,** Melanization in larvae (n=2x20) at 60 minutes post-injury. **Right,** Melanization in adult flies at 16h post-injury (n=1x20 males and 1x20 females). Strong = dark brown or black spot; Weak = light brown or yellow spot; None = no apparent melanization. n=2 independent experiments that yielded similar results, mean of the two experiments is displayed.

In our hands, RNAi against *eiger* in the fat body using two independent drivers and two RNAi constructs showed no effect on melanization (Figure 2), indicating that *eiger* is not required in the fat body for melanization at the adult stage. This is consistent with results showing that *eiger* is not strongly expressed in the fat body (Supplementary S27 - Figure 1).

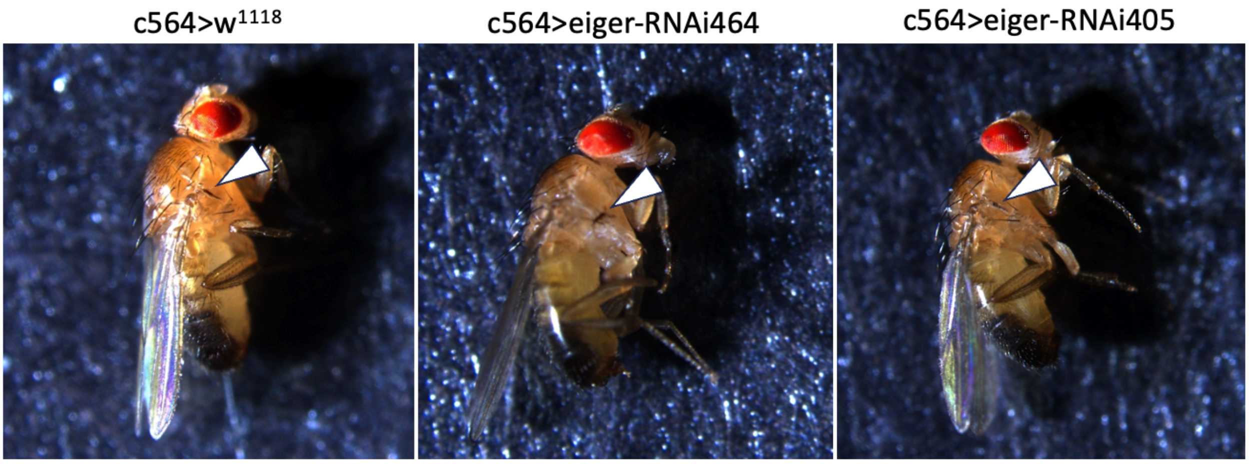

| | Lpp>W | Lpp>464 | Lpp>405 | C564>W | C564>464 | C564>405 |
|---|---|---|---|---|---|---|
| Flies with spots/total flies | 56/60 | 56/60 | 57/60 | 58/60 | 60/60 | 58/60 |

**Figure 2.** Melanization in *eiger*-depleted flies 24h post-infection with *S. typhimurium* ($OD_{600}$ = 200). Arrowheads indicate melanized spot produced by pricking. Two RNAi constructs (464, 405) against *eiger* in the fat body (*c564-Gal4, Lpp-Gal4*) showed no difference in melanization from controls.

#### Conclusion

Our results support the claim that *eiger* is required for wild-type melanization in adult flies and larvae as found by (Bidla et al., 2007; Mabery and Schneider, 2010). Animals bearing the *egr*$^{1}$ and *egr*$^{3}$ mutations had impaired blackening and increased susceptibility to *S. aureus* similar to PPO mutants with reduced hemolymph phenoloxidase activity. However, contrary to (Mabery and Schneider, 2010), we find that *eiger* is likely not required in the fat body for melanization.

### Supplementary S27: Role of *eiger* in the humoral response

**Abstract**

Previous publications indicated that *eiger* may be upregulated in the fat body upon infection, and that expression of the Imd target *Diptericin* was elevated in *eiger* mutants. However, these widely-used *eiger* mutants were later found to bear a secondary mutation of *NimC1,* a gene with independent roles in immunity. Contrary to (Mabery and Schneider, 2010; Schneider et al., 2007), we find that Eiger is not markedly upregulated in the fat body following infection using an *Eiger-GFP* construct, and that depletion of *eiger* either ubiquitously or in the fat body does not significantly affect *Diptericin* expression upon infection.

**Introduction**

A series of results implicating the TNF related gene *eiger* in several aspects of *Drosophila* immune function have recently been called into question due to the discovery that widely used *eiger* mutants bore a secondary mutation of the phagocytic receptor *NimC1* (Kodra et al., 2020), which has independent roles in immunity, particularly in phagocytosis (Melcarne et al., 2019).

(Brandt et al., 2004) state that *eiger* is not upregulated in response to *Salmonella typhimurium* infection, whereas a follow up paper by the same group (Schneider et al., 2007) states that this gene is upregulated in the fat body upon infection. (Mabery and Schneider, 2010) found that RNAi against *eiger* in the fat body (*Isp-Gal4*), but not the hemocytes (*hml-Gal4*) suppressed induction of *eiger* as shown by RT-qPCR upon infection. Despite these findings, upregulation of *eiger* is not detectable on a whole-body level upon infection by *Beauveria bassiana* or mixed *Micrococcus luteus* and *Escherichia coli* (De Gregorio et al., 2001). FlyAtlas indicates enrichment of *eiger* in the fat body of unchallenged larvae but not adults.

Additionally, (Schneider et al., 2007) found that *Drs* induction in *eiger* mutants was normal, although some pathogens (*Burkholderia cepacia, S. typhimiurum*) and LB injection resulted in higher *Dpt* induction in *eiger* mutants than wild type flies. (Mabery and Schneider, 2010) found that RNAi of *eiger* in the fat body but not hemocytes resulted in a 2- to 3-fold increase in *Dpt* expression compared to parents following *S. typhimurium* infection, but unchanged *Drs* expression. Expression was compared to a $w^{1118}$ wild-type strain; this is a different background from the one the *eiger* mutants was generated in, and expression of antimicrobial peptides (AMPs) is known to differ strongly between backgrounds.

Because of these discrepancies, and because most literature on the immune functions of *eiger* has not been independently verified, we tested several claims related to the roles of *eiger* in immunity. Data reported in the ReproSci website show that contrary to (Mabery and Schneider, 2010), *eiger* mutant flies do not have reduced feeding compared to appropriate controls. In addition, depletion of *eiger* in the fat body does not suppress melanization, although a null mutation of *eiger* does (Supplementary S26). Only some of the survival (resistance/tolerance) effects previously attributed to loss of *eiger* are replicable, whereas other survival effects appear to be due to the contaminating mutation in *NimC1* (Supplementary S25).

Here, we tested whether Eiger is induced in the fat body upon infection using an *Eiger-GFP* construct, and whether depletion of *eiger* affects infection-induced expression of the AMP *Diptericin.*

**Methods**

*Fly lines*

Flies were reared at 25°C on cornmeal fly medium (6% cornmeal, 6% yeast, 0.6% agar, 0.1% fruit juice, 10.6 g/L moldex and 4.9 mL/L propionic acid), and flipped to new medium every 2-3 days. $w^{1118}$ flies were used as a wild-type control (Bloomington #3605). RNAi lines (*UAS-eiger-RNAi* II VDRC #108814, *UAS-eiger-RNAi* III VDRC #45253) and an *eiger-GFP* FlyFos construct (#318615, 2XTY1-SGFP-V5-preTEV-BLRP-3XFLAG, 3rd chromosome (Mirzoyan et al., 2023)) were obtained from VDRC. Drivers used are *Lpp-Gal4*$^{ts}$*/TM3* (Brankatschk et al., 2018), $w^{1118}$*; c564-Gal4* (Bloomington #6982), and *yw; Act5C-Gal4/CyO-GFP* (Bloomington #25374).

*Bacterial culture and infection*

*S. typhimurium* SL1344 was cultured as in (Schneider et al., 2007). *Pectobacterium carotovorum Ecc15* was cultured overnight at 29°C shaking in LB. Cultures were concentrated by centrifugation and diluted to $OD_{600}$=200 in PBS prior to septic injury. 4 to 6-day old male flies were anaesthetized with $CO_2$ and pricked on the side of the thorax anterior and ventral to the wing joint with a 0.2 mm tungsten needle dipped in a concentrated bacterial pellet. RNAi crosses were incubated at 29°C to promote driver activity. ~20 flies were used per genotype per experiment, and three independent experiments were performed.

*RT-qPCR experiments*

Male *eiger* RNAi flies were collected at 6h and 24h collected following septic infection with *S. typhimurium*. For each treatment, 6-8 flies were frozen at -80°C at 6h, 24h, and 48h post infection. Three independent experiments were performed for each treatment. Total RNA was extracted from samples using TRIzol reagent (Invitrogen), and re-suspended in Milli-Q water. DNase treatment was performed using TURBO DNA-free kit (Invitrogen). Reverse transcription was performed using 0.5 mg total RNA in 10 ul reactions using Prime-Script RT (Takara) with random hexamer and oligo dT primers. Quantitative PCR was performed on a LightCycler 480 (Roche) in 96-well plates using PowerUp SYBR Green Master Mix (Applied Biosystems). *Diptericin* expression was normalized to expression of the ribosomal protein *rp49* (*rpl32*). For graphical display, *Diptericin* expression levels were normalized to the unchallenged (UC) condition in wild type $w^{1118}$ flies.

- *Rp49* Forward: 5' – GCC GCT TCA AGG GAC AGT ATC TG – 3'
- *Rp49* Reverse: 5' – AAA CGC GGT TCT GCA TGA G – 3'
- *Diptericin A* Forward: 5' – GCT GCG CAA TCG CTT CTA CT – 3'
- *Diptericin A* Reverse: 5' – TGG TGG AGT GGG CTT CAT G – 3'
-

#### Results

*Eiger is not upregulated in the fat body upon infection*

Flies expressing an *Eiger-GFP* construct showed no particular expression of the GFP construct in the fat body either in unchallenged conditions or upon infection with the Gram-negative bacterium *Ecc15* (Figure 1). Our data do not exclude that Eiger may be upregulated in other tissues upon infection, or that the *Eiger-GFP* construct does not fully capture endogenous Eiger expression.

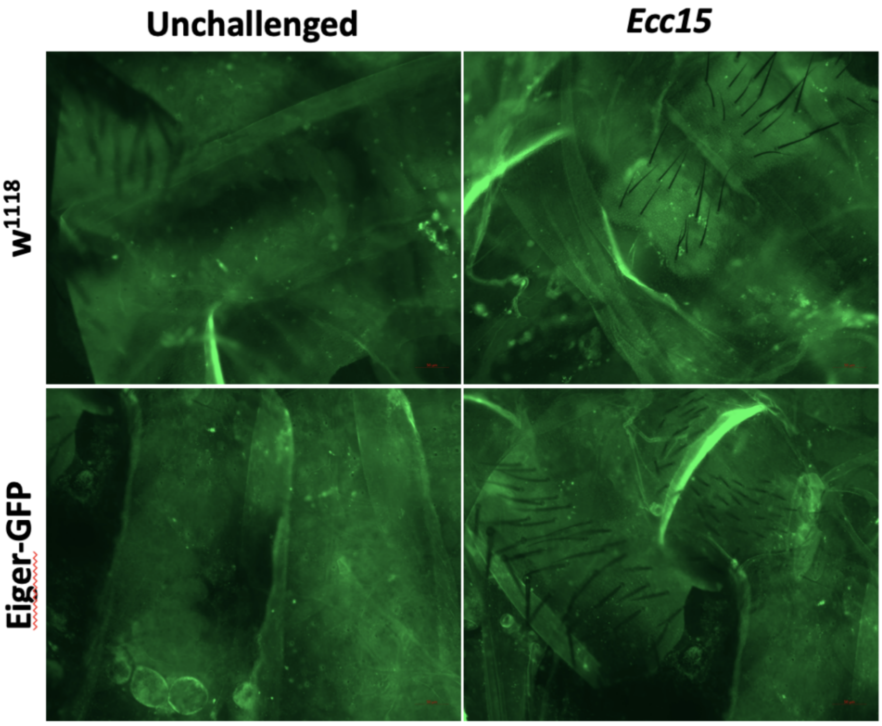

**Figure 1.** Flies expressing an Eiger-GFP construct showed no marked expression of the construct in the fat body either in unchallenged conditions or 24h following infection with *Ecc15.* Flies were torn open with forceps in PBS to expose fat body tissues prior to imaging.

*Loss of eiger in the fat body does not affect the humoral immune response*

In our hands, *Dpt* expression was overall similar to wild type in *eiger-RNAi* flies 6h and 24h following *S. typhimurium* infection. Of the driver and RNAi combinations tested, only *c564>eiger-RNAi405* flies at the 24h time point were significantly different from the appropriate control (*c564>*$w^{1118}$) (Figure 2). These flies showed a reduction in *Dpt* expression, rather than an increase as found by (Mabery and Schneider, 2010; Schneider et al., 2007). Furthermore, a second *eiger-RNAi* construct (*eiger-RNAi464*) did not show a similar trend, and an independent fat body driver (*Lpp-Gal4*) and a ubiquitous driver (*Act5C-Gal4*) also showed no difference in *Dpt* expression upon infection.

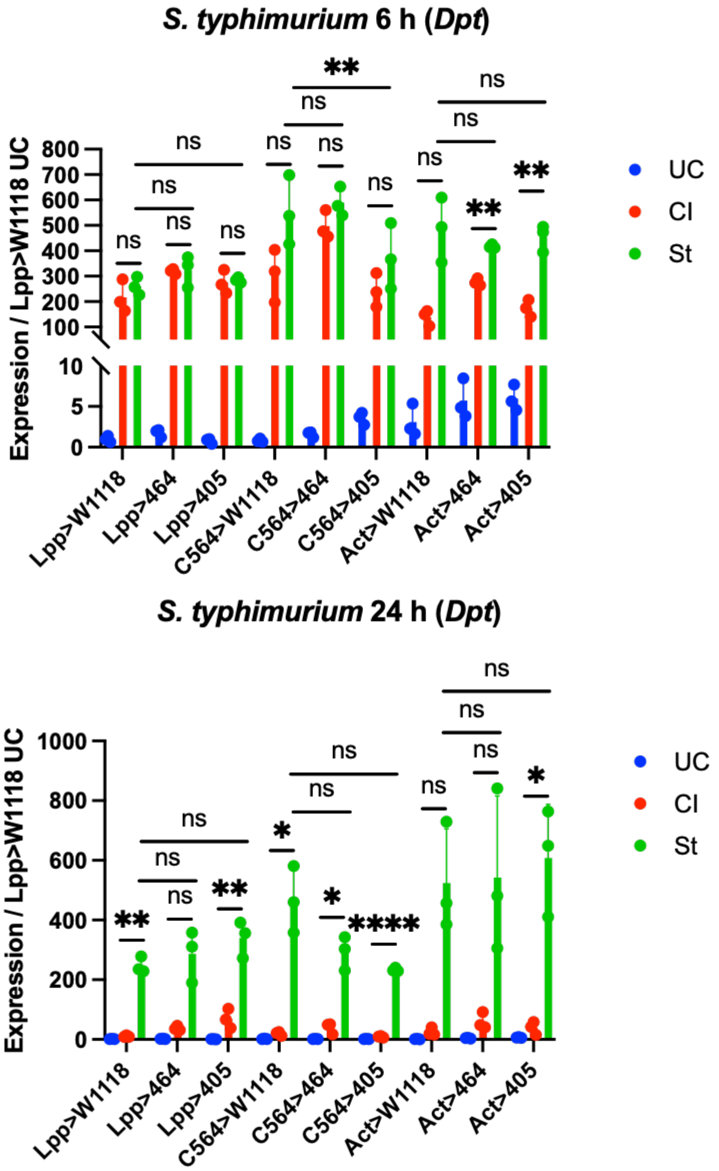

**Figure 2.** *Dpt* expression in controls and *eiger* RNAi flies at 6h and 24h following septic infection with *S. typhimurium* (St). Expression in unchallenged flies (UC) and flies subjected to clean injury (CI) are also shown. Only *c564>eiger-RNAi405* flies show a significant difference in *Diptericin* expression from the appropriate control. n=3 independent experiments, data were analyzed using Two-Way ANOVA followed by Tukey's mutiple comparison tests. 405: *UAS-eiger-RNAi* II VDRC #108814; 464: *UAS-eiger-RNAi* III VDRC #45253. Error bars represent the standard deviation.

#### Discussion and Conclusion

Contrary to (Mabery and Schneider, 2010; Schneider et al., 2007), our results did not find that *eiger* is not strongly expressed in the fat body and is not upregulated upon infection. We cannot exclude that the expression of the *eiger-GFP* reporter is weak and does not fully capture endogenous expression, but our data are consistent with microarray studies that did not reveal any clear induction of *eiger* on the whole-body level (De Gregorio et al., 2001, Troha et al., 2018). Furthermore, we found no consistent effect of *eiger* depletion on expression of the AMP *Diptericin* upon infection, and conclude that loss of *eiger* in the fat body or ubiquitously does not strongly affect the humoral immune response.

### Supplementary S28: Gr28b in feeding and immunity

**Abstract**

(Ayres et al., 2008) identified a *P*-element insertion in the gustatory receptor *Gr28b* in a screen for mutations that increase susceptibility to *L. monocytogenes.* The same authors (Ayres and Schneider, 2009) further explored effects of the *Gr28b* mutation and found that these flies were constitutively anorexic, which affected survival to a number of bacteria. Subsequently published evidence and experiments performed for the ReproSci suggest that phenotypes of the *Gr28b* mutant used by (Ayres and Schneider, 2009) are largely attributable to reduced melanization linked to the genetic background, as an independent *Gr28b* mutant does not replicate these phenotypes. Furthermore, we find that feeding rates are highly variable among wild type lines, and that *Gr28b* mutant flies do not have reduced feeding when considered in a broader context.

**Introduction**

(Ayres et al., 2008) identified a P-element insertion in the gustatory receptor *Gr28b* in a screen for mutations that increase susceptibility to *Listeria monocytogenes.* The same authors further characterized this *Gr28b*$^{c01884}$ line and found that these flies were constitutively anorexic, which affected survival to a number of bacteria (Ayres and Schneider, 2009). These flies had increased tolerance and survival to *Salmonella typhimurium,* reduced survival and resistance to *L. monocytogenes,* and wild-type survival to *Enterococcus faecalis* infection. Authors also found that these flies had reduced melanization and altered antimicrobial peptide (AMP) gene expression compared to wild-type flies.

In contrast to this, (Sang et al., 2019) found that a line with an independent insertion in *Gr28b* (*Gr28b*$^{MB03888}$) had no difference in intake of standard food compared to wild type, suggesting that the reduced feeding and immune phenotype observed by (Ayres et al., 2008; Ayres and Schneider, 2009) in *Gr28b*$^{c01884}$ mutants may have been due to a secondary mutation or genetic background effects. (Sang et al., 2019) further found that *Gr28b* was required for flies to detect and avoid saponins in food, and that *Gr28b* mutants in fact had increased feeding on saponin-laced foods compared to wild-type flies.

Other authors have found that starvation or reduced insulin signaling typically enhances survival to bacterial infections (see e.g. (Lee et al., 2017) *Pseudomonas aeruginosa,* (Libert et al., 2008) *Pseudomonas aeruginosa* and *E. faecalis*, (McCormack et al., 2016) *Photorhabdus luminescens* and *Escherichia coli*)). However, increased nutrient intake appears to be protective in response to other infections, such as *Burkholderia cepacia* (Allen et al., 2016). (McCormack et al., 2016) found that increased survival to pathogenic infections upon reduced insulin signaling was due to increased melanization and phenoloxidase (PO) activity, although these flies also had higher AMP expression and phagocytic activity. In contrast (Libert et al., 2008) found that AMP expression was not increased by starvation. Reduced survival of anorexic flies in response to *L. monocytogenes* infection may be consistent with results showing that this infection is metabolically demanding and causes rapid depletion of energy stores, resulting in death (Chambers et al., 2012).

In order to resolve these conflicting studies, we performed experiments with both alleles of *Gr28b* in tandem, and investigated their feeding, melanization, and survival to a variety of pathogens.

**Methods**

*Fly lines and husbandry:*

Two alleles of *Gr28b* were ordered from Bloomington: PBac{PB}Gr28b[c01884] (BDSC #10743) is the line originally used by (Ayres et al., 2008; Ayres and Schneider, 2009), while w[1118]; Mi{GFP[E.3xP3]=ET1}Gr28b[MB03888] (BDSC #24190) was used by (Sang et al., 2019). For simplicity, we refer to these lines as *Gr28b*$^{P}$ (as in **P**Bac) and *Gr28b*$^{M}$ (as in **M**inos) respectively. We employed several wild-type lines including $w^{iso}$ *Drosdel* (Ryder et al., 2004), $w^{1118}$ (Bloomington #3605) and *OR-R* lines to provide a range of comparisons. *PPO1*$^{D}$ single mutants, *PPO1*$^{D}$*, PPO2*$^{D}$ double mutants, and *PPO1/2/3 (PPO1*$^{D}$*, PPO2*$^{D}$*, PPO3*$^{1}$) triple mutants (Dudzic et al., 2015) were used as melanization-deficient strains. *Relish*$^{E20}$ (*Rel*$^{E20}$) and *Dif*$^{1}$ (BDSC #36559) lines were used as Imd and Toll-deficient comparisons, respectively. Flies were reared at 25°C on cornmeal fly medium (6% cornmeal, 6% yeast, 0.6% agar, 0.1% fruit juice, 10.6 g/L moldex and 4.9 mL/L propionic acid), and flipped to new medium every 2-3 days.

*RNA extraction and qPCR*

Total RNA was extracted from pooled samples of 7-10 flies (5-7 days old) each using TRIzol reagent (Invitrogen), and re-suspended in MilliQ water. DNase treatment was performed using TURBO DNA-free kit (Invitrogen). Reverse transcription was performed using 0.5 mg total RNA in 10 µl reactions using Prime-Script RT (Takara) with random hexamer and oligo dT primers. Quantitative PCR was performed on a LightCycler 480 (Roche) in 96-well plates using PowerUp SYBR Green Master Mix (Applied Biosystems). AMP expression was normalized to expression of the ribosomal protein *rp49* (*RpL32*).

*Survival assays*

5 to 7-day old male and female flies were anaesthetized with $CO_2$ and pricked on the thorax anterior and ventral to the wing joint with a 0.2 mm tungsten needle dipped in a concentrated overnight (shaking, 29°C, BHI) culture of *L. monocytogenes* (CIP 107776, EGDe) or *E. faecalis* (ATCC 29212) diluted to the indicated OD in PBS, or injected with *S. typhimurium SL1344* (cultured as in Schneider et al., 2007) diluted to the indicated OD in LB medium. Flies were incubated at temperatures indicated in figures and scored daily for survival; flies were flipped to new medium during the survival every 2-3 days. At least 20 flies per genotype were scored in 2-3 independent experiments. Data were analyzed in R using a Cox Proportional Hazard test.

*FlyPad feeding assays*

The FlyPad apparatus (Itskov et al., 2014) was set up according to the manufacturer instructions and loaded with 4 µL cornmeal fly medium (6% cornmeal, 6% yeast, 0.6% agar, 0.1% fruit juice, 10.6 g/L moldex and 4.9 mL/L propionic acid) on each electrode. 5-7 day-old flies were sorted at least 24h prior to the feeding assays, as exposure to $CO_2$ can strongly alter metabolism in *Drosophila* (Colinet and Renault, 2012). All feeding experiments were performed in the afternoon (2-5pm) without prior starvation similar to (Ayres and Schneider, 2009). Flies were quickly distributed into the arenas using a pooter (1 fly per arena) and feeding behavior was measured for 30 minutes. Male and female flies from the same cohorts were separated and measured in consecutive runs. Three experiments were performed, for a total of at least 50 flies per genotype per sex.

*Blue dye feeding assay*

Standard fly food was prepared with the addition of 1% FD&C Blue No. 1 dye. Food was dispensed into standard fly vials and allowed to cool completely. To avoid flies becoming externally covered in dye by walking in the food, a small hole was punched in the centre of filter papers and these were used to completely cover the surface of the food except for a ~2mm circle where flies could feed. 5-7 day-old male and female flies were sorted with $CO_2$ anaesthesia at least 24h prior to the feeding assay. Flies were flipped onto the food containing blue dye and allowed to feed for 30 minutes, then flies were rapidly anaesthetized and scored for presence/absence of dye in the gut. 20-40 flies per sex per genotype were scored. Ten flies per sex per genotype that had blue dye in the gut were selected, rinsed briefly in PBS to remove external dye, and homogenized in 200 µL PBS. The homogenate was centrifuged and 100 µL of supernatant was transferred to a transparent 96 well plate. Absorbance of the homogenates were read on a TECAN spectrophotometer at 629nm.

*Melanization in adult flies and larvae*

5 to 7-day old male and female flies were anaesthetized with $CO_2$ and pricked on the left side of the thorax anterior and ventral to the wing joint with a 0.2 mm tungsten needle. The needle was sterilized with ethanol prior to clean injury or dipped in bacterial pellets ($OD_{600}$=200) concentrated from overnight cultures of *Micrococcus luteus* (overnight, shaking, 29°C, LB) prior to septic injury. Flies were incubated at rearing conditions for 16h after pricking and melanization of

the injury site was scored. At least 20 flies per genotype were scored, with dark brown or black spot indicating “strong” melanization; light brown or yellow spot indicating “weak” melanization; and no apparent spot indicating no melanization.

Larval melanization was scored in two independent experiments with 10-20 larvae per genotype per experiment. Larvae were pricked at the posterior end with a 0.1 mm needle dipped in a bacterial pellet ($OD_{600}$=200) concentrated from overnight cultures of *M. luteus*. Larvae were then placed on grape juice agar in a humidification chamber at 25°C for 30 minutes and melanization was scored visually. Larvae were scored as ‘Strong’ if they produced a black spot similar to that typically produced in wild-type larvae, ‘Weak’ if only a light yellow or brown spot was visible, and ‘None’ if no melanization was apparent.

#### Results and Discussion

Gr28b *mutant flies are not anorexic*

(Ayres and Schneider, 2009) found that $Gr28b^P$ flies had decreased feeding rates and ingestion volumes compared to $w^{1118}$ wild-type flies. However, (Sang et al., 2019) found that an independent insertion in the gene, $Gr28b^M$, showed no difference in intake of standard food in these flies compared to the $w^{1118}$ wild type. (Sang et al., 2019) further found that *Gr28b* was required for flies to detect and avoid saponins in food, and that *Gr28b* mutants had increased feeding on saponin-laced foods compared to wild-type flies. To clarify this contradiction, we tested feeding of both *Gr28b* mutants in comparison to two wild type lines (*OR-R* and $w^{1118}$) on cornmeal fly medium using the FlyPad apparatus (Itskov et al., 2014).

Three experiments using the FlyPad to measure feeding do not support the notion put forward by (Ayres and Schneider, 2009) that $Gr28b^P$ male flies have lower feeding compared to the $w^{1118}$ wild type. Both $w^{1118}$ and $Gr28b^P$ genotypes had similar feeding in terms of rate and cumulative feeding events (Figure 1A). Feeding was comparatively extremely low in the $Gr28b^M$ line and in the *OR-R* wild type. Furthermore, females from the $Gr28b^P$ line had very high feeding compared to both wild types (Figure 1B). Variance is high for the $Gr28b^P$ line as two out of three experiments showed excessively high feeding compared to wild types, while in one experiment feeding of $Gr28b^P$ flies was similar to the wild type lines. We conclude that it is not appropriate to characterize the $Gr28b^P$ line as ‘anorexic’, because male flies from this line had similar feeding to the $w^{1118}$ control while females were variably hyperphagic. Furthermore, while the $Gr28b^M$ line had low feeding, this was regardless similar to the feeding of *OR-R* male flies. Both *Gr28b* mutant lines had feeding rates that were within the distribution of the wild types tested. These results suggest that feeding varies strongly with genetic background, and that background must be carefully controlled in order to make meaningful comparisons between genotypes.

Although the paper that pioneered the FlyPad demonstrated that the apparatus is an accurate way of recording the amount of food ingested, as ingestion correlated strongly with the duration and number of sips taken by the fly (Itskov et al., 2014), we reasoned that feeding behavior and ingestion volume may be decoupled in $Gr28b^P$ flies. We therefore attempted to measure ingestion in a manner similar to (Ayres and Schneider, 2009), by allowing flies to feed on blue dye for 30 minutes, selecting flies that had visible dye in the abdomen, and homogenizing groups of these flies to estimate ingestion by recording absorbance of the blue dye in homogenates.

One replicate of this experiment did suggest that fewer male $Gr28b^P$ flies fed enough in the 30-minute period to have visible blue dye in the abdomen, and that the amount of blue dye ingested in these flies was less than the $w^{1118}$ wild type (Figure 1C, D). We note however that there are strong biases associated with these methods, as even an attempt to select feeding flies in an unbiased way leads primarily to favoring those flies that have ingested the greatest amount of blue dye, as they are the most noticeable and easiest to select. This biases the result towards flies with the maximum ingestion for each genotype. Calculation of a ‘feeding index’ by multiplying the proportion of flies feeding for a genotype by the absorbance obtained for that genotype suggested that $w^{1118}$ males had high overall feeding behavior compared to all other treatments of both sexes (Figure 1E), similar to the FlyPad data (Figure 1A). Since the $w^{1118}$ line was used as a comparison to the $Gr28b^P$ flies by (Ayres and Schneider, 2009), this may have led them to conclude that these mutants were anorexic. However, it seems more accurate to say that $w^{1118}$ males are hyperphagic.

An alternate explanation for the discrepancy between our data and that of (Ayres and Schneider, 2009) is that genotypes may have different food preferences, due to mutations in taste receptors or other factors. Indeed, (Sang et al., 2019) found that *Gr28b$^{M}$* mutant flies do not avoid saponin-laced food as wild type flies do. Therefore, apparent 'absolute' feeding rates could differ between genotypes depending on the food recipe used. (Ayres and Schneider, 2009) used a dextrose based medium different from ours, which may have been comparatively unappealing to the *Gr28b$^{P}$* flies.

In conclusion, comparison of two *Gr28b* insertion mutants indicates that mutation of this gene does not cause anorexia in flies, and that the feeding patterns observed by (Ayres and Schneider, 2009) were due to genetic background or environmental effects.

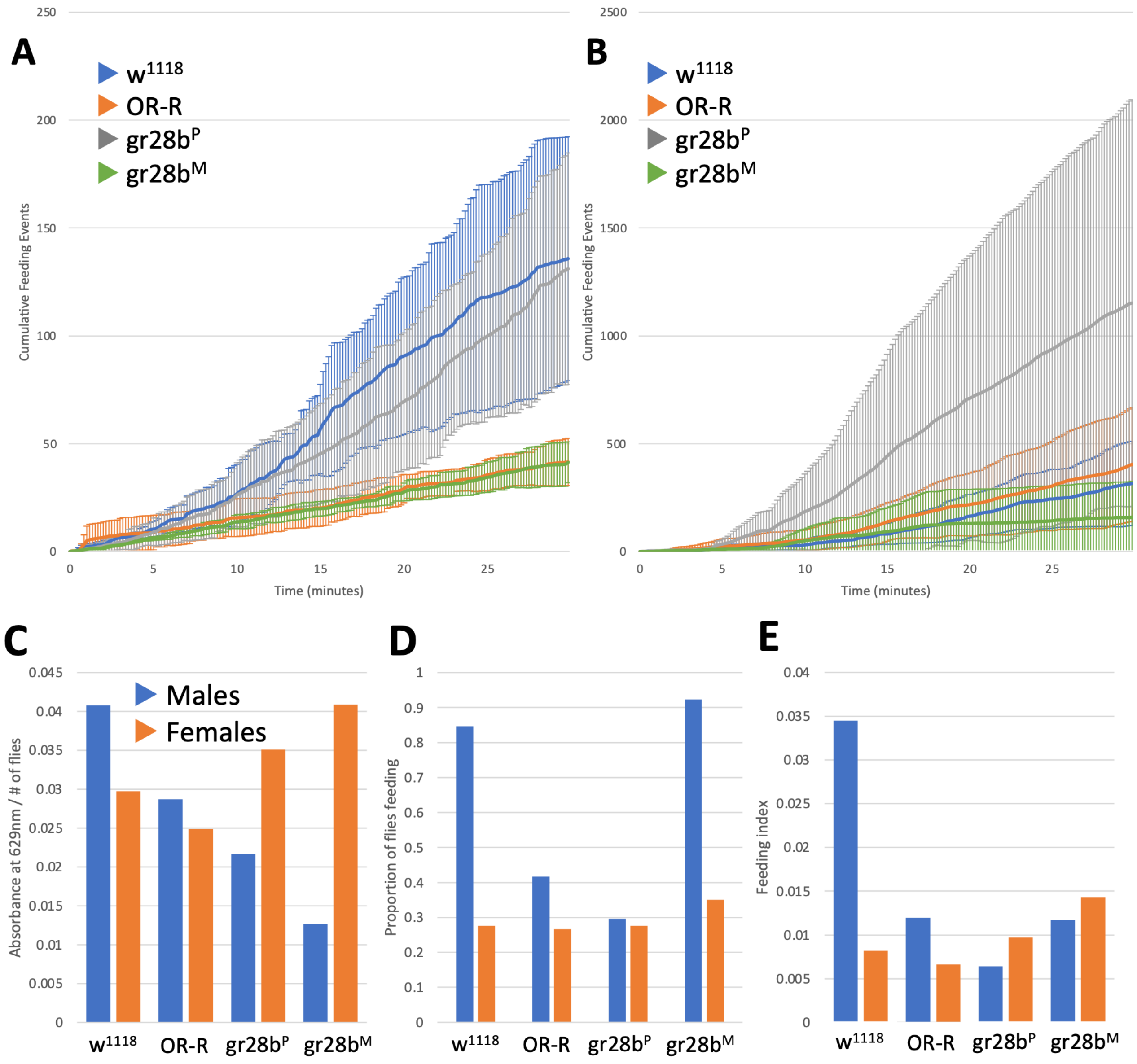

**Figure 1.** Feeding characteristics of wild-type and *Gr28b* mutant flies as shown by **(A, B)** the FlyPad apparatus over 30 minutes ($n^{exp}$=3) and **(C-E)** by ingestion of food containing FD&C Blue No. 1 dye over 30 minutes ($n^{exp}$=1). FlyPad data shows cumulative feeding events in **A)** male and **B)** female flies over 30 minutes of feeding. The trendlines show the average number of times a fly of that genotype fed over the measured period. Shaded areas show standard deviation from the mean for each genotype over time. **A)** Male *w$^{1118}$* and *Gr28b$^{P}$* flies had similarly high feeding rates, whereas male *OR-R* and *Gr28b$^{M}$* flies had low feeding. **B)** Female *Gr28b$^{P}$* flies had very high feeding compared to other genotypes, while *Gr28b$^{M}$* flies had low feeding. Feeding of *OR-R* and *w$^{1118}$* females was similar and intermediate. **C)** Ingestion of blue dye suggests that male *Gr28b$^{P}$*

and $Gr28b^M$ flies have low feeding compared to $w^{1118}$ flies as proposed by (Ayres and Schneider, 2009). When the proportion of flies feeding in each genotype is taken into account **(D)** and used to calculate a feeding index **(E)**, feeding of *Gr28b* flies does not differ strongly from wild-type. Both FlyPad and feeding index data suggest that male $w^{1118}$ flies are hyperphagic compared to other genotypes, including the *OR-R* wild type.

Gr28b *mutant flies do not have altered tolerance and resistance to* S. typhimurium *and* L. monocytogenes

(Ayres and Schneider, 2009) found that 'anorexic' $Gr28b^P$ mutants or diet restricted wild type flies (24h starvation or lifelong 0.5x diet) had increased survival but similar bacterial loads when infected with *S. typhimurium*, indicating increased tolerance to this infection. These flies also had reduced survival and increased bacterial loads of *L. monocytogenes,* indicating reduced resistance to this infection. Finally, survival to *E. faecalis* was unchanged (see Ayres and Schneider, 2009 – Figure 2). The authors note that reduced resistance to *L. monocytogenes* is likely due to reduced melanization seen in $Gr28b^P$ and diet restricted flies, as their previous results showed that melanization is important in defense against this pathogen (Ayres and Schneider, 2008).

Other authors have typically found that starvation or reduced insulin signaling enhances survival to bacterial infections (see e.g. (Lee et al., 2017) *P. aeruginosa,* (Libert et al., 2008) *P. aeruginosa* and *E. faecalis*, (McCormack et al., 2016) *P. luminescens* and *E. coli*)). Reduced survival of anorexic flies in response to *L. monocytogenes* infection may be consistent with results showing that this infection is highly metabolically demanding and causes rapid depletion of energy stores, resulting in death (Chambers et al., 2012). However, the results of (Sang et al., 2019) and our own results (see above) indicate that *Gr28b* mutants are not anorexic, and therefore this is an insufficient explanation for differences in survival to bacterial infections.

To test whether loss of *Gr28b* has consistent effects on resistance or tolerance, we employed both available *Gr28b* mutants and subjected them to survivals with various bacteria.

Males of both *Gr28b* mutant lines had similar survival to *L. monocytogenes* as the *OR-R* wild type. However, *OR-R* had significantly lower survival compared to other wild-types tested ($P<0.05$), intermediate between the other wild types ($w^{iso}$, $w^{1118}$) and immune-deficient lines ($Rel^{E20}$, $Dif^1$). (Ayres and Schneider, 2009) backcrossed the $Gr28b^P$ mutant (generated in a $w^{1118}$ background) into their isogenic $w^{1118}$ background for four generations and found that the $Gr28b^P$ mutation conferred a susceptibility to *L. monocytogenes* compared to the $w^{1118}$ wild type line (males only). Our data shows a similar trend for both $Gr28b^P$ and $Gr28b^M$ mutant males compared to the $w^{1118}$ background ($P<0.05$) (Figure 3A). Females do not show a similar reduction in survival (Figure 3B). Overall, these results are consistent with (Ayres and Schneider, 2009) in that males of both *Gr28b* mutant lines had reduced survival to *L. monocytogenes* compared to the $w^{1118}$ wild type. This pattern did not hold true in female flies, where only the $Gr28b^M$ line had reduced survival compared to wild types. Differing phenotypes between the two mutant lines indicates that genetic background may be a significant factor in susceptibility. Further experiments using RNAi or re-expression of *Gr28b* in these mutant lines would be required to confidently assign this phenotype to mutation of *Gr28b.*

We were unable to show increased survival of $Gr28b^P$ males compared to the $w^{1118}$ background to *S. typhimurium* as found by (Ayres and Schneider, 2009). Instead, the $Gr28b^M$ line was somewhat susceptible compared to wild-types ($P<0.0005$) (Figure 3C, D). Female $Gr28b^P$ flies had a slight but significant increase in survival to *S. typhimurium* over the $w^{1118}$ wild-type ($P<0.0005$), whereas $Gr28b^M$ female survival was similar to wild-type (Figure 3D). Increased susceptibility of male $Gr28b^M$ mutants and increased resistance of female $Gr28b^P$ flies was somewhat visible with LB injection alone (Figure 3E, F), suggesting that these effects are not specific to bacterial infection. We did not measure bacterial loads and are unable to comment on whether the *Gr28b* mutation increases tolerance to *S. typhimurium* as claimed by (Ayres and Schneider, 2009).

Survivals to *E. faecalis* showed a significant reduction in survival of $Gr28b^P$ ($P<0.0005$) but not $Gr28b^M$ compared to the $w^{1118}$ wild type (Figure 3G). This is in contrast to (Ayres and

Schneider, 2009), who found that survival to this pathogen was unaltered in the *Gr28b^P* mutant. The strong difference in survival between the two mutant lines suggest that this is due to a genetic background effect.

Finally, we note that our survival methods differed from (Ayres and Schneider, 2009), who injected bacterial suspensions into the ventrolateral abdomen, whereas we employed septic injury (*E. faecalis, L. monocytogenes*) and thoracic injection (*S. typhimurium*). In our hands, attempts to inject in the abdomen resulted in excessive damage to soft tissues and high levels of mortality unrelated to infection.

In conclusion, there are some differences in survival between the *Gr28b^P* mutant and its appropriate control, but these are largely not replicated in an independent *Gr28b^M* mutant. We conclude that contrary to the findings of (Ayres and Schneider, 2009), mutation of *Gr28b* alone does not affect strongly affect survival to infection with *E. faecalis, L. monocytogenes* or *S. typhimurium.*

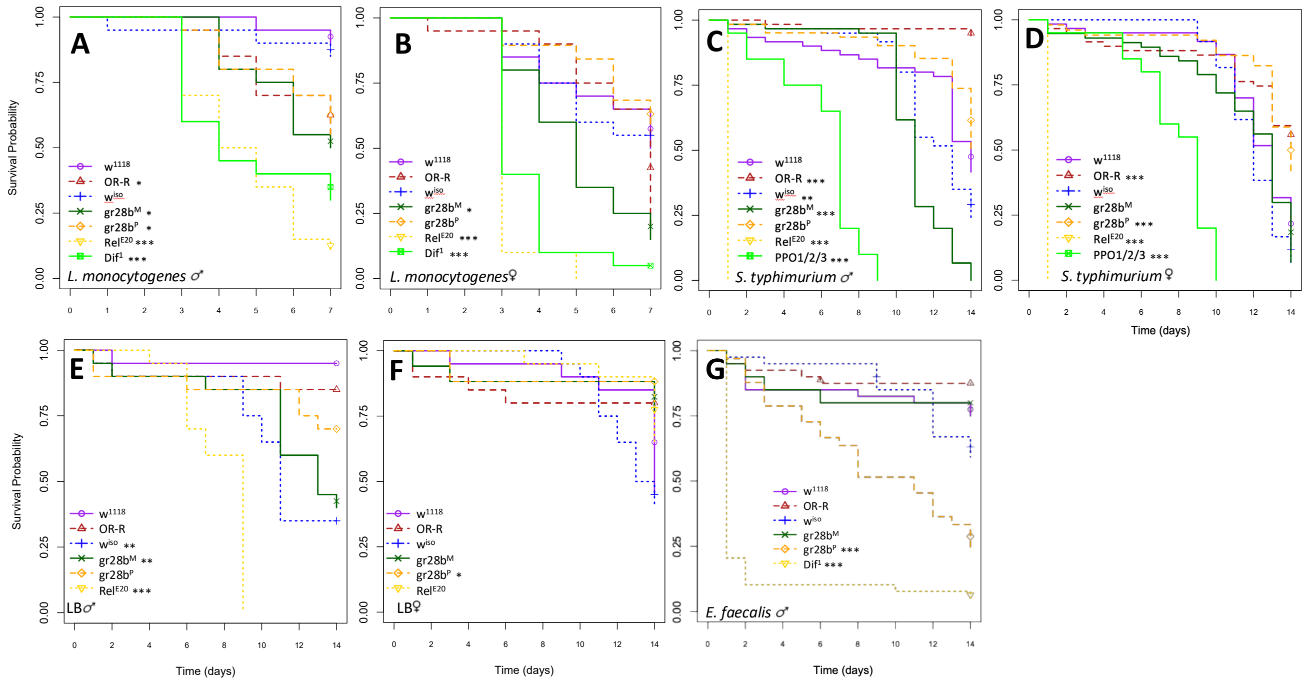

**Figure 3.** Survivals of two *Gr28b* mutant lines: *Gr28b^P*, used by (Ayres and Schneider, 2009) and *Gr28b^M*, used by (Sang et al., 2019)) to a range of bacterial infections. **A, B:** *L. monocytogenes* OD=0.5, septic infection, 25°C; **C, D:** *S. typhimurium* OD=0.01 in LB, injection, 29°C; **E, F:** Injection of LB alone 29°C; and **G:** *E. faecalis* OD=0.5, septic infection, 29°C. Asterisks beside legends indicate significant differences compared to the *w^1118* wild type as determined by a Cox Proportional Hazards test. * P<0.05, ** P<0.01, *** P<0.001. At least 20 flies per genotype were scored in 2-3 independent experiments per survival. Data for male and female flies were combined only where sexes did not differ significantly (indicated on figure).

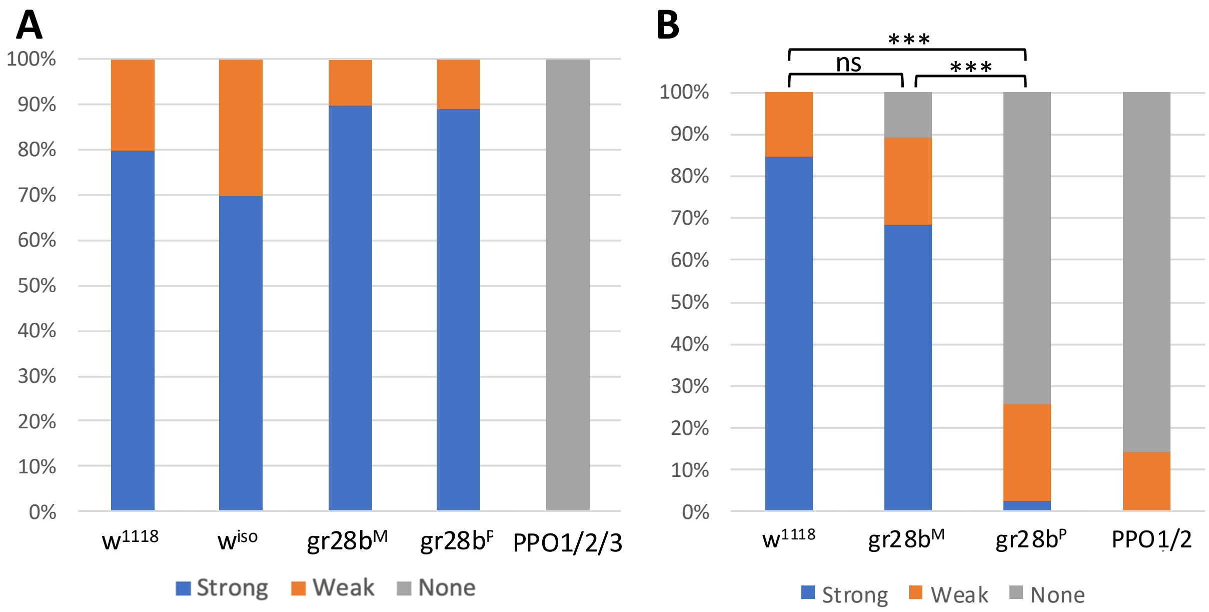

**Figure 4.** Melanization in *Gr28b* mutant adults **(A)** and larvae **(B)**. Adults of both *Gr28b* lines have wild-type melanization in response to septic injury with *M. luteus* (16h post-infection, $n^{exp}$=1, 20 flies per genotype). Only *Gr28b^P^* larvae had reduced melanization similar to *PPO1/2* mutants, with most larvae completely failing to melanize in response to injury ($n^{exp}$=2, ~20 larvae per genotype per experiment). Chi-squared test, ns, not significant; ***, P < 0.001

*Altered AMP expression and reduced melanization in* Gr28b *mutants is due to genetic background*

(Ayres and Schneider, 2009) found that ‘anorexic’ *Gr28b$^P$* mutants or diet-restricted wild type flies (24h or lifelong 0.5x diet) had reduced melanization compared to normally fed wild-type flies. The same group also found that both melanization and feeding rate were reduced in *eiger* RNAi flies, and based on this similarity to *Gr28b$^P$* flies inferred that reduced food uptake may impair melanization (Mabery and Schneider, 2010). In contrast to this, (McCormack et al., 2016) found that *chico* mutants with reduced insulin pathway activity and poor nutrition instead had increased melanization and PO activity, higher AMP expression and phagocytic activity. Furthermore, contrary to (Ayres and Schneider, 2009), our data and that of (Sang et al., 2019) indicate that disruption of *Gr28b* does not cause anorexia (Figure 1). Thus, even if reduced insulin pathway activity can reduce melanization as suggested by (Ayres and Schneider, 2009), this is unlikely to be the cause of reduced melanization in *Gr28b$^P$* flies.

(Ayres and Schneider, 2009) further found that *Gr28b$^P$* mutants had altered AMP expression at 6h post-infection in response to *L. monocytogenes* infection. Diet-restricted flies were not assessed for this parameter. They state that, “AMPs that are usually described to be coordinately regulated were regulated independently in anorexic *Gr28b* flies infected with *L. monocytogenes*: these flies had reduced expression of *Drs* and *Drc*, but higher levels of *Att* and similar *Dpt* to wild type flies.” Note that the *Attacin* primers used in both (Ayres and Schneider, 2009) and in our experiments co-amplify *AttA* and *AttB*.

To test whether melanization and AMP expression are affected by loss of *Gr28b,* we assayed melanization and expression of these AMPs in two *Gr28b* insertion mutant lines.

We found that although cuticle melanization was wild-type at the adult stage in both the *Gr28b$^P$* and *Gr28b$^M$* lines (Figure 4A), blackening was strongly impaired in *Gr28b$^P$* larvae, similar to *PPO1/2* mutants lacking hemolymph phenoloxidases (Figure 4B). In contrast, *Gr28b$^M$* larvae had wild-type melanization. Melanization, and particularly degree of hemolymph blackening apparent upon larval injury, is known to be important in survival to certain infections (Ayres et al., 2008; Binggeli et al., 2014; Dudzic et al., 2019). These results indicate that although melanization is reduced in *Gr28b$^P$* line, this is likely due to background effects and not disruption of *Gr28b* itself, as the *Gr28b$^M$* line has normal melanization. Survivals performed with these lines support this conclusion, as the survival phenotypes found by (Ayres and Schneider, 2009) were largely only replicable with the *Gr28b$^P$* line, and not the *Gr28b$^M$* alternate line (Figure 3). The survival phenotypes of the *Gr28b$^P$* line are consistent with (Binggeli et al., 2014) who show that loss of PO activity strongly reduces survival to *E. faecalis* but has little effect on survival to *L. monocytogenes* or *S. typhimurium*.

In contrast to (Ayres and Schneider, 2009), we found that in response to *L. monocytogenes* infection male flies of both *Gr28b* lines had similar or higher expression of both *Drs* and *Drc* compared to wild types, and similar expression of *AttA/B* and *Dpt* to wild types (Figure 5, upper panel). Expression of AMPs in unchallenged *Gr28b* flies was also within the distribution of wild-type expression (Figure 5, lower panel). (Ayres and Schneider, 2009) concluded that anorexia in the *Gr28b$^P$* mutants was responsible for variation in AMP expression; however, our data indicate that *Gr28b$^P$* flies are not anorexic (Figure 1A, B). Furthermore, this interpretation does not agree with independent data on the effect of starvation on AMP expression: (Libert et al., 2008) found that AMP expression was not increased by starvation, whereas (McCormack et al., 2016) found that overall AMP expression increased in insulin pathway mutants.

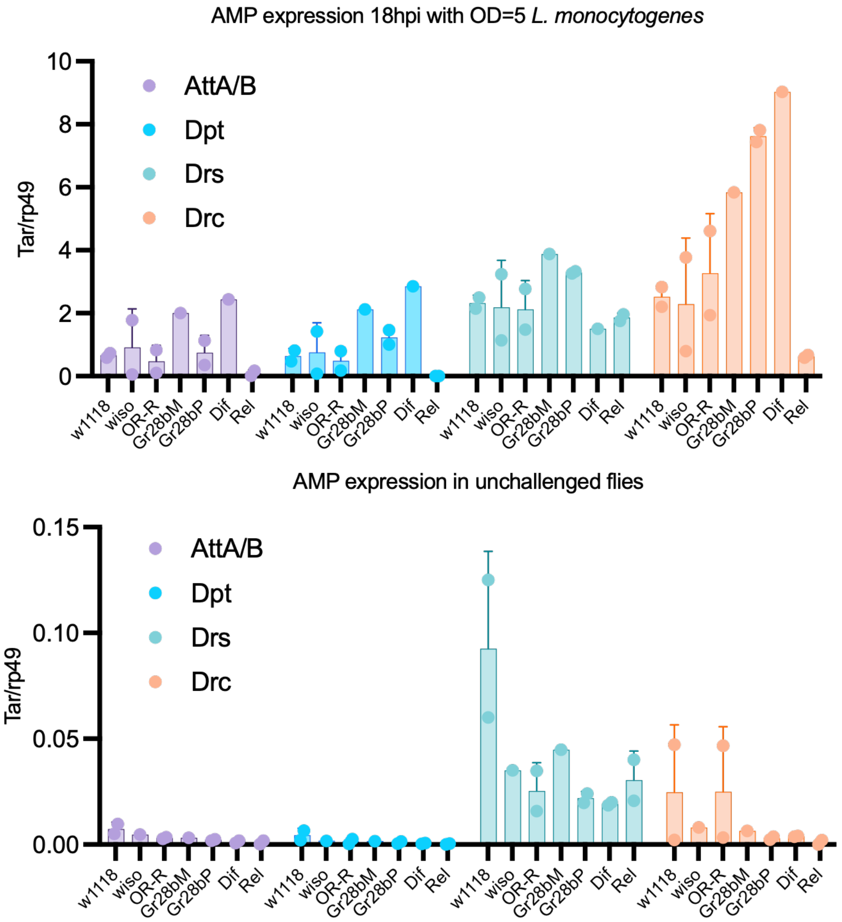

**Figure 5.** Expression of the AMPs *AttA/B, Dpt, Drs* and *Drc* in *Gr28b$^M$* and *Gr28b$^P$* flies 18h following infection with $OD_{600}$=5 *L. monocytogenes* (upper panel) and in unchallenged conditions (lower panel). Data points indicate independent experiments, error bars show standard deviation.

**Conclusion**

Initial results using a P-element insertion mutant of *Gr28b* suggested that loss of this gene causes a range of phenotypes including anorexia, increased tolerance to *S. typhimurium* and reduced resistance to *L. monocytogenes,* altered expression of several AMPs, and reduced melanization. Experiments with an independent insertion mutant and a range of wild-types instead shows that the wild-type used for comparison (*w$^{1118}$*) is hyperphagic, while feeding of *Gr28b* mutants is within the normal distribution. Apparent survival, AMP and melanization effects are also due to genetic background rather than mutation of the *Gr28b* gene, as an independent *Gr28b* mutant does not display these phenotypes. We therefore conclude that *Gr28b* does not have a significant role in fly immunity.
